# Selective inhibition reveals the regulatory function of DYRK2 in protein synthesis and calcium entry

**DOI:** 10.1101/2021.02.12.430909

**Authors:** Tiantian Wei, Jue Wang, Ruqi Liang, Wendong Chen, An He, Yifei Du, Wenjing Zhou, Zhiying Zhang, Mingzhe Ma, Jin Lu, Xing Guo, Xiaowei Chen, Ruijun Tian, Junyu Xiao, Xiaoguang Lei

## Abstract

The dual-specificity tyrosine phosphorylation-regulated kinase DYRK2 has emerged as a key regulator of cellular processes such as proteasome-mediated protein degradation. To gain further insights into its function, we took a chemical biology approach and developed C17, a potent small-molecule DYRK2 inhibitor, through multiple rounds of structure-based optimization guided by a number of co-crystallized structures. C17 displayed an effect on DYRK2 at a single-digit nanomolar IC_50_ and showed outstanding selectivity for the human kinome containing 467 other human kinases. Using C17 as a chemical probe, we further performed quantitative phosphoproteomic assays and identified several novel DYRK2 targets, including eukaryotic translation initiation factor 4E-binding protein 1 (4E-BP1) and stromal interaction molecule 1 (STIM1). DYRK2 phosphorylated 4E-BP1 at multiple sites, and the combined treatment of C17 with AKT and MEK inhibitors showed synergistic 4E-BP1 phosphorylation suppression. The phosphorylation of STIM1 by DYRK2 substantially increased the interaction of STIM1 with the ORAI1 channel, and C17 impeded the store-operated calcium entry process. Collectively, these studies further expand our understanding of DYRK2 and provide a valuable tool to further pinpoint its biological function.

---

Dual-specificity tyrosine phosphorylation-regulated kinases (DYRKs) belong to the CMGC group of kinases together with other important human kinases, such as cyclin-dependent kinases (CDKs) and mitogen-activated protein kinases (MAPKs)^1–3^. DYRKs uniquely phosphorylate tyrosine residues within their activation loops *in cis* during biosynthesis, although mature proteins display exclusive serine/threonine kinase activities^4^. There are five DYRKs in humans: DYRK1A, DYRK1B, DYRK2, DYRK3, and DYRK4. DYRK1A has been extensively studied due to its potential function in the pathogenesis of Down syndrome and neurodegenerative disorders^5, 6^. DYRK3 has been shown to function as a central “dissolvase” to regulate the formation of membraneless organelles^7, 8^. On the other hand, DYRK2 is a key regulator of 26S proteasome activity^9^.

The 26S proteasome degrades the majority of proteins in human cells and plays a central role in many cellular processes, including the regulation of gene expression and cell division^10, 11^. Recent discoveries have revealed that the 26S proteasome is subjected to intricate regulation by reversible phosphorylation^9, 12, 13^. DYRK2 phosphorylates the Rpt3 subunit in the regulatory particle of the proteasome at Thr25, leading to the upregulation of proteasome activity^9^. DYRK2 is overexpressed in several tumours, including triple-negative breast cancer and multiple myeloma, which are known to rely heavily on proteasome activity for progression, and perturbation of DYRK2 activity impedes cancer cell proliferation and inhibits tumour growth^14, 15^.

Our knowledge of the physiological functions of DYRK2 remains in its infancy, and it is likely that DYRK2 has cellular targets in addition to Rpt3. In fact, substrates of many kinases, especially Ser/Thr kinases, remain insufficiently identified. A major obstacle to discovering physiologically relevant substrates of a kinase is the lack of highly specific chemical probes that allow precise modulation of kinase function. Some DYRK2 inhibitors have been reported; however, these compounds also inhibit other kinases, mostly other DYRK family members, to various degrees^16, 17^. We have recently identified LDN192960 as a selective DYRK2 inhibitor and showed that LDN192960 can alleviate multiple myeloma and triple-negative breast cancer progression by inhibiting DYRK2-mediated proteasome phosphorylation^15^. To obtain even more potent and selective DYRK2 inhibitors, we applied a structure-guided approach to further engineer chemical compounds based on the LDN192960 scaffold. One of the best compounds we generated, compound C17 (C17), displays an effect on DYRK2 at a single-digit nanomolar IC_50_ with moderate to excellent selectivity against kinases that are closely related to DYRK2. Using this potent DYRK2 inhibitor as a tool, we treated U266 cells with C17 and performed quantitative phosphoproteomic analyses, which led to the identification of several novel DYRK2 targets, including eukaryotic translation initiation factor 4E-binding protein 1 (4E-BP1) and stromal interaction molecule 1 (STIM1). These results demonstrate that DYRK2 plays important regulatory roles in multiple cellular processes, including protein translation and store-operated calcium entry, and indicate that C17 can serve as a valuable probe for the study of DYRK2 function.

## Results

### Structure-based optimization of DYRK2 inhibitors

LDN192960 was identified as a DYRK2 inhibitor^15, 18, 19^. It occupies the ATP-binding pocket of DYRK2 and mediates extensive hydrophobic and hydrogen bond interactions^15^. Nevertheless, LDN192960 also inhibits other DYRK2-related kinases, especially Haspin and DYRK3^15^. To generate DYRK2 inhibitors with better selectivity, we synthesized a series of new compounds based on the same acridine core structure (Table 1). The amine side chain was first changed to a protected amine (compounds 1-3), a cyano group (compound 4), or a cyclic amine (compounds 5-6) (Extended Data Fig. 1a, Table 1). Among these candidates, compound 6 exhibited the strongest inhibitory effect towards DYRK2, with an *in vitro* IC_50_ of 17 nM, while LDN192960 showed an IC_50_ of 53 nM when the same protocol was used (Table 1). Treating HEK293T cells transiently expressing DYRK2 with increasing concentrations of compound 6 efficiently inhibited Rpt3-Thr25 phosphorylation, with the maximal effect observed at an inhibitor concentration of less than 3 μM (Extended Data Fig. 1b). Importantly, compound 6 also displays good selectivity towards DYRK2 compared to other kinases, including DRYK1A, DRYK1B, DYRK3, Haspin, and MARK3 (IC_50_ values of 889, 697, 121305, 45, and 100 nM, respectively; Table 1). Therefore, compound 6 was chosen as the lead compound for further chemical modification.

**Table 1.**
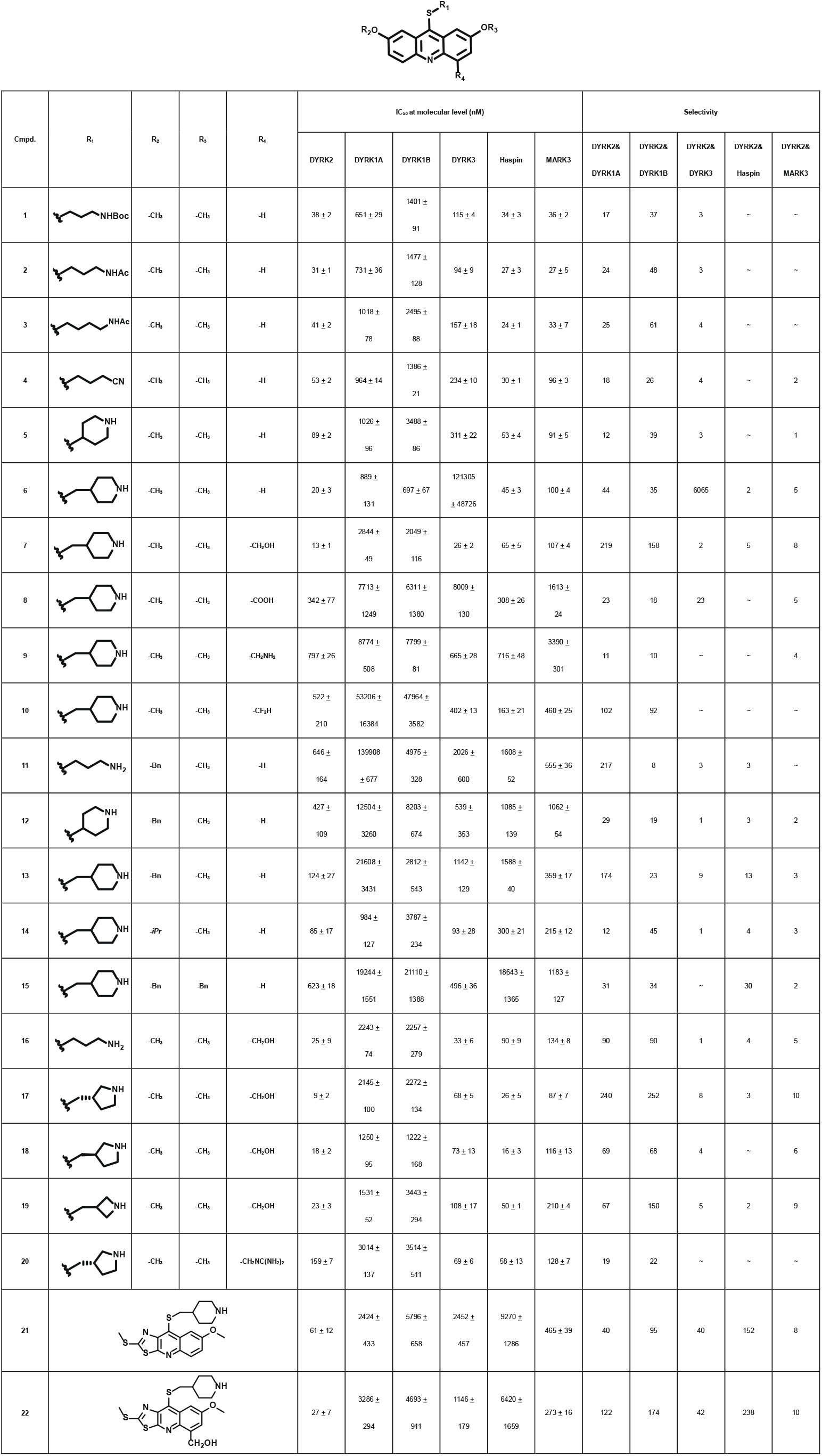
The Inhibitory activity and selectivity of acridine analogues of DYRK2.

We subsequently crystallized DYRK2 in complex with compound 6 and determined the structure at a resolution of 2.2 Å (Fig. 1a, Extended Data Fig. 1c). Not surprisingly, compound 6 binds the ATP-binding site of DYRK2 in a manner similar to LDN192960. A water molecule is located deep inside the binding pocket acting as a bridge in the interactions between LDN192960 and the protein. The newly added amino side chain displays clear densities and adopts an extended conformation. An in-depth analysis of the crystal structure revealed several additional sites for chemical expansion that may further strengthen its interaction with DYRK2 (Extended Data Fig. 2a). First, a hydrophilic group can be introduced into the acridine core to functionally replace the aforementioned water molecule and maintain a constant contact with DYRK2. Second, a bulky functional group can be used to replace the methoxy groups to mediate additional interactions with DYRK2. Finally, the amine side chain can be further altered to stabilize its conformation (Extended Data Fig. 2a). To this end, we synthesized 9 new compounds (compounds 7-15) and evaluated their inhibitory effects on DYRK2 and related kinases (Fig. 2c, Extended Data Fig. 2c). We also determined the co-crystallized structures of several of these compounds with DYRK2 to visualize their detailed interactions (Fig. 1, Extended Data Fig. 3). Compound 7, introducing a hydroxymethyl group to the acridine core, inhibits DRYK2 efficiently as compound 6 while displaying better selectivity against other DRYK family members (Table 1). The co-crystallized structure shows that the hydroxymethyl group directly contacts the main chain amide group of Ile367 and indirectly coordinates Glu266 and Phe369 via a water molecule (Fig. 1d). Compared to compound 7, compounds 8-10, which contain a carboxyl, aminomethyl, and fluoromethyl group, respectively, instead of a hydroxymethyl group, display reduced inhibition towards DYRK2. Compounds 11-15, which were designed to replace the methoxy group with a bulkier side chain, also showed significantly decreased activity and selectivity and therefore were not further pursued (Table 1).

**Fig. 1.**
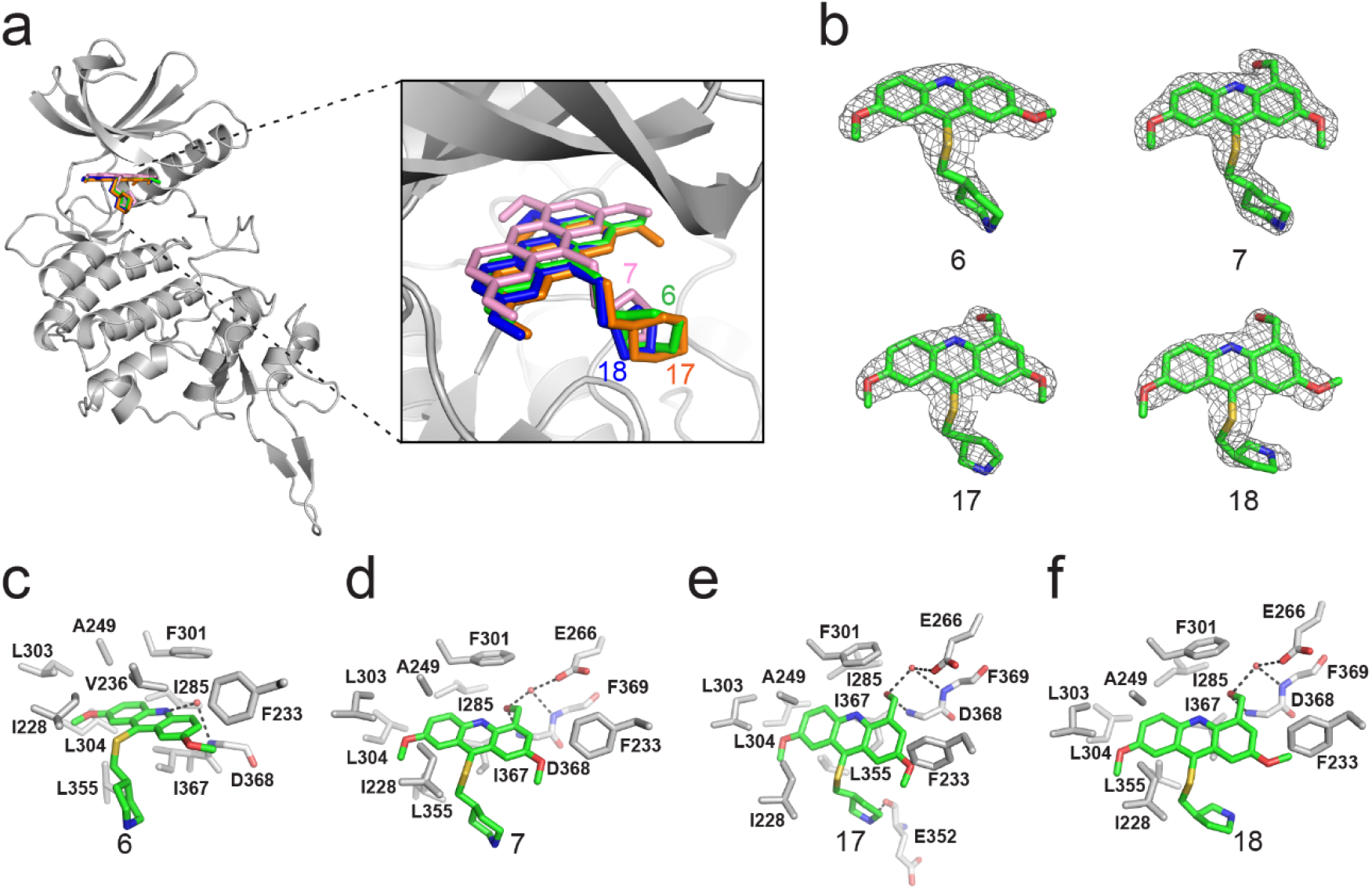
Crystal structures of DYRK2 bound to novel agonists. **a**, Overall structure of DYRK2 (grey) bound to 6 (green), 7 (orange), C17 (blue) and 18 (pink). **b**, Composite omit maps are contoured at 1.5σ and shown as grey meshes to reveal the presence of compounds 6, 7, 17 and 18 in the respective crystal structures. **c-f**, Close-up view of the DYRK2 binding pocket with compounds 6, 7, 17 and 18. Hydrogen bonds are shown as dashed lines. Water molecules are indicated with red spheres.

Further chemical modification was carried out based on compound 7. By changing the 6-membered ring to a straight chain or smaller rings, we synthesized compounds 16-19 (Extended Data Fig. 2e, Table 1). Among these compounds, C17 with a (S)-3-methylpyrrolidine side chain exhibited the best potency and selectivity among all the analogues. Interestingly, we noticed that compound 18 containing a (R)-3-methylpyrrolidine side chain was not as good as C17, indicating that the chirality of the 3-methylpyrrolidine motif plays an important role in both potency and selectivity. Further modification of compound 17 (compound 20) aimed at promoting further hydrogen bond interactions with DYRK2 failed to improve the inhibitory effect. We also wondered whether acridine was the best aromatic core structure and synthesized two new compounds (compounds 21 and 22) by changing one side of the benzene group to a sulfur-containing thiazole structure, which we thought might facilitate hydrophobic interactions with DYRK2 within the ATP-binding pocket; however, they did not have as effective an inhibitory effect as compound 17 (Table 1). The details of the chemical syntheses of all the intermediates and tested compounds are provided in the Supporting Information.

### C17 is a potent and selective DYRK2 inhibitor

We set to comprehensively characterize the inhibitory function of compound 17 (Fig. 2a), referred to as C17 hereafter. *In vitro,* C17 displays an effect on DYRK2 at a single-digit nanomolar IC_50_ value (9 nM) (Fig. 2b, Extended Data Fig. 3a). To further evaluate the selectivity of C17, we performed kinome profiling analyses. Among the 468 human kinases tested, C17 targeted only DYRK2, Haspin, and MARK3 at a concentration of 500 nM (Fig. 2c). Nonetheless, the *in vitro* IC_50_ values of C17 for Haspin and MARK3 (26 nM and 87 nM, respectively) were 3-10-fold higher than that for DYRK2 (Fig. 2b, Extended Data Fig. 3b-f). Similarly, C17 also inhibited DYRK3 to a lesser extent (IC_50_ of 68 nM). In contrast, LDN192960 inhibited DYRK3 and Haspin to a greater extent than it inhibited DYRK2^15, 18^. Importantly, C17 also efficiently suppressed DYRK2 activity in the cell and abolished Rpt3-Thr25 phosphorylation at an inhibitor concentration of less than 1 μM (Fig. 2d). Taken together, these data demonstrate that C17 is a highly potent and selective DYRK2 inhibitor both *in vitro* and *in vivo.*

**Fig. 2.**
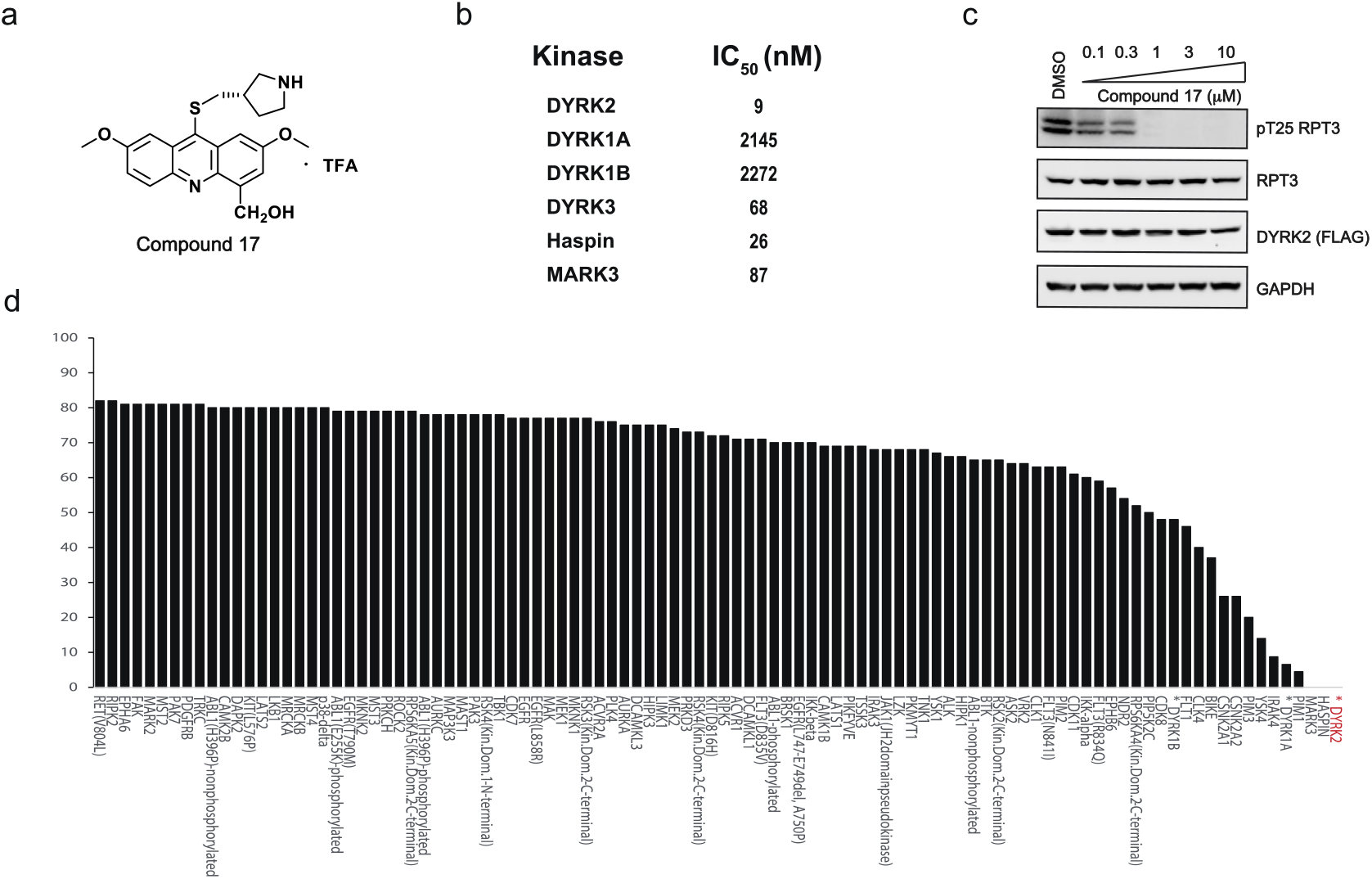
C17 is a potent and selective inhibitor of DYRK2. **a**, Chemical structure of C17. **b**, IC_50_ values of C17 against DYRK1A, DYRKIB, DYRK3, Haspin and MARK3. **c**, Kinome profiling of C17 at 500 nM was carried out using 468 human kinases (https://www.discoverx.com/). **d**, C17 inhibits Rpt3-Thr25 phosphorylation. HEK293T cells stably expressing FLAG-DYRK2 were treated with the indicated concentrations of C17 for 1 h. The cells were lysed, and immunoblotting was carried out with the indicated antibodies.

### DYRK2 substrate profiling by quantitative phosphoproteomic analyses

Quantitative phosphoproteomic approaches have significantly expanded the scope of phosphorylation analysis, enabling the quantification of changes in thousands of phosphorylation sites simultaneously^20^. To obtain a comprehensive list of potential DYRK2 targets, we treated the myeloma U266 cells with C17 and carried out quantitative phosphoproteomic analyses^21, 22^. We prepared lysates of U266 cells treated with C17 or the DMSO control and trypsinized them. Phosphorylated peptides were then enriched using Ti^4+^-immobilized metal ion affinity chromatography (IMAC) tips and analysed by LC-MS/MS (Fig. 3a). A total of 15,755 phosphosites were identified, among which 12,818 (81%) were serine and 2,798 (18%) were threonine. A total of 10,647 (68%) phosphosites were Class I (localization probability >0.75), 2,557 (16%) were Class II (0.5< localization probability ≤0.75), and 2,401 (16%) were Class III ( 0.25< localization probability ≤0.5) (Fig. 3b). To the best of our knowledge, this is the largest phosphoproteomic dataset prepared to date for DYRK2 substrate profiling. Extended Data Fig. 5 shows that a good Pearson correlation coefficient of 0.9 was obtained for the phosphosite intensities among the treatment and control samples. The coefficient of variance of the intensities of the majority of the phosphosites was lower than 20% (Extended Data Fig. 6), demonstrating the high quantification precision of our label-free phosphoproteomic analysis. Remarkably, C17 treatment led to significant downregulation of 337 phosphosites. These sites may all be potential DYRK2 targets (Fig. 3c). Interestingly, another 445 phosphosites were upregulated. DYRK2 likely inhibits some downstream kinases or activates some phosphatases, and therefore, suppressing its activity reversed these effects. In any event, these data demonstrate that DYRK2 is involved in a network of phosphorylation events and can directly or indirectly regulate the phosphorylation status of many proteins. The top pathways with which DYRK2 may participate were revealed by a global analysis of the significantly up- and downregulated phosphoproteins (Fig. 3e).

**Fig. 3.**
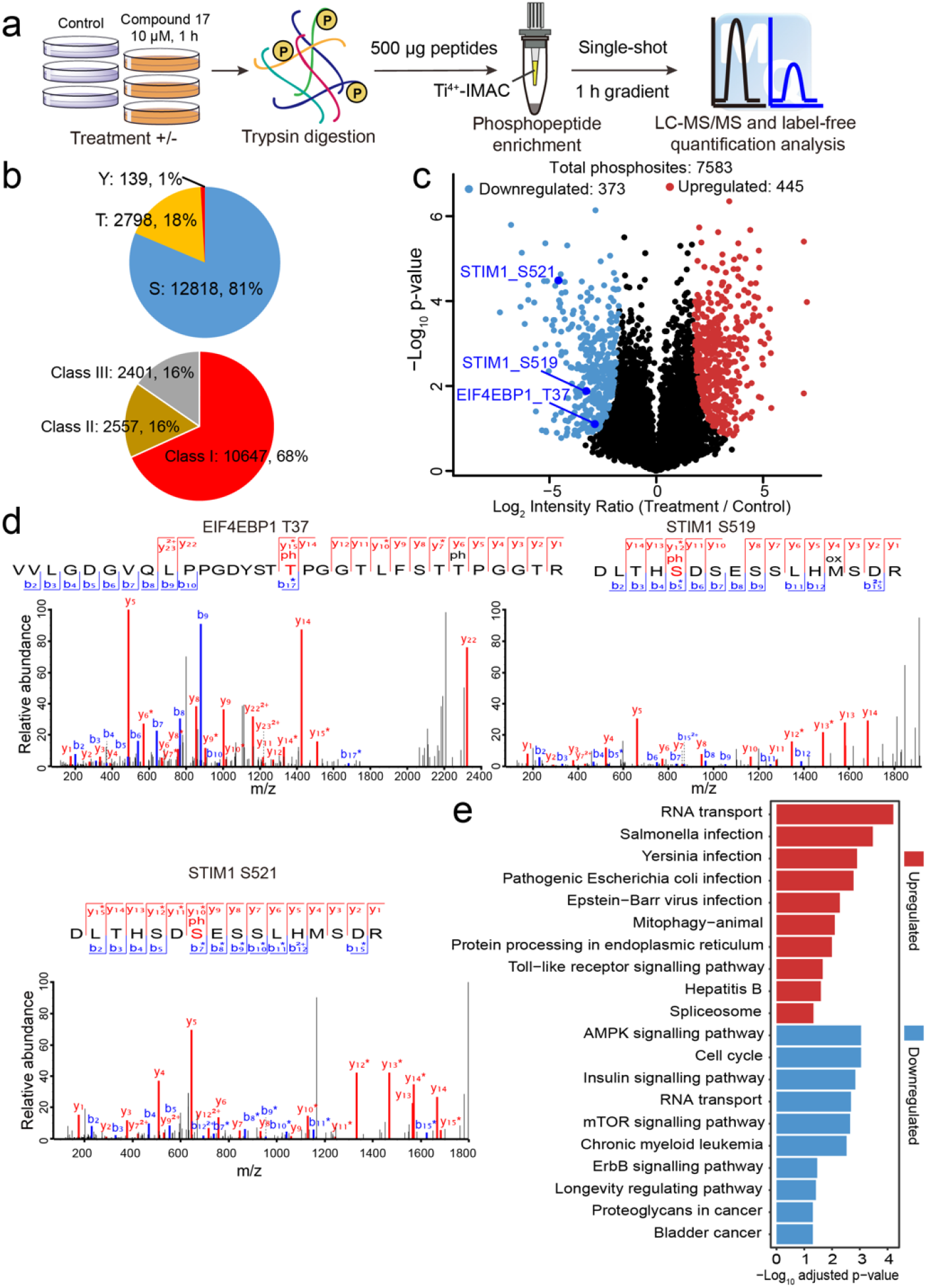
Quantitative phosphoproteomic analysis of U266 cells treated with C17. **a**, Workflow of the phosphoproteomic approach. Triplicate samples treated with/without 10 μM C17 for 1 h were separately lysed and digested, and the phosphorylated peptides were enriched by the Ti^4+^-IMAC tip and analysed by LC-MS/MS. **b**, Distribution of the assigned amino acid residues and their localization probabilities (Class I >0.75, Class II >0.5 and ≤0.75, Class III >0.25 and ≤0.5) for all identified phosphorylation sites. **c**, Volcano plot (FDR < 0.05 and S0 = 2) shows the significantly up- and downregulated phosphosites after C17 treatment. **d**, MS/MS spectra of the phosphosites of two potential DYRK2 substrates, pT37 of 4E-BP1 and pS519 and pS521 of STIM1. **e**, Global canonical pathway analysis of the significantly up- and downregulated phosphoproteins. −Log_10_ adjusted *p*-values associated with a pathway are presented.

### 4E-BP1 is a direct cellular target of DYRK2

We set out to determine whether some of the 337 downregulated phosphosites are genuine DYRK2 targets. We first focused on eukaryotic translation initiation factor 4E-binding protein 1 (4E-BP1), which is a master regulator of protein synthesis. Phosphorylation of 4E-BP1 by mTORC1 leads to its dissociation from eukaryotic initiation factor 4E (eIF4E), allowing mRNA translation^23, 24^. The MS/MS spectrum of pThr37 of 4E-BP1 is shown in Fig. 3d with clear fragmentation pattern for confident peptide identification and site localization. And Extended Data Fig. 7 shows that the intensity for pThr37 of 4E-BP1 is confidently quantified with significant changes. C17 treatment significantly reduced the phosphorylation of Thr37 of both endogenous and overexpressed 4E-BP1 in HEK293T cells (Fig. 4a), and the findings are consistent with the mass spectrometry analyses performed using U266 cells. To examine whether DYRK2 can directly phosphate 4E-BP1, we performed an *in vitro* kinase assay using purified DYRK2 and 4E-BP1 proteins. As shown in Fig. 4b, DYRK2 efficiently phosphorylated 4E-BP1 at multiple sites, including Thr37, whereas the kinase-deficient DYRK2 mutant (D275N) had no activity. These findings are also consistent with a previous study showing that Ser65 and Ser101 in 4E-BP1 can be phosphorylated by DYRK2^25^. C17 suppressed DYRK2-mediated 4E-BP1 phosphorylation in a dose-dependent manner (Fig. 4c). These results demonstrate that 4E-BP1 is a genuine substrate of DYRK2 and that C17 can inhibit DYRK2 activity both *in vivo* and *in vitro*.

**Fig. 4.**
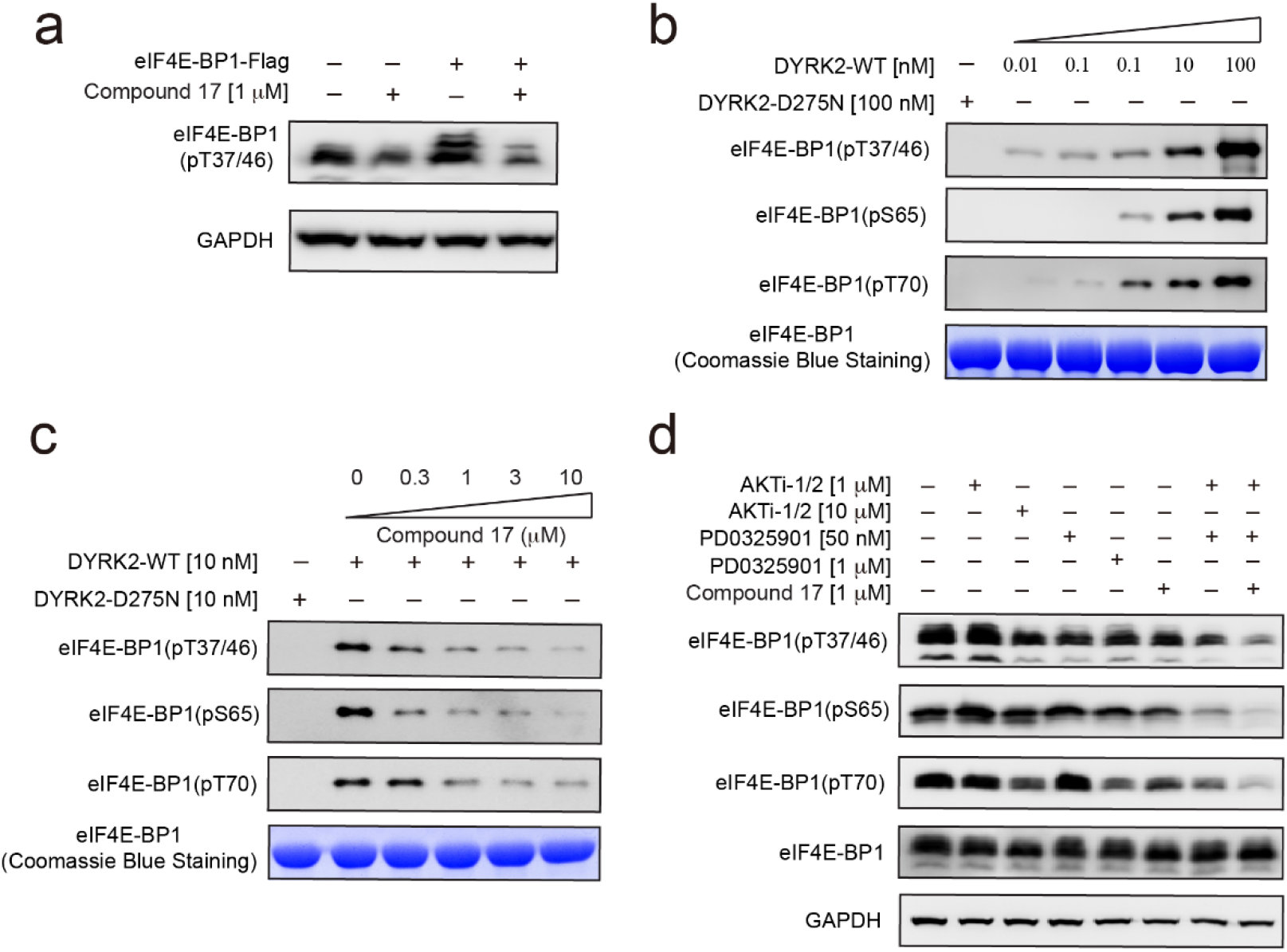
4E-BP1 is a substrate of DYRK2. **a**, C17 treatment reduced the phosphorylation of endogenous and overexpressed 4E-BP1 in HEK293T cells. The phosphorylation status of 4E-BP1 was analysed by immunoblotting of cell lysates using an anti-4E-BP1 pThr37/46 antibody. **b**, DYRK2 directly phosphorylated 4E-BP1 at multiple sites. **c**, C17 inhibited DYRK2-mediated 4E-BP1 phosphorylation in a concentration-dependent manner. **d**, C17 displayed a synergistic effect with AKT and MEK inhibitors to suppress 4E-BP1 phosphorylation. HTC116 cells were treated with the indicated concentrations of PD032590, AKTi, and C17 alone or in combination for 6 h. Cell lysates were immunoblotted with the indicated antibodies.

C17 decreased but did not totally abolish the phosphorylation of 4E-BP1 (Fig. 4a). Indeed, 4E-BP1 can be targeted by multiple kinases^26^. A previous study showed that the combined inhibition of AKT and MEK inhibited 4E-BP1 phosphorylation and tumour growth^27^. To assess whether C17 can elicit a greater synergistic effect with these kinase inhibitors, we treated HTC116 cells with AKTi (an AKT1/AKT2 inhibitor), PD0325901 (a MEK inhibitor), and C17, either alone or in combination, and examined the phosphorylation status of endogenous 4E-BP1 (Fig. 4d). Indeed, while AKTi and PD0325901 alone decreased 4E-BP1 phosphorylation to a certain degree, the combined treatment resulted in more marked suppression. The presence of C17 further potentiated the inhibitory effect (Fig. 4d). These results confirm that 4E-BP1 is a direct cellular target of DYRK2 and suggest the potential use of DYRK2 inhibitors in combination with other kinase inhibitors for cancer therapy.

### DYRK2 phosphorylates STIM1 to modulates SOCE

In addition to 4E-BP1, another potential target of DYRK2 is STIM1 (stromal interaction molecule 1), as indicated by the phosphorylation levels of both Ser519 and Ser521 in STIM1 being significantly reduced upon DYRK2 inhibition in our mass spectrometry analyses (Fig. 3c). The MS/MS spectra for pSer519 and pSer521 of STIM1 are shown in Fig. 3d with clear fragmentation pattern for confident peptide identification and their intensities are confidently quantified with significant changes (Extended Data Fig. 7). STIM1 is a single-pass transmembrane protein residing in the endoplasmic reticulum (ER) and plays a key role in the store-operated calcium entry (SOCE) process ^28^. The luminal domain of STIM1 senses calcium depletion in the ER and induces protein oligomerization. Oligomerized STIM1 then travels to the ER-plasma membrane contact site and activates the ORAI1 calcium channel. The cytosolic region of STIM1 contains multiple phosphorylation sites, and it has been shown that the function of STIM1 is regulated by several kinases, including ERK1/2^29, 30^.

To verify that DYRK2 can directly phosphorylate STIM1, we performed an *in vitro* kinase experiment (Fig. 5a). Purified wild-type (WT) DYRK2, but not the kinase-dead (KD) mutant D275N, induced a dose-dependent mobility change of the cytosolic region of STIM1 (STIM1^235-END^) in SDS-PAGE gels, suggesting that STIM1 is effectively phosphorylated by DYRK2. To assess the functional outcome of STIM1 phosphorylation by DYRK2, we examined the interaction between STIM1 and ORAI1 in HEK293A cells. The expression of WT DYRK2 significantly increased the interaction between STIM1 and ORAI1 (Fig. 5b). In contrast, the expression of DYRK2-KD exerted no such effect. Treating cells with C17 effectively abolished the DYRK2-dependent STIM1-ORAI1 interaction. C17 also decreased the interaction between STIM1 and ORAI1 in the absence of exogenously expressed DYRK2 (Fig. 5c), suggesting that C17 can inhibit endogenous DYRK2 activity, in accordance with our mass spectrometry results.

**Fig. 5.**
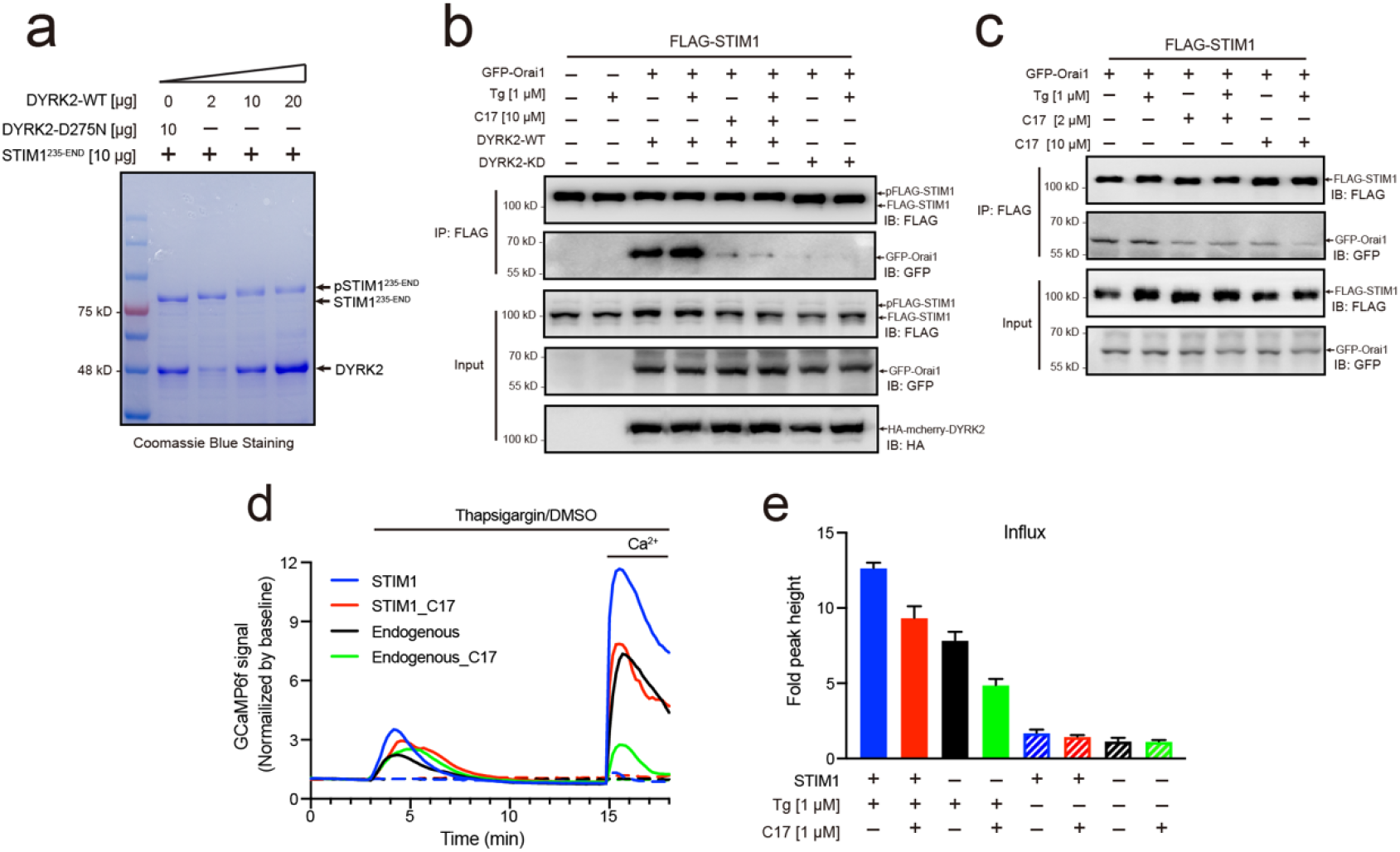
Phosphorylation of STIM1 by DYRK2 modulates SOCE. **a**, DYRK2 directly phosphorylated STIM1. GST-STIM1^235-END^ was incubated with wild-type or kinase-deficient DYRK2 in the presence of Mn-ATP for 30 minutes. Phosphorylation of GST-STIM1^235-END^ was indicated by the mobility change of STIM1 in SDS-PAGE gel. **b**, DYRK2 promoted the interaction between STIM1 and OraiI1. HEK293A cells were co-transfected with FLAG-STIM1, GFP-Orai1, and wild-type or kinase-deficient DYRK2 for 36 h and then treated with 10 μM C17 for 1 h. Then, the cells were incubated in Ca^2+^-free medium with or without 1 μM Tg for 10 minutes. STIM1 was immunoprecipitated with FLAG agarose, and the associated proteins were analysed with the indicated antibodies. **c**, HEK293A cells were co-transfected with FLAG-STIM1 and GFP-Orai1 for 36 h and then treated with the indicated amounts of C17 for 1 h. Flag IP was performed similarly as in **b**. **d**, C17 inhibited SOCE in HEK293A cells. HEK293A cells were transfected with GCAMP6f or GCAMP6f plus STIM1 for 24 h and then treated with 1 μM C17 for 1 h. The medium of the cell culture was switched to Ca^2+^-free medium before thapsigargin treatment. Thapsigargin (1 μM, solid lines) or DMSO (dashed lines) was added to the cells, and 2 mM Ca^2+^ was added 12 minutes later. The red and green lines correspond to C17-treated cells. Blue and black lines represent untreated cells. GCAMP6f fluorescence was monitored by a Zeiss LSM 700 laser scanning confocal microscope. **e**, Quantification of **d**. The following number of cells were monitored: STIM1, 44 cells on 3 coverslips (blue solid line); STIM1 + C17 (1 μM), 59 cells on 3 coverslips (red solid line); endogenous, 47 cells on 3 coverslips (black solid line); endogenous + C17 (1 μM), 42 cells on 3 coverslips (green solid line). STIM1(-Tg), 43 cells on 3 coverslips (blue dashed line). STIM1 + C17 (1 μM) (-Tg), 43 cells on 3 coverslips (red dashed line); endogenous (-Tg), 43 cells on 3 coverslips (black dashed line); and endogenous + C17 (1 μM) (-Tg), 43 cells on 3 coverslips (green dashed line). Error bars represent the means ± s.e.m.

We did not observe a significant difference in the interaction between STIM1 and ORAI1 in the presence or absence of thapsigargin, which induces calcium depletion in the ER. This result is likely related to the overexpression of both STIM1 and ORAI1 in our assay to evaluate their interaction since we could not procure antibodies that recognize the endogenous proteins. To further examine the physiological relevance of our results, we performed SOCE analyses with HEK293A cells expressing GCaMP6f, a genetically encoded calcium sensor^31^. Treating cells grown in calcium-free medium containing thapsigargin, but not DMSO, resulted in a transient increase in GCaMP6f fluorescence due to the release of calcium from the ER to the cytosol (Fig. 5d). The subsequent addition of calcium to the cell culture medium resulted in a marked increase in GCaMP6f signalling, indicating calcium entry into the cells, which was further augmented by STIM1 overexpression (Fig. 5d, 5e). Pretreating cells with C17 for 1 h substantially reduced SOCE into cells with either endogenous or overexpressed STIM1 levels (Fig. 5d, 5e). Together, these results demonstrated that DRYK2 can directly phosphorylate STIM1, which leads to enhanced STIM1-ORAI1 interaction and SOCE.

## Discussion

Here, we used a structure-based approach to design, synthesize and evaluate a series of new analogues based on the acridine core structure and eventually identified C17 as a potent and selective DYRK2 inhibitor. We showed that C17 displays an effect on DYRK2 at a single-digit nanomolar IC_50_ and inhibits DYRK2 more potently than closely related kinases such as DYRK3, Haspin and MARK3. The crystal structure of DYRK2 bound to C17 revealed critical interactions that explain its high selectivity, including a hydrogen bond between the (S)-3-methylpyrrolidine ring and Glu352 in DYRK2 (Fig. 1e).

Acridine derivatives have traditionally been used as antibacterial, antiparasitic and anticancer agents since these compounds usually show strong DNA intercalating effects^36^. Considering the potential toxicity of C17 due to its possible DNA-binding capacity, we also assessed the DNA-binding effect of C17 (Extended Data Fig. 11). Isothermal titration calorimetry revealed that C17 (Kd=36 μM) binds to DNA with significantly lower affinity than LDN192960 binds to DNA (Kd=200 nM), possibly because of the introduction of hydroxymethyl group on the acridine core, which are not present in LDN192960.

C17 provided us with a unique tool to interrogate the physiological functions of DYRK2. We treated U266 cells with C17 and performed quantitative phosphoproteomic analyses. We found that the cellular phosphorylation pattern is significantly altered by C17, suggesting that DYRK2 likely has multiple cellular targets and is involved in a network of biological processes. We then identified several leading phosphosites that are downregulated and demonstrated that 4E-BP1 and STIM1 are bona fide substrates of DYRK2. We showed that DYRK2 efficiently phosphorylated 4E-BP1 at multiple sites, including Thr37, and combined treatment of C17 with AKT and MEK inhibitors resulted in marked suppression of 4E-BP1 phosphorylation. Therefore, DYRK2 likely functions synergistically with other kinases to regulate the protein synthesis process. For the first time, we also discovered that STIM1 can be efficiently phosphorylated by DYRK2, and phosphorylation of STIM1 by DYRK2 substantially increased the interaction between STIM1 and ORAI1. Treating cells with C17 suppressed SOCE, validating the important role of DYRK2 in regulating calcium entry into cells. Taken together, these results reveal that DYRK2 is implicated in multiple cellular processes, and it will be interesting to further explore the complicated phosphorylation network that involves DYRK2.

Recently, Mehnert M. et al. developed a multilayered proteomic workflow and determined how different pathological-related DYRK2 mutations altered protein conformation, substrates modification and biological function^37^. Therefore, DYRK2 is also of great significance for regulating many unknown protein-protein interactions. Accordingly, the selective DYRK2 inhibitor C17 that we have developed may also serve as a useful tool in dissecting complex protein-protein interactions regulated by DYRK2.

## Methods

### IC_50_ determination

IC_50_ determination was carried out using the ADP-Glo^TM^ kinase assay system (Promega, Madison, WI). Active DYRK1A, DYRK1B, DYRK2, DYRK3, Haspin and MARK3 were purified as reported previously. C17 IC_50_ measurements were carried out against the kinases with final concentrations between 0.01 nM to 100 μM *in vitro* (C17 was added to the kinase reaction prior to ATP master mix). The values were expressed as a percentage of the DMSO control. DYRK isoforms (1 ng/μL diluted in 50 mM Tris-HCl pH7.5, 2 mM DTT) are assayed against Woodtide (KKISGRLSPIMTEQ) in a final volume of 5 μL containing 50 mM Tris pH 7.5, 150 μM substrate peptide, 5 mM MgCl_2_ and 10 −50 μM ATP (10μM for DYRK2 and DYRK3, 25 μM for DYRK1A and 50 μM for DYRK1B) and incubated for 60 min at room temperature. Haspin (0.2 ng/μL diluted in 50 mM Tris-HCl pH7.5, 2 mM DTT) was assayed against a substrate peptide H3(1-21) (ARTKQTARKSTGGKAPRKQLA) in a final volume of 5 μL containing 50 mM Tris pH 7.5, 200 μM substrate peptide, 5 mM MgCl_2_ and 200 μM ATP and incubated for 120 min at room temperature. MARK3 (1 ng/μL diluted in 50 mM Tris-HCl pH7.5, 2 mM DTT) was assayed against Cdc25C peptide (KKKVSRSGLYRSPSMPENLNRPR) in a final volume of 5 μL 50 mM Tris pH 7.5, 200 μM substrate peptide, 10 mM MgCl_2_ and 5 μM ATP and incubated for 120 min at room temperature. After incubation, the ADP-Glo^TM^ kinase assay system was used to determine kinase activity following the manufacturer’s protocol. IC_50_ curves were developed as % of DMSO control and IC_50_ values were calculated using GraphPad Prism software.

### Quantitative phosphoproteomic analysis

Triplicate U266 cells treated with/without compound 17 were lysed by the lysis buffer containing 1% (v/v) Triton X-100, 7M Urea, 50 mM Tris-HCl, pH 8.5, 1 mM pervanadate, protease inhibitor mixture (Sigma-Aldrich, USA), and phosphatase inhibitor mixtures (Roche, Switzerland). The cell lysates were firstly digested with trypsin (Promega, USA) by the in-solution digestion method^22^. After desalting, the Ti^4+^-IMAC tip was used to purify the phosphopeptides. The phosphopeptides were desalted by the C18 StageTip prior to the LC MS/MS analysis^22^. An Easy-nLC 1200 system coupled with the Q-Exactive HF-X mass spectrometer (Thermo Fisher Scientific, USA) was used to analyze the phosphopeptide samples with 1 h LC gradient. The raw files were searched against Human fasta database (downloaded from Uniprot on March 27, 2018) by MaxQuant (version 1.5.5.1). The oxidation (M), deamidation (NQ), and Phospho (STY) were selected as the variable modifications for the phosphopeptide identification, while the carbamidomethyl was set as the fixed modification. Label-free quantification (LFQ) and match between runs were set for the triplicate analysis data. The MaxQuant searching file “Phospho (STY)Sites.txt” was loaded into the Perseus software (version 1.5.5.3) to make volcano plots using student’s t-test and cutoff of “FDR< 0.05 and S0=2”. The pathway analysis was performed using the Kyoto Encyclopedia of Genes and Genomes (KEGG) database with cutoff of adjusted p-value < 0.05.

### KINOMEscan^®^ kinase profiling

The KINOMEscan^®^ kinase profiling assay was carried out at The Largest Kinase Assay Panel in the world for Protein Kinase Profiling (https://www.discoverx.com). Compound 17 kinase selectivity was determined against a panel of 468 protein kinases. Results are presented as a percentage of kinase activity in DMSO control reactions. Protein kinases were assayed *in vitro* with 500 nM final concentration of compound 17 and the results are presented as an average of triplicate reactions ± SD or in the form of comparative histograms.

### Statistics and data presentation

Most experiments were repeated 3 times with multiple technical replicates to be eligible for the indicated statistical analyses, and representative image has been shown. All results are presented as mean ± SD unless otherwise mentioned. Data were analysed using Graphpad Prism statistical package.

## Data availability

The structural coordinates of DYRK2 in complex with compounds 5, 6, 7, 8, 10, 13, 14, 17, 18, 19, and 20 have been deposited in the Protein Data Bank with accession codes 7DH3, 7DG4, 7DH9, 7DHV, 7DHC, 7DHK, 7DHO, 7DJO, 7DL6, 7DHH, and 7DHN, respectively. All the raw mass spectrometry data have been deposited in the public proteomics repository MassIVE and are accessible at ftp://massive.ucsd.edu/MSV000086745/.

## Acknowledgements

We thank the National Center for Protein Sciences at Peking University for assistance with crystal screening, and Shanghai Synchrotron Radiation Facility and KEK Photon Factory for assistance with X-ray data collection. This work is supported by National Key Research & Development Plan (2017YFA0505200 to X.L. and J.X. and 2020YFE0202200 to R. T.), the National Natural Science Foundation of China (91853202, 21625201,21961142010, 21661140001, and 21521003 to X.L., 31822014 to J.X., 31700088 to W. C., and 91953118 to R.T.).

## Author contributions

T.W., J.W., R.L., W.C., R.T., J.X. and X.L. designed the experiments. T.W. conducted the biological and biochemical experiments with the help of Y.D., W.Z., Z.Z. and performed the crystal structure study under the guidance of J.X. J.W. and M.M. synthesized small molecules under the guidance of X.L. R.L. conducted the chemical library screens and IC_50_ determination under the guidance of X.L. W.C. and A.H. performed the quantitative phosphoproteomic analysis under the guidance of R.T. X.G. and X.C. provided critical reagents and suggestions. T.W., J.W., R.L., W.C., R.T., J.X. and X.L. wrote the manuscript, with inputs from all the authors.

## Extended Data Figures and Tables

**Extended Data Figures 1-11**

**Extended Data Fig.1.**
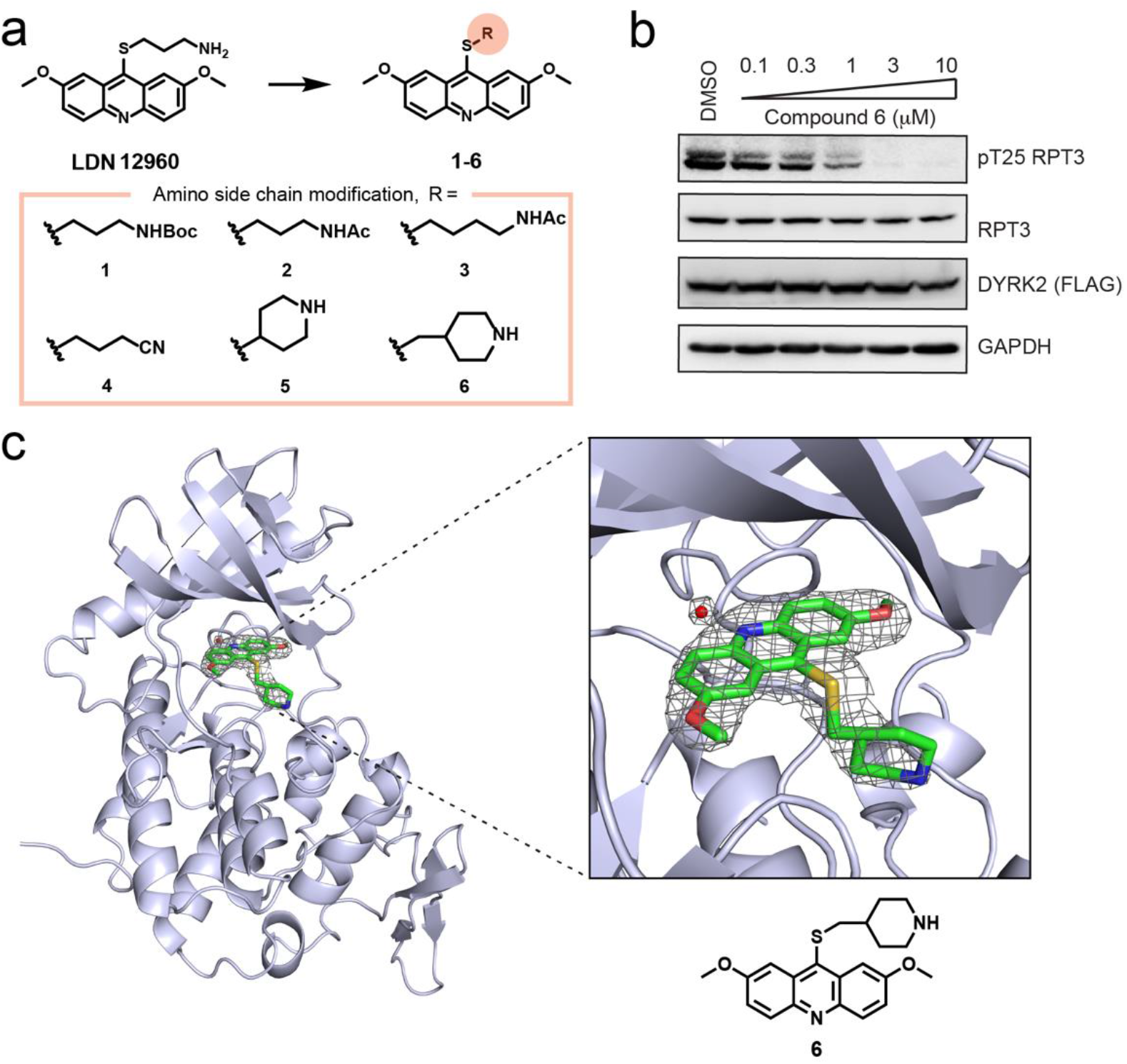
Chemical compounds derived from LDN192960. **a**, Structure of amino side chain change analogues 1-6 based on LDN192960. **b**, HEK293T cells stably expressing FLAG-DYRK2 were treated with the indicated concentrations of compound 6 in 1h. Cells were lysed and immunoblotting was carried out with the indicated antibodies. **c**, Structure of DYRK2 in complex with compound 6. DYRK2 is shown as ribbons and colored in blue white. The 2Fo-Fc difference electron density map (1.5σ which reveals the presence of 6 and water is shown as a gray mesh. The 6 and water are omitted to calculate the map).

**Extended Data Fig.2.**
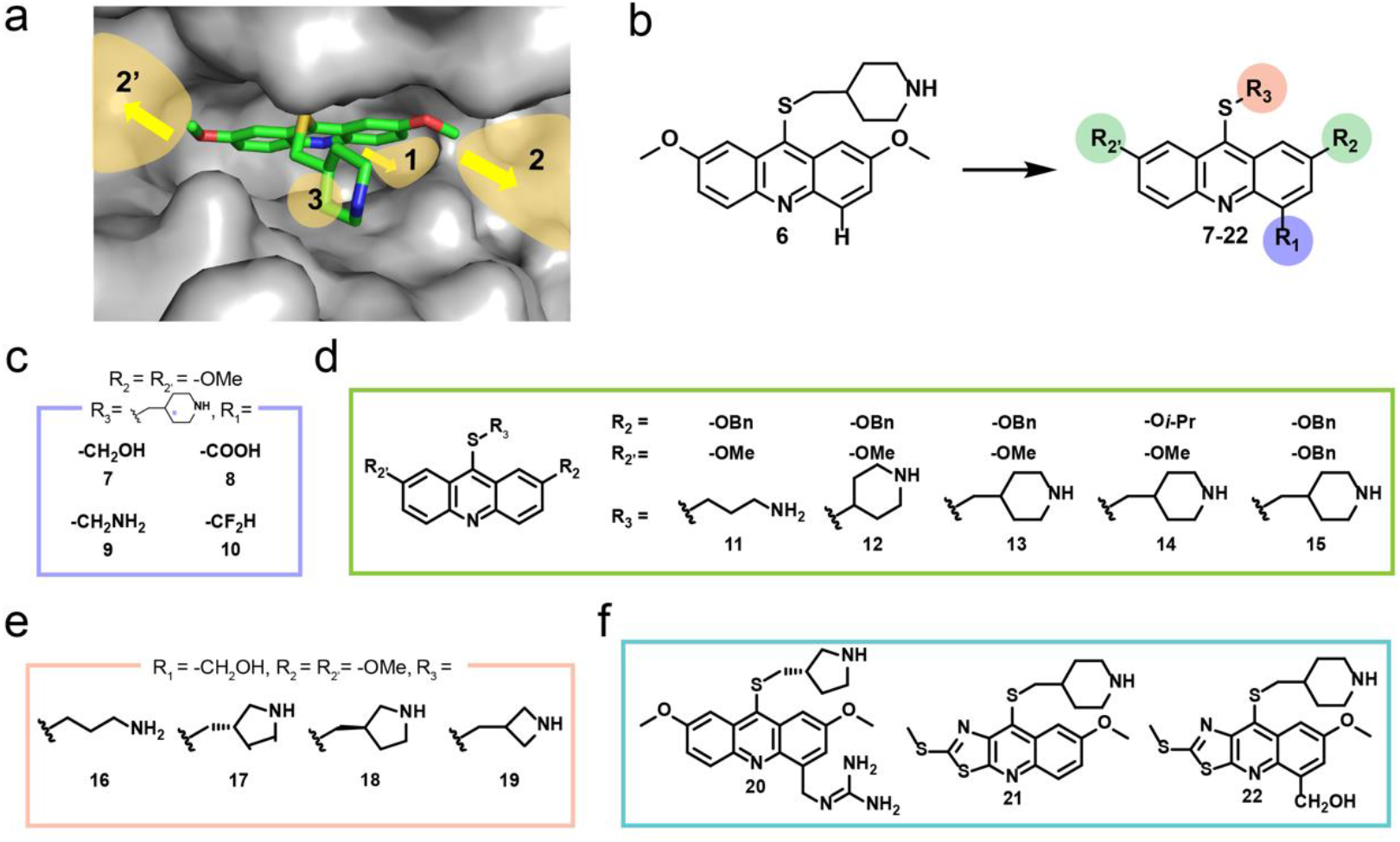
Structure-guided engineering of DYRK2 inhibitors based on compound 6. **a**, The possible sites for further expansion based on the co-crystal structure of 6 and DYRK2. **b**, Overview of modification of compound 6. **c**, Modifications for inner space 1. **d**, Modifications for cavity around ATP-binding pocket. **e**, Modifications of amine side chain based on compound 7. **f**, Modifications based on compound 17.

**Extended Data Fig.3.**
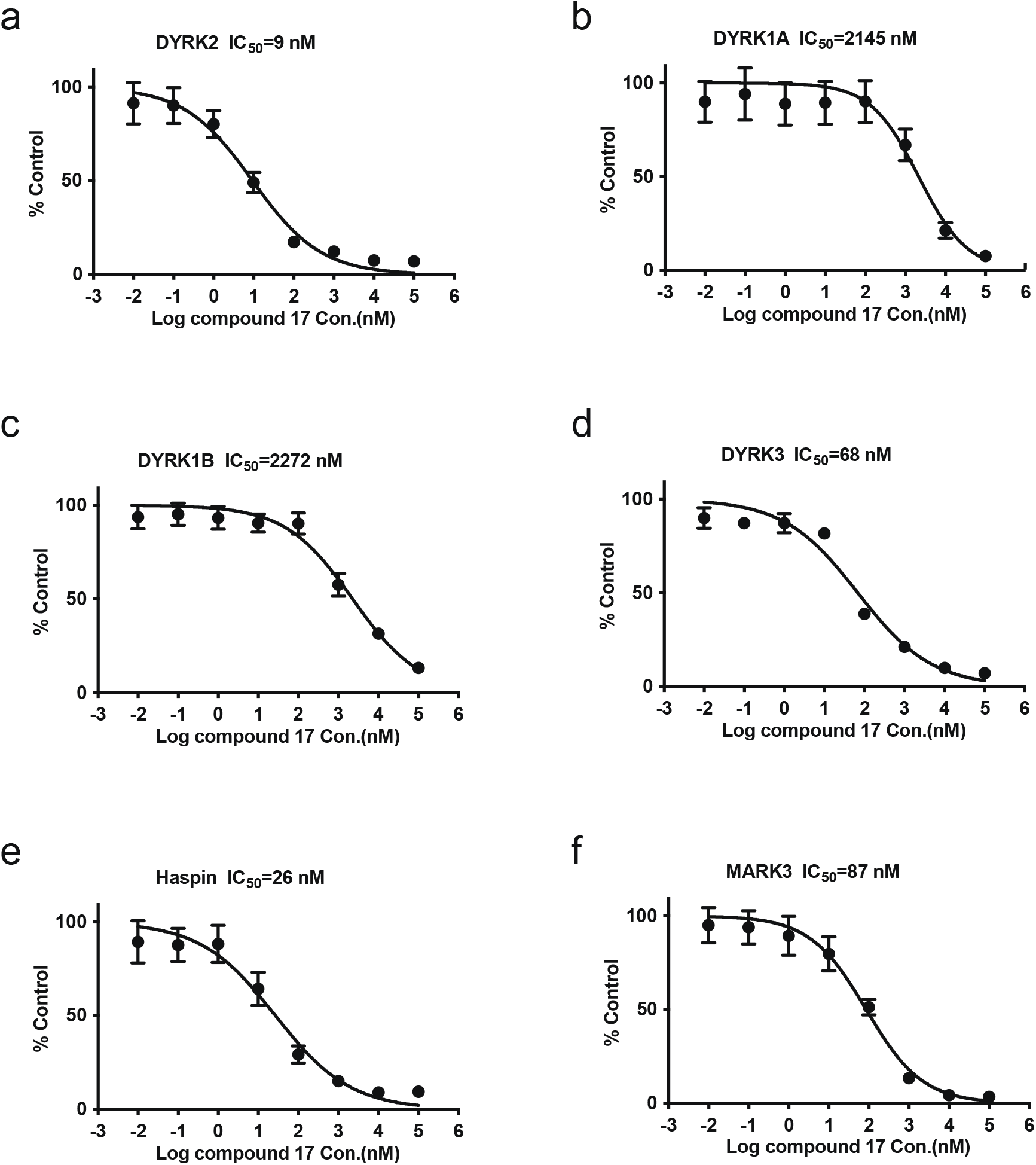
IC_50_ of C17 on DYRK2 and its main off targets. **a**, IC_50_ of C17 on DYRK2. DYRK2 was assayed using the indicated concentrations of C17. The IC_50_ graph was plotted using GraphPad Prism software. **b**, IC_50_ of C17 on DYRK1A. DYRK1A was assayed as in **a**. **c**, IC_50_ of C17 on DYRK1B. DYRK1B was assayed as in **a**. **d**, IC_50_ of C17 on DYRK3. DYRK3 was assayed as in **a**. **e**, IC_50_ of C17 on Haspin. Haspin was assayed as in **a**. **f**, IC_50_ of C17 on MARK3. MARK3 was assayed as in **a**.

**Extended Data Fig.4.**
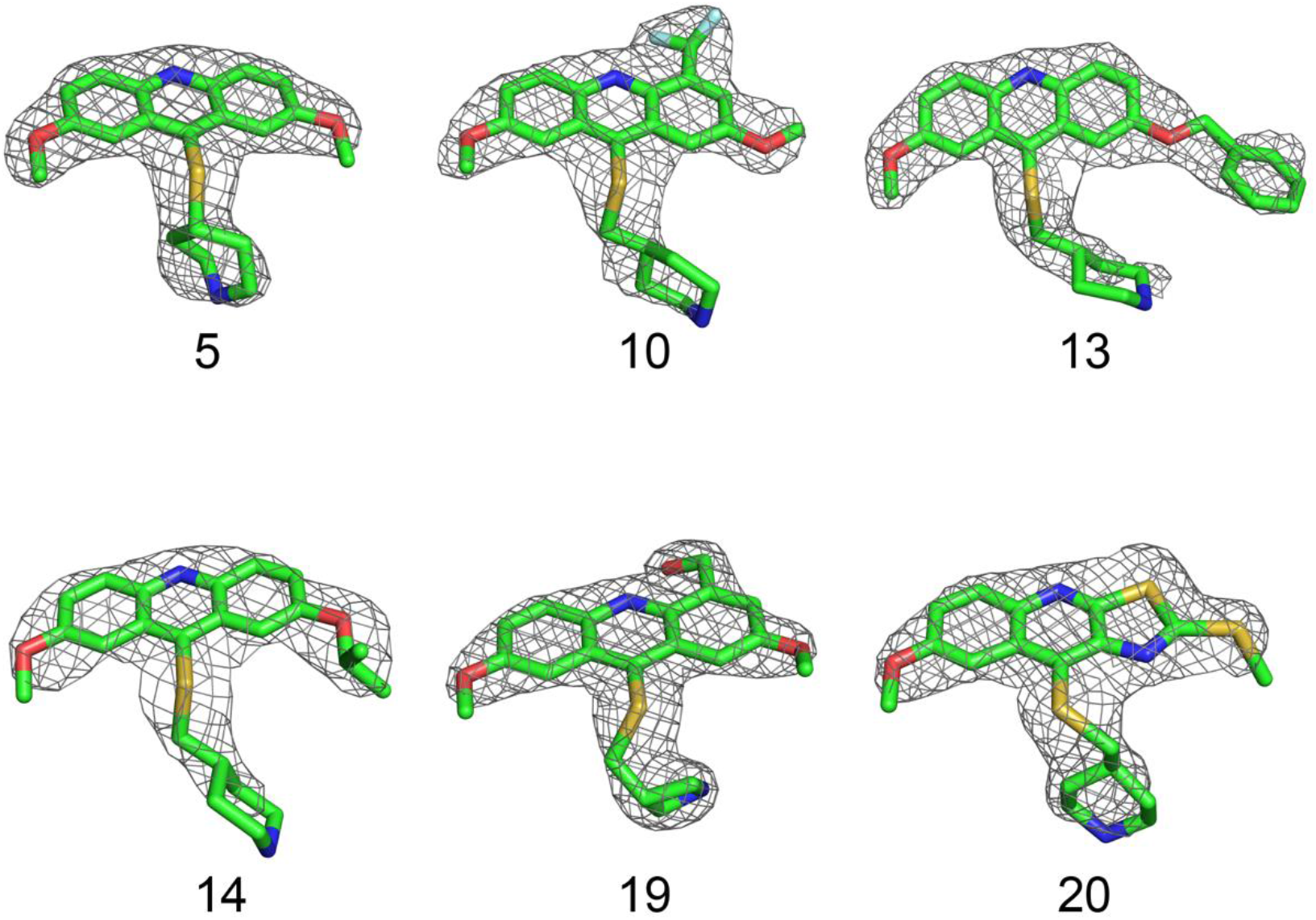
The 2Fo-Fc composite omit maps (1.5σ surrounding compounds 5, 10, 13, 14, 19 and 20 are shown in the co-crystal structures with DRYK2 respectively.

**Extended Data Fig.5.**
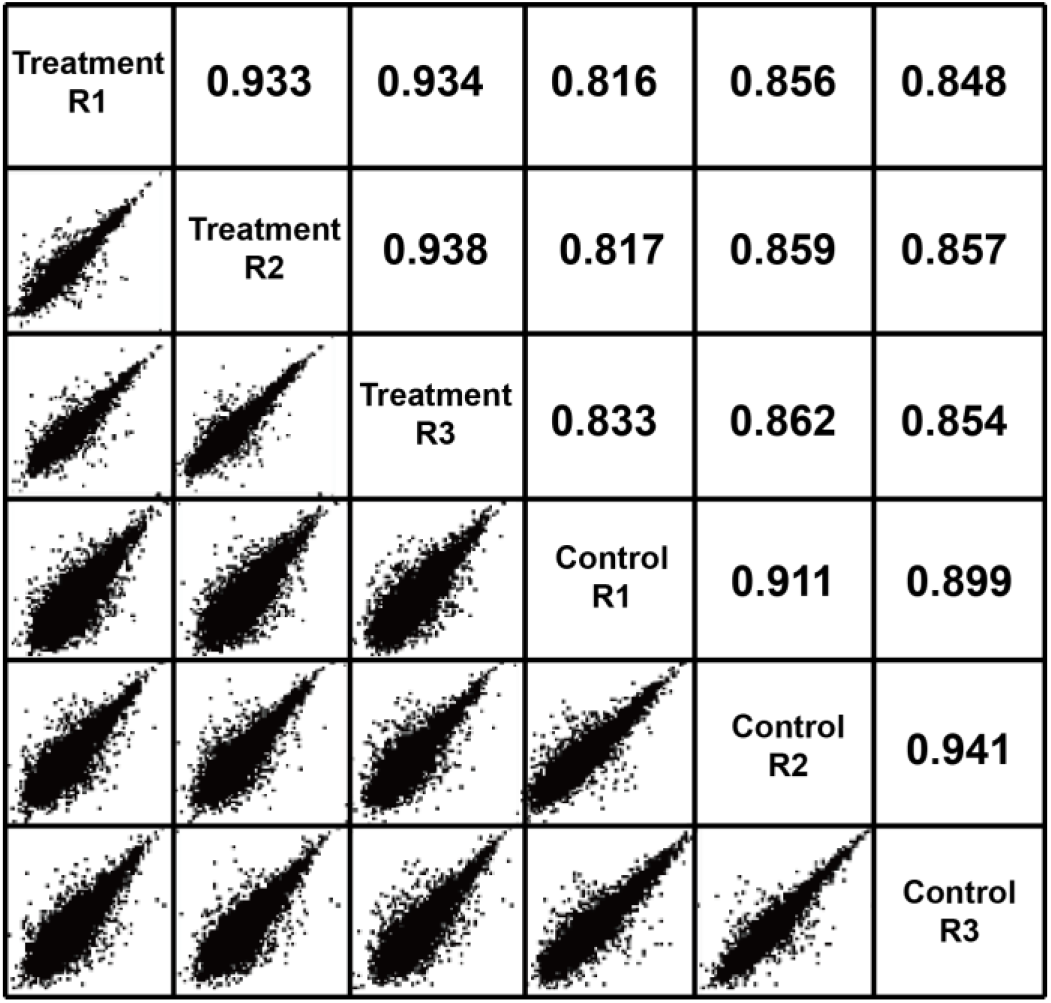
Correlation of the intensities of phosphosites between any two samples in phosphoproteomic analysis of U266 cells treated with/without C17.

**Extended Data Fig.6.**
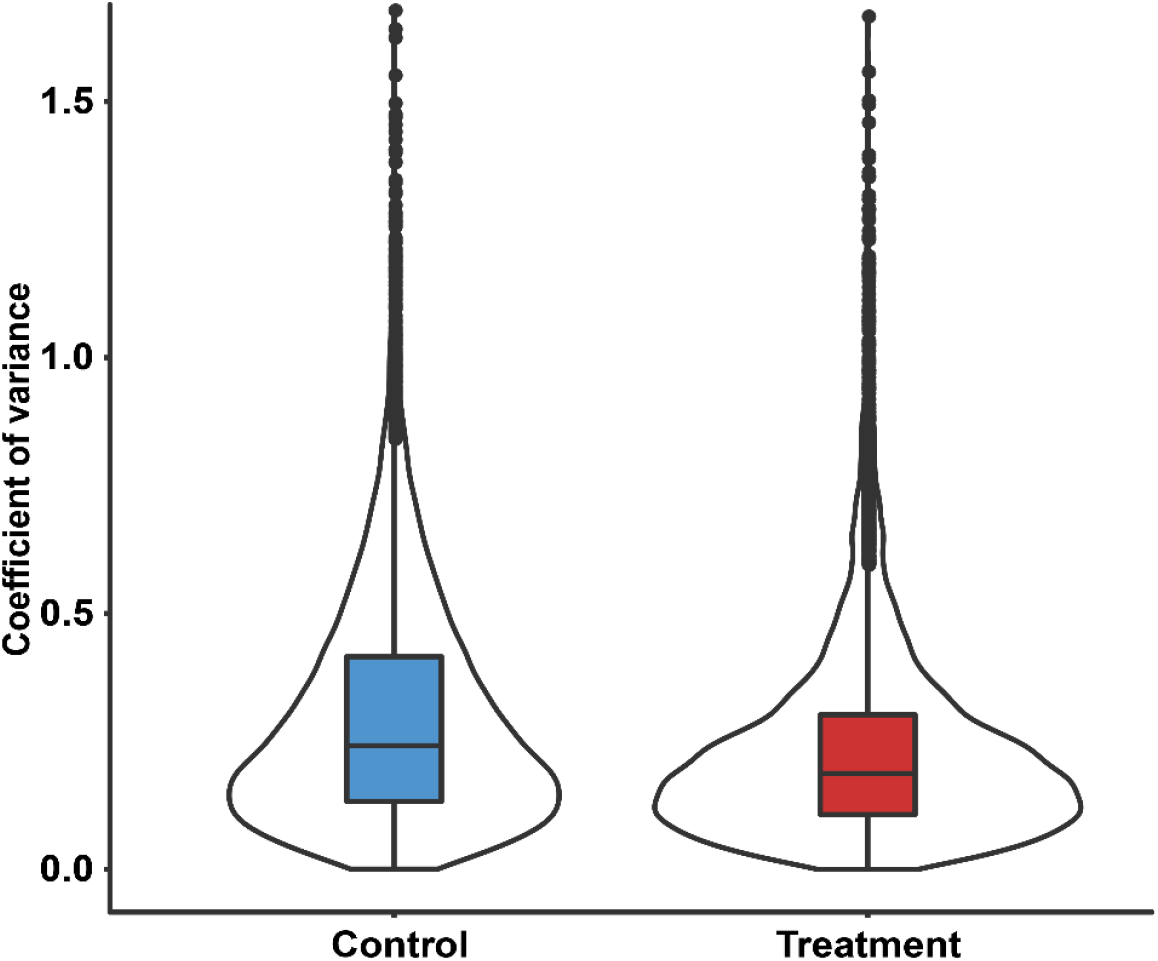
Coefficient of variance of the intensities of phosphosites in phosphoproteomic analysis of U266 cells treated with/without C17.

**Extended Data Fig.7.**
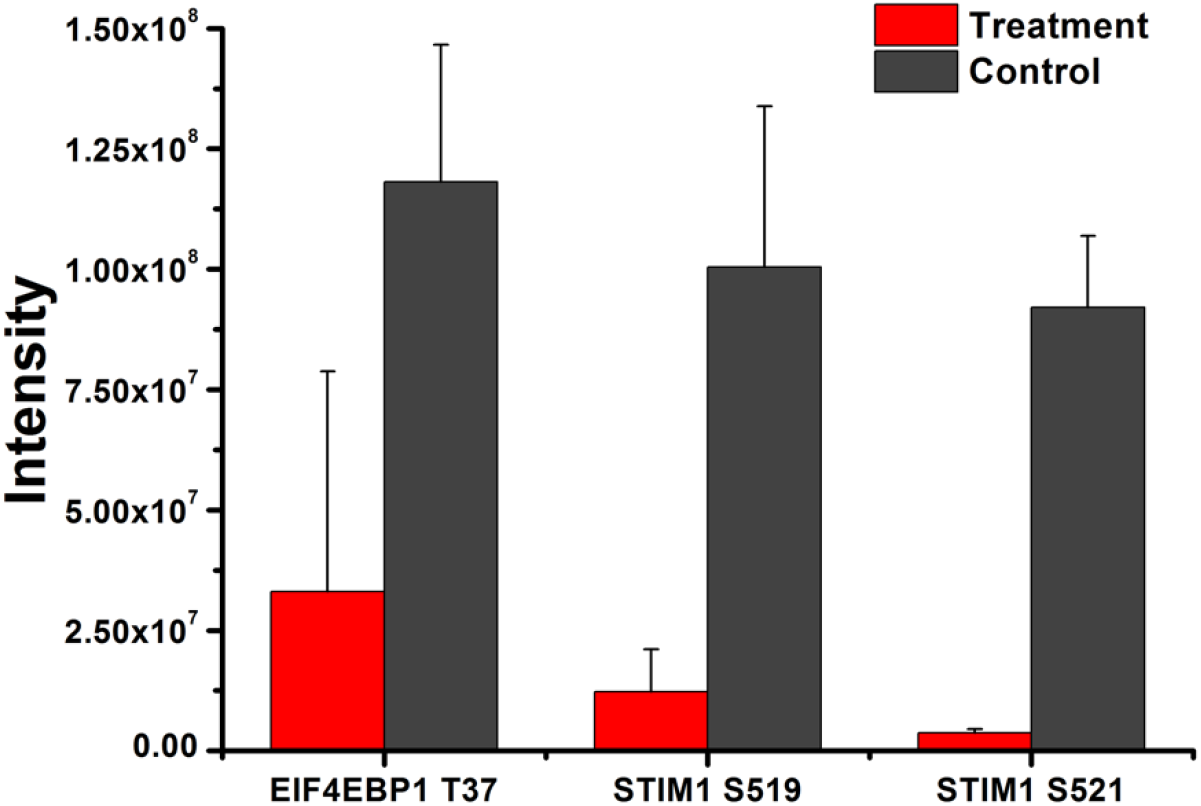
The intensities for pT37 phosphosite of EIF4E-BP1 and pS519, pS521 phosphosites of STIM1.

**Extended Data Fig.8.**
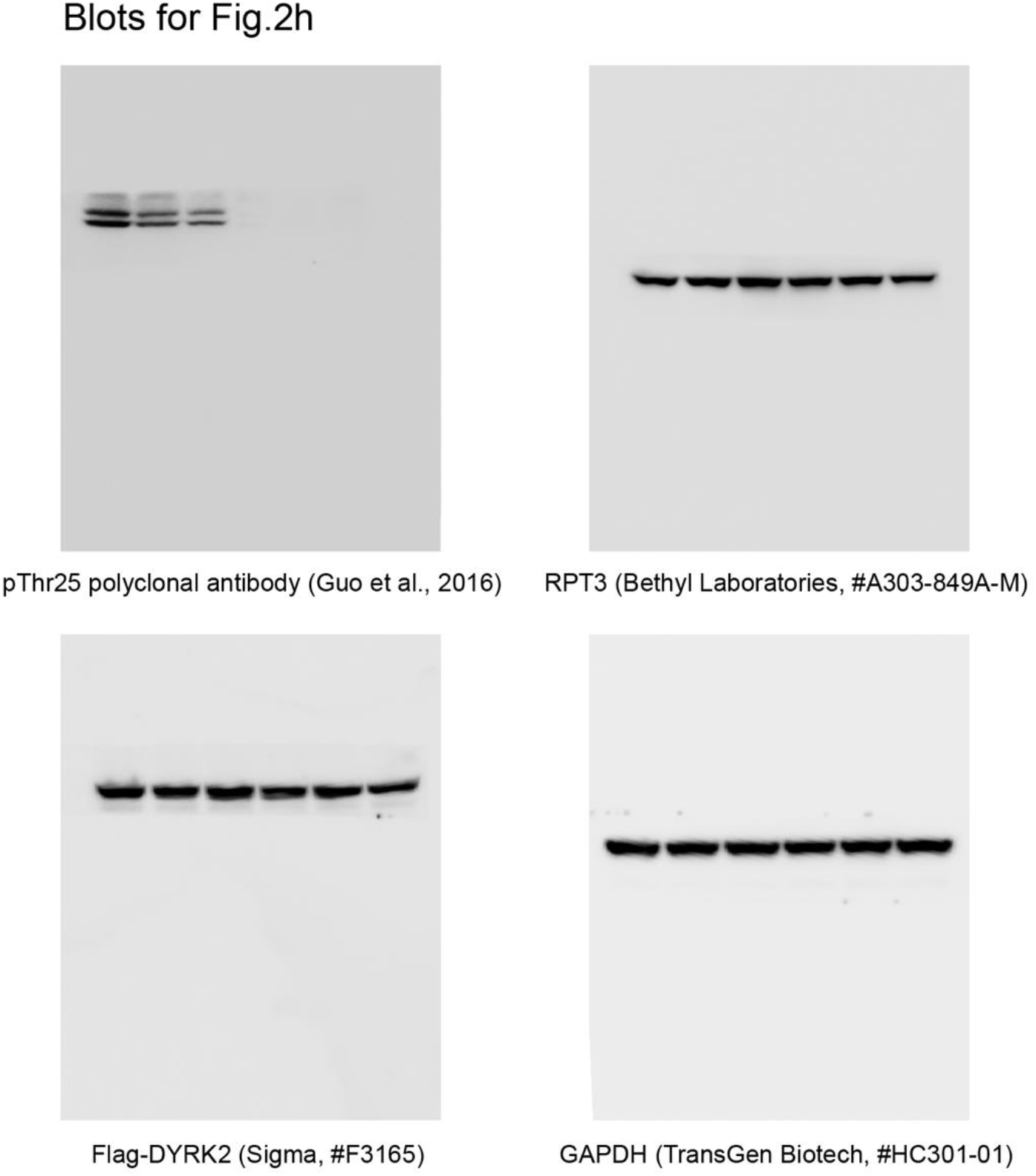
Uncropped immunoblots from supplementary figure 2. In some cases, membranes were cut and the two halves probed with separate antibodies.

**Extended Data Fig.9.**
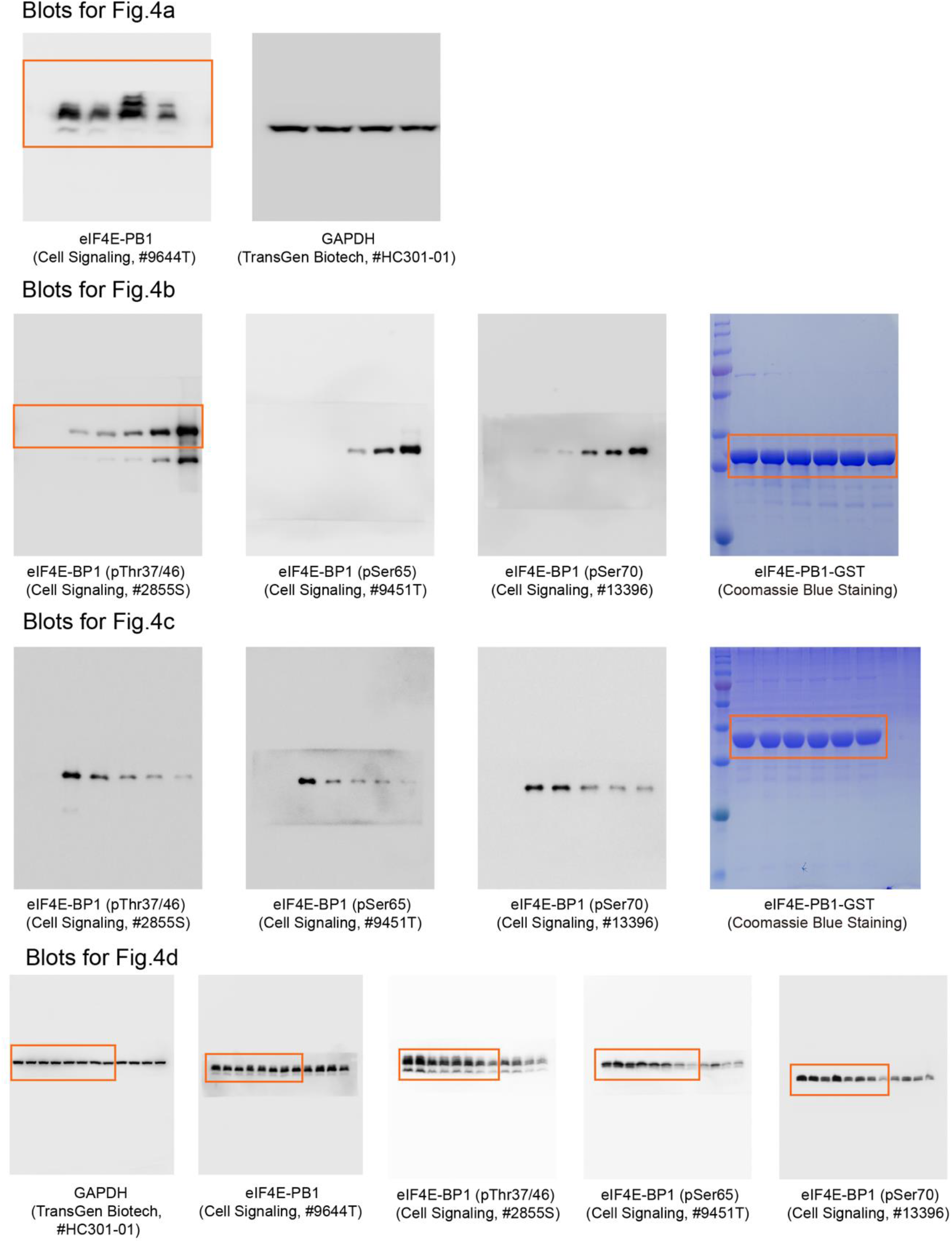
Uncropped immunoblots from supplementary figure 4. In some cases, membranes were cut and the two halves probed with separate antibodies.

**Extended Data Fig.10.**
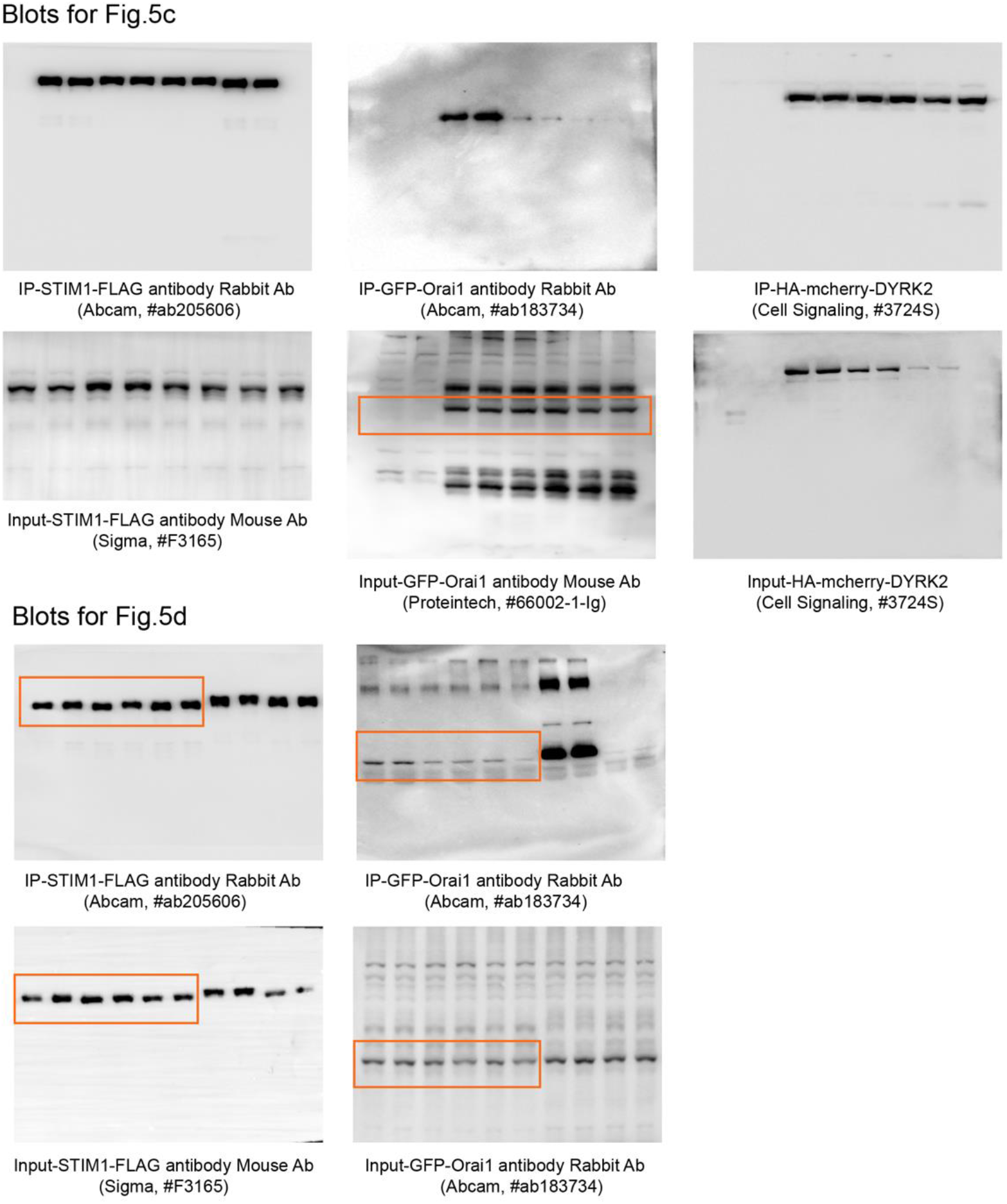
Uncropped immunoblots from supplementary figure 5.

**Extended Data Fig.11.**
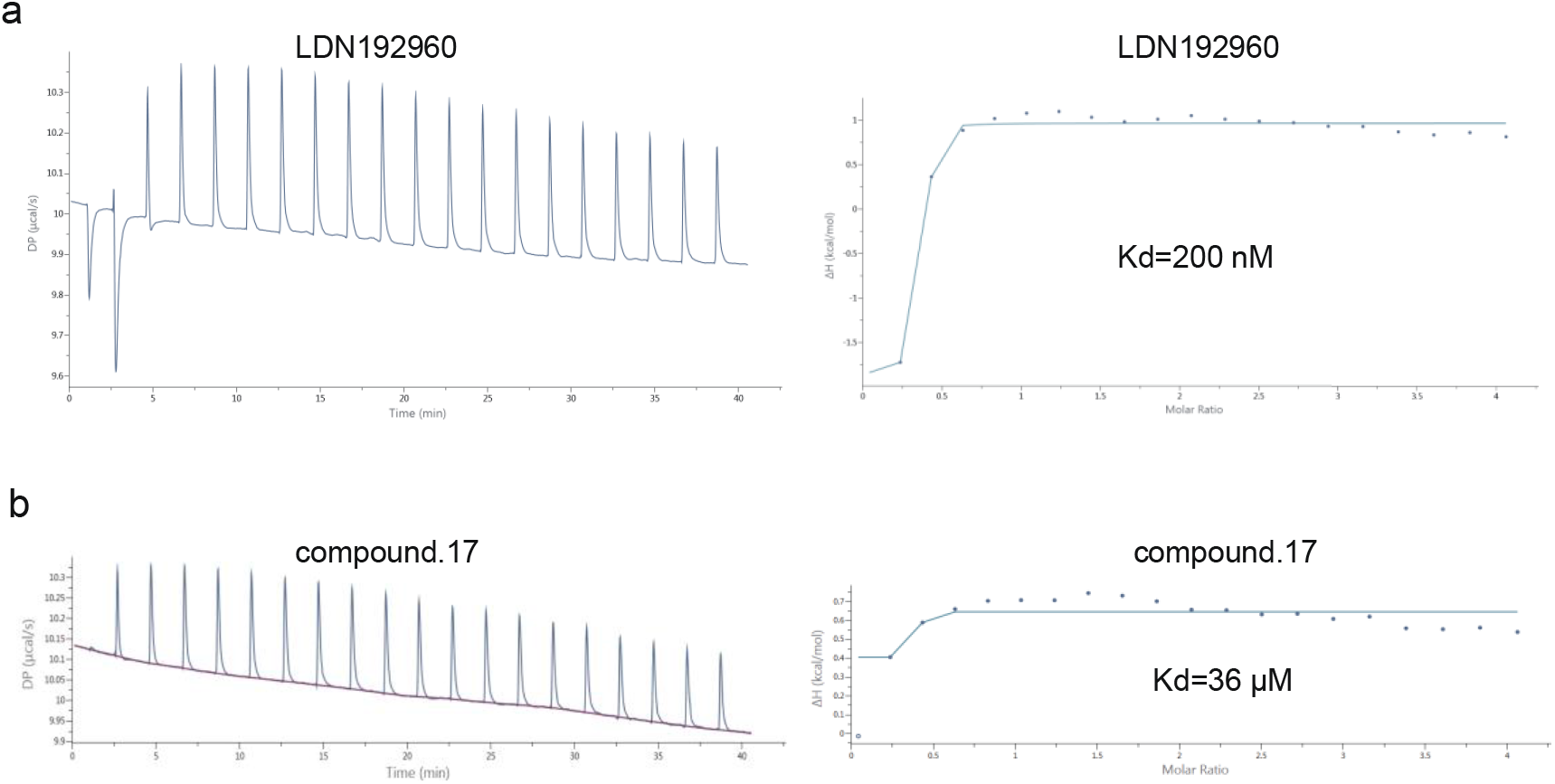
Binding strength of LDN192960 and C17 with calf thymus DNA. **a**, Binding strength of LDN192960 with calf thymus DNA tested by Isothermal titration calorimetry. **b**, Binding strength of compound 17 with calf thymus DNA tested by Isothermal titration calorimetry.

