## Supplementary Information for "Selective inhibition reveals the regulatory function of DYRK2 in protein synthesis and calcium entry"

| | DYRK2-Compound 5 | DYRK2-Compound 6 | DYRK2-Compound 7 | DYRK2-Compound 8 | DYRK2-Compound 10 |
|---|---|---|---|---|---|
| PDB code | 7DH3 | 7DG4 | 7DH9 | 7DHV | 7DHC |
| **Data collection** | | | | | |
| Space group | C 2 2 21 | C 2 2 21 | C 2 2 21 | C 2 2 21 | C 2 2 21 |
| Cell dimensions (Å) | 64.56 128.83 132.45 | 64.58 128.88 132.48 | 64.142 128.44 134.106 | 64.546 128.782 132.735 | 64.427 128.382 132.446 |
| α, β, γ (°) | 90, 90, 90 | 90, 90, 90 | 90, 90, 90 | 90, 90, 90 | 90, 90, 90 |
| Wavelength (Å) | 1.0000 | 1.0000 | 1.0000 | 1.0000 | 1.0000 |
| Resolution (Å) | 50.00-2.33(2.41-2.33) | 50.00-2.58 (2.67-2.58) | 50.00-2.19 (2.27-2.19) | 50.00-2.68(2.78-2.68) | 50.00-2.59(2.69-2.59) |
| CC1/2 | 0.999(0.807) | 0.99(0.818) | 0.998(0.821) | 0.985(0.689) | 0.956(0.786) |
| Rpim | 0.028(0.311) | 0.073(0.389) | 0.037(0.296) | 0.054(0.420) | 0.066(0.448) |
| $R_{merge}$ | 0.093(1.006) | 0.183(0.945) | 0.127(1.001) | 0.186(1.257) | 0.138(0.993) |
| $I$ / σ$I$ | 15.9(2.2) | 7.9(2.3) | 24.4(3.0) | 18.3(2.3) | 11.6(2.4) |
| Completeness (%) | 96.4(99.8) | 99.9 (99.8) | 100.0 (100.0) | 100.0 (100.0) | 99.4 (99.8) |
| redundancy | 12.9(13.5) | 7.7(7.3) | 12.7(12.3) | 12.7(9.7) | 5.4(5.6) |
| Wilson B-factor | 51.65 | 62.02 | 32.11 | 49.20 | 40.65 |
| **Refinement** | | | | | |
| No. reflections | 23126 | 17733 | 28701 | 14996 | 16633 |
| $R_{work}$ / $R_{free}$ | 0.193 / 0.217 | 0.201 / 0.248 | 0.186 / 0.219 | 0.190 / 0.250 | 0.179 / 0.223 |
| No. of atoms | | | | | |
| Protein | 2646 | 2646 | 2646 | 2646 | 2646 |
| Ligand/ion | 26 | 26 | 28 | 29 | 29 |
| Water | 65 | 23 | 215 | 34 | 38 |
| *B*-factors | | | | | |
| Macromolecules | 56.73 | 66.21 | 37.25 | 52.04 | 41.31 |
| Ligand/ion | 54.40 | 59.48 | 33.68 | 55.95 | 39.52 |
| Water | 52.34 | 59.97 | 41.79 | 42.67 | 35.84 |
| R.m.s deviations | | | | | |
| Bond lengths (Å) | 0.009 | 0.009 | 0.008 | 0.008 | 0.009 |
| Bond angles (°) | 1.35 | 1.31 | 1.18 | 1.10 | 1.32 |
| Ramachandran | | | | | |
| Favored (%) | 93.73 | 94.36 | 94.36 | 93.73 | 93.10 |
| Allowed (%) | 6.27 | 5.64 | 5.64 | 5.96 | 6.9 |
| Outliers (%) | 0 | 0 | 0 | 0.31 | 0 |

**Table S1.** Data collection and refinement statistics.

| | DYRK2-C13 | DYRK2-C14 | DYRK2-C17 | DYRK2-C 18 | DYRK2-C19 | DYRK2-C20 |
|---|---|---|---|---|---|---|
| PDB code | 7DHK | 7DHO | 7DJO | 7DL6 | 7DHH | 7DHN |
| **Data collection** | | | | | | |
| Space group | C 2 2 21 | C 2 2 21 | C 2 2 21 | C 2 2 21 | C 2 2 21 | C 2 2 21 |
| Cell dimensions (Å) | 64.557 127.838 132.027 | 64.61 128.87 133.74 | 64.955 128.929 133.548 | 64.662 128.495 132.553 | 64.984 128.466 132.676 | 64.76 128.55 132.64 |
| α, β, γ (°) | 90, 90, 90 | 90, 90, 90 | 90, 90, 90 | 90, 90, 90 | 90, 90, 90 | 90, 90, 90 |
| Wavelength (Å) | 1.0000 | 1.0000 | 1.0000 | 1.0000 | 1.0000 | 1.0000 |
| Resolution (Å) | 50.00-2.34(2.43-2.34) | 50.00-3.29(3.41-3.29) | 50.00-2.49(2.58-2.49) | 50.00-2.65(2.74-2.65) | 50.00-2.21(2.29-2.21) | 50.00-2.38(2.47-2.38) |
| CC1/2 | 0.971(0.588) | 0.925(0.676) | 0.994(0.903) | 0.991(0.816) | 0.997(0.755) | 0.999(0.800) |
| Rpim | 0.120(0.456) | 0.203(0.424) | 0.039(0.247) | 0.053(0.219) | 0.034(0.405) | 0.025(0.334) |
| $R_{merge}$ | 0.257(1.017) | 0.483(1.013) | 0.134(0.829) | 0.149(0.452) | 0.106(1.379) | 0.083(1.133) |
| $I$ / σ$I$ | 6.0(2.2) | 3.6(2.2) | 24.1(4.1) | 13.0(4.0) | 17.3(2.2) | 18.9(2.3) |
| Completeness (%) | 99.8 (99.9) | 99.9(99.8) | 100.0 (100.0) | 99.2 (93.2) | 99.7 (100.0) | 99.8 (99.8) |
| redundancy | 6.4(6.2) | 6.6(6.8) | 13.0(12.3) | 8.3(4.9) | 12.8(13.6) | 13.2(13.3) |
| Wilson B-factor | 33.31 | 52.26 | 35.49 | 42.16 | 45.92 | 58.58 |
| **Refinement** | | | | | | |
| No. reflections | 17793 | 8788 | 19677 | 15541 | 20545 | 22545 |
| $R_{work}$ / $R_{free}$ | 0.195 / 0.257 | 0.184 / 0.254 | 0.199/ 0.239 | 0.173 / 0.238 | 0.213 / 0.213 | 0.185 / 0.223 |
| No. of atoms | | | | | | |
| Protein | 2646 | 2646 | 2646 | 2646 | 2646 | 2646 |
| Ligand/ion | 32 | 28 | 27 | 27 | 26 | 25 |
| Water | 69 | 0 | 112 | 71 | 45 | 40 |
| *B*-factors | | | | | | |
| Macromolecules | 43.72 | 45.46 | 43.94 | 46.27 | 57.45 | 68.62 |
| Ligand/ion | 46.42 | 49.17 | 37.16 | 39.71 | 50.17 | 64.31 |
| Water | 38.66 | 0 | 41.16 | 43.68 | 48.11 | 57.95 |
| R.m.s deviations | | | | | | |
| Bond lengths (Å) | 0.008 | 0.011 | 0.009 | 0.008 | 0.009 | 0.009 |
| Bond angles (°) | 1.14 | 1.40 | 1.08 | 1.00 | 1.05 | 1.09 |
| Ramachandran | | | | | | |
| Favored (%) | 94.36 | 90.28 | 94.04 | 91.85 | 91.22 | 93.73 |
| Allowed (%) | 5.02 | 9.09 | 5.64 | 7.84 | 8.46 | 6.27 |
| Outliers (%) | 0.63 | 0.63 | 0.31 | 0.31 | 0.31 | 0 |

**Table S2.** Data collection and refinement statistics.

Each dataset was collected from a single crystal. Values in parentheses are for highest-resolution shell.

#### General information for chemical synthesis

NMR spectra were recorded on a Varian 400 MHz spectrometer, Bruker 400 MHz NMR spectrometer (ARX400), Bruker 400 MHz NMR spectrometer (AVANCE III), Bruker-500M Hz NMR spectrometer (500M) and Bruker-600M Hz NMR spectrometer (600M) at ambient temperature with CDCl3 as the solvent unless otherwise stated. Chemical shifts are reported in parts per million relative to CDCl3 (1H, δ 7.26; 13C, δ 77.16) and MeOD-d4 (1H, δ 3.31; 13C, δ 49.00). Data for 1H NMR are reported as follows: chemical shift, integration, multiplicity (s = singlet, d = doublet, t = triplet, q = quartet, quint = quintet, sixt = sixtet, m = multiplet) and coupling constants. High-resolution mass spectra were obtained at Peking University Mass Spectrometry Laboratory using a Bruker APEX Flash chromatography. The samples were analyzed by UPLC/MS on a Waters Auto Purification LC/MS system (Waters C18 5 μm 150 X 4.6 mm SunFire separation column) or prepared by HPLC/MS on a Waters Auto Purification LC/MS system (ACQUITY UPLC ® BEH C18 17 μm 2.1X50 mm column). Analytical thin layer chromatography was performed using 0.25 mm silica gel 60-F plates, using 250 nm UV light as the visualizing agent and a solution of phosphomolybdic acid and heat as developing agents. Flash chromatography was performed using 200-400 mesh silica gel. Yields refer to chromatographically pure materials, unless otherwise stated. All reagents were used as supplied by Sigma-Aldrich, J&K, Alfa Aesar Chemicals, TCI. Tetrahydrofuran and diethyl ether were distilled from sodium/ benzophenone ketyl prior to use; the other solvents were distilled from calcium hydride unless otherwise noted. All reactions were carried out in oven-dried glassware under an argon atmosphere unless otherwise noted. Reactions using microwave irradiation were performed on CEM Intellivent Explorer Microwave Reactor.

#### Experimental procedures:

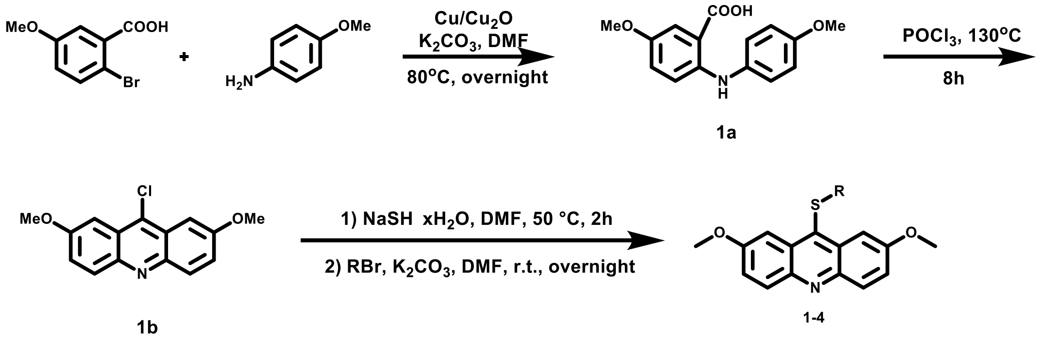

**5-methoxy-2-((4-methoxyphenyl)amino) benzoic acid (1a).** 2-bromo-5-methoxybenzoic acid (9.26 g, 40 mmol), 4-methoxyaniline (6.90 g, 56 mmol), copper (0.73 g, 11 mmol), cuprous oxide (0.82 g, 5.7 mmol) and potassium carbonate (7.74 g, 56 mmol) were added to 100 ml DMF, the mixture was stirred at 80 °C overnight. The resulting slurry was cooled to room temperature, and 2M HCl was added into the mixture until the system became acidic and a large amount of solid was precipitated. After filtration, the precipitate was washed with water and dried to give compound **1a** (4.02 g, 37%) as a dark green solid. **1a** was used in next step without further purification.

**9-chloro-2,7-dimethoxyacridine (1b).** Compound **1a** (2.58 g, 9.45 mmol) was added in a sealed tube, and 30 ml of phosphorus oxychloride was added under argon atmosphere. The reaction was heated at 130 °C for 8 h. The resulting slurry was poured onto ice with vigorous stirring, and a large amount of a yellow solid was precipitated. After filtration, the precipitate was washed with water and dried to give compound **1b** (1.65 g, quant.) as an orange solid. **1b** was used in next step without further purification.

***tert*-butyl (3-((2,7-dimethoxyacridin-9-yl)thio)propyl)carbamate (1).** To a solution of **1b** (50 mg, 0.183 mmol) in 5 ml of anhydrous DMF, sodium hydrogen hydride hydrate powder (67% 22.9 mg, 0.274 mmol) was added under argon atmosphere, and the reaction was stirred at 50 °C for 2 h until full conversion of **1b**. *N*-Boc-3-aminopropyl bromide (60.9 mg, 0.274 mmol) and potassium carbonate (50.5 mg, 0.365 mmol) were added into the slurry and the reaction was allowed to react at room temperature overnight. The solvent was then evaporated and the residue was dissolved with dichloromethane and washed with water. The combined organic extracts were dried over anhydrous $Na_2SO_4$ and concentrated *in vacuo*. The residue was purified by chromatography on a silica gel column to give compound **1** as a light yellow solid (47.4 mg, 61%). $^1$H NMR (400 MHz, CDCl3) δ 8.10 (d, *J* = 9.3 Hz, 2H), 7.94 (s, 2H), 7.42 (d, *J* = 9.3 Hz, 2H), 4.04 (s, 6H), 3.20 (d, *J* = 5.9 Hz, 2H), 2.95 (t, *J* = 7.2 Hz, 2H), 1.70 - 1.61 (m, 2H), 1.39 (s, 9H). $^{13}$C NMR (151 MHz, Chloroform-d) δ 158.5, 156.0, 144.3, 136.0, 132.1, 130.7, 124.1, 102.2, 79.5, 55.8, 39.6, 33.6, 30.8, 28.5. HRMS(ESI) [M + H]$^+$ calculated for $C_{23}H_{29}N_2O_4S$: 429.1843, found: 429.1831.

**Compounds 2-4**. By employment of the above-described procedure, starting from **1b** and using suitable bromides, compounds 2-4 were prepared.

***N*-(3-((2,7-dimethoxyacridin-9-yl)thio)propyl)acetamide (2). Yield 58%.** $^1$H NMR (400 MHz, CDCl3) δ 8.10 (d, *J* = 9.4 Hz, 2H), 7.94 (d, *J* = 2.7 Hz, 2H), 7.42 (dd, *J* = 9.4, 2.8 Hz, 2H), 4.04 (s, 6H), 3.30 (dd, *J* = 13.2, 6.7 Hz, 2H), 2.96 (t, *J* = 7.3 Hz, 2H), 1.88 (s, 3H), 1.69 -

1.59 (m, 2H). [13]C NMR (151 MHz, Chloroform-d) δ 170.2, 158.6, 144.3, 135.8, 132.2, 130.7, 124.1, 102.2, 55.8, 38.7, 33.6, 30.4, 23.4. HRMS(ESI) [M + H]+ calculated for $C_{20}H_{23}N_2O_3S$: 371.1424, found: 371.1428.

***N*-(4-((2,7-dimethoxyacridin-9-yl)thio)butyl)acetamide (3). Yield 57%.** [1]H NMR (400 MHz, CDCl3) δ 8.10 (d, *J* = 9.3 Hz, 2H), 7.95 (d, *J* = 2.7 Hz, 2H), 7.42 (dd, *J* = 9.3, 2.7 Hz, 2H), 4.04 (s, 6H), 3.15 (dd, *J* = 13.1, 6.7 Hz, 2H), 2.94 (t, *J* = 7.1 Hz, 2H), 1.86 (s, 3H), 1.59 (m, 2H), 1.49 (m, 2H). [13]C NMR (101 MHz, Chloroform-d) δ 170.1, 158.4, 144.2, 136.2, 132.0, 130.7, 124.0, 102.2, 55.8, 39.0, 35.9, 28.9, 27.5, 23.3. HRMS(ESI) [M + H]+ calculated for $C_{21}H_{25}N_2O_3S$: 385.1580, found: 385.1572.

**4-((2,7-dimethoxyacridin-9-yl)thio)butanenitrile (4). Yield 52%.** [1]H NMR (400 MHz, CDCl3) δ 8.11 (d, *J* = 9.4 Hz, 2H), 7.89 (d, *J* = 2.7 Hz, 2H), 7.43 (dd, *J* = 9.4, 2.8 Hz, 2H), 4.04 (s, 6H), 3.07 (t, *J* = 7.0 Hz, 2H), 2.47 (t, *J* = 7.0 Hz, 2H), 1.82 - 1.72 (m, 2H). [13]C NMR (101 MHz, Chloroform-d) δ 158.7, 144.3, 134.4, 132.3, 130.6, 124.2, 118.8, 101.7, 55.8, 34.4, 25.8, 16.4. HRMS(ESI) [M + H]+ calculated for $C_{19}H_{19}N_2O_2S$: 339.1162, found: 339.1159.

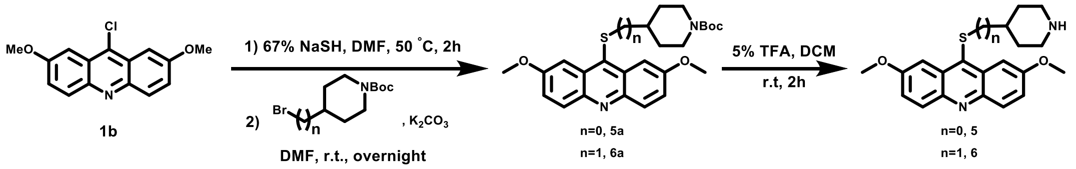


**tert-butyl 4-((2,7-dimethoxyacridin-9-yl)thio)piperidine-1-carboxylate (5a).** To a solution of **1b** (50 mg, 0.183 mmol) in 5 ml of anhydrous DMF, sodium hydrogen hydride hydrate powder (67% 22.9 mg, 0.274 mmol) was added under argon atmosphere, and the reaction was stirred at 50 °C for 2 h until full conversion of **1b**. 4-bromopiperidine-1-carboxylic acid tert-butyl ester (72.4mg，0.274mmol) and potassium carbonate (50.5 mg, 0.365 mmol) were added into the slurry and the reaction was allowed to react at room temperature overnight. The solvent was then evaporated and the residue was dissolved with dichloromethane and washed with water. The combined organic extracts were dried over anhydrous $Na_2SO_4$ and concentrated *in vacuo*. The residue was purified by chromatography on a silica gel column to give compound **5a** as a light yellow solid (45.4mg, 55%). [1]H NMR: (400 MHz, $CDCl_3$) δ 8.08 (d, *J* = 9.3 Hz, 2H), 7.93 (d, *J* = 2.7 Hz, 2H), 7.40 (dd, *J* = 9.3, 2.8 Hz, 2H), 4.01 (s, 6H), 3.98-3.84 (m, 2H), 3.25 - 3.17 (m, 1H), 2.81 (ddd, *J* = 13.5, 10.5, 3.0 Hz, 2H), 1.86-1.72 (m, 2H), 1.72-1.58 (m, 2H), 1.42 (s, 9H). [13]C NMR (101 MHz, Chloroform-d) δ 158.4, 154.7, 144.3, 134.8, 132.0, 131.1, 124.0, 102.4, 79.8, 55.7, 46.9, 43.2, 33.1, 28.5. HRMS(ESI) [M + H]+ calculated for $C_{25}H_{31}N_2O_4S$: 455.1999, found: 455.1995.

**Compounds 6a**. By employment of the above-described procedure, starting from **1b** and using suitable bromide, compound **6a** were prepared.

**tert-butyl 4-(((2,7-dimethoxyacridin-9-yl)thio)methyl)piperidine-1-carboxylate (6a). Yield 79%.** $^{1}$H NMR (400 MHz, $CDCl_3$) δ 8.10 (d, *J* = 9.4 Hz, 2H), 7.94 (d, *J* = 2.7 Hz, 2H), 7.42 (dd, *J* = 9.4, 2.8 Hz, 2H), 4.03 (d, *J* = 48.7 Hz, 6H), 4.15 - 3.98 (overlapped, m, 2H), 2.82 (d, *J* = 6.8 Hz, 2H), 2.59 (t, *J* = 12.1 Hz, 2H), 1.86 (br s, 2H), 1.53 - 1.46 (m, 1H), 1.43 (s, 9H), 1.23-1.14 (m, *J* = 10.9 Hz, 2H). $^{13}$C NMR (101 MHz, Chloroform-d) δ 158.4, 154.8, 144.2, 136.7, 132.0, 130.4, 123.9, 102.0, 79.5, 55.6, 43.6, 42.9, 36.8, 28.5. HRMS(ESI) $[M + H]^+$ calculated for $C_{26}H_{36}N_2O_4S$: 469.2156, found: 469.2153.

**2,7-dimethoxy-9-(piperidin-4-ylthio)acridine (5).** Compound **5a** (45.4mg, 0.100 mmol) was dissolved in a 5% trifluoroacetic acid dichloromethane solution and the mixture was allowed to react at room temperature for 2h. The solvent was then evaporated and the residue was dissolved with methanol, then purified by HPLC/MS on a Waters Auto Purification LC/MS system (ACQUITY UPLC ® BEH C18 17 µm 2.1X50 mm column) to afford **5** as a dark red solid (35.2mg, 75%). $^{1}$H NMR (400 MHz, MeOD) δ 8.05 (d, *J* = 9.4 Hz, 2H), 8.00 (d, *J* = 2.7 Hz, 2H), 7.49 (dd, *J* = 9.4, 2.8 Hz, 2H), 4.05 (s, 6H), 3.36-3.33 (m, 1H), 2.99 (d, *J* = 6.8 Hz, 2H), 2.85 (td, *J* = 12.8, 2.9 Hz, 2H), 2.13 (d, *J* = 16.5 Hz, 2H), 1.70-1.65 (m, 2H). $^{13}$C NMR (151 MHz, Methanol-d4) δ 160.9, 144.0, 138.6, 132.9, 129.7, 126.7, 104.0, 56.0, 45.5, 44.4, 30.9. HRMS(ESI) $[M + H]^+$ calculated for $C_{20}H_{23}N_2O_2S$: 355.1475, found: 355.1467.

**Compounds 6**. By employment of the above-described procedure, starting from **6a**, compounds **6** was prepared.

**2,7-dimethoxy-9-((piperidin-4-ylmethyl)thio)acridine (6). Yield 56%.** $^{1}$H NMR (400 MHz, MeOD) δ 8.24 (d, *J* = 9.4 Hz, 2H), 8.17 (d, *J* = 2.3 Hz, 2H), 7.85 (dd, *J* = 9.4, 2.4 Hz, 2H), 4.14 (s, 6H), 3.75-3.68 (m, 1H), 3.39 (d, *J* = 13.3 Hz, 2H), 3.01 (t, *J* = 10.9 Hz, 2H), 2.18 - 2.09 (m, 2H), 2.05 - 2.01 (m, 2H), 1.37 - 1.32 (m, 2H). $^{13}$C NMR (101 MHz, Methanol-d4) δ 61.0, 151.0, 135.6, 131.8, 131.1, 124.0, 104.0, 56.8, 44.7, 44.2, 35.9, 29.1. HRMS(ESI) $[M + H]^+$ calculated for $C_{21}H_{25}N_2O_2S$: 369.1631, found: 369.1638.

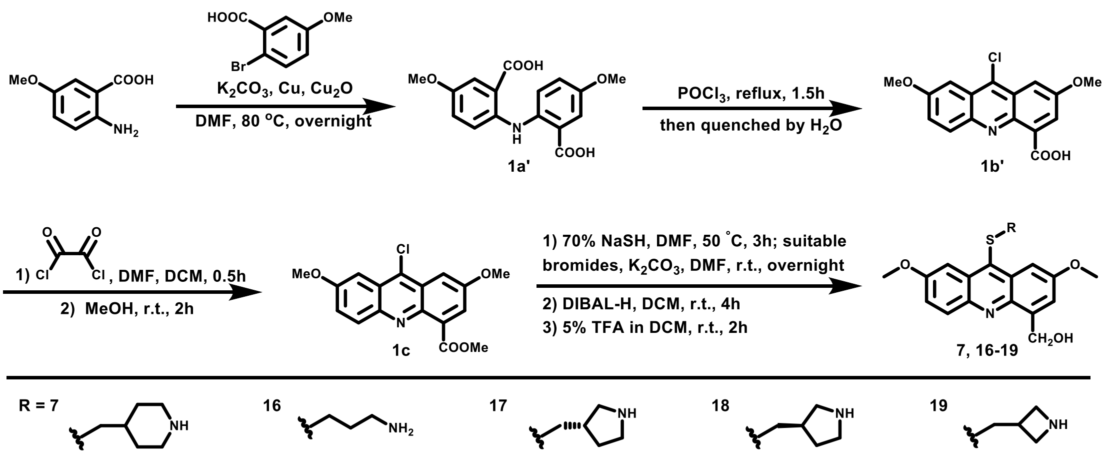


**6,6'-azanediylbis-3-methoxybenzoic acid (1a').** 2-amino-5-methoxybenzoic acid (2.00g, 8.66mmol), 2-bromo-5-methoxybenzoic acid (0.11g, 1.73mmol), copper (0.11g, 1.73mmol), cuprous oxide (0.12g, 0.87mmol) and potassium carbonate (7.74 g, 56 mmol) were added to 15ml DMF, the mixture was stirred at 80 °C overnight. The resulting slurry was cooled to room temperature, and 2M HCl was added into the mixture until the system became acidic and a large amount of solid was precipitated. After filtration, the precipitate was washed with water and dried to give compound **1a'** (2.00 g, 73%) as a green solid. **1a'** was used in next step without further purification.

**9-chloro-2,7-dimethoxyacridine-4-carboxylic acid (1b').** Compound **1a'** (1.00g, 3.16mmol) was added in a sealed tube, and 10 ml of phosphorus oxychloride was added under argon atmosphere. The reaction was heated at 130 °C for 8 h. The resulting slurry was poured onto ice with vigorous stirring, and a large amount of a yellow solid was precipitated. After filtration, the precipitate was washed with water and dried to give compound **1b** (1.00g, 95%) as an orange solid. **1b'** was used in next step without further purification.

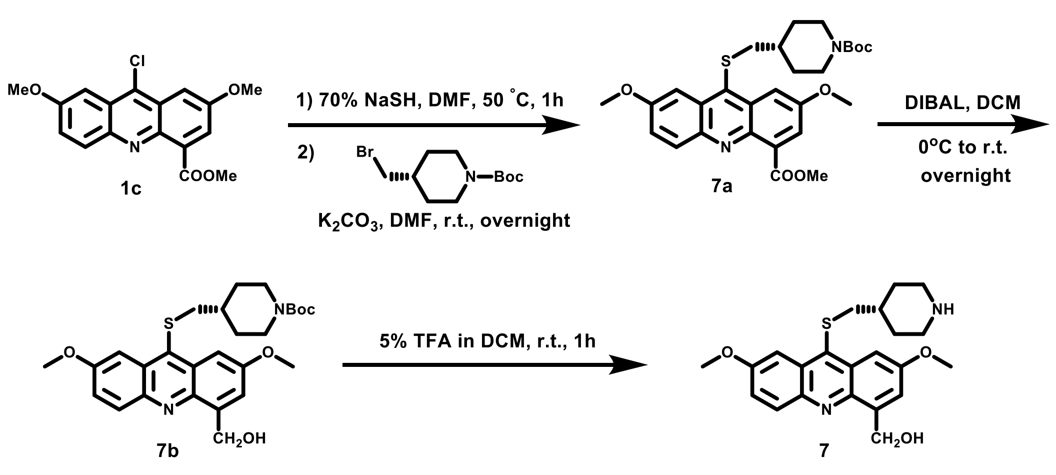


**methyl 9-chloro-2,7-dimethoxyacridine-4-carboxylate (1c).** To a suspension of **1b'** (0.89g, 2.67mmol) in 10 ml of dry dichloromethane, 0.40ml of oxalyl chloride was added followed by one drop of DMF, and a large amount of bubbles was generated. The mixture was allowed to react at room temperature for 0.5 h until the system became a brownish black solution. Then the reaction was quenched by dry menthol at room temperature for 2 h, followed by the addition of triethylamine until the mixture became neutral. The system was diluted with dichloromethane, washed twice with brine, dried over anhydrous $Na_2SO_4$ and concentrated *in vacuo*. The residue was purified by chromatography on a silica gel column (dichloromethane / ethyl acetate = 95/5) to gave compound **1c** as a yellow solid (0.74 g, 80%).$^1$H NMR (400 MHz, $CDCl_3$) δ 8.12 (d, *J* = 9.3 Hz, 1H), 7.70 (s, 1H), 7.64 (s, 1H), 7.46 (s, 1H), 7.42 (d, *J* = 9.4 Hz, 1H), 4.09 (s, 3H), 4.03 (s, 6H). $^{13}$C NMR (101 MHz, Chloroform-d) δ 167.9, 159.0, 157.1, 144.8, 141.3, 135.9, 134.1, 132.6, 125.8, 125.7, 125.0, 124.9, 103.0, 99.6, 56.0, 55.8, 52.3. HRMS(ESI) $[M + H]^+$ calculated for $C_{17}H_{15}ClNO_4$: 332.0684, found: 332.0680.

**methyl-9-(((1-(tert-butoxycarbonyl)piperidin-4-yl)methyl)thio)-2,7-dimethoxyacridine-4-caboxylate (7a).** To a solution of **1c** (0.38g, 1.16mmol) in 10 ml of anhydrous DMF, sodium hydrogen hydride hydrate powder (70%, 0.10g，1.21mmol) was added under argon atmosphere, and the reaction was stirred at 50 °C for 2 h until full conversion of **1c**. 1-BOC-4-bromomethylpiperidine (0.64g，2.31mmol) and potassium carbonate (0.40g，2.89mmol) were added into the slurry and the reaction was allowed to react at room temperature overnight. The solvent was then evaporated and the residue was dissolved with dichloromethane and washed with water. The combined organic extracts were dried over anhydrous $Na_2SO_4$ and concentrated *in vacuo*. The residue was purified by chromatography on a silica gel column (dichloromethane / ethyl acetate = 95/5) to give compound **7a** as a yellow solid (0.38g, 63%). $^1$H NMR (400 MHz, $CDCl_3$) δ 8.13 (d, *J* = 9.4 Hz, 1H), 8.08 (d, *J* = 2.0 Hz, 1H), 7.89 (s, 1H), 7.68 (d, *J* = 1.9 Hz, 1H), 7.41 (dd, *J* = 9.4, 1.9 Hz, 1H), 4.09 (s, 3H), 4.02 (s, 6H), 2.78 (d, *J* = 6.7 Hz, 2H), 2.55 (t, *J* = 12.2 Hz, 2H), 1.82 (br s, 2H), 1.45-1.35(m, 3H), 1.43 (s, 9H), 1.20-1.09 (m, 2H). $^{13}$C NMR (101 MHz, Chloroform-d) δ 168.2, 158.9, 157.0, 154.8, 144.7, 141.2, 137.0, 134.6, 133.0, 130.7, 130.5, 124.4, 124.3, 105.11, 101.74, 79.6, 56.0, 55.7, 52.8, 43.8, 43.1 (br s), 36.8, 31.8, 28.5. HRMS(ESI) $[M + H]^+$ calculated for $C_{28}H_{35}N_2O_6S$: 527.2210, found: 527.2212.

**Compounds 16a**-**19a**. By employment of the above-described procedure, starting from **1c** and using suitable bromides, compounds **16a**-**19a** were prepared.

**methyl 9-((3-((tert-butoxycarbonyl)amino)propyl)thio)-2,7-dimethoxyacridine-4-carboxylate (16a). Yield 76%.** $^1$H NMR (400 MHz, Chloroform-d) δ 8.12 (d, *J* = 9.4 Hz, 1H),

8.09 (d, *J* = 2.8 Hz, 1H), 7.89 (d, *J* = 2.8 Hz, 1H), 7.68 (dd, *J* = 2.9, 0.8 Hz, 1H), 7.41 (dd, *J* = 9.3, 2.8 Hz, 1H), 4.40 (br s, 1H), 4.09 (s, 3H), 4.04 (s, 3H), 4.03 (s, 3H), 3.17 (q, *J* = 6.6 Hz, 2H), 2.92 (t, *J* = 7.3 Hz, 2H), 1.61-1.59 (m, 2H), 1.39 (s, 9H). $^{13}$C NMR (101 MHz, Chloroform-d) δ 168.2, 158.8, 157.0, 155.9, 144.57, 141.08, 136.2, 134.5, 132.9, 130.8, 130.6, 124.3, 124.3, 105.1, 101.8, 55.9, 55.7, 52.8, 39.5, 33.7, 30.7, 28.4. HRMS(ESI) [M + H]$^+$ calculated for $C_{25}H_{31}N_2O_6S$: 487.1889, found: 487.1897.

**methyl-(S)-9-(((1-(tert-butoxycarbonyl)pyrrolidin-3-yl)methyl)thio)-2,7-dimethoxyacridine-4-carboxylate (17a). Yield 76%.** $^1$H NMR (400 MHz, $CDCl_3$) δ 8.13 (d, *J* = 9.4 Hz, 1H), 8.08 (d, *J* = 2.4 Hz, 1H), 7.88 (d, *J* = 2.6 Hz, 1H), 7.68 (d, *J* = 2.7 Hz, 1H), 7.42 (dd, *J* = 9.4, 2.6 Hz, 1H), 4.10 (s, 3H), 4.05 (s, 3H), 4.04 (s, 3H), 3.61-3.14 (m, 2H), 3.27-2.1 (m, 4H), 2.10-2.01 (m, 1H), 1.90-1.85 (m, 1H), 1.68-1.59 (m, 1H), 1.43 (s, 9H). $^{13}$C NMR (101 MHz, Chloroform-d) δ 168.2, 159.0, 157.1, 154.5, 144.6, 141.1, 135.9, 134.7, 133.1, 130.7, 130.5, 124.4, 124.3, 104.9, 101.6, 79.4, 55.9 (d, *J* = 96 Hz), 52.8, 51.0 (d, *J* = 104 Hz), 45.2 (d, *J* = 104 Hz), 39.6-39.3 (m), 38.4, 31.3, 30.7, 28.6. HRMS(ESI) [M + H]$^+$ calculated for $C_{27}H_{33}N_2O_6S$: 513.2052, found: 513.2054.

**methyl-(R)-9-(((1-(tert-butoxycarbonyl)pyrrolidin-3-yl)methyl)thio)-2,7-dimethoxyacridine-4-carboxylate (18a). Yield 84%.** $^1$H NMR (400 MHz, $CDCl_3$) δ 8.13 (d, *J* = 9.4 Hz, 1H), 8.07 (s, 1H), 7.88 (s, 1H), 7.68 (s, 1H), 7.41 (d, *J* = 8.8 Hz, 1H), 4.09 (s, 3H), 4.04 (s, 6H), 3.62-3.31 (m, 2H), 3.26-3.18 (m, 1H), 3.10-2.79 (m, 3H), 2.11-1.99 (m, 1H), 1.97-1.86 (m, 1H), 1.65-1.56 (m, 1H), 1.43 (s, 9H). $^{13}$C NMR (126 MHz, Chloroform-d) δ 168.2, 159.1, 157.2, 154.5, 144.7, 141.3, 135.9, 134.8, 133.2, 130.8, 130.6, 124.4, 124.3, 105.0, 101.7, 79.4, 55.9(d, *J* = 105 Hz), 52.8, 51.1 (d, *J* = 115 Hz), 45.2 (d, *J* = 135 Hz), 39.7-39.4 (m), 38.5, 31.4, 30.7, 28.6. HRMS(ESI) [M + H]$^+$ calculated for $C_{27}H_{33}N_2O_6S$: 513.2052, found: 513.2054.

**methyl-9-(((1-(tert-butoxycarbonyl)azetidin-3-yl)methyl)thio)-2,7-dimethoxyacridine-4-carboxylate (19a). Yield 75%.** $^1$H NMR (400 MHz, $CDCl_3$) δ 8.14 (d, *J* = 9.4 Hz, 1H), 8.03 (s, 1H), 7.84 (s, 1H), 7.68 (d, *J* = 1.6 Hz, 1H), 7.42 (d, *J* = 9.4 Hz, 1H), 4.10 (s, 3H), 4.04 (s, 6H), 3.85 (t, *J* = 8.3 Hz, 2H), 3.60 (s, 2H), 3.10 (d, *J* = 7.8 Hz, 2H), 2.26-2.16 (m, 1H), 1.40 (s, 9H). $^{13}$C NMR (126 MHz, Chloroform-d) δ 168.2, 159.2, 157.3, 156.3, 144.7, 141.2, 134.9, 134.8, 133.2, 130.9, 130.7, 124.5, 124.3, 104.9, 101.6, 79.7, 56.0, 55.8, 54.0(br s), 52.8, 39.9, 29.0, 28.5. HRMS(ESI) [M + H]$^+$ calculated for $C_{26}H_{31}N_2O_6S$: 499.1897, found: 499.1895.

***tert*-butyl-4-(((4-(hydroxymethyl)-2,7-dimethoxyacridin-9-yl)thio)methyl)piperidine-1-carboxylate (7b).** 3.1ml of 1.5M DIBAL-H solution in toluene was added to a solution of **7a**

(348mg，0.661mmol) in 10 ml of dry dichloromethane at 0 °C, and the reaction was stirred at room temperature for 4h. The reaction was quenched by adding saturated potassium hydrogen tartrate solution, diluted with dichloromethane, washed twice with brine, dried over anhydrous $Na_2SO_4$ and concentrated *in vacuo*. The residue was purified by chromatography on a silica gel column (dichloromethane / ethyl acetate = 90/10) to gave compound **7b** (180mg, 55%) as yellow foam. $^1$H NMR (400 MHz, $CDCl_3$) δ 8.03 (d, *J* = 9.3 Hz, 1H), 7.88 (s, 1H), 7.81 (s, 1H), 7.39 (d, *J* = 9.3 Hz, 1H), 7.27 (s, 1H), 5.41 (s, 1H), 5.21 (s, 2H), 4.02 (s, 3H), 4.00 (s, 3H) 2.79 (d, *J* = 6.7 Hz, 2H), 2.57 (t, *J* = 11.7 Hz, 2H), 1.84 (br s, 2H), 1.47-1.25 (m, 3H), 1.43 (s, 9H), 1.20-1.09 (m, 2H). $^{13}$C NMR (101 MHz, Chloroform-d) δ 158.5, 157.9, 154.8, 143.3, 142.7, 140.4, 137.6, 131.9, 131.1, 130.5, 124.1, 122.1, 102.0, 101.4, 79.6, 65.0, 55.7, 55.7, 43.6, 43.1, 36.8, 31.8, 28.5. HRMS(ESI) $[M + H]^+$ calculated for $C_{27}H_{35}N_2O_5S$: 499.2258, found: 499.2261.

**Compounds 16b-19b**. By employment of the above-described procedure, starting from **16a-19a**, compounds **16b-19b** were prepared.

***tert*-butyl (3-((4-(hydroxymethyl)-2,7-dimethoxyacridin-9-yl)thio)propyl)carbamate (16b). Yield 68%.** $^1$H NMR (400 MHz, Chloroform-d) δ 8.04 (d, *J* = 9.3 Hz, 1H), 7.92 (d, *J* = 2.8 Hz, 1H), 7.85 (d, *J* = 2.8 Hz, 1H), 7.41 (dd, *J* = 9.3, 2.8 Hz, 1H), 7.27 (d, *J* = 2.7 Hz, 1H), 5.42 (br s, 1H), 5.21 (s, 2H), 4.43 (br s, 1H), 4.04 (s, 5H), 4.01 (s, 5H), 3.19 (d, *J* = 6.7 Hz, 2H), 2.93 (t, *J* = 7.3 Hz, 2H), 1.62 (t, *J* = 7.0 Hz, 3H), 1.39 (s, 9H). 13C NMR (101 MHz, Chloroform-d) δ 158.6, 158.0, 156.0, 143.4, 142.7, 132.0, 131.3, 130.8, 124.2, 122.2, 102.1, 101.6, 65.1, 55.8, 55.8, 39.6， 33.7, 30.8, 28.5. HRMS(ESI) $[M + H]^+$ calculated for $C_{24}H_{31}N_2O_5S$: 459.1948, found: 459.1950.

***tert*-butyl-(S)-3-(((4-(hydroxymethyl)-2,7-dimethoxyacridin-9-yl)thio)methyl)pyrrolidine-1-carboxylate (17b). 68%.** $^1$H NMR (400 MHz, Chloroform-d) δ 8.06 (d, *J* = 9.3 Hz, 1H), 7.92 (d, *J* = 2.6 Hz, 1H), 7.85 (d, *J* = 2.3 Hz, 1H), 7.42 (dd, *J* = 9.3, 2.5 Hz, 1H), 7.29 (s, 1H), 5.36 (br s, 1H), 5.21 (s, 2H), 4.05 (s, 3H), 4.02 (s, 3H), 3.62-3.32 (m, 2H), 3.27 - 2.84 (m, 4H), 2.15-2.05 (d, *J* = 6.4 Hz, 1H), 2.01 - 1.92 (m, 1H), 1.70-1.60 (m, 1H), 1.43 (d, *J* = 4.8 Hz, 9H). $^{13}$C NMR (126 MHz, Chloroform-d) δ 158.6, 158.0, 154.4, 143.4, 142.7, 140.5, 136.5, 131.9, 131.1, 130.5, 124.1, 122.1, 101.9, 101.3, 79.3, 64.9, 55.7, 51.1 (d, *J* = 115 Hz ), 45.1 (d, *J* = 135 Hz), 39.4, 38.4, 31.2, 30.6, 28.5. HRMS(ESI) $[M + H]^+$ calculated for $C_{26}H_{33}N_2O_5S$: 485.2105, found: 485.2110.

***tert*-butyl-(R)-3-(((4-(hydroxymethyl)-2,7-dimethoxyacridin-9-yl)thio)methyl)pyrrolidine-1-carboxylate (18b). 64%.** $^1$H NMR (400 MHz, Chloroform-d) δ 8.05 (d, *J* = 9.4 Hz, 1H), 7.90

(d, $J$ = 2.7 Hz, 1H), 7.83 (d, $J$ = 2.6 Hz, 1H), 7.41 (dd, $J$ = 9.3, 2.7 Hz, 1H), 7.28 (s, 1H), 5.38 (br s, 1H), 5.21 (s, 2H), 4.04 (s, 3H), 4.02 (s, 3H), 3.64 - 3.20 (m, 4H), 3.08 - 2.87 (m, 2H), 2.14 - 2.03 (m, 1H), 1.99-1.90 (m, 1H), 1.68 - 1.57 (m, 1H), 1.42 (d, $J$ = 5.2 Hz, 9H). $^{13}$C NMR (126 MHz, Chloroform-d) δ 158.6, 158.0, 154.4, 143.3, 142.7, 140.5, 136.4, 131.9, 131.1, 130.5, 124.1, 122.0, 101.9, 101.3, 79.3, 64.9, 55.7, 50.9 (d, $J$ = 115 Hz ), 45.1 (d, $J$ = 140 Hz), 39.4, 38.4, 31.2, 30.6, 28.5. HRMS(ESI) [M + H]$^+$ calculated for $C_{26}H_{33}N_2O_5S$: 485.2105, found: 485.2102.

***tert*-butyl-3-(((4-(hydroxymethyl)-2,7-dimethoxyacridin-9-yl)thio)methyl)azetidine-1-carboxylate (19b). 50%.** $^1$H NMR (400 MHz, Chloroform-d) δ 8.08 (d, $J$ = 9.3 Hz, 1H), 7.87 (d, $J$ = 2.7 Hz, 1H), 7.80 (d, $J$ = 2.7 Hz, 1H), 7.43 (dd, $J$ = 9.3, 2.7 Hz, 1H), 7.30 (d, $J$ = 2.6 Hz, 1H), 5.22 (s, 2H), 4.04 (s, 3H), 4.02 (s, 3H), 3.85 (t, $J$ = 8.4 Hz, 2H), 3.59 (dd, $J$ = 8.8, 5.2 Hz, 2H), 3.12 (d, $J$ = 7.9 Hz, 2H), 2.24 (td, $J$ = 7.9, 4.1 Hz, 1H), 1.39 (s, 9H). $^{13}$C NMR (126 MHz, Chloroform-d) δ 158.7, 158.1, 156.2, 143.0, 142.5, 140.5, 135.6, 131.8, 131.2, 130.7, 124.3, 122.3, 101.8, 101.2, 79.6, 64.8, 55.7, 55.7, 39.7, 29.0, 28.3. HRMS(ESI) [M + H]$^+$ calculated for $C_{25}H_{31}N_2O_5S$: 471.1948, found: 471.1939.

**(2,7-dimethoxy-9-((piperidin-4-ylmethyl)thio)acridin-4-yl)methanol (7).** Compound **7b** (24.2mg, 0.048mmol) was dissolved in 2ml of 5% trifluoroacetic acid dichloromethane solution and the mixture was allowed to react at room temperature for 2h. The solvent was then evaporated and the residue was dissolved with methanol, then purified by HPLC/MS on a Waters Auto Purification LC/MS system (ACQUITY UPLC ® BEH C18 17 μm 2.1X50 mm column) to afford **7** (11.2mg, 45%) as a dark red solid. $^1$H NMR (400 MHz, MeOD) δ 8.09 (d, $J$ = 9.4 Hz, 1H), 7.76 (d, $J$ = 2.2 Hz, 1H), 7.71 (d, $J$ = 2.2 Hz, 1H), 7.54 (d, $J$ = 1.1 Hz, 1H), 7.50 (dd, $J$ = 9.4, 2.5 Hz, 1H), 5.23 (s, 2H), 4.04 (s, 3H), 4.03 (s, 3H), 3.33 - 3.30 (m, 2H), 2.95 (d, $J$ = 6.7 Hz, 2H), 2.83 (td, $J$ = 12.8, 1.6 Hz, 2H), 2.07-1.99 (m, 2H), 1.66 - 1.55 (m, 1H), 1.51 - 1.40 (m, 2H). $^{13}$C NMR (126 MHz, Methanol-d4) δ 160.0, 159.8, 144.0, 143.1, 143.0, 137.7, 132.9, 131.7, 131.4, 124.9, 121.8, 102.8, 101.7, 62.2, 56.2, 56.1, 44.8, 42.9, 35.7, 29.4. HRMS(ESI) [M + H]$^+$ calculated for $C_{22}H_{27}N_2O_3S$: 399.1737, found: 399.1732.

**Compounds 16-19**. By employment of the above-described procedure, starting from **16b-19b**, compounds **16-19** were prepared.

**(9-((3-aminopropyl)thio)-2,7-dimethoxyacridin-4-yl)methanol (16). Yield 88%.** $^1$H NMR (400 MHz, MeOD) δ 8.13 (d, $J$ = 9.4 Hz, 1H), 7.94 (d, $J$ = 26.6 Hz, 2H), 7.58 - 7.54 (m, 1H), 7.50 (dd, $J$ = 9.4, 2.7 Hz, 1H), 5.31 (s, 2H), 4.05 (s, 3H), 4.05 (s, 3H), 3.09 (t, $J$ = 7.4 Hz, 2H), 2.95 - 2.88 (t, $J$ = 7.4 Hz, 2H), 1.79 - 1.70 (m, 2H). $^{13}$C NMR (151 MHz, Methanol-d4) δ 160.3,

160.0, 141.4, 140.9, 140.6, 139.8, 132.1, 131.7, 129.9, 127.0, 124.3, 103.2, 102.3, 62.1, 56.4, 56.3, 39.5, 34.1, 29.4.HRMS(ESI) [M + H]$^+$ calculated for $C_{19}H_{23}N_2O_3S$: 359.1424, found: 359.1428.

**(S)-(2,7-dimethoxy-9-((pyrrolidin-3-ylmethyl)thio)acridin-4-yl)methanol (17). Yield 93%.** $^1$H NMR (400 MHz, MeOD) δ 8.11 (d, *J* = 9.4 Hz, 1H), 7.81 (s, 1H), 7.75 (s, 1H), 7.57 (s, 1H), 7.51 (d, *J* = 9.4 Hz, 1H), 5.27 (s, 2H), 4.07 (s, 3H), 4.07 (s, 3H), 3.36 - 3.27 (m, 2H), 3.19 - 3.05 (m, 3H), 2.96 (dd, *J* = 11.4, 8.3 Hz, 1H), 2.25 (hept, *J* = 7.6 Hz, 1H), 2.17 - 2.05 (m, 1H), 1.80 - 1.70 (m, 1H). $^{13}$C NMR (126 MHz, Methanol-d4) δ 160.3, 160.1, 141.8, 141.4, 140.7, 139.9, 131.8, 131.4, 130.8, 126.4, 123.6, 103.0, 102.0, 62.1, 56.3, 56.2, 50.7, 46.2, 40.0, 39.5, 30.9. HRMS(ESI) [M + H]$^+$ calculated for $C_{21}H_{25}N_2O_3S$: 385.1580, found: 385.1580.

**(R)-(2,7-dimethoxy-9-((pyrrolidin-3-ylmethyl)thio)acridin-4-yl)methanol (18). Yield 64%.** $^1$H NMR (400 MHz, MeOD) δ 8.14 (d, *J* = 9.3 Hz, 1H), 7.96 (s, 1H), 7.89 (s, 1H), 7.58 (s, 1H), 7.53 - 7.48 (m, 1H), 5.32 (s, 2H), 4.06 (s, 6H), 3.35 - 3.26 (m, 2H), 3.20 - 3.08 (m, 3H), 2.97 - 2.92 (m, 1H), 2.26 (hept, *J* = 7.6 Hz, 1H), 2.17 - 2.07 (m, 1H), 1.80 - 1.71 (m, 1H). $^{13}$C NMR (126 MHz, Methanol-d4) δ 160.2, 160.0, 141.4, 141.1, 140.3, 140.2, 131.7, 131.3, 130.5, 126.5, 123.7, 103.0, 102.0, 62.1, 56.3, 56.2, 50.6, 46.1, 39.9, 39.6, 30.9. HRMS(ESI) [M + H]$^+$ calculated for $C_{21}H_{25}N_2O_3S$: 385.1580, found: 385.1579.

**(9-((azetidin-3-ylmethyl)thio)-2,7-dimethoxyacridin-4-yl)methanol (19). Yield 56%.** $^1$H NMR (400 MHz, MeOD) δ 8.07 (d, *J* = 9.4 Hz, 1H), 7.80 (d, *J* = 2.2 Hz, 1H), 7.73 (d, *J* = 2.1 Hz, 1H), 7.53 (s, 1H), 7.46 (dd, *J* = 9.3, 2.2 Hz, 1H), 5.28 (s, 2H), 4.04 (s, 3H), 4.03 (s, 3H), 3.82 (t, *J* = 9.7 Hz, 2H), 3.69 - 3.62 (m, 2H), 3.26 (d, *J* = 7.9 Hz, 2H), 2.71 - 2.58 (m, 1H). $^{13}$C NMR (101 MHz, Methanol-d4) δ 160.4, 160.2, 142.4, 142.0, 141.3, 138.0, 132.0, 131.7, 131.5, 126.2, 123.2, 102.7, 101.6, 62.0, 56.3, 56.2, 51.9, 38.9, 33.9. HRMS(ESI) [M + H]$^+$ calculated for $C_{20}H_{23}N_2O_3S$: 371.1424, found: 371.1431.

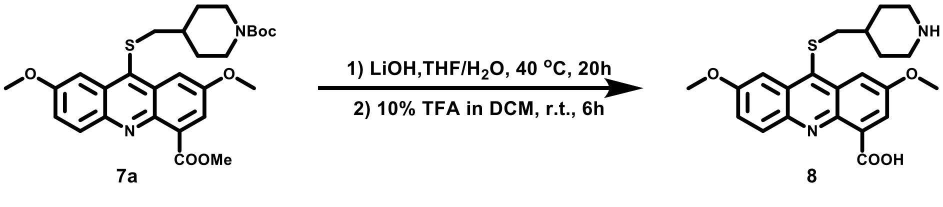


**2,7-dimethoxy-9-((piperidin-4-ylmethyl)thio)acridine-4-carboxylic acid (8).** 0.4 ml of aqueous1 M lithium hydroxide solution of was added into a solution of compound **7a** (42mg, 0.080mmol) 2ml of tetrahydrofuran, and the mixture was reacted at 40 ° C for 20 hours. After cooling to room temperature, 5 ml of a 10% solution of trifluoroacetic acid in dichloromethane was added to the system, and the mixture was reacted at room temperature for 6 h. The solvent was then evaporated and the residue was dissolved with methanol, then purified by

HPLC/MS on a Waters Auto Purification LC/MS system (ACQUITY UPLC ® BEH C18 17 μm 2.1X50 mm column) to afford **8** (21.6 mg，51%) as a dark red foam. $^1$H NMR (600 MHz, MeOD) δ 8.39 (d, *J* = 2.8 Hz, 1H), 8.23 (d, *J* = 2.7 Hz, 1H), 8.12 (d, *J* = 9.3 Hz, 1H), 7.97 (d, *J* = 2.5 Hz, 1H), 7.62 (dd, *J* = 9.3, 2.6 Hz, 1H), 4.09 (s, 3H), 4.07 (s, 3H), 3.36-3.33 (m, 2H), 3.02 (d, *J* = 6.8 Hz, 2H), 2.89 - 2.82 (m, 2H), 2.11 (d, *J* = 14.0 Hz, 2H), 1.70 - 1.64 (m, 1H), 1.50 - 1.41 (m, 2H). $^{13}$C NMR (126 MHz, Methanol-d4) δ 168.7, 160.7, 158.9, 141.9, 141.4, 140.9, 131.6, 131.5, 130.7, 130.5, 128.0, 126.7, 108.6, 103.1, 56.6, 56.5, 44.8, 43.4, 35.7, 29.4. HRMS(ESI) [M + H]$^+$ calculated for $C_{22}H_{25}N_2O_4S$: 413.1530, found: 413.1526.

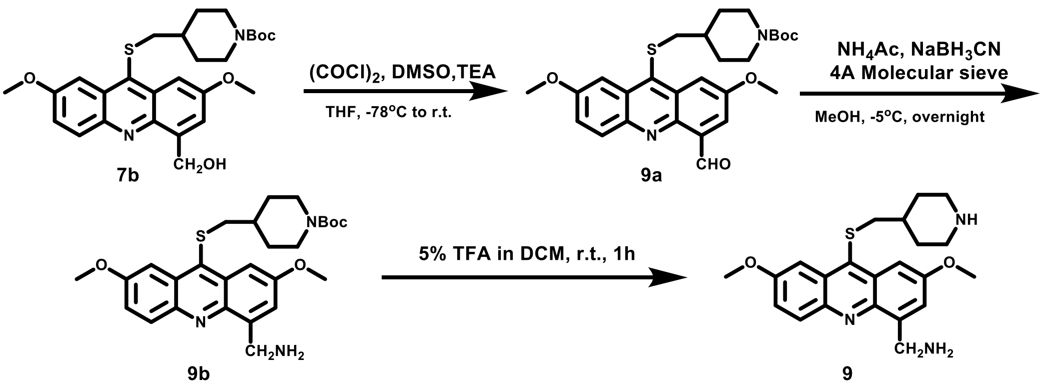


***tert*-butyl 4-(((4-formyl-2,7-dimethoxyacridin-9-yl)thio)methyl)piperidine-1-carboxylate (9a).** 20.4 μl of oxalyl chloride was dissolved in 1 ml of dry tetrahydrofuran and the mixture was cooled to -78 ° C. 25.6 μl of dimethyl sulfoxide was slowly added to the system, the reaction was stirred at -78 ° C for 30 min, followed by addition of a solution of compound **7b** (60 mg, 0.12 mmol) in 1 ml of tetrahydrofuran solution, and the reaction was continued at -78 ° C for 1 h. 0.10 ml of triethylamine was then added, and the reaction was allowed to return to room temperature for 2 h. The solvent was then evaporated and the residue was purified by chromatography on a silica gel column (dichloromethane / ethyl acetate = 95/5) to gave compound **9a** (56mg，94%) as a light yellow solid. $^1$H NMR (400 MHz, CDCl3) δ 11.54 (s, 1H), 8.19 (d, *J* = 2.9 Hz, 1H), 8.09 (d, *J* = 9.3 Hz, 1H), 7.98 (d, *J* = 2.9 Hz, 1H), 7.86 (d, *J* = 2.6 Hz, 1H), 7.44 (dd, *J* = 9.4, 2.7 Hz, 1H), 4.10-3.98 (m, 2H), 4.04 (s, 3H), 4.03 (s, 3H), 2.80 (d, *J* = 6.8 Hz, 2H), 2.58 (t, *J* = 12.2 Hz, 2H), 1.83 (m, 2H), 1.52 - 1.44 (m, 1H), 1.43 (s, 9H), 1.23-1.08 (m, 2H). $^{13}$C NMR (101 MHz, Chloroform-d) δ 192.8, 158.9, 157.6, 154.8, 144.5, 142.9, 137.3, 133.5, 132.5, 130.7, 130.7, 125.0, 123.0, 109.0, 101.9, 79.7, 56.0, 55.8, 43.8 (br s), 43.2, 36.9, 31.8, 28.5. HRMS(ESI) [M + H]$^+$ calculated for $C_{27}H_{33}N_2O_5S$: 497.2105, found: 497.2101.

***tert*-butyl-4-(((4-(aminomethyl)-2,7-dimethoxyacridin-9-yl)thio)methyl)piperidine-1-carboxylate (9b).** Under argon atmosphere, compound **9a** (55 mg, 0.11 mmol), ammonium acetate (85 mg, 1.11 mmol), sodium cyanoborohydride (6.9 mg, 0.11 mmol), 4 mg of 4A molecular sieves was dissolved in 3 ml of dry methanol at -5 °C, then the reaction was stilled at the same temperature overnight. After the removal of solid by filtration, the solvent was diluted with methylene chloride, washed with brine, and then evaporated under vacuum. The residue was purified by chromatography on a silica gel column (dichloromethane /methanol = 95/5) to gave compound **9b** (25 mg, 46%) as a light yellow solid. $^{1}H$ NMR (400 MHz, $CDCl_3$) δ 7.54 (d, *J* = 2.6 Hz, 1H), 7.52 (d, *J* = 2.5 Hz, 1H), 7.30 (d, *J* = 9.3 Hz, 1H), 7.08 (d, *J* = 2.3 Hz, 1H), 6.90 (dd, *J* = 9.3, 2.6 Hz, 1H), 4.86 (s, 2H), 4.06 - 3.95 (m, 2H),4.04 (s, 3H), 3.96 (s, 3H), 2.62 (d, *J* = 6.7 Hz, 2H), 2.52 (t, *J* = 12.0 Hz, 2H), 1.85-1.70 (m, 2H), 1.41 (s, 9H), 1.35-1.30 (m, 1H), 1.18-1.04 (m, 2H). $^{13}C$ NMR (151 MHz, Chloroform-d) δ 158.7, 157.2, 154.8, 142.5, 140.9, 137.2, 131.2 (overlapped), 130.0, 129.6, 126.0, 124.0, 103.2, 101.5, 79.7, 55.7 (overlapped), 50.4, 43.6 (br s), 43.1, 36.8, 31.8, 28.5. HRMS(ESI) $[M + H]^+$ calculated for $C_{27}H_{36}N_3O_4S$: 498.2421, found: 498.2420.

**(2,7-dimethoxy-9-((piperidin-4-ylmethyl)thio)acridin-4-yl)methanamine (9).** Compound **9b** (13.5mg，0.027mmol) was dissolved in 2ml of 5% trifluoroacetic acid dichloromethane solution and the mixture was allowed to react at room temperature for 2h. The solvent was then evaporated and the residue was dissolved with methanol, then purified by HPLC/MS on a Waters Auto Purification LC/MS system (ACQUITY UPLC ® BEH C18 17 μm 2.1X50 mm column) to afford **9** (6.3mg, 45%) as a dark red foam. $^{1}H$ NMR (400 MHz, MeOD) δ 7.55 (dd, *J* = 18.2, 2.6 Hz, 2H), 7.47 (d, *J* = 9.3 Hz, 1H), 7.17 (d, *J* = 2.5 Hz, 1H), 6.95 (dd, *J* = 9.3, 2.7 Hz, 1H), 4.14 - 4.07 (m, 3H), 4.02 (s, 3H), 2.86 - 2.72 (m, 4H), 2.07 - 1.98 (m, 3H), 1.57 - 1.31 (m, 4H). $^{13}C$ NMR (151 MHz, Methanol-d4) δ 160.2, 158.7, 143.5, 141.6, 137.9, 132.0, 131.7, 130.9, 130.4, 127.6, 125.2, 104.1, 102.5, 56.3, 56.2, 50.7, 44.8, 42.9, 35.6, 29.4. HRMS(ESI) $[M + H]^+$ calculated for $C_{22}H_{28}N_3O_2S$: 398.1897, found: 398.1896.

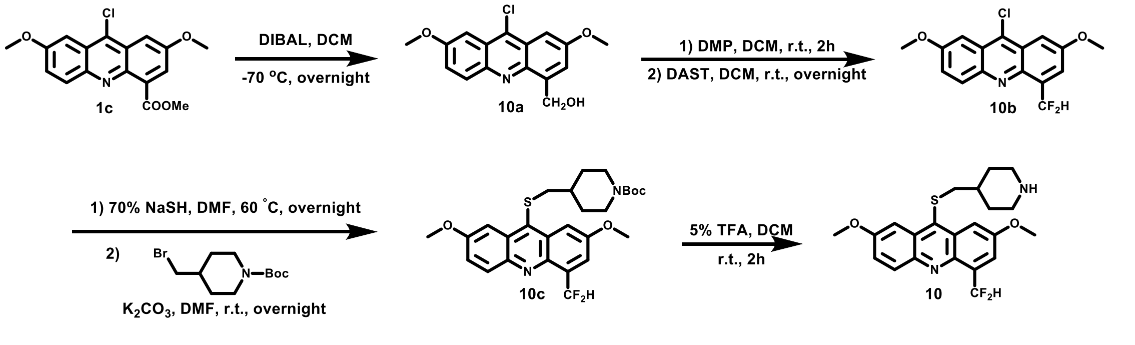


**(9-chloro-2,7-dimethoxyacridin-4-yl)methanol (10a).** 0.28ml of 1.0M DIBAL-H solution in toluene was added to a solution of **1c** (40mg，0.12mmol) in 5 ml of dry dichloromethane at -70 °C, then the reaction was stilled at the same temperature overnight. The reaction was quenched by adding saturated potassium hydrogen tartrate solution, diluted with dichloromethane, washed twice with brine, dried over anhydrous $Na_2SO_4$ and concentrated *in vacuo*. The residue was purified by chromatography on a silica gel column (dichloromethane / ethyl acetate = 90/10) to gave compound **10a** (180mg, 55%) as a light yellow solid. $^1H$ NMR (400 MHz, CDCl3) δ 8.00 (d, *J* = 9.3 Hz, 1H), 7.44 - 7.34 (m, 3H), 7.27 (s, 1H), 5.28 (br s, 1H), 5.19 (s, 2H), 4.02 (s, 3H), 4.00 (s, 3H). $^{13}C$ NMR (101 MHz, Chloroform-d) δ 158.6, 158.0, 143.4, 142.9, 140.1, 136.3, 131.5, 126.0, 125.6, 124.7, 122.5, 99.8, 99.2, 64.8, 55.8, 55.8. HRMS(ESI) $[M + H]^+$ calculated for $C_{16}H_{15}ClNO_3$: 304.0735, found: 304.0732.

**9-chloro-4-(difluoromethyl)-2,7-dimethoxyacridine (10b).** To a solution of compound **10a** (30mg，0.10mmol) in 3ml of dichloromethane, Dess-Martin oxidant (60 mg, 0.14 mmol) was added and the reaction was allowed to stir at room temperature for 2h. The solution was diluted with dichloromethane, washed twice with brine, dried over anhydrous $Na_2SO_4$ and concentrated *in vacuo*. Under argon atmosphere, the crude product was dissolved in 2 ml of dry dichloromethane, 0.1 ml of diethylaminosulfur trifluoride was added, and the mixture was reacted at room temperature overnight. The reaction was quenched by sodium bicarbonate aqueous solution, then diluted with dichloromethane, washed twice with brine, dried over anhydrous $Na_2SO_4$ and concentrated *in vacuo*. The residue was purified by chromatography on a silica gel column (petroleum ether / ethyl acetate = 5/1) to afford **10b** (7.4mg, 23%) as white solid. $^1H$ NMR (400 MHz, $CDCl_3$) δ 8.08 (d, *J* = 9.3 Hz, 1H), 7.91 (t, *J* = 55.4 Hz, 1H), 7.76 - 7.73 (m, 1H), 7.59 (d, *J* = 2.6 Hz, 1H), 7.48 - 7.41 (m, 2H), 4.05 (s, 3H), 4.03 (s, 3H). $^{13}C$ NMR (151 MHz, Chloroform-d) δ 158.9, 157.6, 144.3, 141.2 (t, *J* = 19Hz), 135.9, 133.8 (t, *J* = 85 Hz), 132.2, 125.9, 125.6, 125.0, 121.8 (t, *J* = 27 Hz), 112.1 (t, *J* = 942 Hz), 102.4, 99.7, 56.0, 55.9. HRMS(ESI) $[M + H]^+$ calculated for $C_{16}H_{13}ClF_2NO_2$: 324.0597, found: 324.0597.

***tert*-butyl 4-(((4-(difluoromethyl)-2,7-dimethoxyacridin-9-yl)thio)methyl)piperidine-1-carboxylate (10c).** To a solution of **10b** (7.4mg，0.023mmol) in 2 ml of anhydrous DMF, sodium hydrogen hydride hydrate powder (70%, 1.9mg，0.034mmol) was added under argon atmosphere, and the reaction was stirred at 60 °C overnight. 1-BOC-4-bromomethylpiperidine (12.7mg，0.0461mmoll) and potassium carbonate (15.8mg，0.114mmol) were added into the slurry and the reaction was allowed to react at room temperature overnight. The solvent was then evaporated and the residue was dissolved with dichloromethane and washed with water. The combined organic extracts were dried over anhydrous $Na_2SO_4$ and concentrated *in vacuo*. The residue was purified by flash chromatography on a silica gel column (Petroleum ether / ethyl acetate = 3/1) to give compound **10c** as a yellow solid (5.3mg, 45%) and used for next step.

**4-(difluoromethyl)-2,7-dimethoxy-9-((piperidin-4-ylmethyl)thio)acridine (10).** Compound **10c** (5.3mg，0.010mmol) was dissolved in 2ml of 5% trifluoroacetic acid dichloromethane solution and the mixture was allowed to react at room temperature for 2h. The solvent was then evaporated and the residue was dissolved with methanol, then purified by HPLC/MS on a Waters Auto Purification LC/MS system (ACQUITY UPLC ® BEH C18 17 μm 2.1X50 mm column) to afford **10** (3.8mg，70%) as a yellow solid. $^{1}$H NMR (400 MHz, MeOD) δ 8.08 (d, $J$ = 6.3 Hz, 1H), 7.93 (d, $J$ = 2.8 Hz, 1H), 7.89（t, $J$ = 59.6Hz, 1H）7.89 (d, $J$ = 4.1 Hz, 1H), 7.68 (dd, $J$ = 2.8, 1.4 Hz, 1H), 7.47 (dd, $J$ = 9.4, 2.8 Hz, 1H), 4.09-4.05(overlapped, m, 2H), 4.06 (s, 3H), 4.04 (s, 3H), 2.96 (d, $J$ = 6.7 Hz, 2H), 2.90 - 2.78 (m, 2H), 2.11 (d, $J$ = 14.4 Hz, 2H), 1.68 - 58 (m, 1H)，1.52 - 1.43 (m, 2H). $^{13}$C NMR (151 MHz, Methanol-d4) δ 160.4, 158.9, 145.1, 142.0, 137.7, 135.5 (t, $J$ = 85 Hz), 133.4, 131.7, 131.4, 125.6, 122.0 (t, $J$ = 27 Hz), 113.3 (t, $J$ = 937 Hz), 105.3, 102.7, 56.4, 56.2, 44.8, 43.0, 35.7, 29.5. HRMS(ESI) [M + H]$^+$ calculated for $C_{22}H_{25}F_2N_2O_2S$: 419.1599, found: 419.1587.

General procedures of synthesizing **1c'/1d;** To a suspension of compound **1b** (1.65 g, 6.05 mmol) in 200 ml of dry dichloromethane was slowly added 30 ml of boron tribromide, and the mixture was reacted at 0 ° C for 2 h. Methanol was added to the reaction system to quench the reaction, and the solvent was removed under reduced pressure. The crude product was separated and purified with a silica gel column (dichloromethane / methanol = 92/8) to obtain **1c'** (0.22 g, 14%) and the reported compound **1d** (0.80 g, 56%), with 0.70 g of the starting material recovered. **1c':** $^{1}$H NMR (400 MHz, MeOD) δ 8.08 (dd, $J$ = 9.4, 4.8 Hz, 2H), 7.64 (dd, $J$ = 7.6, 4.4 Hz, 4H), 4.07 (s, 3H). $^{13}$C NMR (151 MHz, Methanol-d4) δ 160.8, 159.2, 129.2, 127.8, 127.2, 104.6, 101.3, 56.6. HRMS(ESI) [M + H]$^+$ calculated for $C_{14}H_{11}ClNO_2$: 260.0473, found: 260.0471.

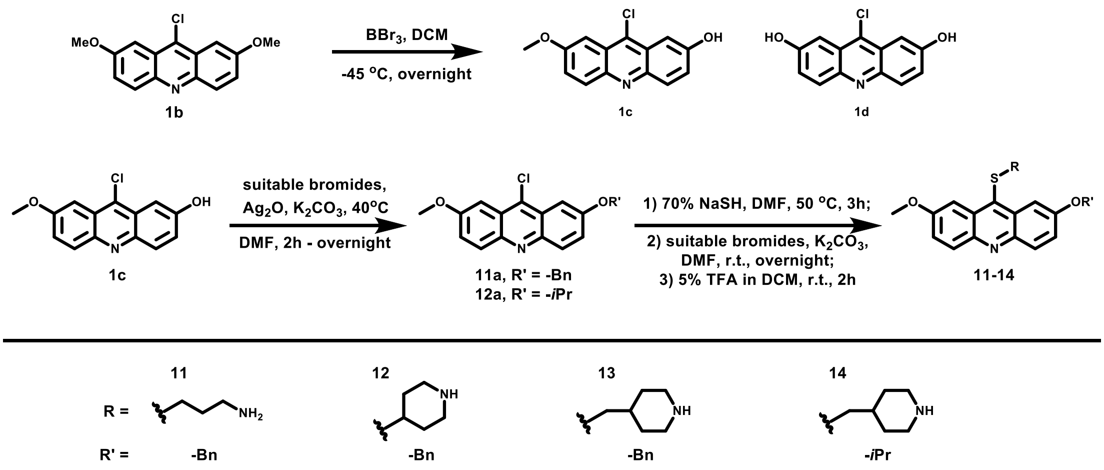

General procedures of synthesizing **11a**,**12a**: To a solution of **1c'** (94.5mg, 0.364mmol, 100 mol%) in 10 ml of anhydrous DMF, potassium carbonate (75.4mg, 0.546mmol, 150 mol%) silver oxide (126.5mg, 0.546mmol, 150 mol%) and suitable bromides (0.546mmol, 150 mol%) was added. The reaction was allowed to react at 40 °C until full conversion. The reaction solution was spin-dried under reduced pressure, and water / dichloromethane was separated. The solvent was then evaporated and the residue was dissolved with dichloromethane and washed with water. The combined organic extracts were dried over anhydrous $Na_2SO_4$ and concentrated *in vacuo*. The residue was purified by chromatography on a silica gel column (Petroleum ether / ethyl acetate = 9/1) to give the desired compound.

**2-(benzyloxy)-9-chloro-7-methoxyacridine (11a). Yield 100%.** $^1$H NMR (400 MHz, CDCl3) δ 8.08 (dd, *J* = 9.3, 7.6 Hz, 2H), 7.60 (d, *J* = 2.6 Hz, 1H), 7.56 - 7.35 (m, 8H), 5.27 (s, 2H), 4.02 (s, 3H). $^{13}$C NMR (101 MHz, Chloroform-d) δ 158.4, 157.5, 144.4, 144.3, 136.2, 135.7, 131.6, 131.5, 128.3, 127.9, 128.0, 125.6, 125.6, 124.7, 124.6, 101.2, 99.9, 70.6, 55.8. HRMS(ESI) $[M + H]^+$ calculated for $C_{21}H_{17}ClNO_2$: 350.0942, found: 350.0941.

**9-chloro-2-isopropoxy-7-methoxyacridine (12a). Yield 78%.** $^1$H NMR (400 MHz, Chloroform-d) δ 8.08 (dd, *J* = 9.4, 2.2 Hz, 2H), 7.51 (dd, *J* = 13.1, 2.7 Hz, 2H), 7.41 (ddd, *J* = 9.3, 7.5, 2.7 Hz, 2H), 4.85 (p, *J* = 6.0 Hz, 1H), 4.03 (s, 3H), 1.48 (d, *J* = 6.0 Hz, 6H). $^{13}$C NMR (151 MHz, Chloroform-d) δ 158.5, 156.7, 144.4, 144.3, 135.5, 131.7, 131.7, 125.8, 125.6, 125.5, 124.5, 101.8, 99.9, 70.5, 55.8, 22.0. HRMS(ESI) $[M + H]^+$ calculated for $C_{17}H_{17}ClNO_2$: 302.0942, found: 302.0942.

General procedures of synthesizing **11-14**: To a solution of suitable acridine chloride (100 mol%) in 2 ml of anhydrous DMF, sodium hydrogen hydride hydrate powder (150 mol%) was added under argon atmosphere, and the reaction was stirred at 50 °C for 3h. Suitable bromides (200 mol%) and potassium carbonate (500 mol%) were added into the slurry and the reaction was allowed to react at room temperature overnight. The solvent was then evaporated and the residue was dissolved with dichloromethane and washed with water. The combined organic extracts were dried over anhydrous $Na_2SO_4$ and concentrated *in vacuo*. The residue was dissolved in 2ml of 5% trifluoroacetic acid dichloromethane solution and the mixture was allowed to react at room temperature for 2h. The solvent was then evaporated and the residue was dissolved with methanol, then purified by HPLC/MS on a Waters Auto Purification LC/MS system (ACQUITY UPLC ® BEH C18 17 μm 2.1X50 mm column) to afford desired product.

**3-((2-(benzyloxy)-7-methoxyacridin-9-yl)thio)propan-1-amine (11). Yield 9.6%.** $^{1}$H NMR (400 MHz, MeOD) δ 8.13 (dd, *J* = 12.1, 9.4 Hz, 2H), 8.01 (dd, *J* = 8.5, 2.6 Hz, 2H), 7.81 (dd, J = 9.4, 2.7 Hz, 1H), 7.72 (dd, J = 9.4, 2.7 Hz, 1H), 7.60-7.34 (m, 5H), 5.42 (s, 2H), 4.08 (s, 3H), 3.00 (t, *J* = 7.6Hz, 2H), 2.85(t, *J* = 7.6Hz, 2H), 1.70 (quint, *J* = 7.6Hz, 2H). $^{13}$C NMR (151 MHz, Methanol-d4) δ 160.9, 159.6, 146.7, 138.5, 138.4, 137.6, 132.4, 132.0, 130.0, 129.9, 129.8, 129.4, 128.7, 126.4, 126.2, 105.7, 103.8, 71.8, 56.7, 39.4, 34.6, 29.4. HRMS(ESI) [M + H]$^+$ calculated for $C_{24}H_{25}N_2O_2S$: 405.1631, found: 405.1630.

**2-(benzyloxy)-7-methoxy-9-(piperidin-4-ylthio)acridine (12). 24%.** $^{1}$H NMR (400 MHz, MeOD) δ 7.99 (dd, *J* = 14.3, 9.4 Hz, 2H), 7.89 (d, *J* = 2.4 Hz, 2H), 7.58-7.50 (m, 3H), 7.47-7.40 (m, 3H), 7.39-7.33 (m, 1H), 5.36 (s, 2H), 4.01 (s, 3H), 3.26 (d, *J* = 13.1 Hz, 2H), 3.19 - 3.10 (m, 1H), 2.81 (t, *J* = 10.8 Hz, 2H), 1.86 (d, *J* = 11.5 Hz, 2H), 1.70 (td, *J* = 14.4, 3.7 Hz, 2H). $^{13}$C NMR (101 MHz, Methanol-d4) δ 159.9, 158.7, 144.7, 144.6, 138.3, 135.9, 132.6, 132.2, 131.8, 131.7, 129.8, 129.1, 128.3, 125.8, 125.6, 105.0, 103.3, 71.2, 56.2, 45.5, 45.1, 32.0. HRMS(ESI) [M + H]$^+$ calculated for $C_{26}H_{27}N_2O_2S$: 431.1788, found: 431.1781.

**2-(benzyloxy)-7-methoxy-9-((piperidin-4-ylmethyl)thio)acridine (13). 19%.** $^{1}$H NMR (400 MHz, MeOD) δ 8.01 - 7.89 (m, 2H), 7.86-7.75 (m, 2H), 7.58 - 7.31 (m, 7H), 5.32 (s, 2H), 3.99 (s, 3H), 3.27 (d, *J* = 13.1Hz, 2H), 2.75 (t, *J* = 12.5 Hz, 2H), 2.68 (d, *J* = 6.4 Hz, 2H), 1.94 (d, *J* = 13.2 Hz, 2H), 1.44 - 1.28 (m, 3H). $^{13}$C NMR (151 MHz, Methanol-d4) δ 160.0, 158.7, 144.7, 144.7, 138.6, 138.3, 132.0, 131.9, 131.7, 131.3, 129.8, 129.2, 125.8, 125.6, 104.9, 103.0, 71.4, 56.2, 45.1, 43.0, 36.0, 30.0. HRMS(ESI) [M + H]$^+$ calculated for $C_{27}H_{29}N_2O_2S$: 445.1944, found: 445.1945.

**2-isopropoxy-7-methoxy-9-((piperidin-4-ylmethyl)thio)acridine (14). 26%.** $^{1}$H NMR (400 MHz, MeOD) δ 8.15 (dd, *J* = 9.4, 3.6 Hz, 2H), 8.06 (t, *J* = 3.0 Hz, 2H), 7.71 (td, *J* = 9.3, 2.6 Hz, 2H), 4.98 - 4.92 (m, 1H), 4.09 (s, 3H), 3.36 (d, *J* = 12.9 Hz, 2H), 3.11 (d, *J* = 6.8 Hz, 2H), 2.88 (t, *J* = 11.9 Hz, 2H), 2.12 (d, *J* = 14.0 Hz, 2H), 1.79 - 1.69 (m, 1H), 1.54-1.41 (overlapped, m, 2H), 1.51 (s, 3H), 1.49 (s, 3H). $^{13}$C NMR (151 MHz, Methanol-d4) δ 160.8, 158.9, 139.5, 139.3, 132.1, 129.9, 129.0, 127.4, 127.3, 105.6, 103.7, 72.2, 56.6, 44.8, 43.8, 36.0, 29.4, 22.0. HRMS(ESI) [M + H]$^+$ calculated for $C_{23}H_{29}N_2O_2S$: 397.1944, found: 397.1943.

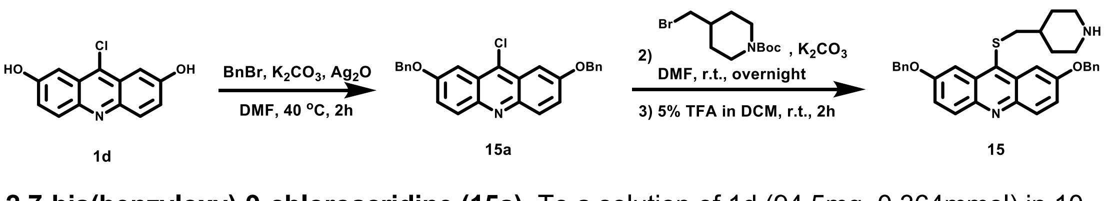


**2,7-bis(benzyloxy)-9-chloroacridine (15a).** To a solution of 1d (94.5mg, 0.364mmol) in 10 ml of anhydrous DMF, potassium carbonate (150.8mg, 1.092mmol) silver oxide (253.0mg,

1.092mmol) and benzyl bromide (100μl) was added. The reaction was allowed to react at 40 °C until full conversion. The reaction solution was spin-dried under reduced pressure, and water / dichloromethane was separated. The solvent was then evaporated and the residue was dissolved with dichloromethane and washed with water. The combined organic extracts were dried over anhydrous $Na_2SO_4$ and concentrated in vacuo. The residue was purified by chromatography on a silica gel column (Petroleum ether / ethyl acetate = 9/1) to give 15a (163.9mg, quant.) as a yellow solid. $^1H$ NMR (400 MHz, Chloroform-d) δ 8.10 (d, J = 9.4 Hz, 2H), 7.62 (d, J = 2.7 Hz, 2H), 7.57 - 7.36 (m, 12H), 5.28 (s, 4H). $^{13}C$ NMR (101 MHz, Chloroform-d) δ 157.6, 144.5, 136.3, 135.9, 131.8, 128.9, 128.5, 128.0, 125.6, 124.8, 101.2, 70.6. HRMS(ESI) $[M + H]^+$ calculated for $C_{27}H_{21}ClNO_2$: 426.1255, found: 426.1255.

**2,7-bis(benzyloxy)-9-((piperidin-4-ylmethyl)thio)acridine (15).** To a solution of **15a** (20mg, 0.047mmol) in 2 ml of anhydrous DMF, sodium hydrogen hydride hydrate powder (70%, 5.6mg，0.070mmol) was added under argon atmosphere, and the reaction was stirred at 50 °C for 3h. *tert*-butyl 4-(bromomethyl)piperidine-1-carboxylate (26.1mg, 0.094mmol) and potassium carbonate (32.4mg, 0.235mol) were added into the slurry and the reaction was allowed to react at room temperature overnight. The solvent was then evaporated and the residue was dissolved with dichloromethane and washed with water. The combined organic extracts were dried over anhydrous $Na_2SO_4$ and concentrated *in vacuo*. The residue was dissolved in 2ml of 5% trifluoroacetic acid dichloromethane solution and the mixture was allowed to react at room temperature for 2h. The solvent was then evaporated and the residue was dissolved with methanol, then purified by HPLC/MS on a Waters Auto Purification LC/MS system (ACQUITY UPLC ® BEH C18 17 μm 2.1X50 mm column) to afford **15** (10.8mg, 36%). $^1H$ NMR (400 MHz, MeOD) δ 8.09 (d, *J* = 9.4 Hz, 2H), 7.85 (d, *J* = 2.5 Hz, 2H), 7.73 (dd, *J* = 9.4, 2.6 Hz, 2H), 7.43 (ddd, *J* = 28.6, 28.0, 7.3 Hz, 10H), 5.37 (s, 4H), 3.22 (d, *J* = 12.8 Hz, 2H), 2.73 - 2.62 (m, 4H), 1.75 (d, *J* = 13.6 Hz, 2H), 1.41 - 1.31 (m, 1H), 1.30 - 1.16 (m, 2H). $^{13}C$ NMR (101 MHz, Methanol-d4) δ 159.4, 137.6, 137.6, 131.5, 130.3, 129.9, 129.4, 128.5, 125.8, 105.5, 71.8, 44.6, 43.6, 35.8, 29.1. HRMS(ESI) $[M + H]^+$ calculated for $C_{33}H_{33}N_2O_2S$: 521.2257, found: 521.2258.

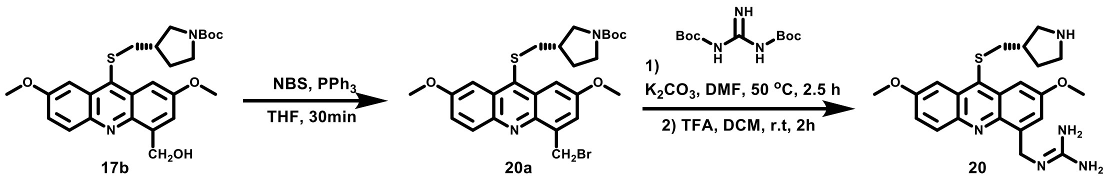


***t*ert-butyl-(S)-3-(((4-(bromomethyl)-2,7-dimethoxyacridin-9-yl)thio)methyl)pyrrolidine-1-carboxylate (20a). 17b** (13mg, 0.027mmol), Triphenylphosphine (10.6mg, 0.040mmol), Nbromosuccinimide (7.2mg, 0.040mmol) was dissolved in 1ml of anhydrous tetrahydrofuran and the reaction was stirred at room temperature for 30min. The reaction was concentrated *in vacuo* and the residue was purified by chromatography on a silica gel column (pure dichloromethane) to give compound **20a** (9.6mg, 65%) as a yellow solid. $^1$H NMR (400 MHz, $CDCl_3$) δ 8.16 (d, *J* = 9.3 Hz, 1H), 7.94 (s, 1H), 7.91 (s, 1H), 7.59 (s, 1H), 7.43 (d, *J* = 9.2 Hz, 1H), 5.31 (d, *J* = 8.9 Hz, 2H), 4.05 (s, 3H), 4.03 (s, 3H), 3.64 - 3.33 (m, 2H), 3.30 - 2.84 (m, 4H), 2.12 (br s, 1H), 2.01 - 1.92 (m, 1H), 1.67-1.60 (m, 1H), 1.43 (s, 9H). $^{13}$C NMR (101 MHz, Chloroform-d) δ 158.8, 157.8, 154.6, 143.8, 141.7, 138.8, 135.8, 132.9, 130.8, 130.7, 124.7, 124.1, 102.9, 101.8, 79.4, 55.8, 51.0 (d, *J* = 108Hz ), 50.9, 45.2 (d, *J* = 112Hz), 39.5, 31.4, 30.7, 29.7, 28.6. HRMS(ESI) $[M + H]^+$ calculated for $C_{26}H_{32}BrN_2O_4S$: 547.1261, found: 547.1255.

**(S)-2-((2,7-dimethoxy-9-((pyrrolidin-3-ylmethyl)thio)acridin-4-yl)methyl)guanidine (20).** Under argon atmosphere, N, N'-di-Boc-guanidine (3.1 mg, 12.1 μmol) and potassium carbonate (1.7 mg, 12.1 μmol) was added into the solution of **20a** (5.5 mg, 10.0 μmol) in 1 ml of anhydrous DMF. The reaction was performed at 50 °C for 2.5 h. The reaction was concentrated *in vacuo* and the residue was dissolved in 4 ml of dichloromethane containing 5% trifluoroacetic acid and reacted at room temperature for 2 h. The solvent was then evaporated and the residue was dissolved with methanol, then purified by HPLC/MS on a Waters Auto Purification LC/MS system (ACQUITY UPLC ® BEH C18 17 μm 2.1X50 mm column) to give compound **20** (2.0mg, 27%). $^1$H NMR (400 MHz, MeOD) δ 8.11 (d, *J* = 9.4 Hz, 1H), 7.96 (d, *J* = 2.7 Hz, 1H), 7.94 (d, *J* = 2.7 Hz, 1H), 7.52 - 7.48 (m, 2H), 5.04 (s, 2H), 4.07 (s, 3H), 4.06 (s, 3H), 3.39 - 3.32 (m, 2H), 3.19 - 2.98 (m, 4H), 2.28 (hept, *J* = 7.6 Hz, 1H), 2.20 - 2.09 (m, 1H), 1.83 - 1.73 (m, 1H). $^{13}$C NMR (151 MHz, Methanol-d4) δ 160.5, 159.6, 159.0, 144.3, 142.9, 138.5, 137.4, 132.8, 132.1, 131.8, 125.7, 125.1, 103.2, 102.8, 56.3, 50.8, 46.2, 43.1, 40.0, 39.4, 31.0. HRMS(ESI) $[M + H]^+$ calculated for $C_{22}H_{28}N_5O_2S$: 426.1958, found: 426.1946.

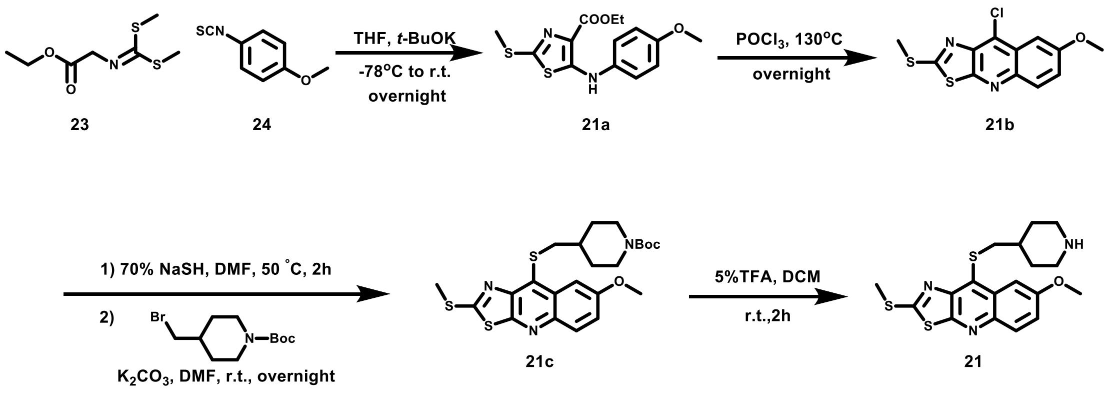

**ethyl 5-((4-methoxyphenyl)amino)-2-(methylthio)thiazole-4-carboxylate (21a).** Under argon atmosphere, a solution of a known compound 23 (1.04 g, 5.02 mmol) in 4 ml of anhydrous tetrahydrofuran was added dropwise to a solution of potassium tert-butoxide (0.79 g, 7.02 mmol) in THF at -78 °C, followed by a solution of compound 24 (0.83 g, 5.02 mmol) in 4 ml of anhydrous tetrahydrofuran. The reaction was performed at -78 °C for 0.5 h and then slowly warmed up to room temperature and reacted overnight. The reaction was quenched by adding saturated ammonium chloride, diluted with dichloromethane, washed twice with brine, dried over anhydrous $Na_2SO_4$ and concentrated *in vacuo*. The residue was purified by chromatography on a silica gel column (pure dichloromethane) to give compound **21a** (0.29 g, 18%) as a light yellow solid. $^1$H NMR (400 MHz, CDCl3) δ 9.39 (s, 1H), 7.17 - 7.11 (m, 2H), 6.91 - 6.83 (m, 2H), 4.38 (q, *J* = 7.1 Hz, 2H), 3.77 (s, 3H), 2.57 (s, 3H), 1.39(t, *J* = 7.1 Hz, 3H). $^{13}$C NMR (101 MHz, Chloroform-d) δ 164.7, 158.5, 156.8, 145.4, 134.2, 121.9, 121.5, 114.9, 60.8, 55.5, 17.7, 14.6. HRMS(ESI) $[M + H]^+$ calculated for $C_{14}H_{17}N_2O_3S_2$: 325.0675, found: 325.0673.

**9-chloro-7-methoxy-2-(methylthio)thiazolo[5,4-b]quinolone (21b).** Compound **21a** (198mg, 0.61mmol) was added in a sealed tube, and 5 ml of phosphorus oxychloride was added under argon atmosphere. The reaction was heated at 130 °C overnight. The resulting slurry was poured onto ice with vigorous stirring to make full quenching. The mixture was diluted with dichloromethane, washed twice with brine, dried over anhydrous $Na_2SO_4$ and concentrated *in vacuo*. The residue was purified by chromatography on a silica gel column (pure dichloromethane) to give compound **21b** (74mg, 41%) as a light yellow solid. $^1$H NMR (400 MHz, CDCl3) δ 7.97 (d, *J* = 9.2 Hz, 1H), 7.55 (d, *J* = 2.8 Hz, 1H), 7.40 (dd, *J* = 9.2, 2.8 Hz, 1H), 4.01 (s, 3H), 2.88 (s, 3H). $^{13}$C NMR (101 MHz, Chloroform-d) δ 172.1, 158.5, 157.6, 142.6, 142.4, 130.2, 129.5, 126.2, 123.1, 101.8, 55.8, 15.5. HRMS(ESI) $[M + H]^+$ calculated for $C_{12}H_{10}ClN_2OS_2$: 296.9918, found: 296.9917.

**tert-butyl-4-(((7-methoxy-2-(methylthio)thiazolo[5,4-b]quinolin-9-yl)thio)methyl)piperidine-1-carboxylate (21c).** To a solution of **21b** (0.020g, 0.067mmo) in 5 ml of anhydrous DMF, sodium hydrogen hydride hydrate powder (70%, 10.8mg, 0.135mmol) was added under argon atmosphere and the reaction was stirred at 50 °C for 2 h until full conversion. 1-BOC-4-bromomethylpiperidine (37.5mg, 0.135mmol) and potassium carbonate (27.9mg, 0.202mmol) were added into the slurry and the reaction was allowed to react at room temperature overnight. The solvent was then evaporated and the residue was dissolved with dichloromethane and washed with water. The combined organic extracts were dried over anhydrous $Na_2SO_4$ and concentrated *in vacuo*. The residue was purified by chromatography on a silica gel column (dichloromethane / ethyl acetate = 95/5) to give compound **21c** (14.9mg, 45%) as a light yellow solid. $^1$H NMR (400 MHz,$CDCl_3$) δ 7.94 (d, *J* = 9.2 Hz, 1H), 7.85 (d, *J* = 2.8 Hz, 1H), 7.38 (dd, *J* = 9.2, 2.8 Hz, 1H), 4.12 - 4.01 (m, 2H), 4.00 (s, 3H), 3.47 (d, *J* = 6.9 Hz, 2H), 2.84 (s, 3H), 2.61 (t, *J* = 12.7 Hz, 2H), 1.86 (d, *J* = 13.1 Hz, 2H), 1.65 - 1.57 (m, 1H), 1.44 (s, 9H), 1.23 - 1.14 (m, 2H). $^{13}$C NMR (101 MHz, Chloroform-d) δ 169.9, 158.1, 158.0, 154.9, 145.1, 141.7, 132.9, 130.3, 128.7, 122.4, 103.5, 79.6, 55.8, 43.9 (br s), 41.1, 36.9, 31.7, 28.6, 15.5. HRMS(ESI) $[M + H]^+$ calculated for $C_{23}H_{30}N_3O_3S_3$: 492.1444, found: 492.1435.

**7-methoxy-2-(methylthio)-9-((piperidin-4-ylmethyl)thio)thiazolo[5,4-b]quinolone (21).** **21c** (14.9mg, 0.030mmol) was dissolved in 2ml of 5% trifluoroacetic acid dichloromethane solution and the mixture was allowed to react at room temperature for 2h. The solvent was then evaporated and the residue was dissolved with methanol, then purified by HPLC/MS on a Waters Auto Purification LC/MS system (ACQUITY UPLC ® BEH C18 17 µm 2.1X50 mm column) to afford **21** (14.7mg, 96%). $^1$H NMR (400 MHz, MeOD) δ 7.85 (d, *J* = 9.2 Hz, 1H), 7.81 (d, *J* = 2.7 Hz, 1H), 7.40 (dd, *J* = 9.2, 2.8 Hz, 1H), 3.98 (s, 3H), 3.62 (d, *J* = 6.9 Hz, 2H), 3.37 - 3.32 (m, 2H), 2.88 (s, 3H), 2.91 - 2.83 (overlapped, m, 2H), 2.12 (d, *J* = 13.8 Hz, 2H), 1.82-1.70 (m, 1H), 1.55-1.42 (m, 2H). $^{13}$C NMR (151 MHz, Chloroform-d) δ 172.0, 159.6, 158.8, 146.1, 142.4, 133.9, 130.7, 129.6, 123.5, 104.3, 97.5, 56.2, 44.9, 40.7, 35.7, 29.3, 15.7. HRMS(ESI) $[M + H]^+$ calculated for $C_{18}H_{22}N_3OS_3$: 392.0920, found: 392.0920.

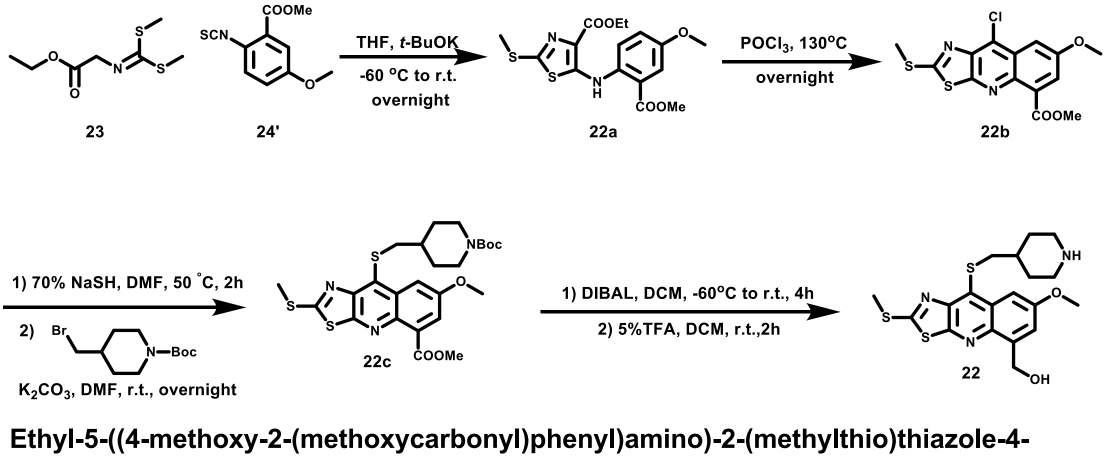


**Ethyl-5-((4-methoxy-2-(methoxycarbonyl)phenyl)amino)-2-(methylthio)thiazole-4-carboxylate (22a).** Under argon atmosphere, a solution of a known compound 23 (150mg, 0.73mmol) in 1 ml of anhydrous tetrahydrofuran was added dropwise to a solution of potassium tert-butoxide (112mg, 0.92mmol) in 8ml of anhydrous tetrahydrofuran at -60 °C, followed by a solution of known compound 24' (150mg, 0.66mmol) in 1 ml of anhydrous tetrahydrofuran. The reaction was performed at -60 °C for 1.5 h and then slowly warmed up to room temperature and reacted overnight. The reaction was quenched by adding saturated ammonium chloride, diluted with dichloromethane, washed twice with brine, dried over anhydrous $Na_2SO_4$ and concentrated *in vacuo*. The residue was purified by chromatography on a silica gel column (pure dichloromethane) to give compound **22a** (69.2mg, 28%) as a yellow solid. $^1$H NMR (400 MHz, CDCl3) δ 11.61 (s, 1H), 7.58 - 7.50 (m, 2H), 7.11 (dd, *J* = 9.0, 3.2 Hz, 1H), 4.49 (q, *J* = 7.1 Hz, 2H), 3.97 (s, 3H), 3.83 (s, 3H), 2.64 (s, 3H), 1.42 (t, *J* = 7.1 Hz, 3H). $^{13}$C NMR (101 MHz, Chloroform-d) δ 167.1, 163.9, 154.3, 153.6, 147.0, 137.0, 125.6, 121.0, 117.9, 117.4, 115.8, 61.1, 55.9, 52.7, 17.7, 14.7. HRMS(ESI) $[M + H]^+$ calculated for $C_{16}H_{19}N_2O_5S_2$: 383.0730, found: 383.0729.

**Methyl-9-chloro-7-methoxy-2-(methylthio)thiazolo[5,4-b]quinoline-5-carboxylate (22b).** Compound **22a** (50mg, 0.14mmol) was added in a sealed tube, and 1 ml of phosphorus oxychloride was added under argon atmosphere. The reaction was heated at 130 °C overnight. The resulting slurry was poured onto ice with vigorous stirring to make full quenching. The mixture was diluted with dichloromethane, washed twice with brine, dried over anhydrous $Na_2SO_4$ and concentrated *in vacuo*. The residue was purified by chromatography on a silica gel column (pure dichloromethane) to give compound **22b** (15mg, 32%) as a light yellow solid. $^1$H NMR (400 MHz, $CDCl_3$) δ 7.69 (s, 2H), 4.05 (s, 3H), 4.02 (s, 3H), 2.88 (s, 3H). $^{13}$C NMR (101 MHz, Chloroform-d) δ 173.4, 167.5, 157.2, 142.6, 139.3, 132.7, 129.4, 126.6, 123.5, 105.1, 56.1, 53.0, 15.5. HRMS(ESI) $[M + H]^+$ calculated for $C_{14}H_{12}ClN_2O_3S_2$: 354.9972, found: 354.9973.

**Methyl-9-(((1-(tert-butoxycarbonyl)piperidin-4-yl)methyl)thio)-7-methoxy-2-(methylthio)thiazolo[5,4-b]quinoline-5-carboxylate (22c).** To a solution of **22b** (15mg, 0.042mmol) in 2 ml of anhydrous DMF, sodium hydrogen hydride hydrate powder (70%, 6.7mg, 0.084mmol) was added under argon atmosphere, and the reaction was stirred at 50 °C for 2 h until full conversion of **1c**. 1-BOC-4-bromomethylpiperidine (29.3mg, 0.105mmol) and potassium carbonate (17.4mg, 0.126mmol) were added into the slurry and the reaction was allowed to react at room temperature overnight. The solvent was then evaporated and the residue was dissolved with dichloromethane and washed with water. The combined organic extracts were dried over anhydrous $Na_2SO_4$ and concentrated *in vacuo*. The residue was purified by chromatography on a silica gel column (dichloromethane / ethyl acetate = 95/5) to give compound **22c** (16.2mg, 70%) as a light yellow solid. $^1$H NMR (400 MHz, CDCl3) δ 8.02 (d, *J* = 2.9 Hz, 1H), 7.65 (d, *J* = 2.8 Hz, 1H), 4.12-3.96 (overlapped, m, 2H), 4.04 (s, 3H), 4.00 (s, 3H), 3.44 (d, *J* = 6.9 Hz, 2H), 2.84 (s, 3H), 2.58 (t, *J* = 12.6 Hz, 2H), 1.82 (d, *J* = 12.9 Hz, 2H), 1.61 - 1.52 (m, 1H), 1.44 (s, 9H), 1.23 - 1.13 (m, 2H). $^{13}$C NMR (151 MHz, Chloroform-d) δ 171.2, 167.8, 158.8, 156.6, 154.8, 145.2, 138.5, 132.7, 132.6, 129.1,

122.5, 106.7, 79.4, 55.9, 52.8, 44.3-43.2 (m), 41.1, 36.7, 31.5, 28.4, 15.4. HRMS(ESI) [M + H]$^+$ calculated for $C_{25}H_{32}N_3O_5S_3$: 550.1499, found: 550.1492.

**(7-methoxy-2-(methylthio)-9-((piperidin-4-ylmethyl)thio)thiazolo[5,4-b]quinolin-5-yl)methaol (22).** 87μl of 1.0M DIBAL-H solution in toluene was added to a solution of **22c** (16.0mg, 0.029mmol) in 1 ml of dry dichloromethane at -60 °C, and the reaction was slowly warmed to room temperature and reacted overnight. The reaction was quenched by adding saturated potassium hydrogen tartrate solution, diluted with dichloromethane, washed twice with brine, dried over anhydrous $Na_2SO_4$ and concentrated *in vacuo*. The residue was dissolved in 4ml of 5% trifluoroacetic acid dichloromethane solution and the mixture was allowed to react at room temperature for 2h. The solvent was then evaporated and the residue was dissolved with methanol, then purified by HPLC/MS on a Waters Auto Purification LC/MS system (ACQUITY UPLC ® BEH C18 17 μm 2.1X50 mm column) to afford **22** (5.3mg, 34%). $^1$H NMR (400 MHz, MeOD) δ 7.78 (d, *J* = 2.8 Hz, 1H), 7.52 (d, *J* = 2.7 Hz, 1H), 5.20 (s, 2H), 3.98 (s, 3H), 3.58 (d, *J* = 6.9 Hz, 2H), 3.37-3.32 (m, 2H), 2.88 (s, 3H), 2.88 - 2.81 (m, 2H), 2.15 - 2.08 (m, 2H), 1.78 - 1.69 (m, 1H), 1.55 - 1.41 (m, 2H). $^{13}$C NMR (151 MHz, Methanol-d4) δ 172.2, 159.5, 157.9, 146.2, 142.2, 140.4, 133.5, 129.8, 120.6, 102.9, 61.7, 56.0, 44.9, 40.8, 35.7, 29.3, 15.6. HRMS(ESI) [M + H]$^+$ calculated for $C_{19}H_{24}N_3O_2S_3$: 422.1025, found: 422.1013.

### NMR data

**1c**

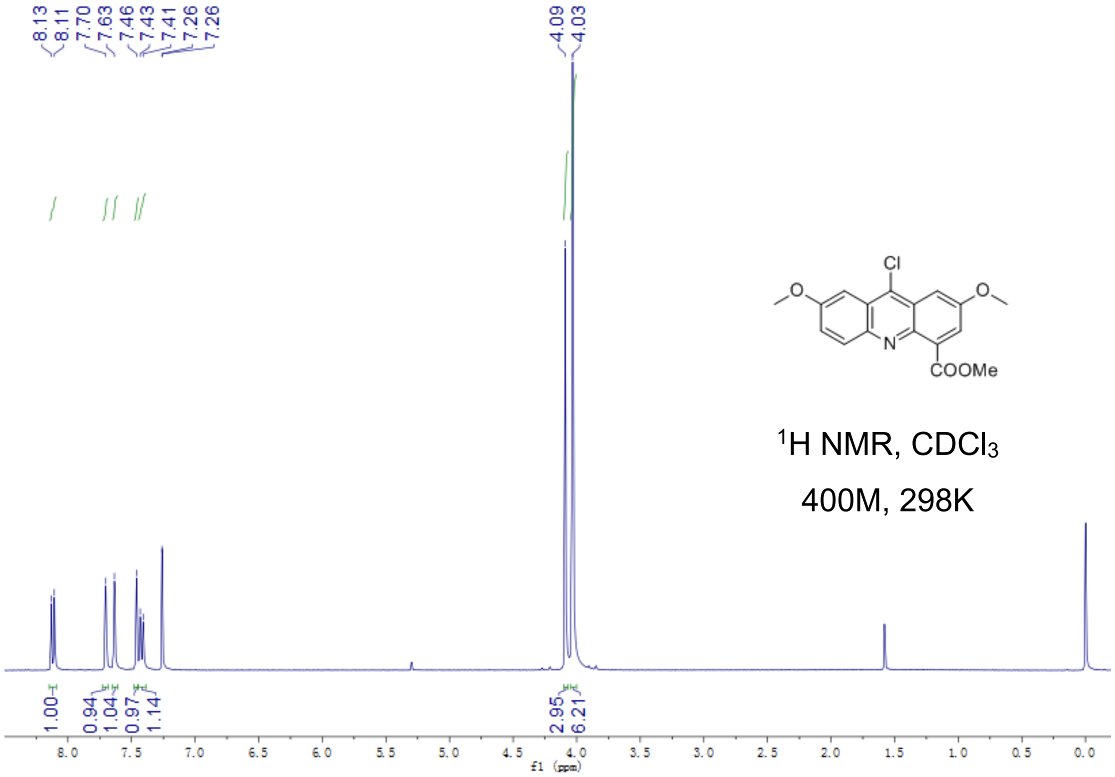


Fig S2.1. $^{1}$H-NMR of **1c**

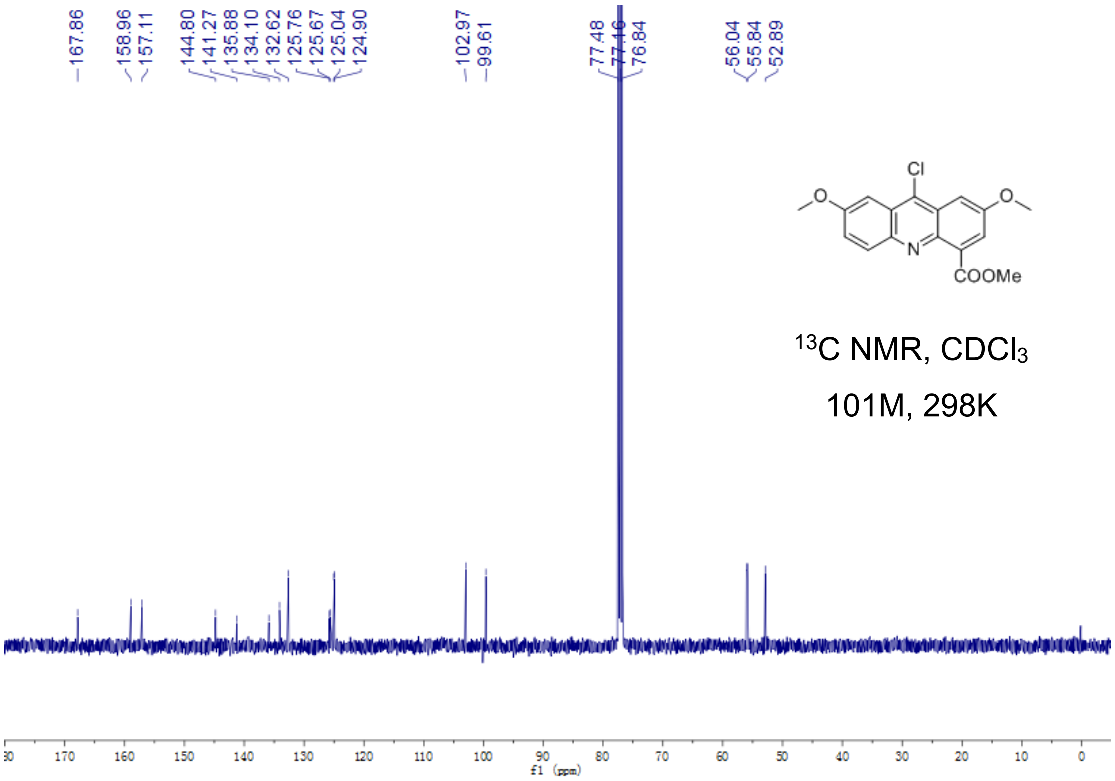


Fig S2.2. $^{13}$C-NMR of **1c**

**1**

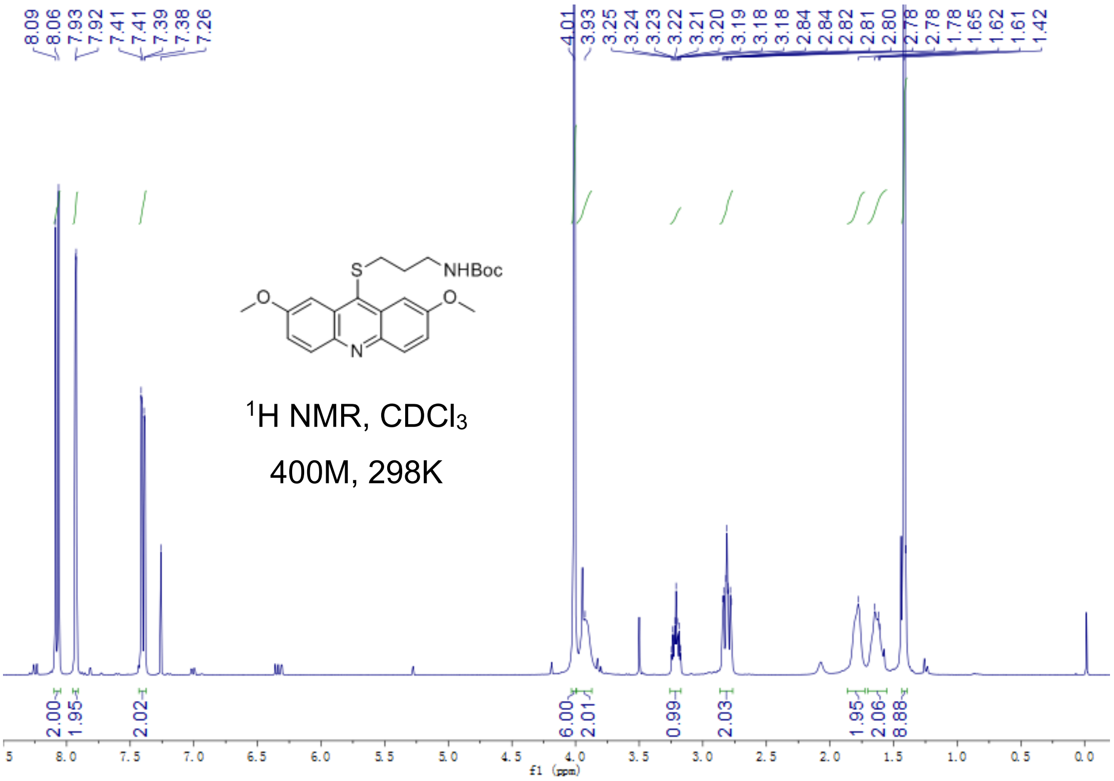


Fig S2.3. $^{1}$H-NMR of **1**

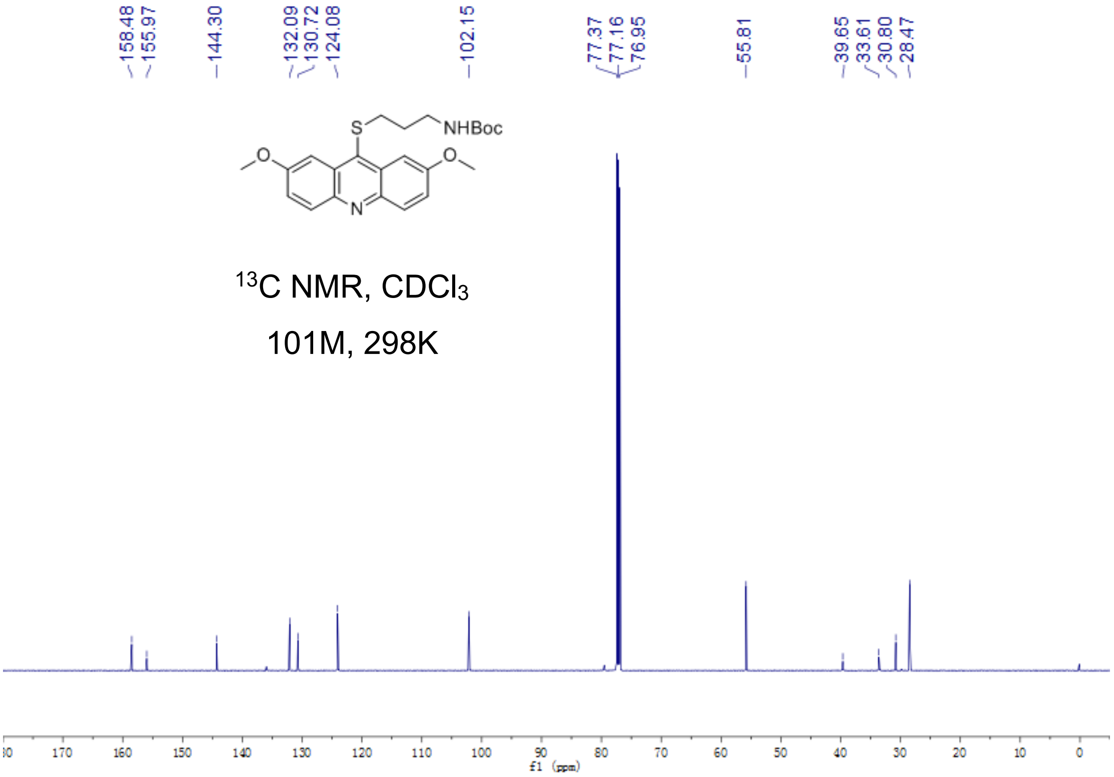


Fig S2.4. $^{13}$C-NMR of **1**

**1c'**

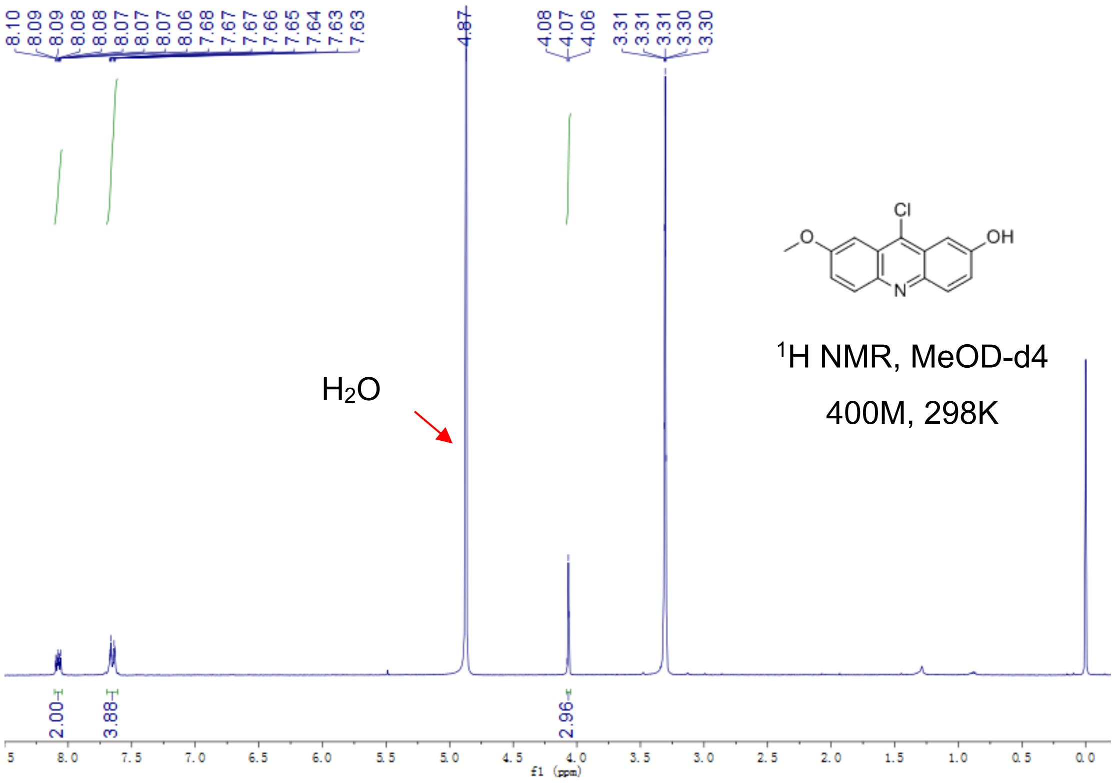


Fig S2.5. $^1$H-NMR of **1c'**

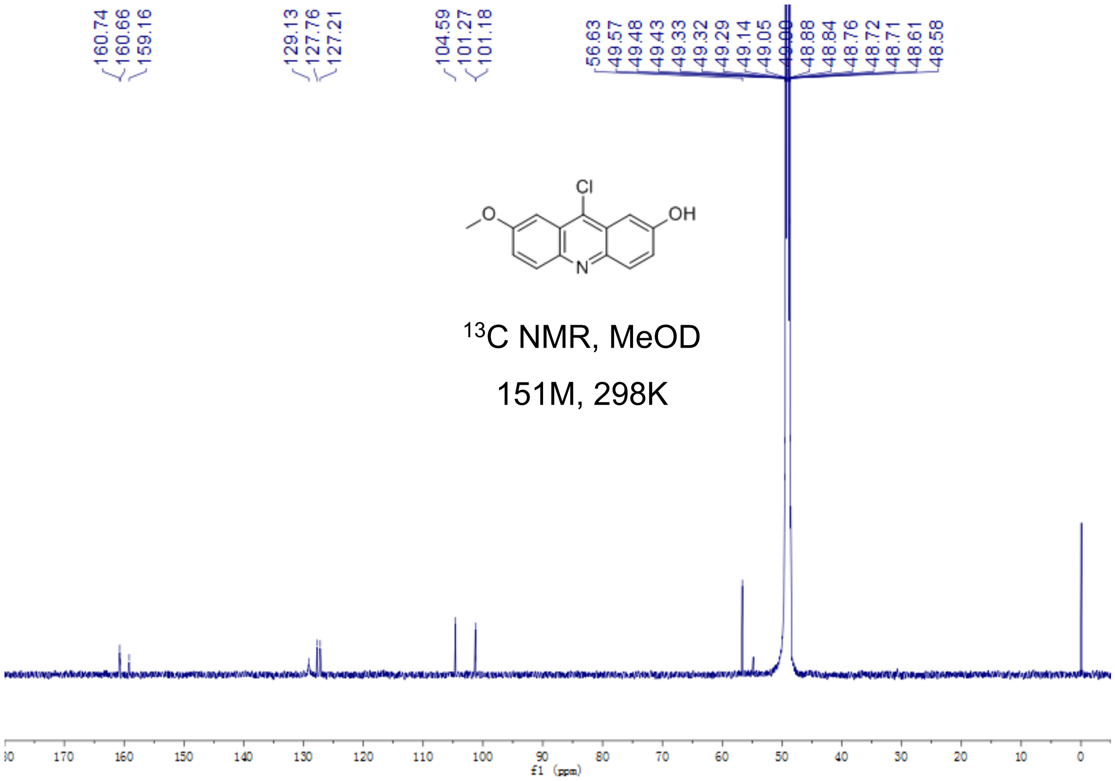


Fig S2.6. $^{13}$C-NMR of **1c'**

**2**

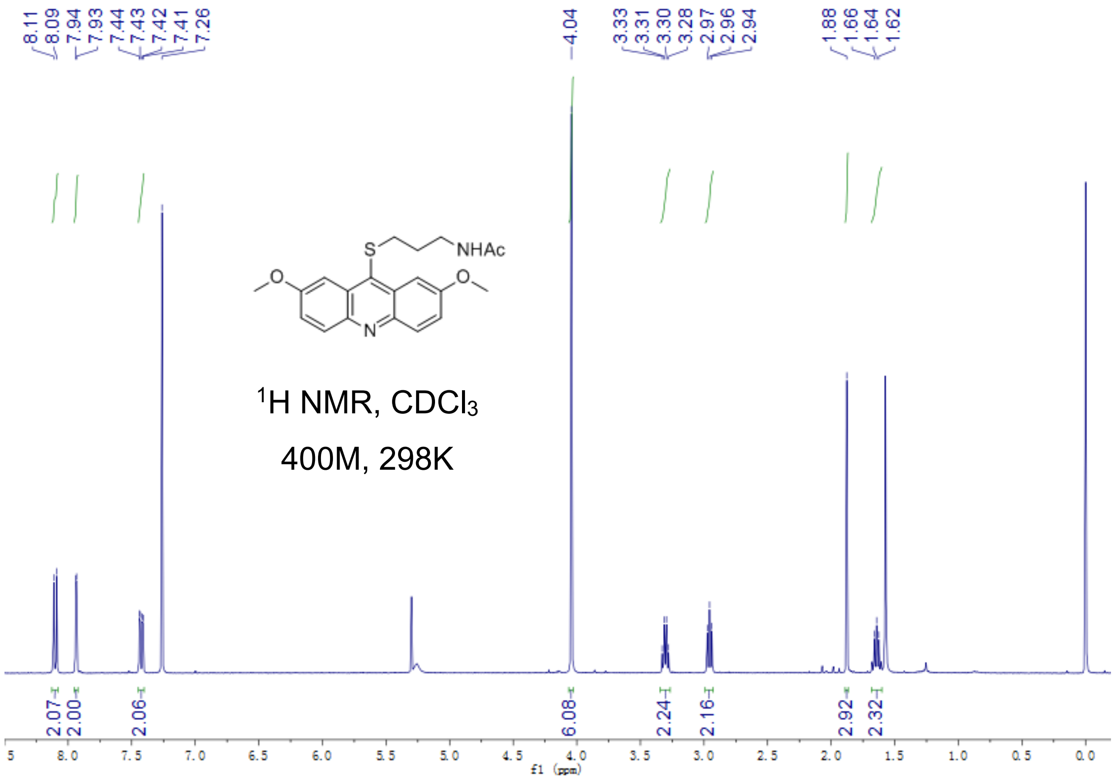


Fig S2.7. $^{1}$H-NMR of **2**

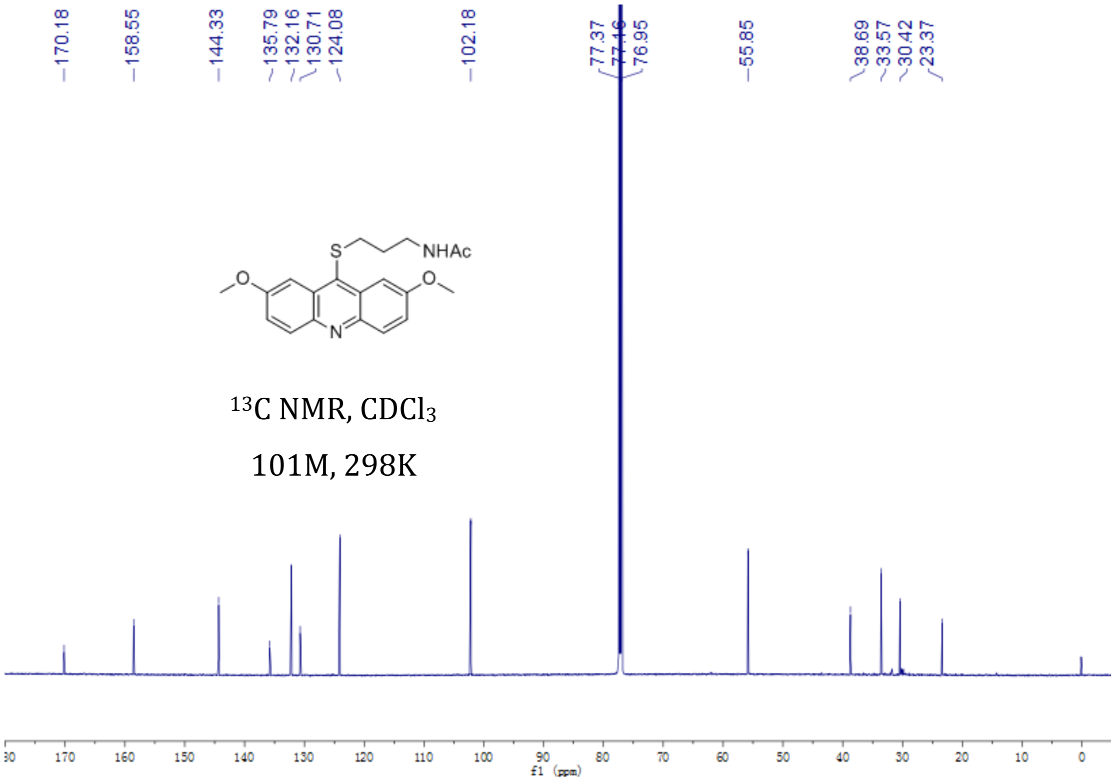

Fig S2.8. $^{13}$C-NMR of **2**

**3**

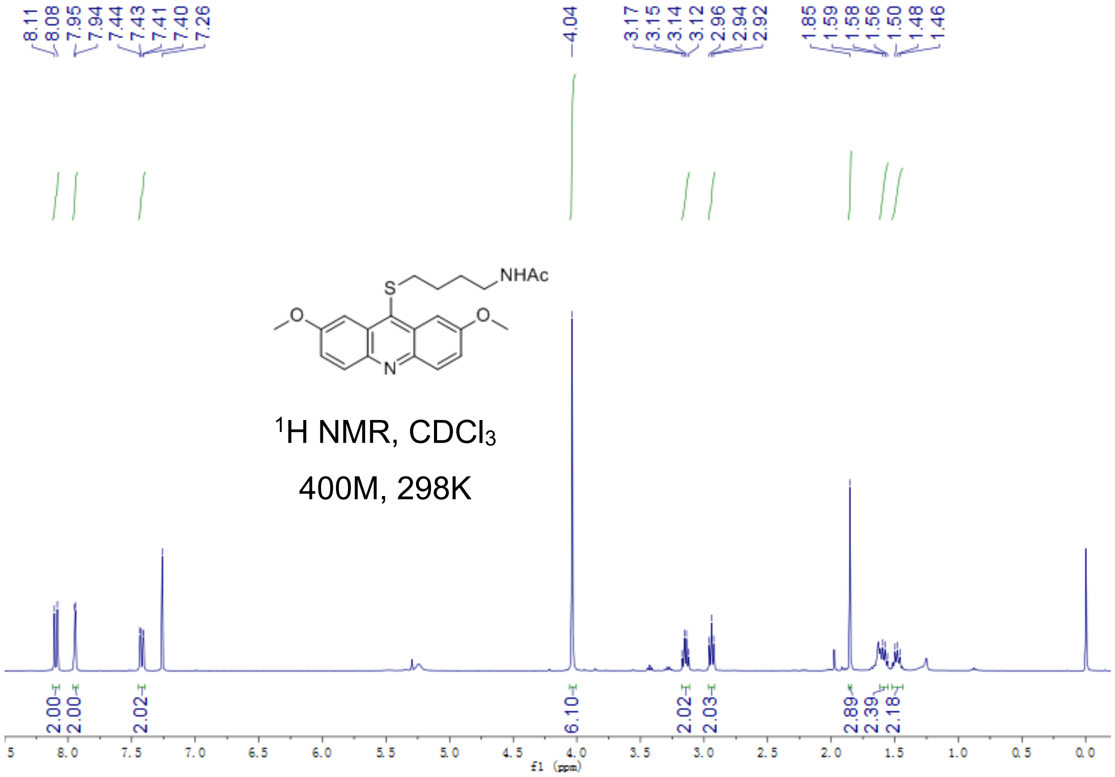


Fig S2.9. $^{1}$H-NMR of **3**

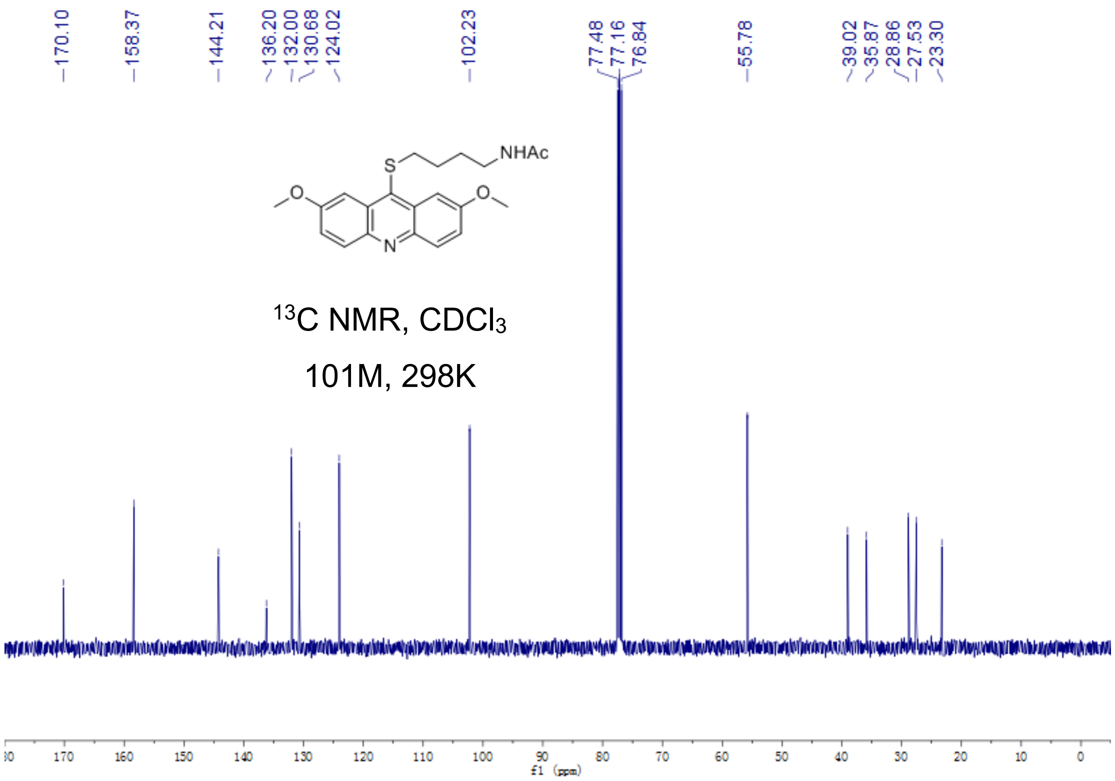


Fig S2.10. $^{13}$C-NMR of **3**

**4**

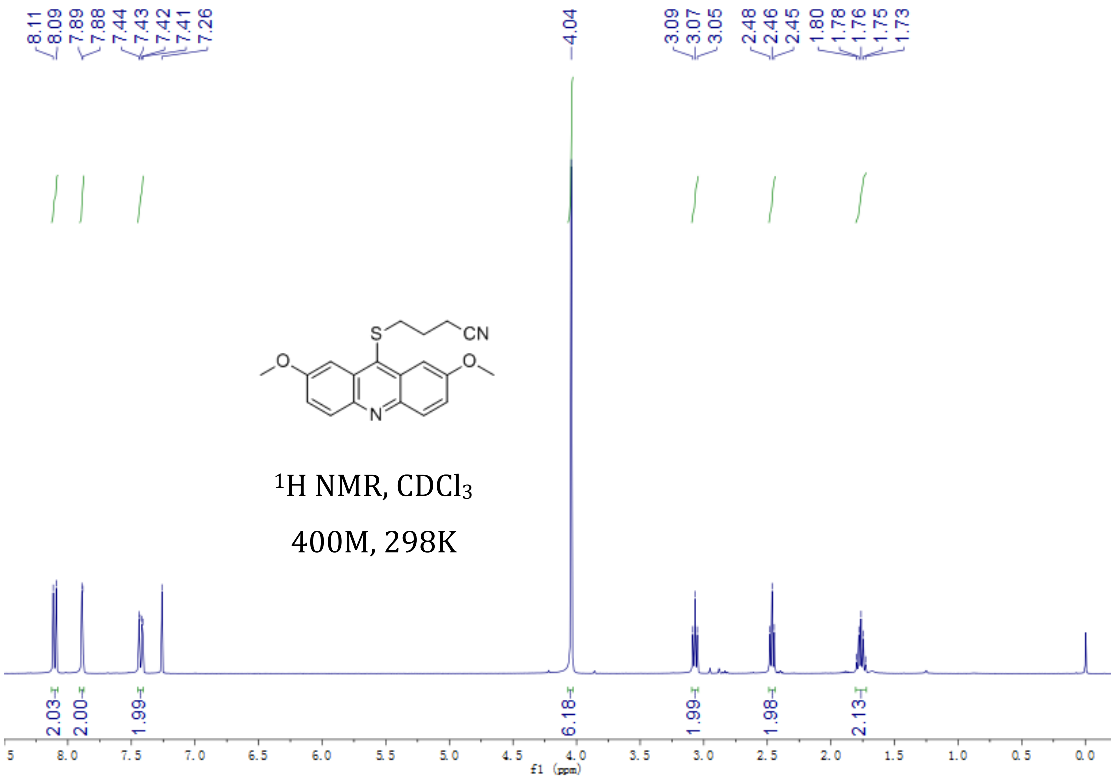


Fig S2.11. $^{1}$H-NMR of **4**

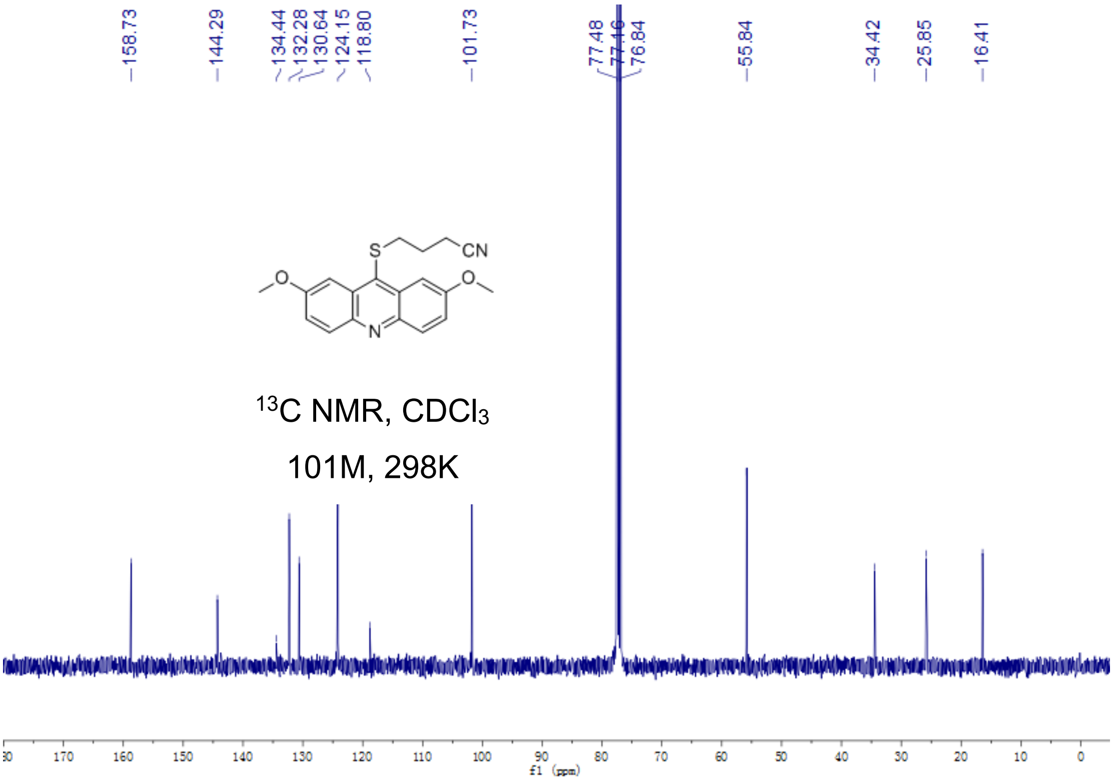


Fig S2.12. $^{13}$C-NMR of **4**

**5a**

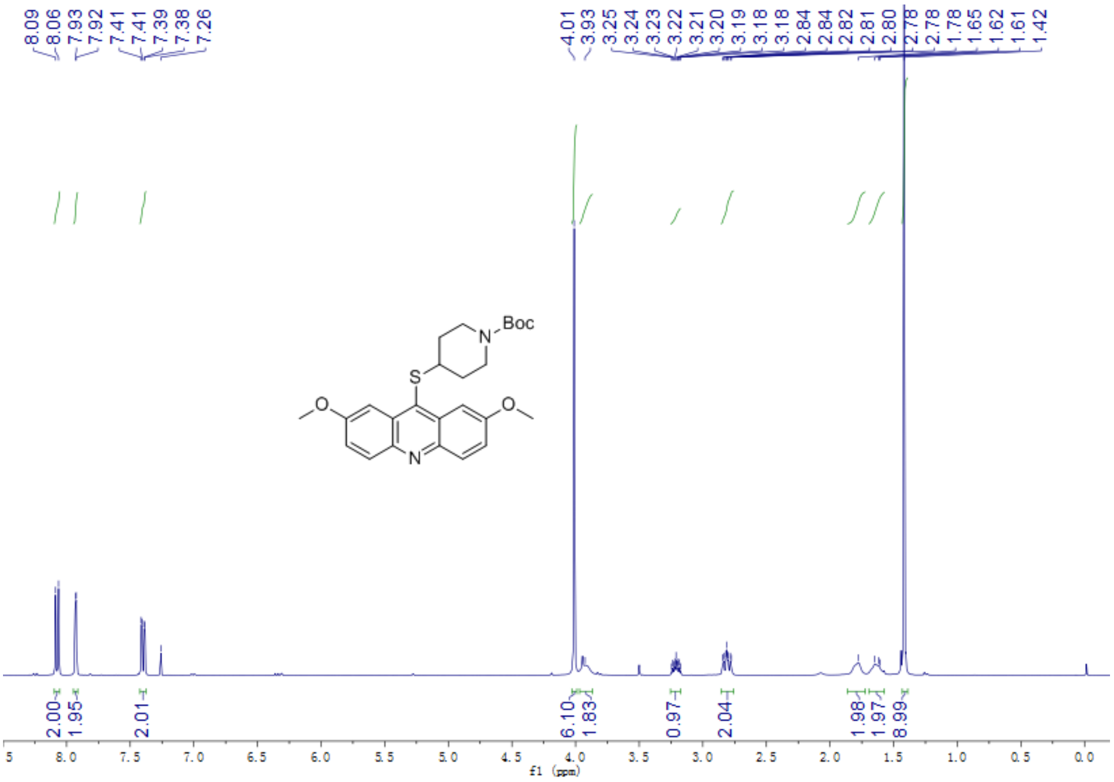

400M, 298K

Fig S2.13. $^{1}H$-NMR of **5a**

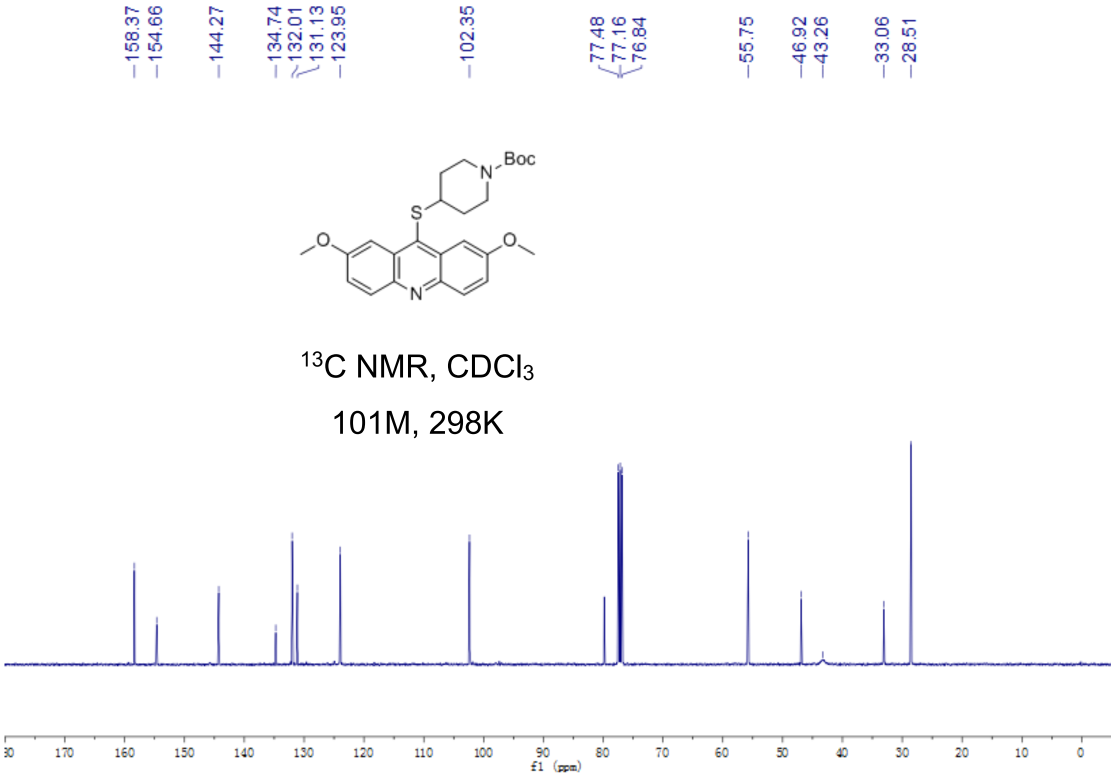


Fig S2.14. $^{13}C$-NMR of **5a**

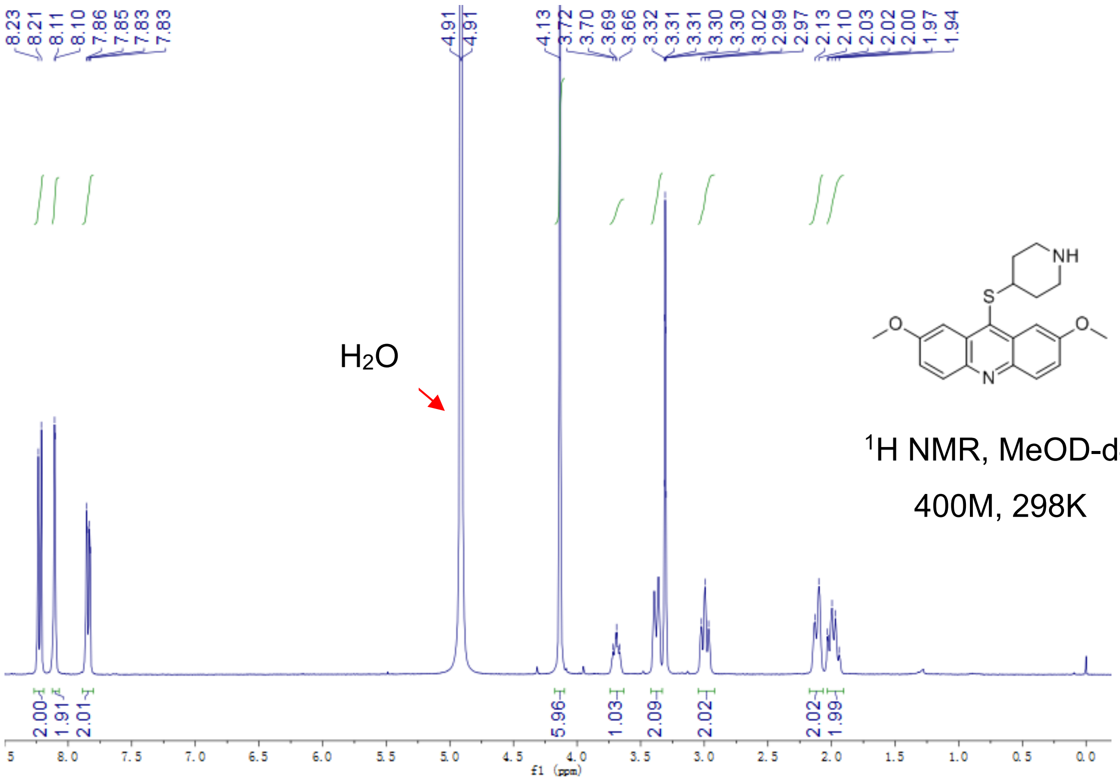


Fig S2.15. $^{1}$H-NMR of **5**

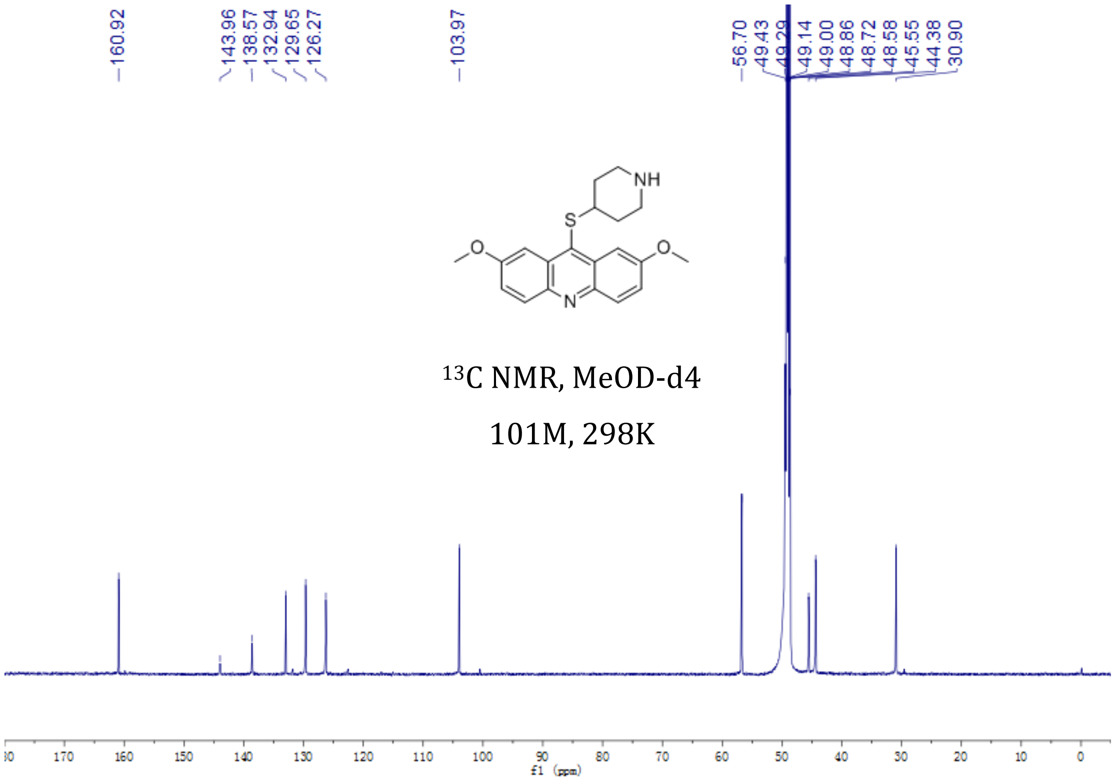


Fig S2.16. $^{13}$C-NMR of **5**

**6a**

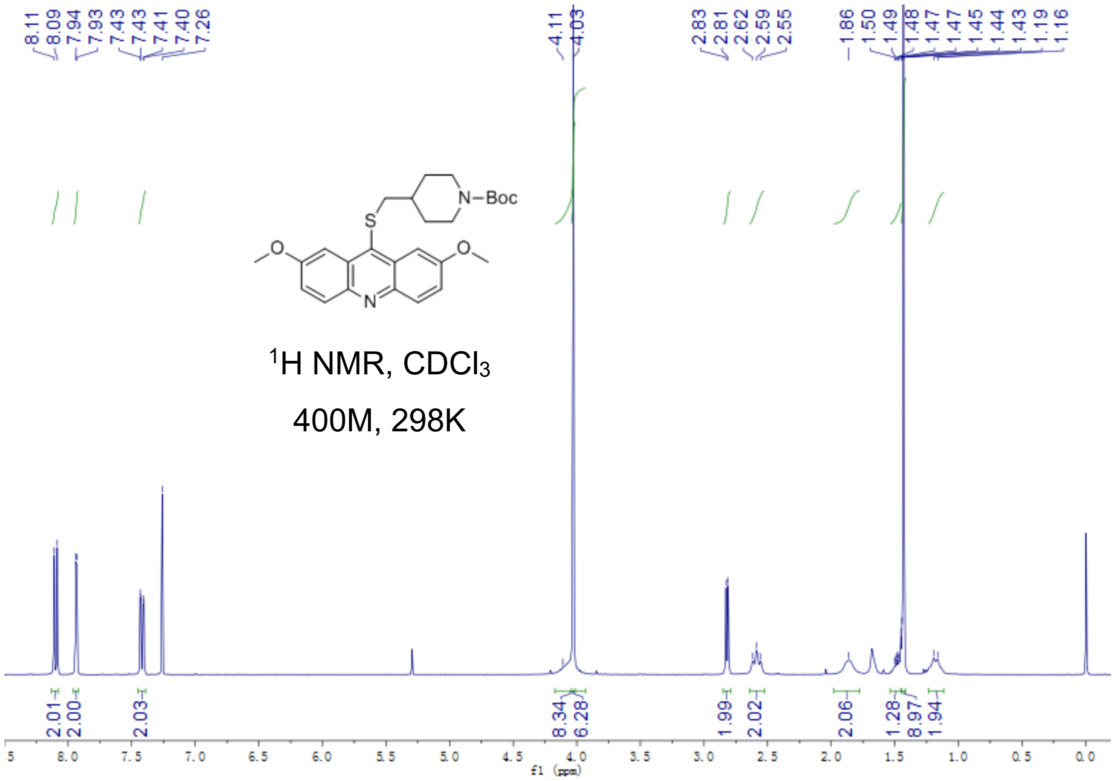


Fig S2.17. $^{1}$H-NMR of **6a**

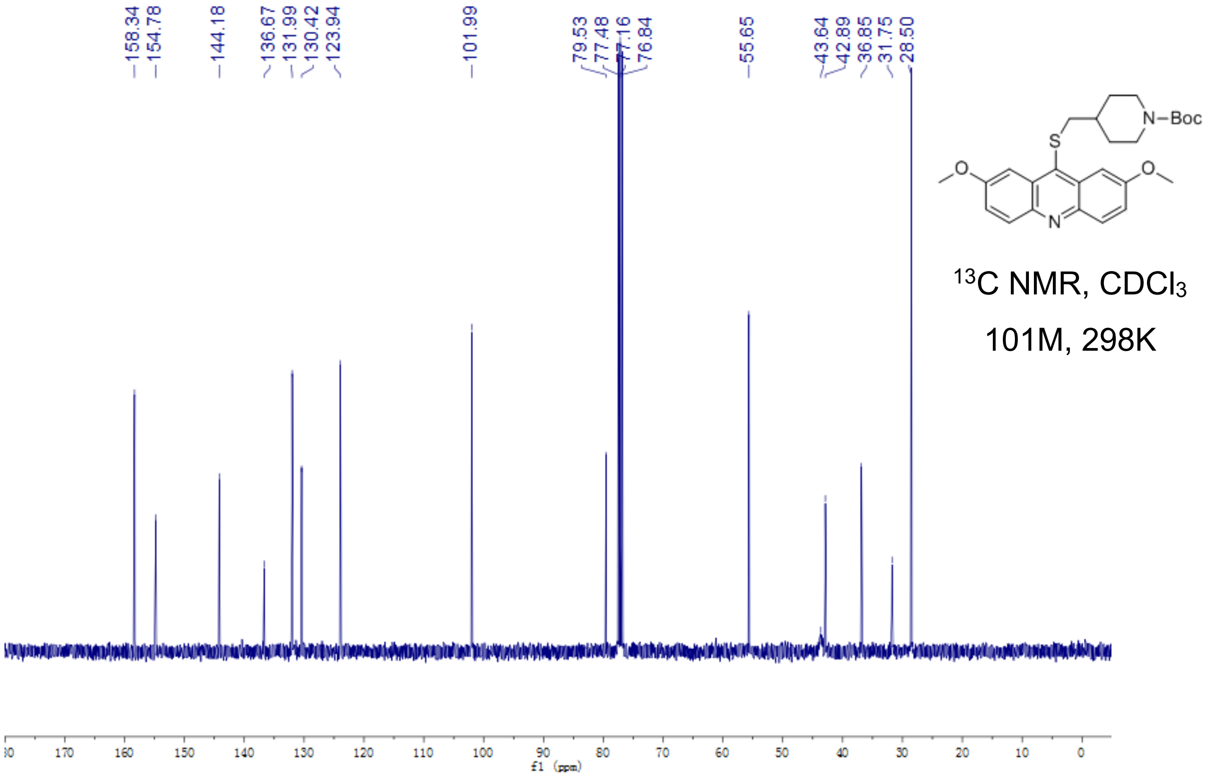


Fig S2.18. $^{13}$C-NMR of **6a**

**6**

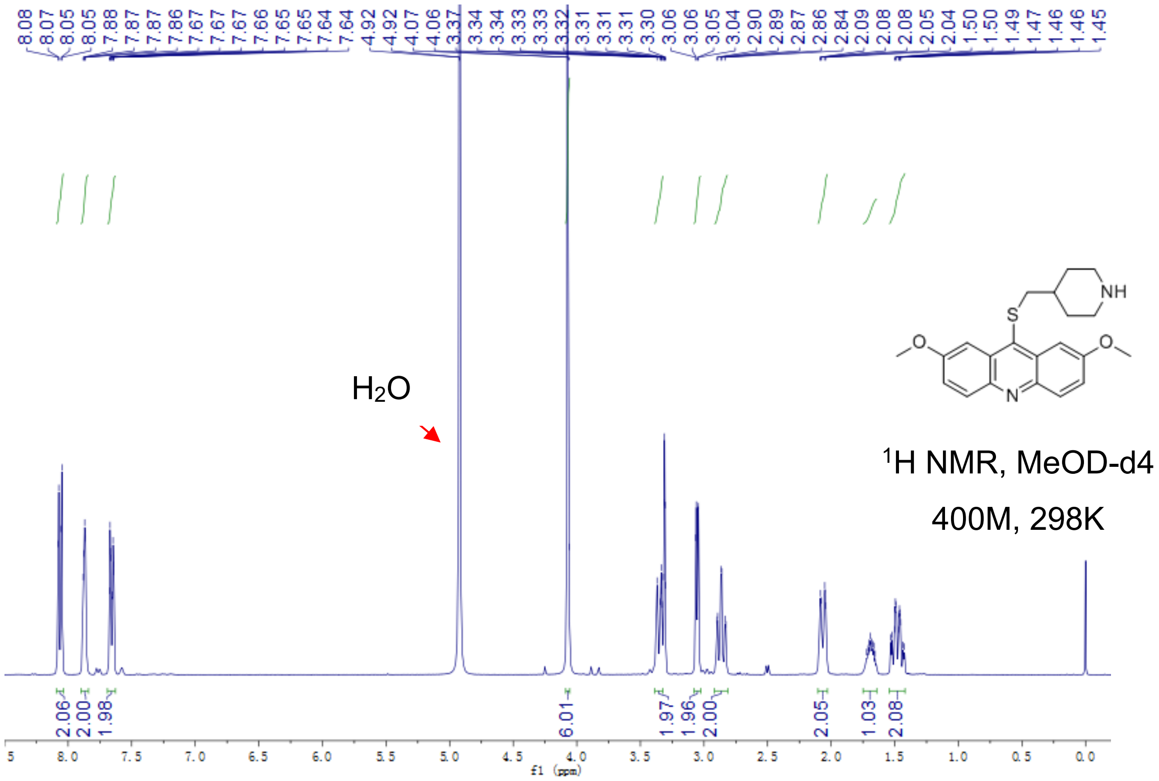


Fig S2.19. $^1$H-NMR of **6**

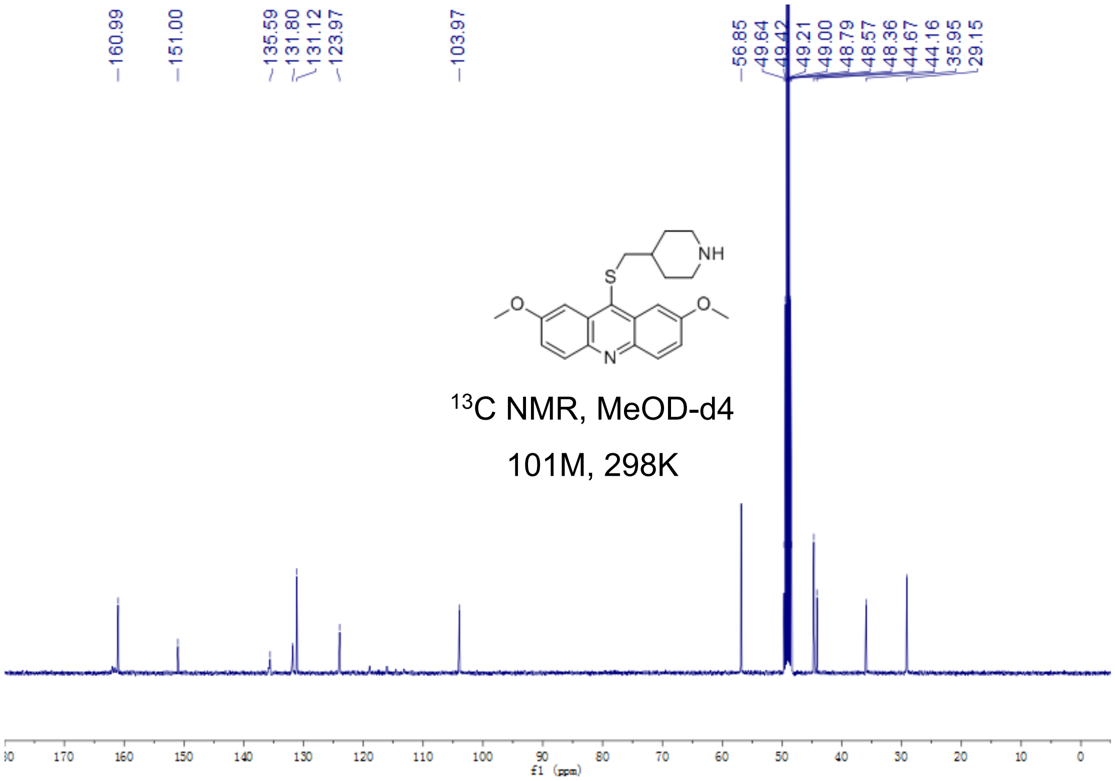


Fig S2.20. $^{13}$C-NMR of **6**

**7a**

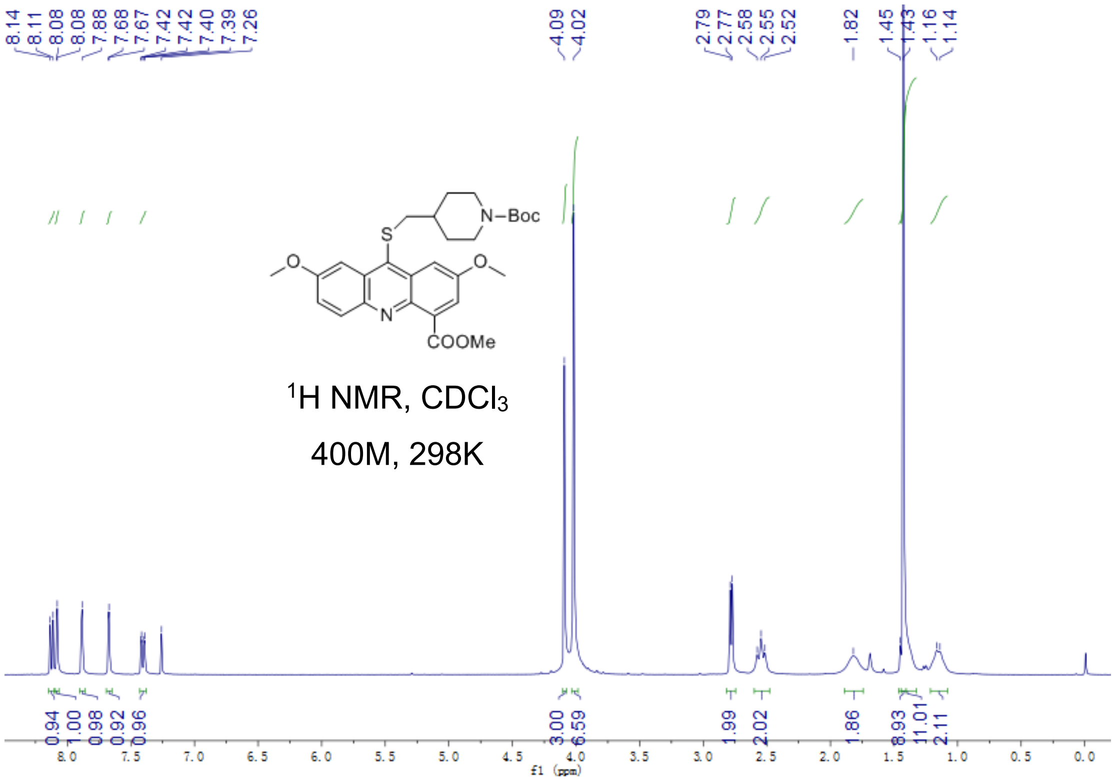


Fig S2.21. $^{1}$H-NMR of **7a**

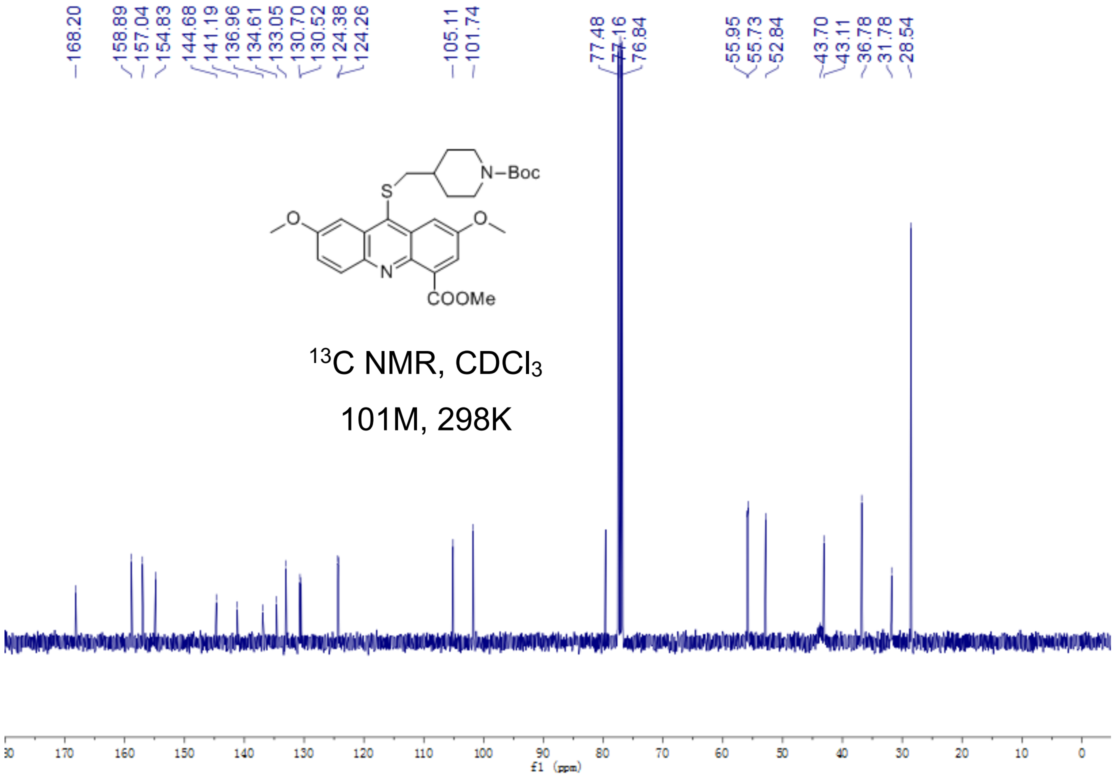

Fig S2.22. $^{13}$C-NMR of **7a**

**7b**

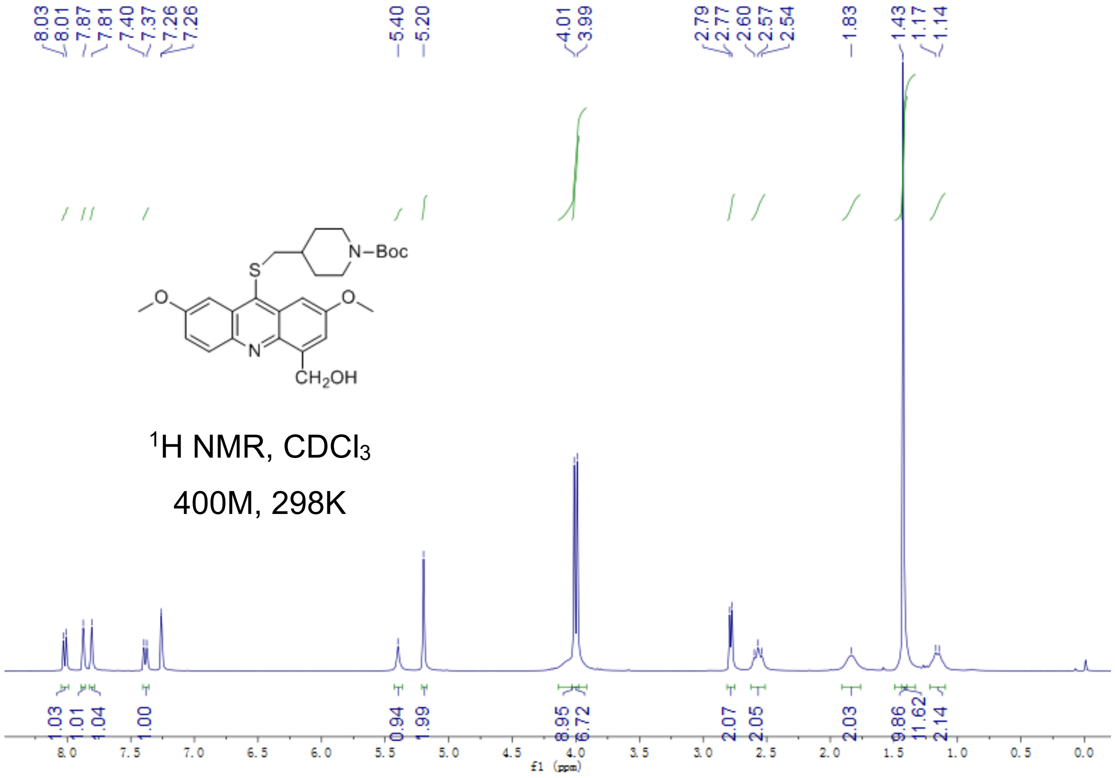


Fig S2.23. $^{1}$H-NMR of **7b**

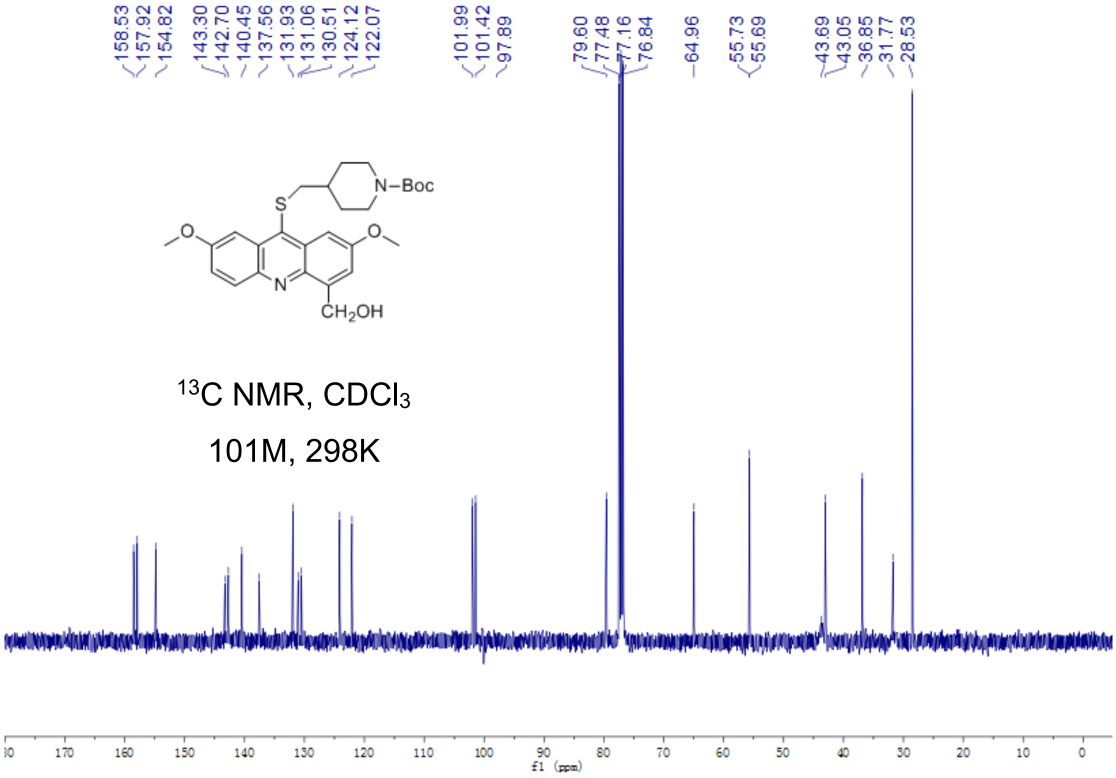


Fig S2.24. $^{13}$C-NMR of **7b**

**7**

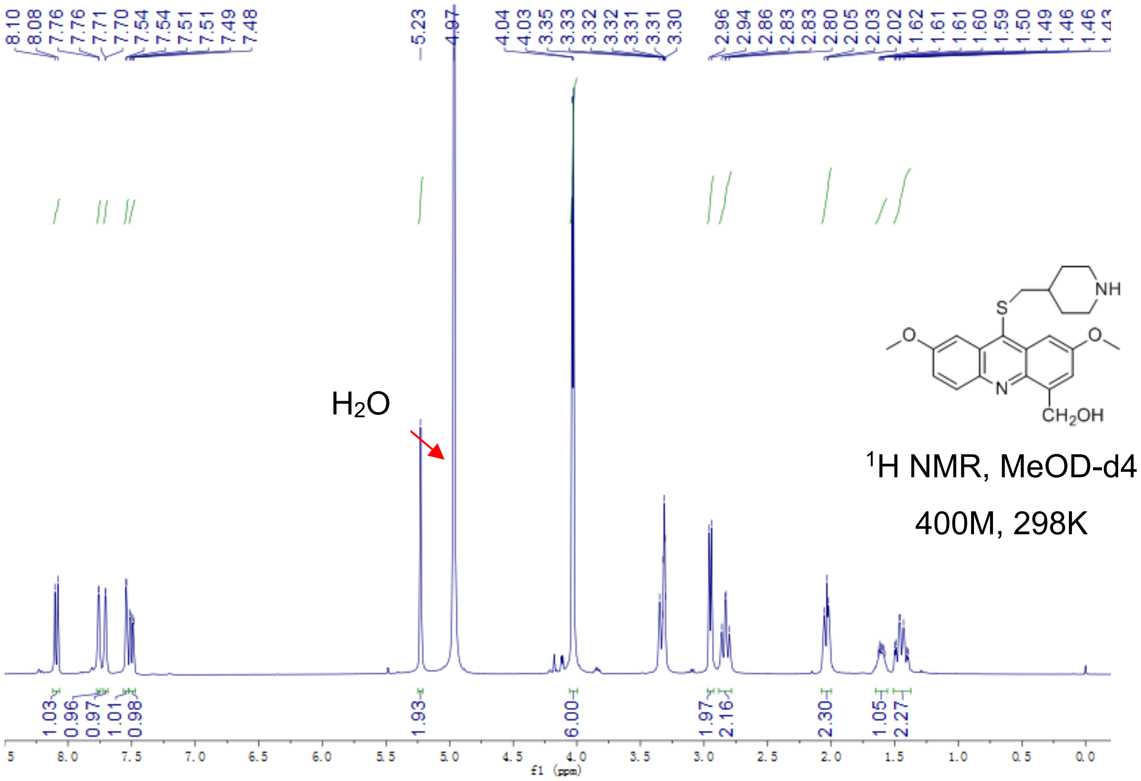


Fig S2.25. $^{1}$H-NMR of **7**

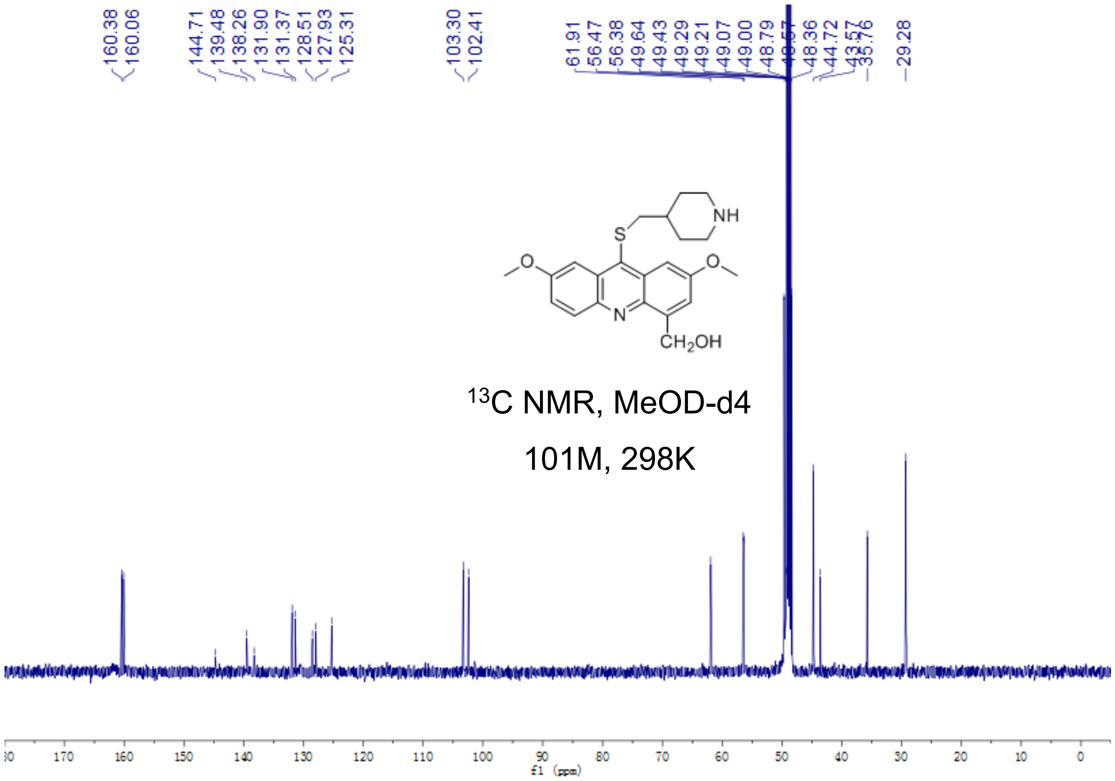


Fig S2.26. $^{13}C$-NMR of **7**

**8**

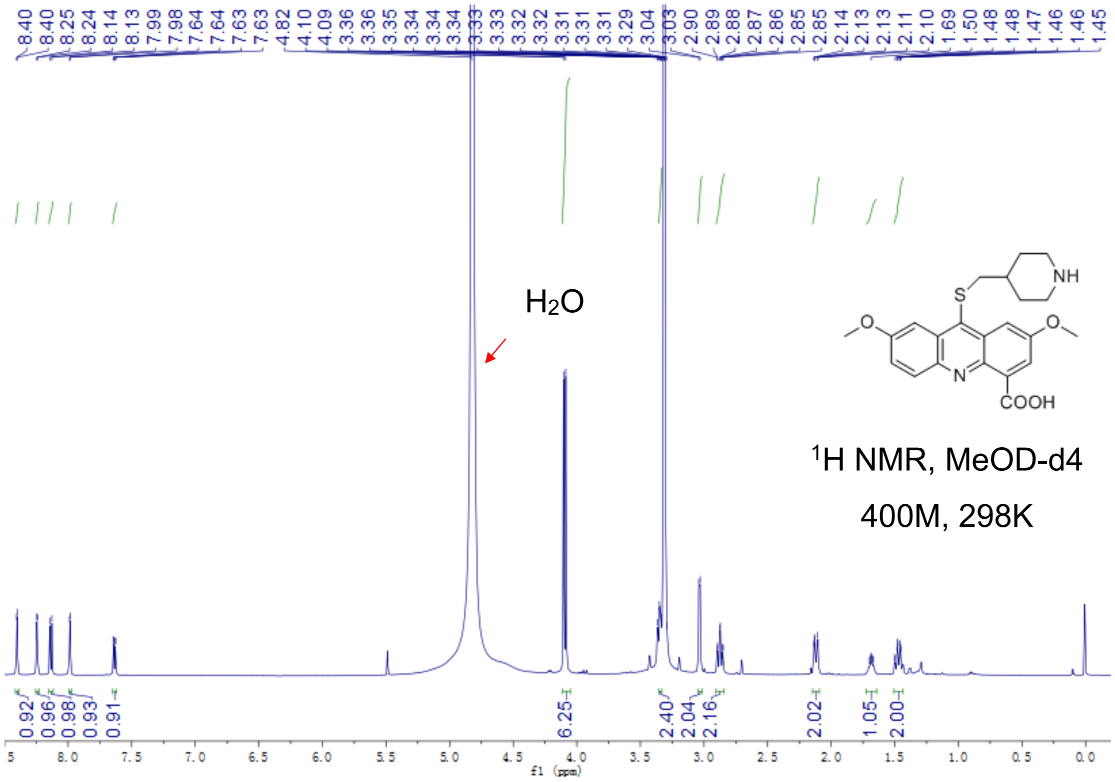

Fig S2.27. $^{1}$H-NMR of **8**

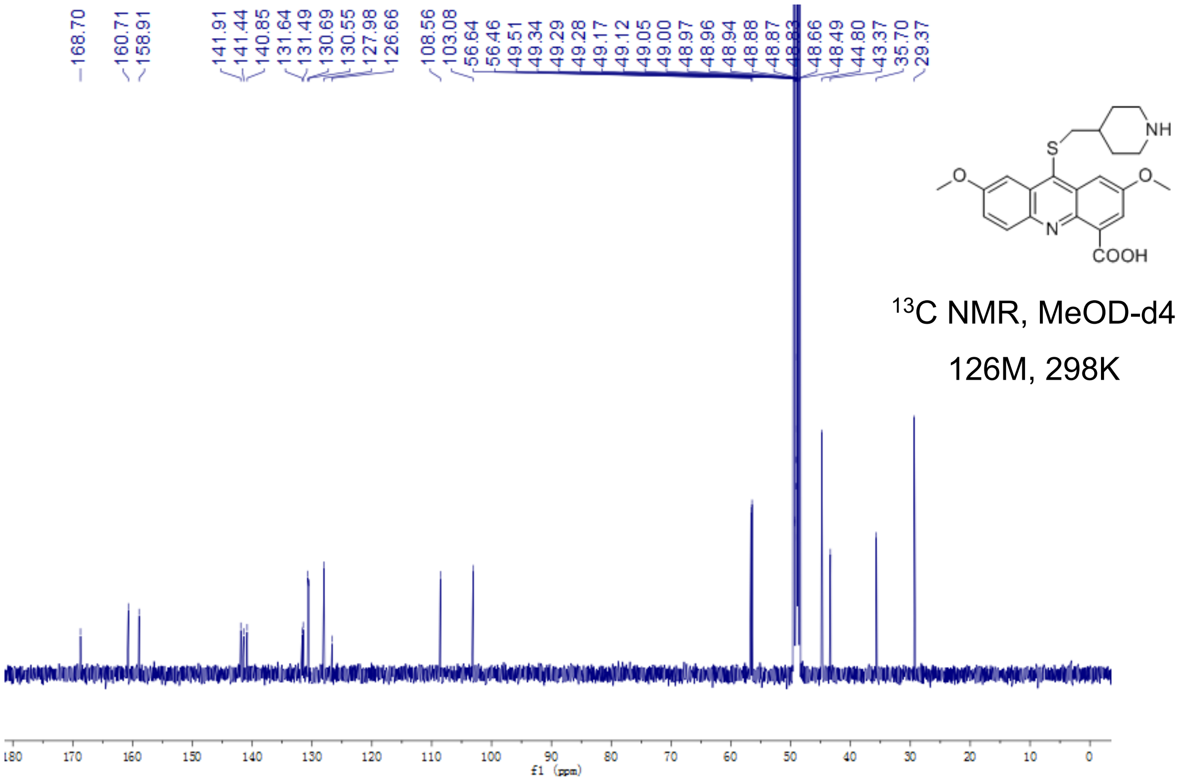


Fig S2.28. $^{13}$C-NMR of **8**

**9a**

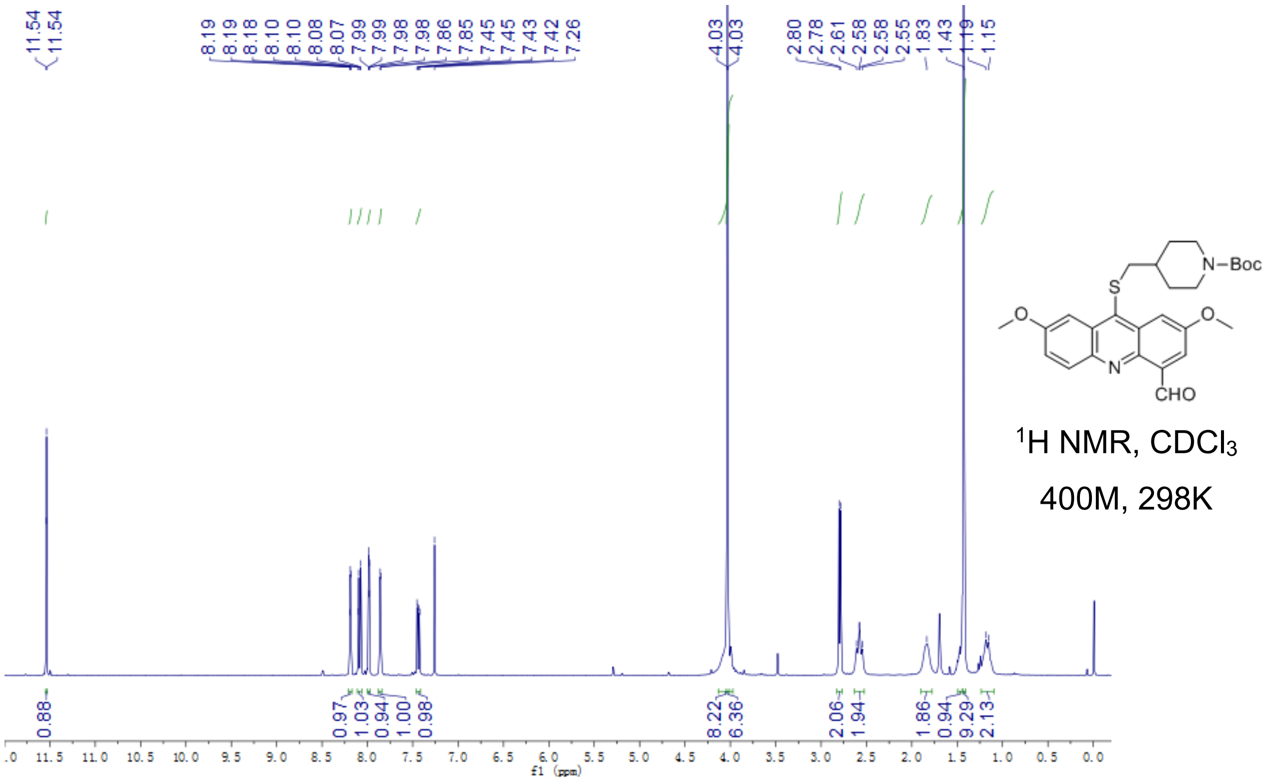


Fig S2.29. $^1$H-NMR of **9a**

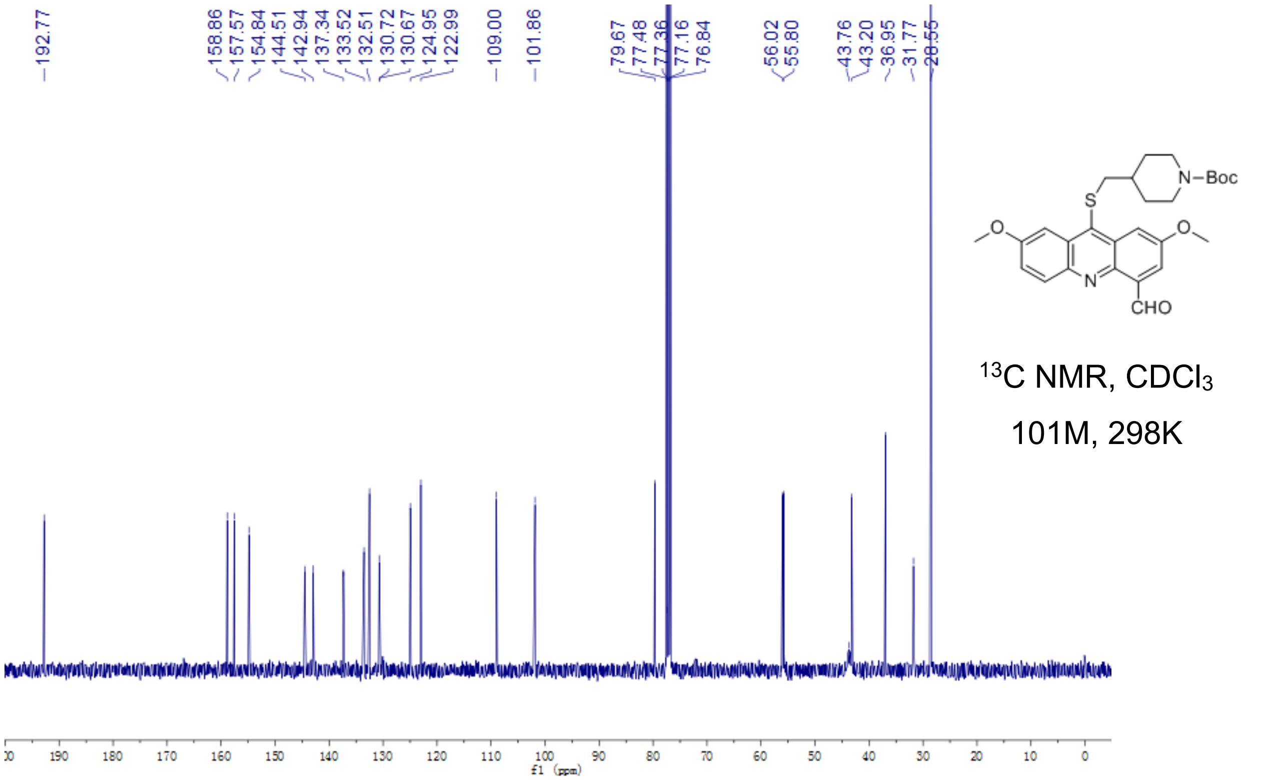


Fig S2.30. $^{13}$C-NMR of **9a**

**9b**

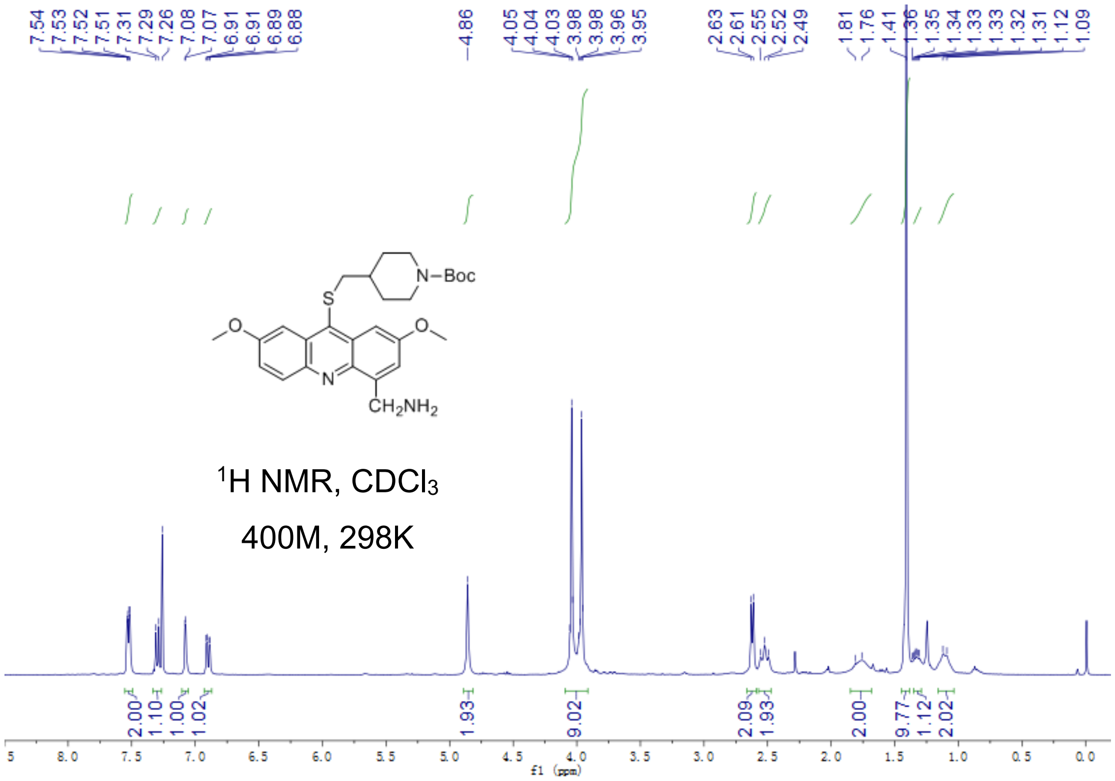


Fig S2.31. $^{1}$H-NMR of **9b**

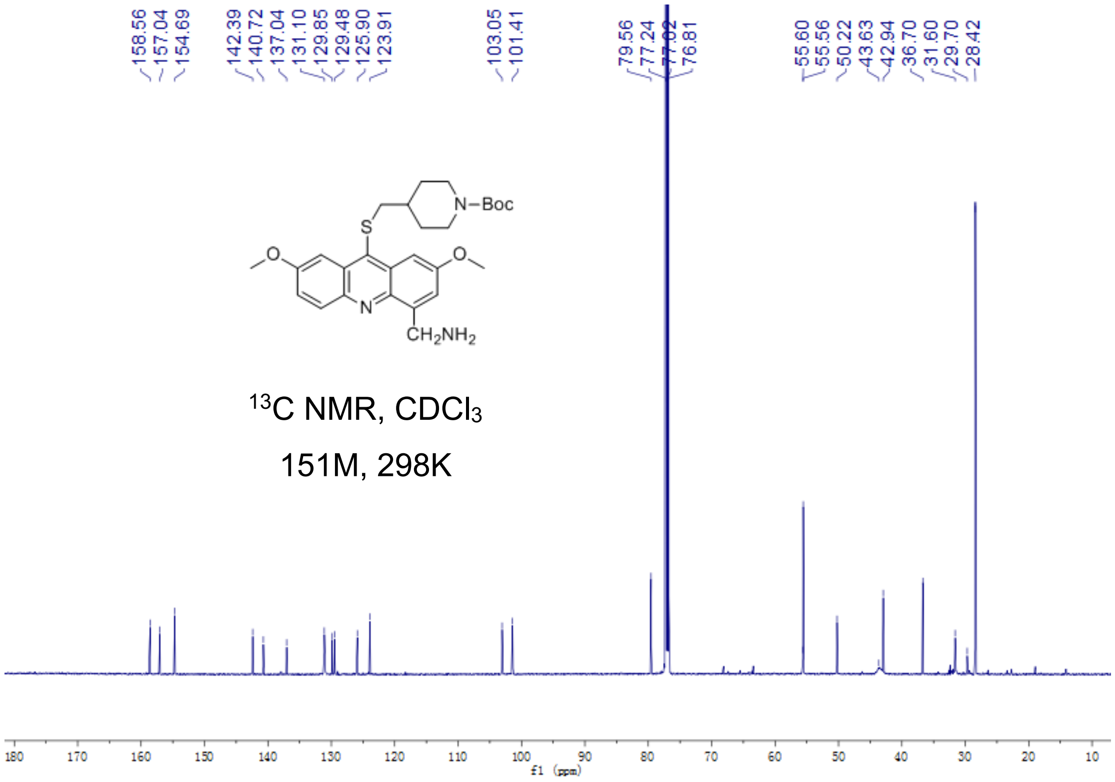


Fig S2.32. $^{13}$C-NMR of **9b**

**9**

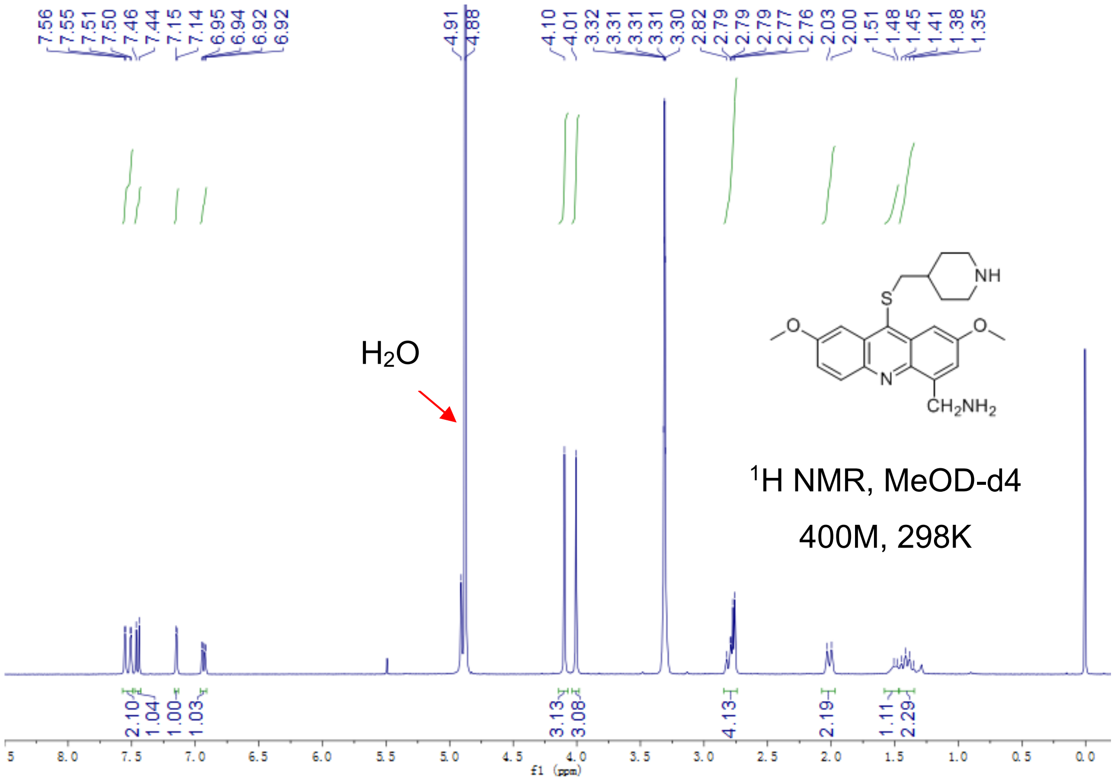


Fig S2.33. $^{1}$H-NMR of **9**

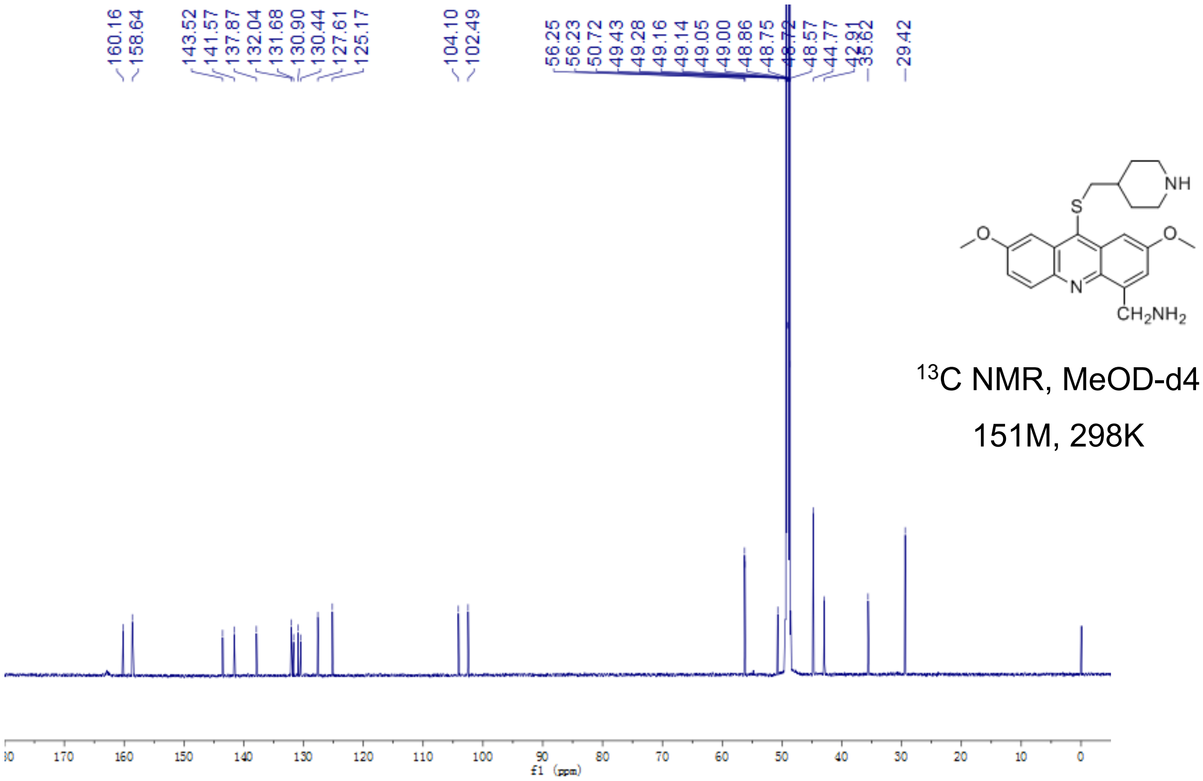


Fig S2.34. $^{13}$C-NMR of **9**

**10a**

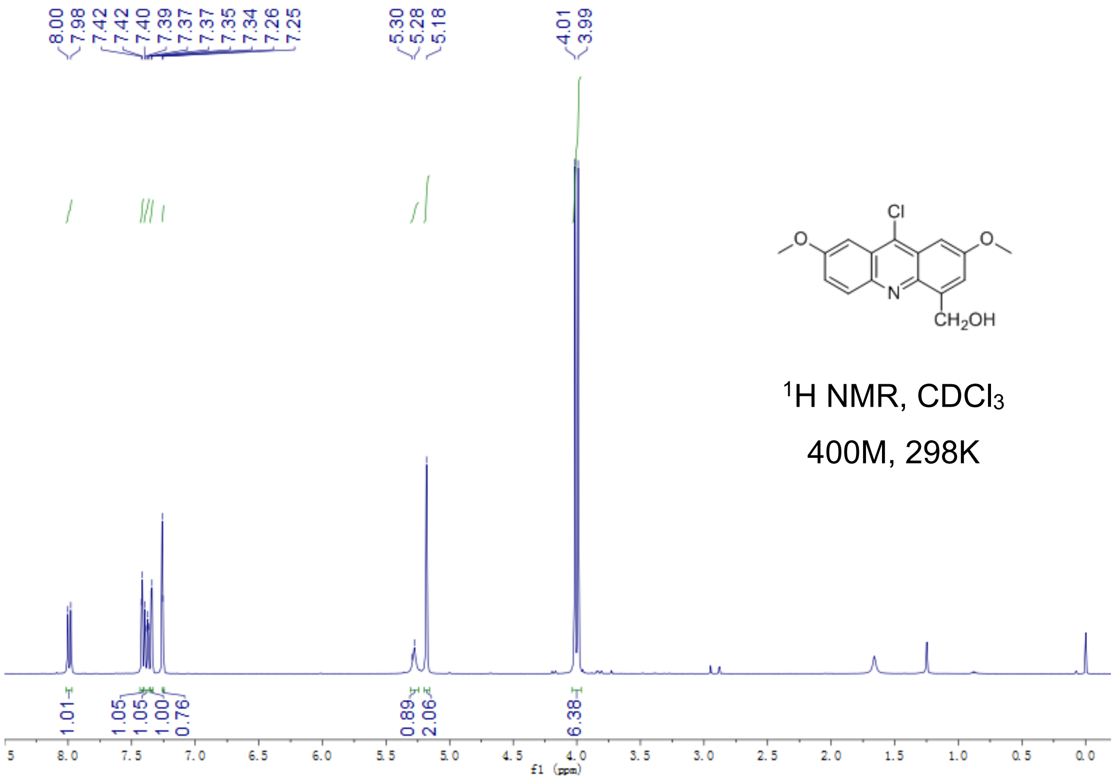


Fig S2.35. $^{1}$H-NMR of **10a**

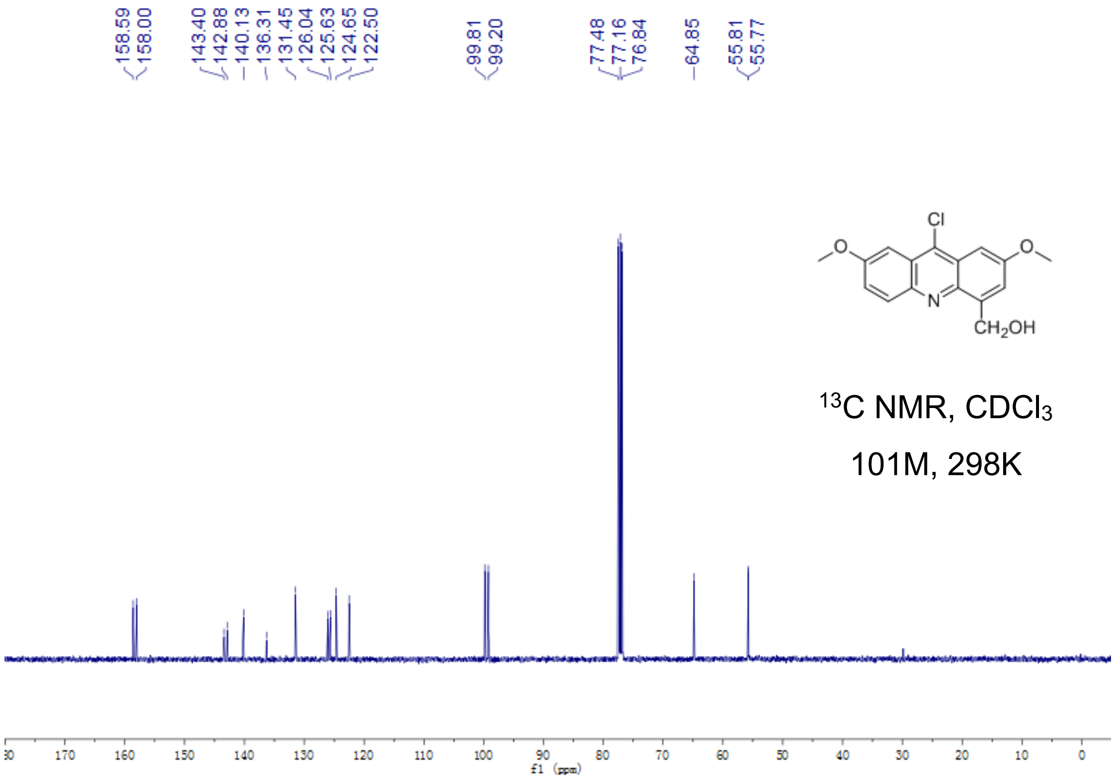

Fig S2.36. $^{13}$C-NMR of **10a**

**10b**

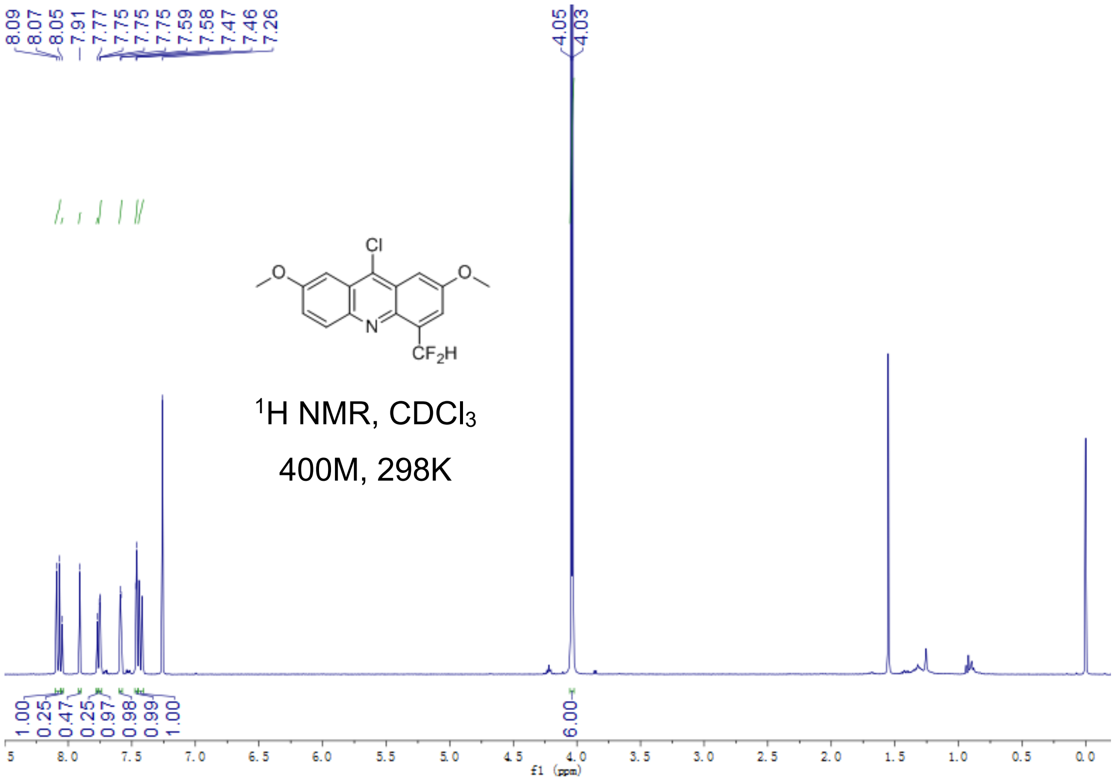


Fig S2.37. $^{1}$H-NMR of **10b**

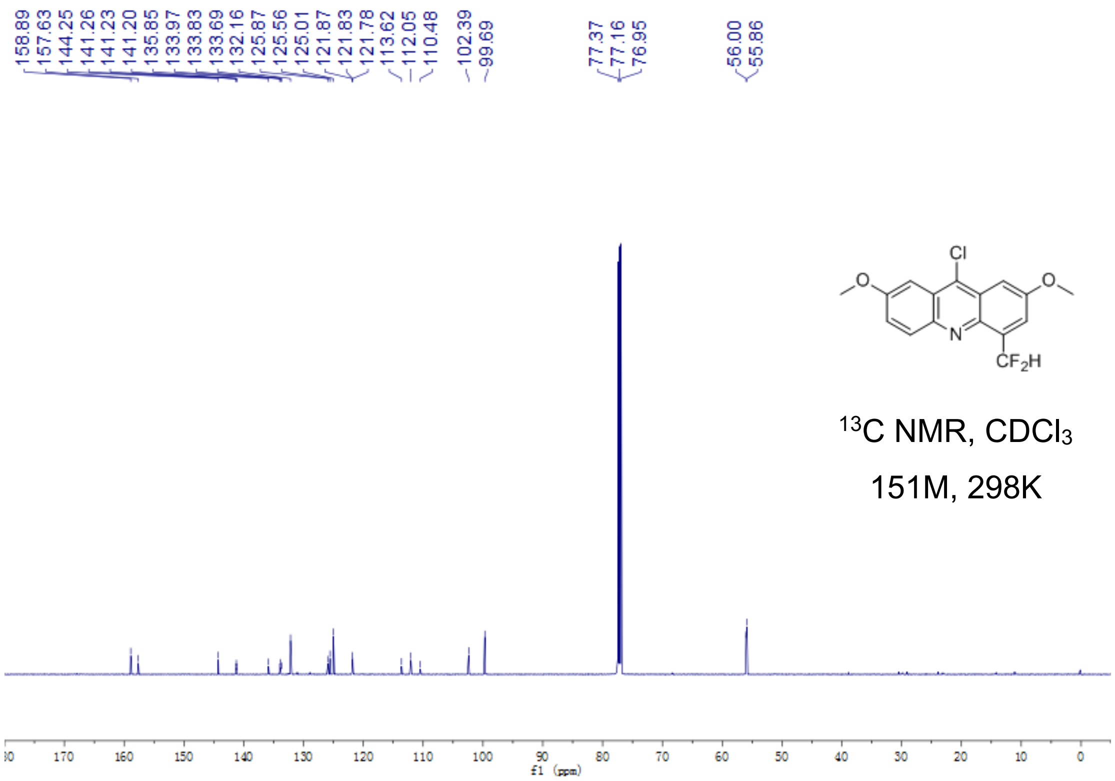


Fig S2.38. $^{13}$C-NMR of **10b**

**10**

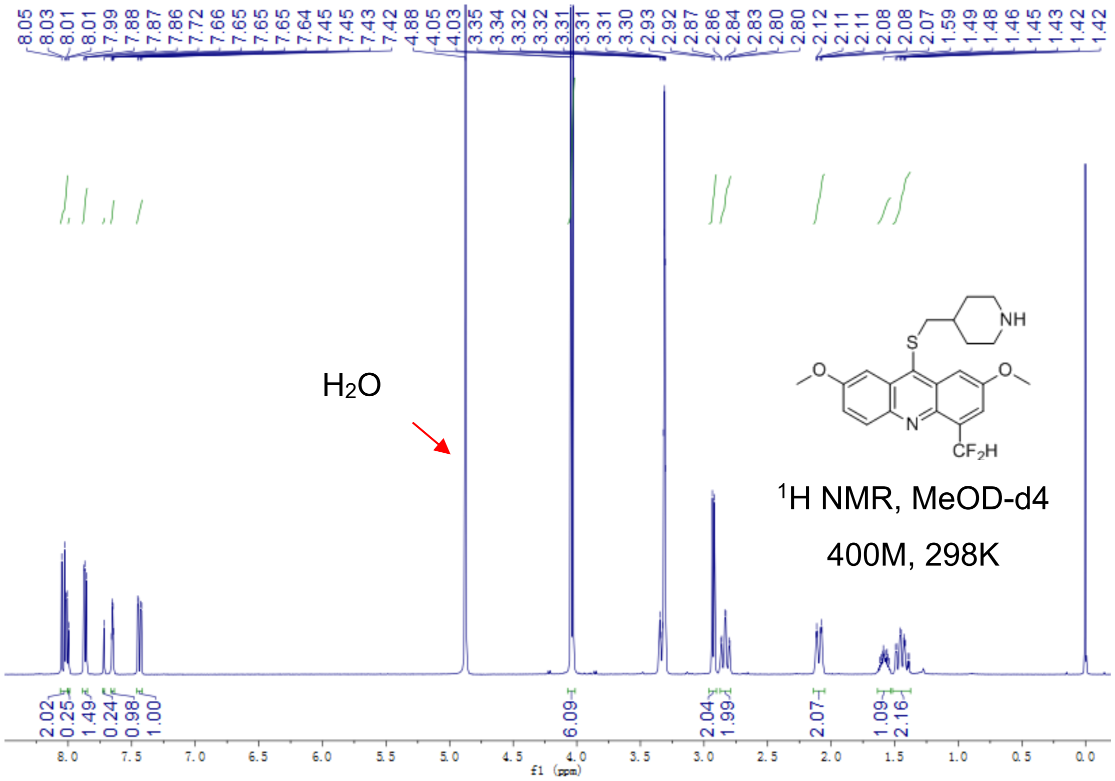


Fig S2.39. $^{1}$H-NMR of **10**

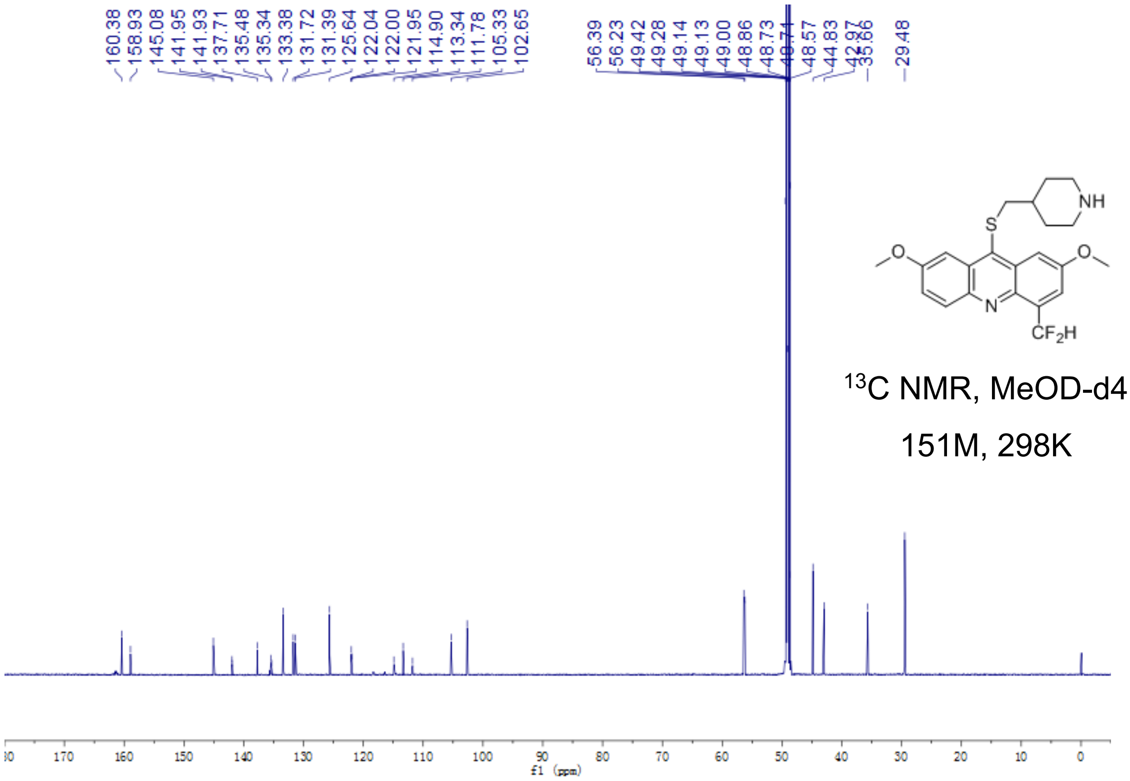


Fig S2.40. $^{13}$C-NMR of **10**

**11a**

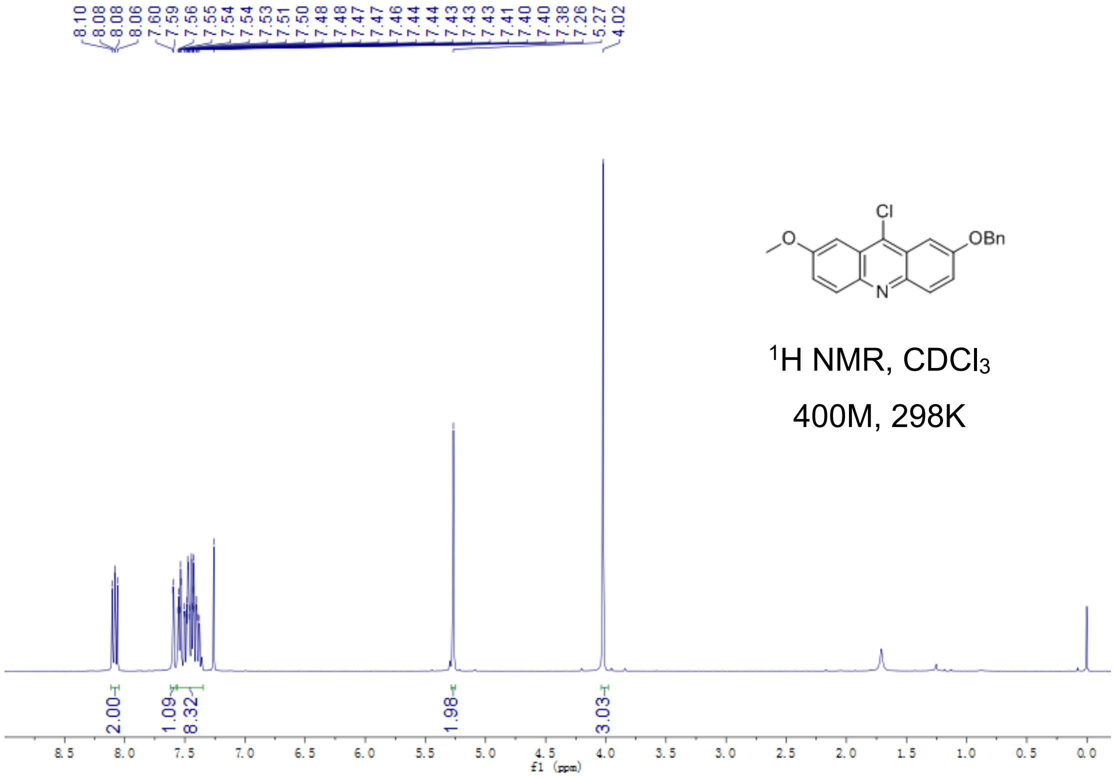

Fig S2.41. $^{1}$H-NMR of **11a**

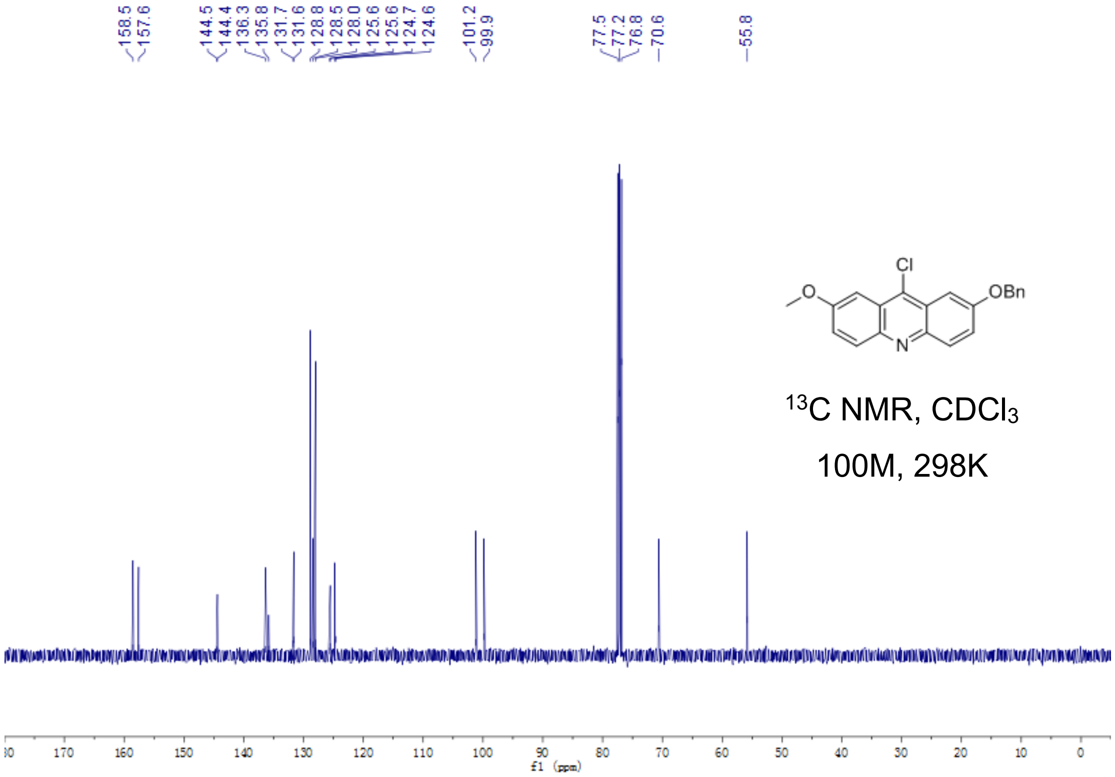


Fig S2.42. $^{13}$C-NMR of **11a**

**11**

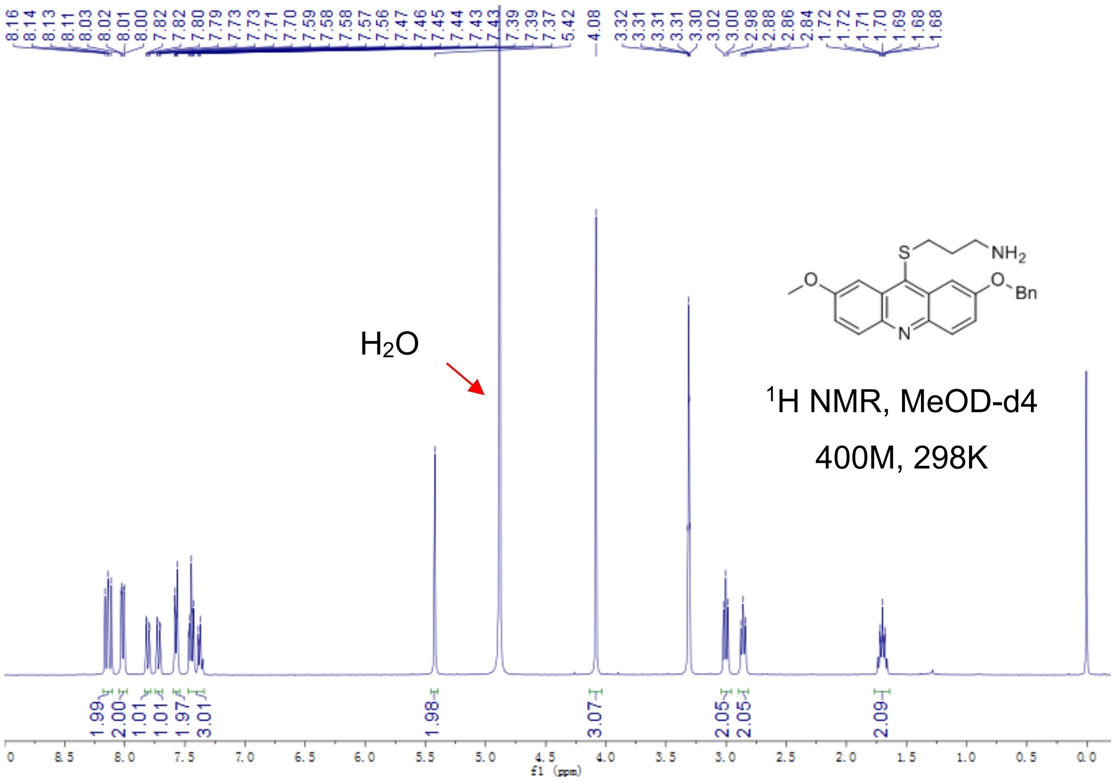


Fig S2.43. $^{1}$H-NMR of **11**

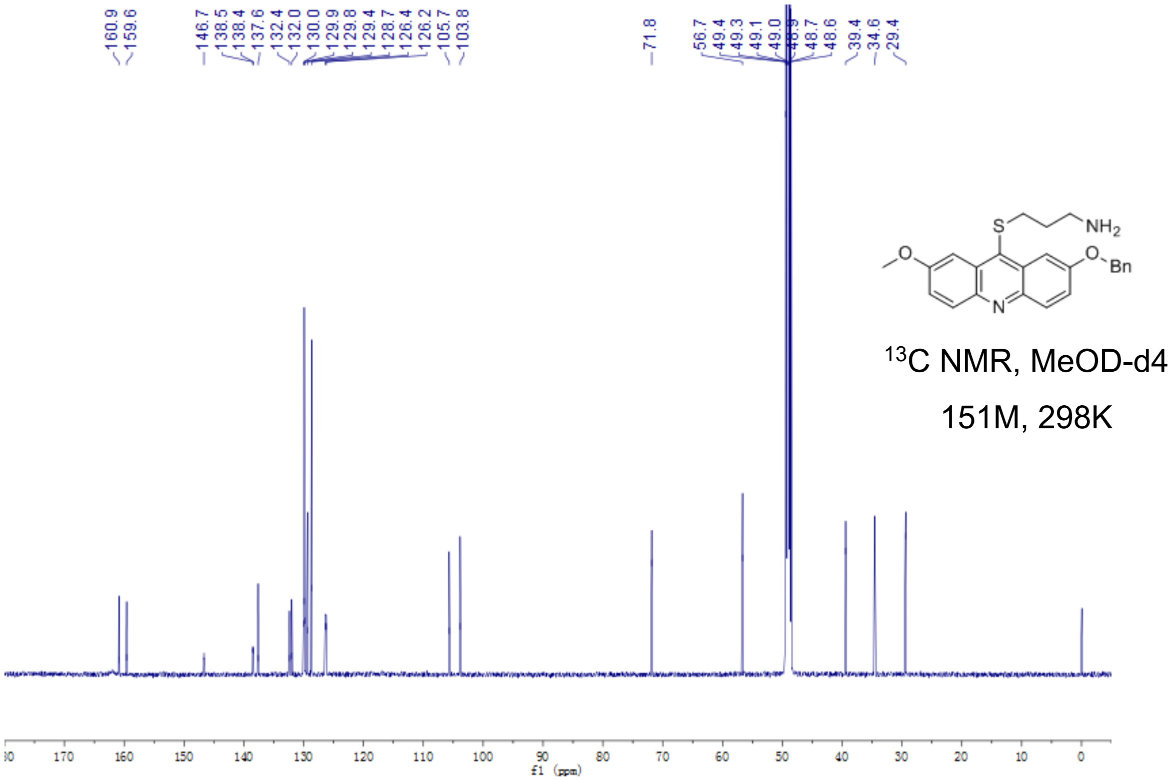


Fig S2.44. $^{13}$C-NMR of **11**

**12a**

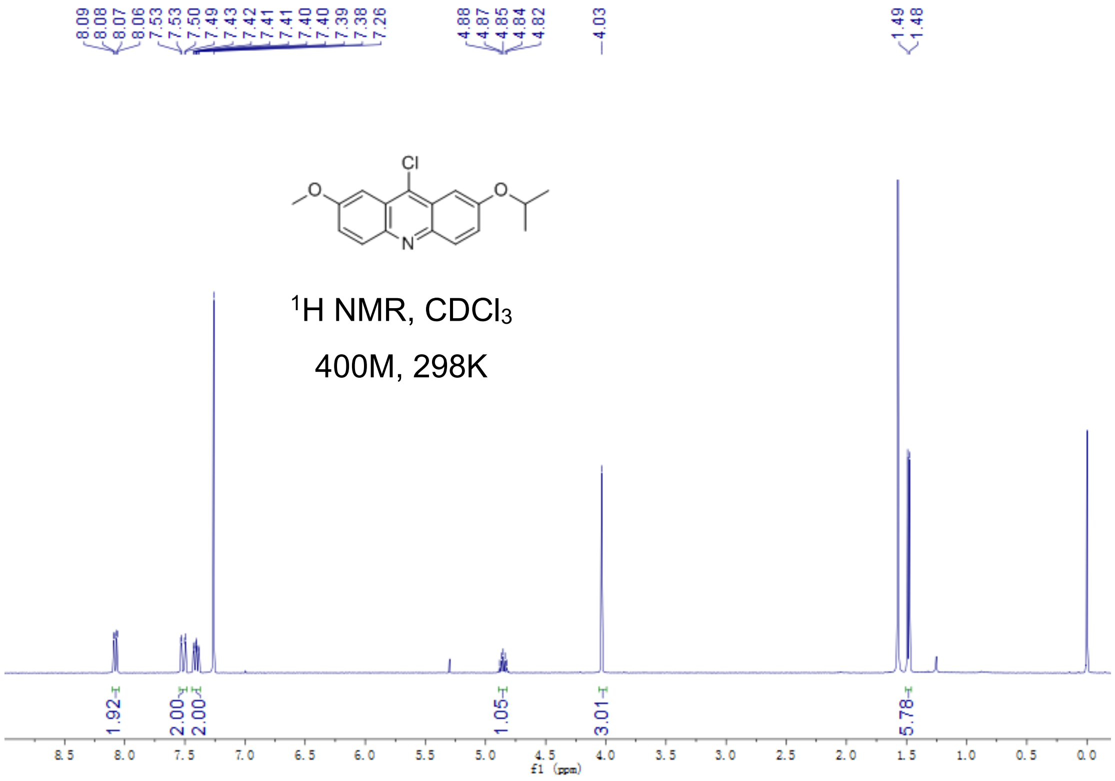


Fig S2.45. $^{1}$H-NMR of **12a**

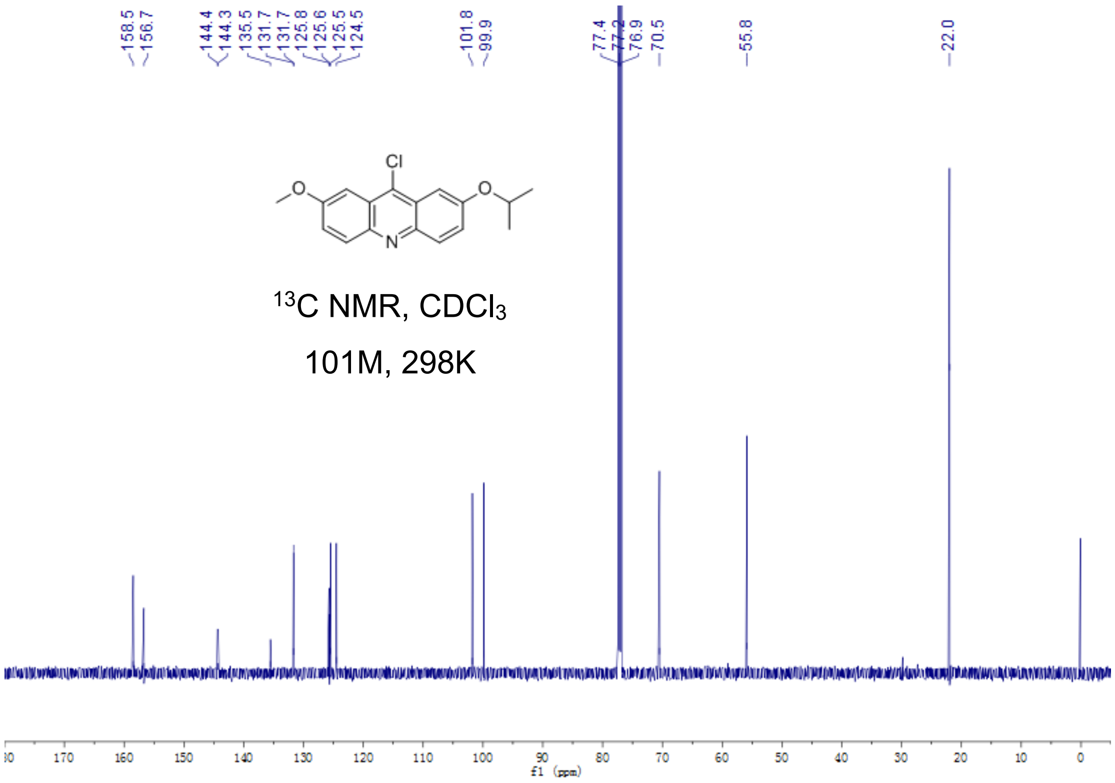


Fig S2.46. $^{13}$C-NMR of **12a**

**12**

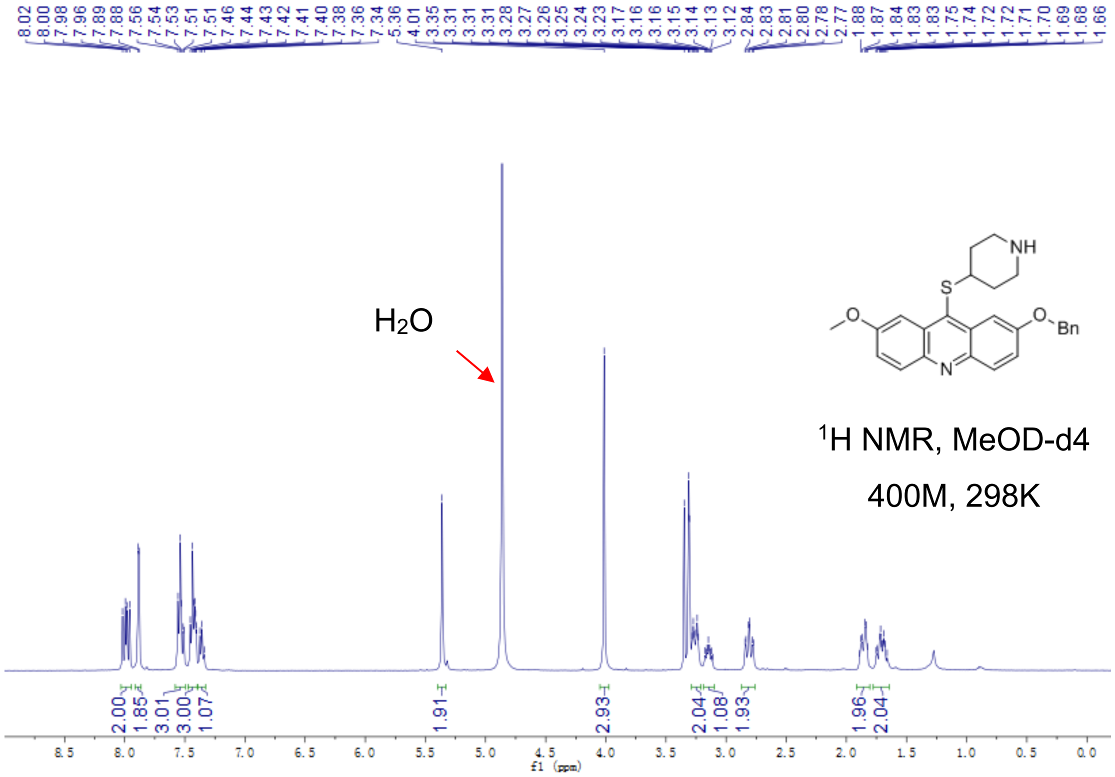


Fig S2.47. $^1$H-NMR of **12**

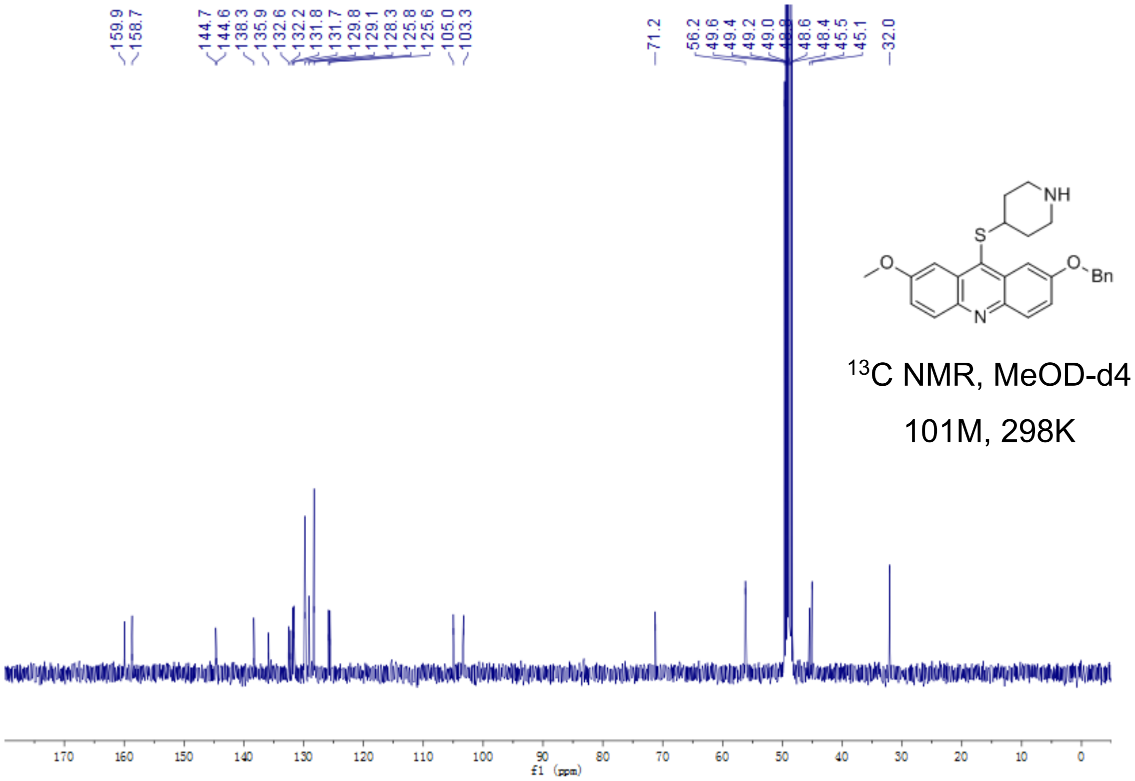


Fig S2.48. $^{13}$C-NMR of **12**

**13**

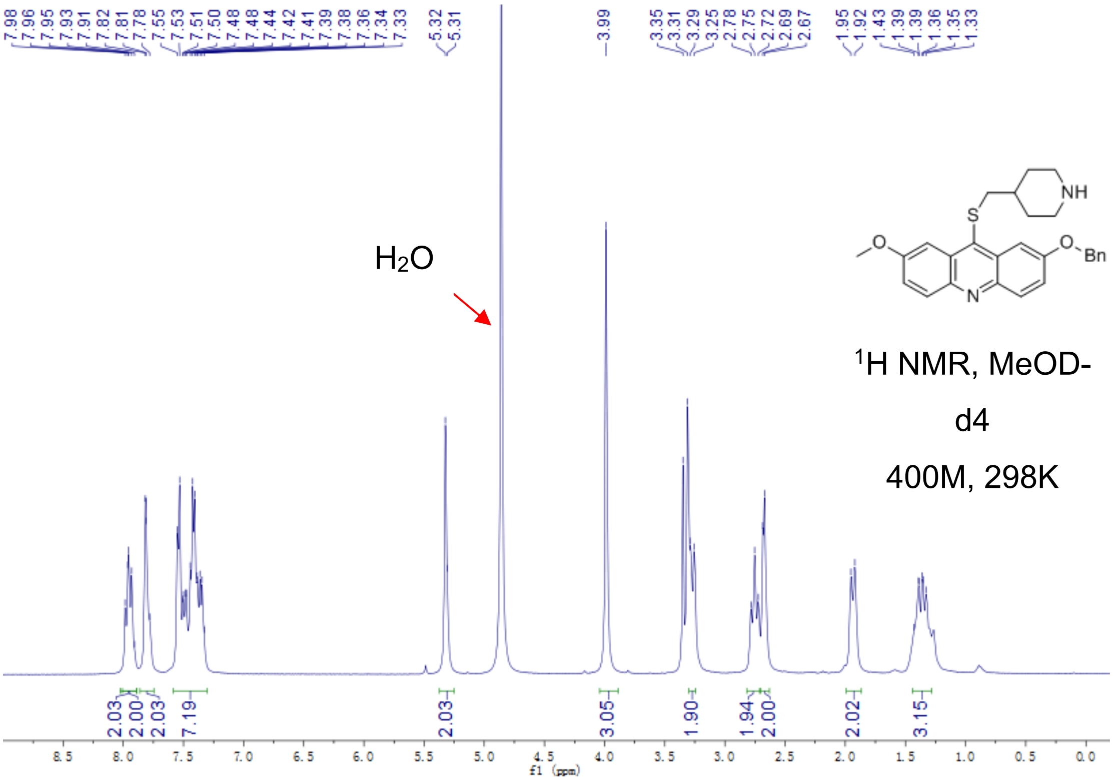


Fig S2.49. $^{1}$H-NMR of **13**

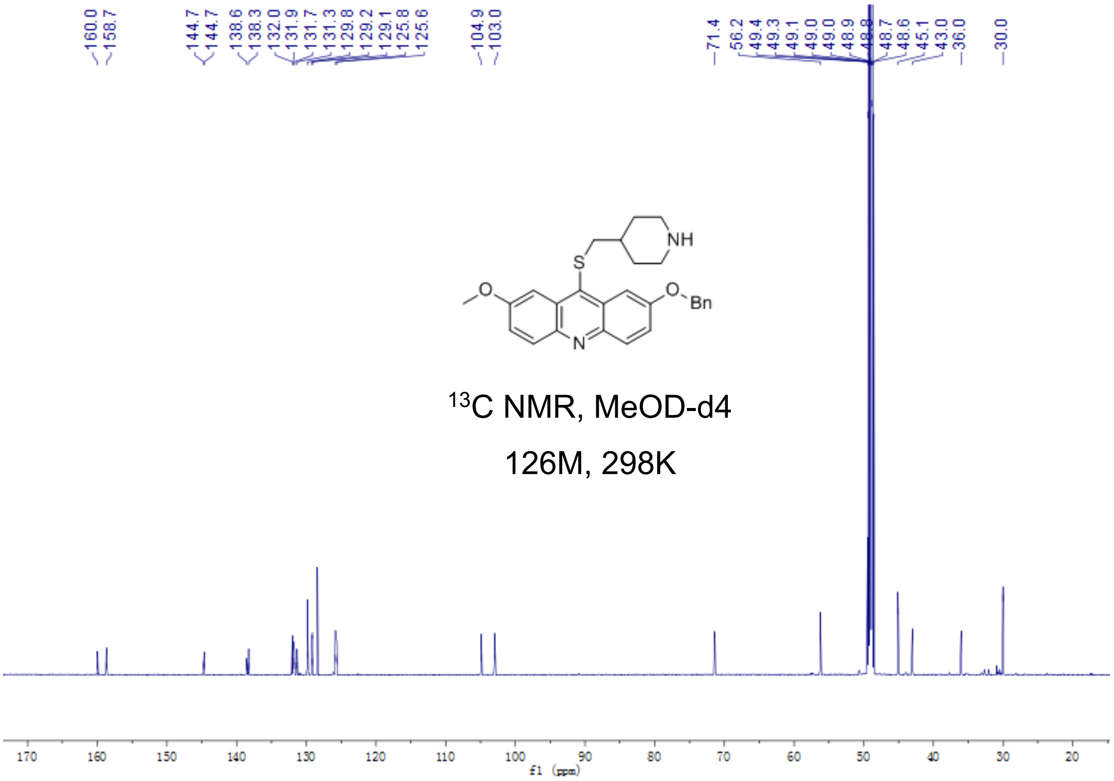

Fig S2.50. $^{13}$C-NMR of **13**

**14**

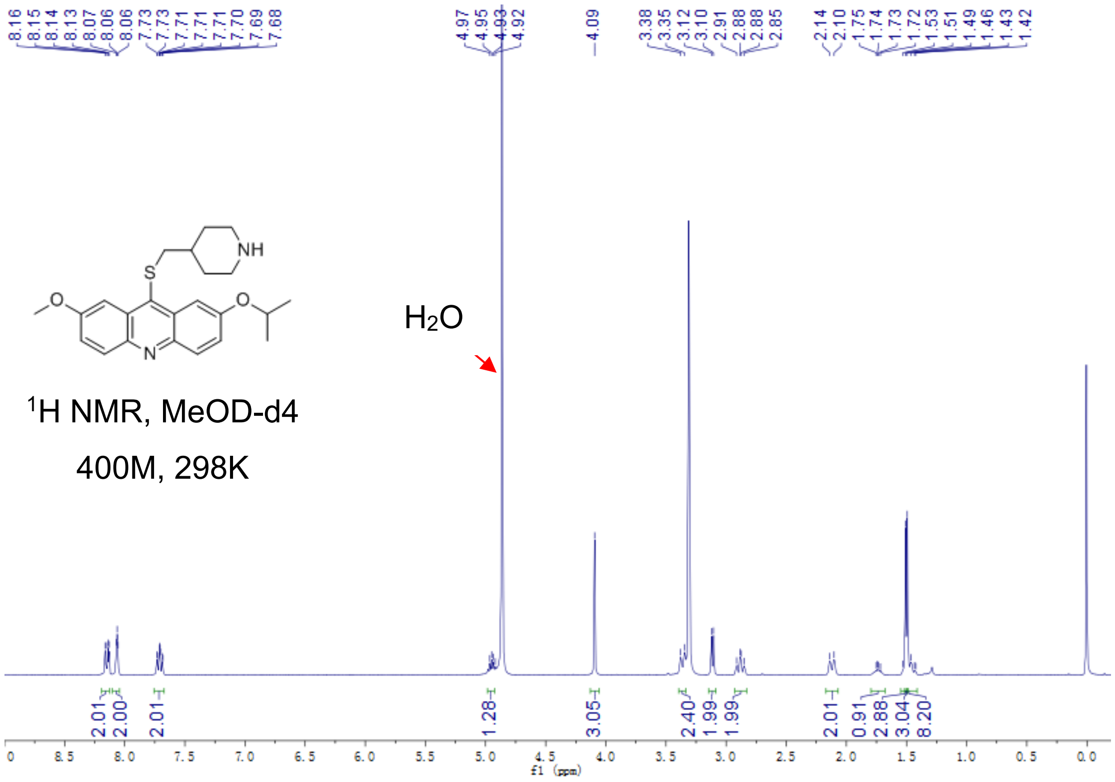


Fig S2.51. $^{1}$H-NMR of **14**

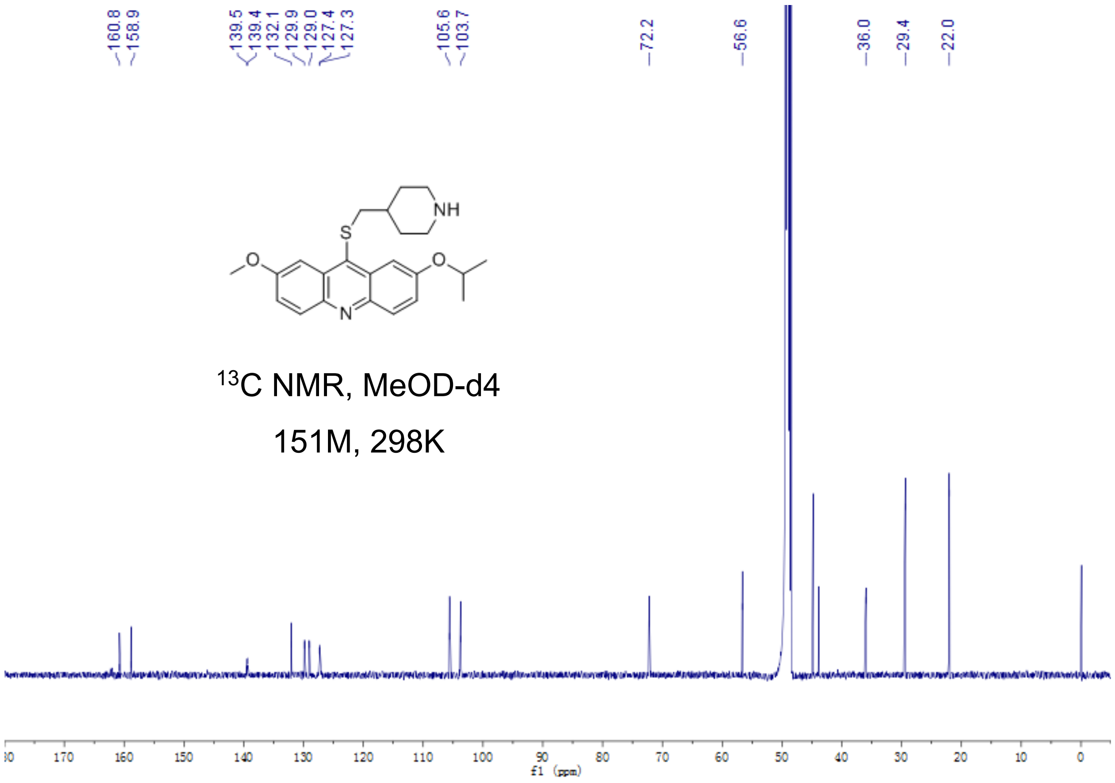


Fig S2.52. $^{13}$C-NMR of **14**

**15a**

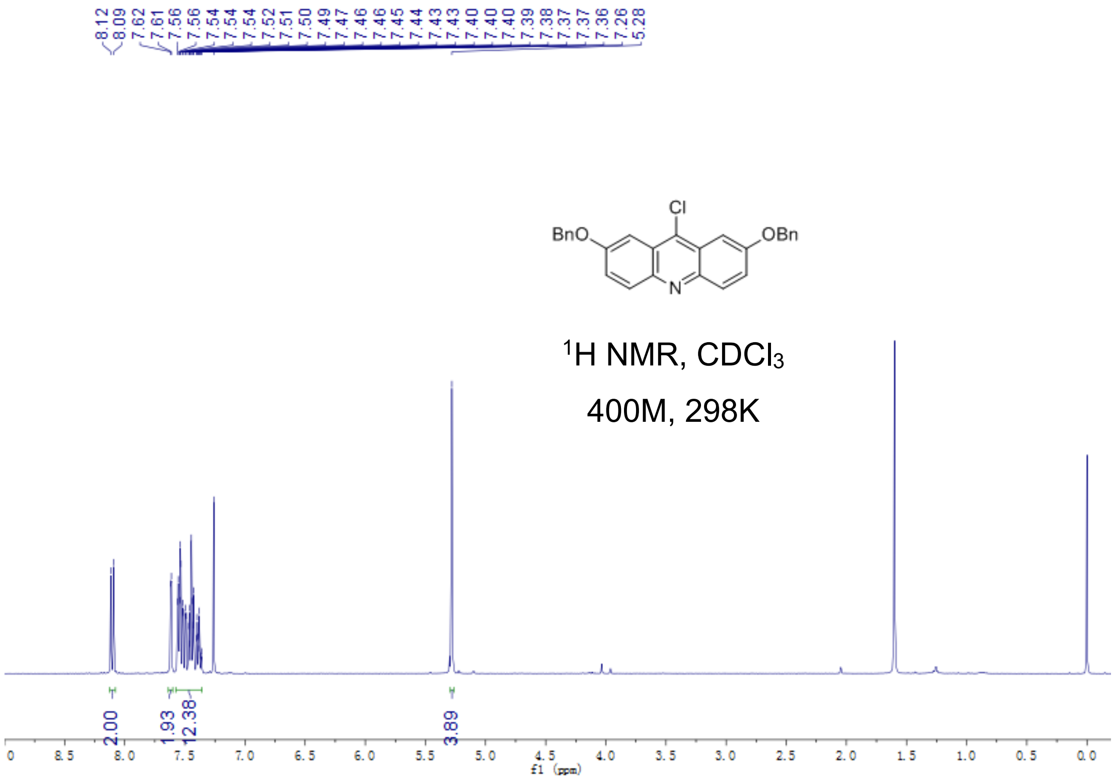


Fig S2.53. $^{1}$H-NMR of **15a**

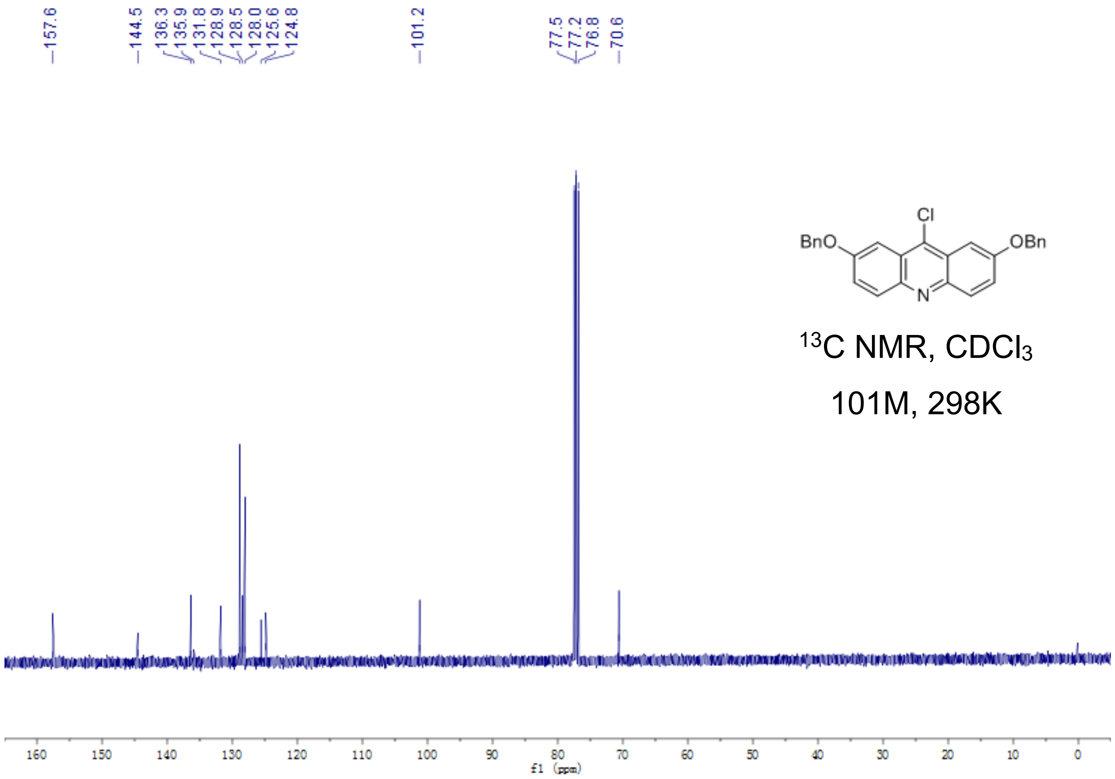


Fig S2.54. $^{13}$C-NMR of **15a**

**15**

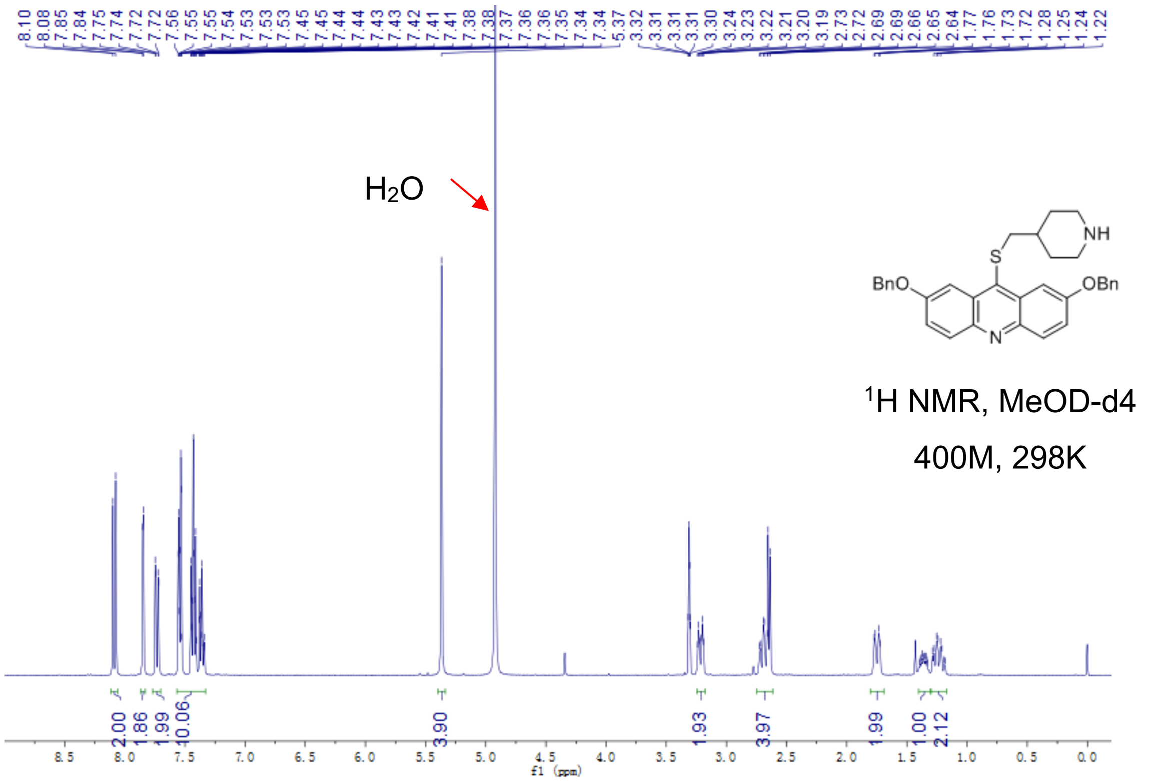


Fig S2.55. [1]H-NMR of **15**

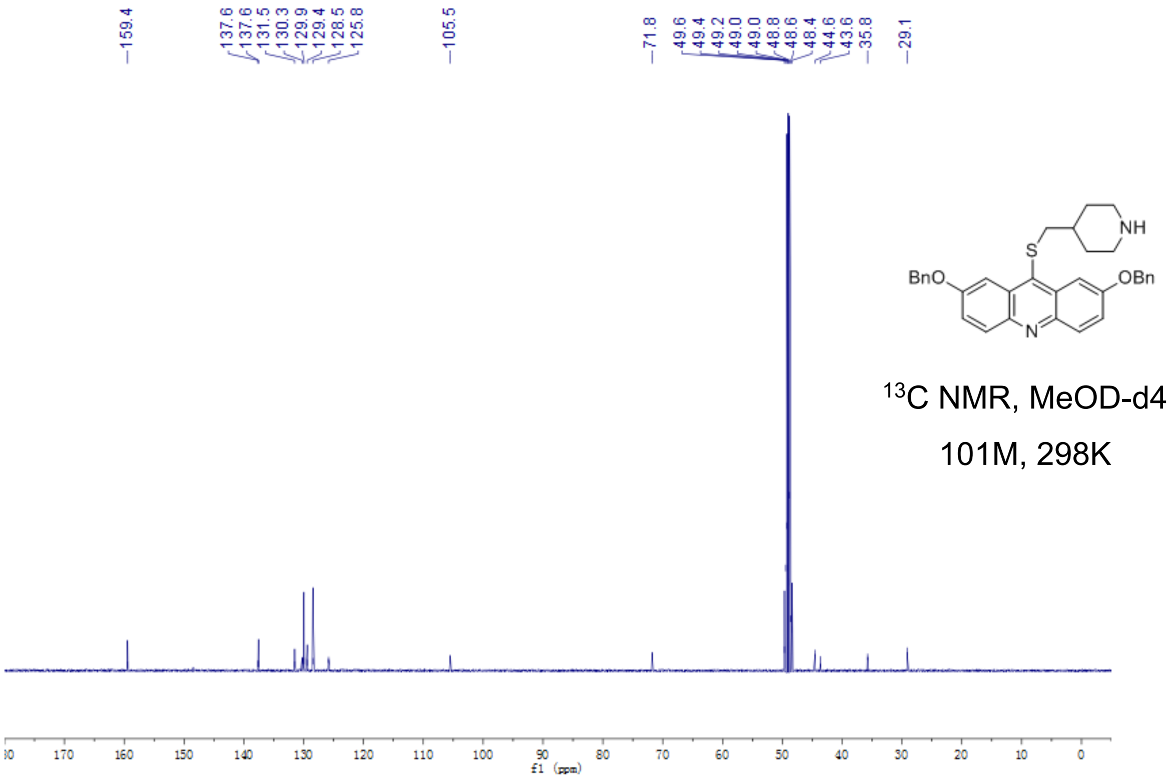


Fig S2.56. [13]C-NMR of **15**

**16a**

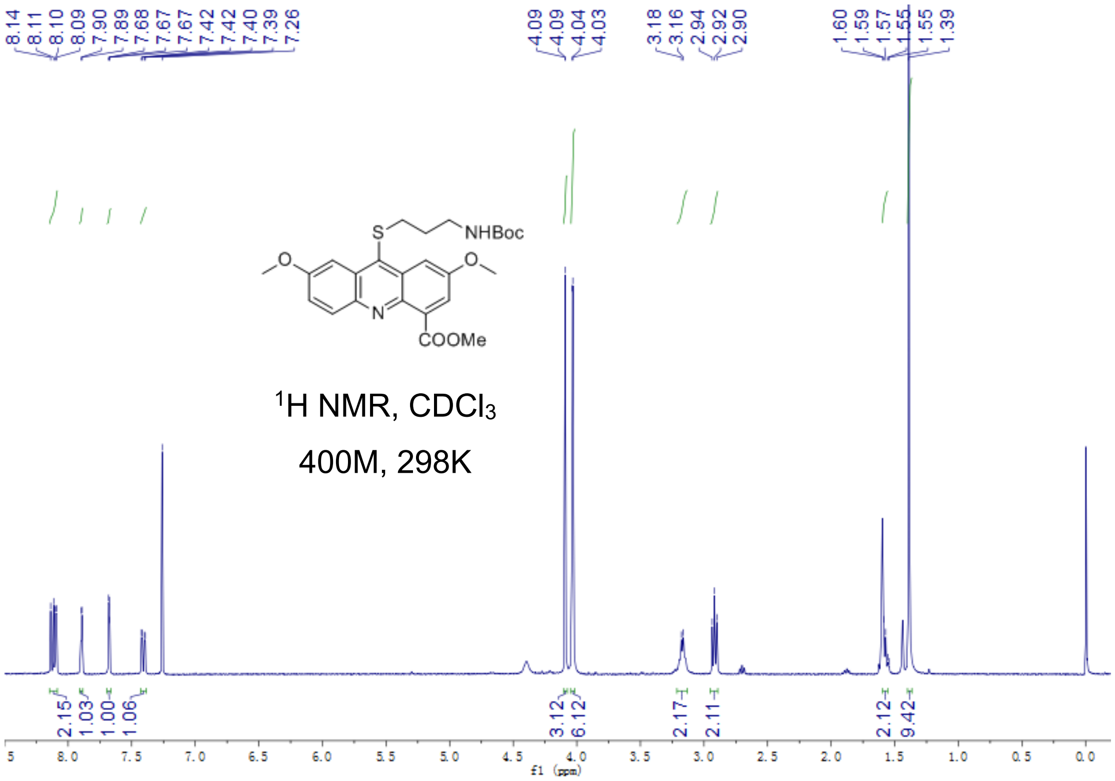


Fig S2.57. $^{1}$H-NMR of **16a**

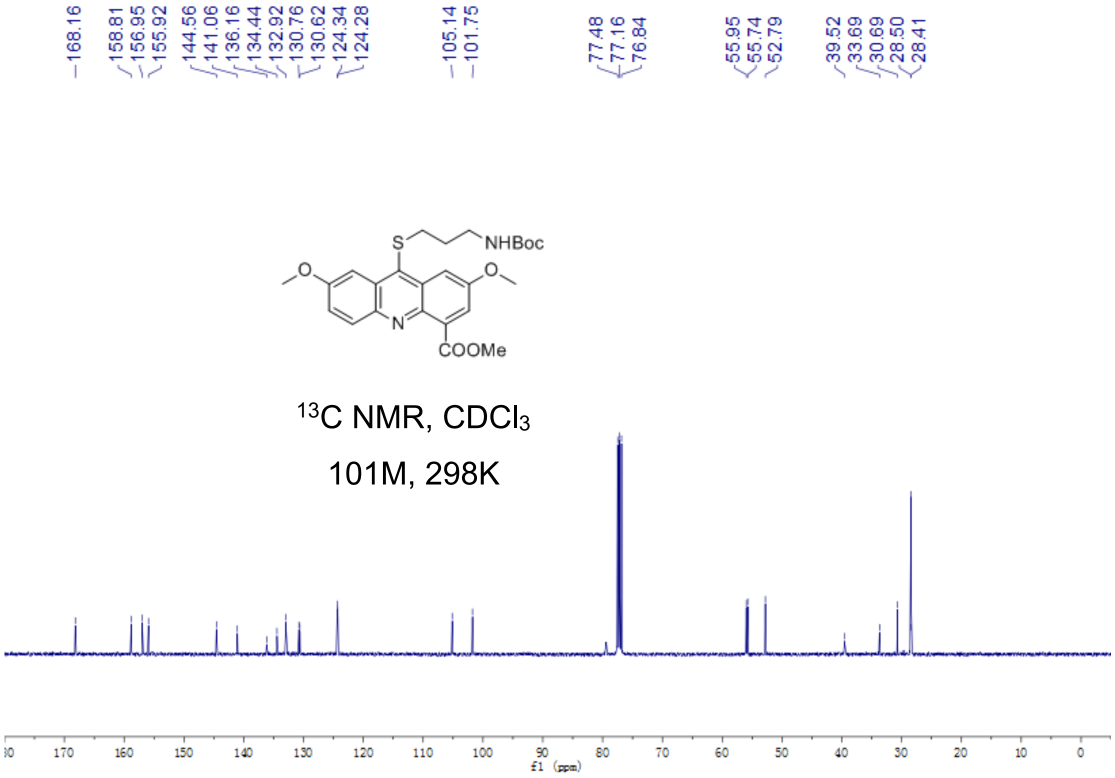


Fig S2.58. $^{13}$C-NMR of **16a**

**16b**

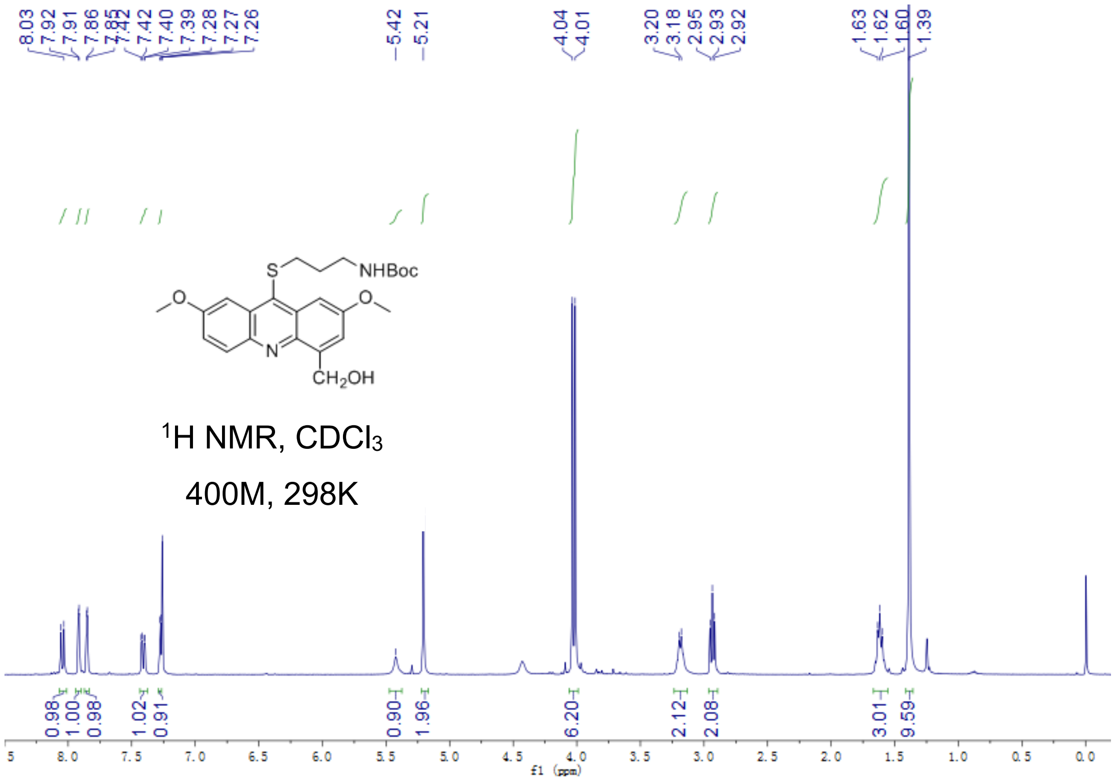


Fig S2.59. $^1$H-NMR of **16b**

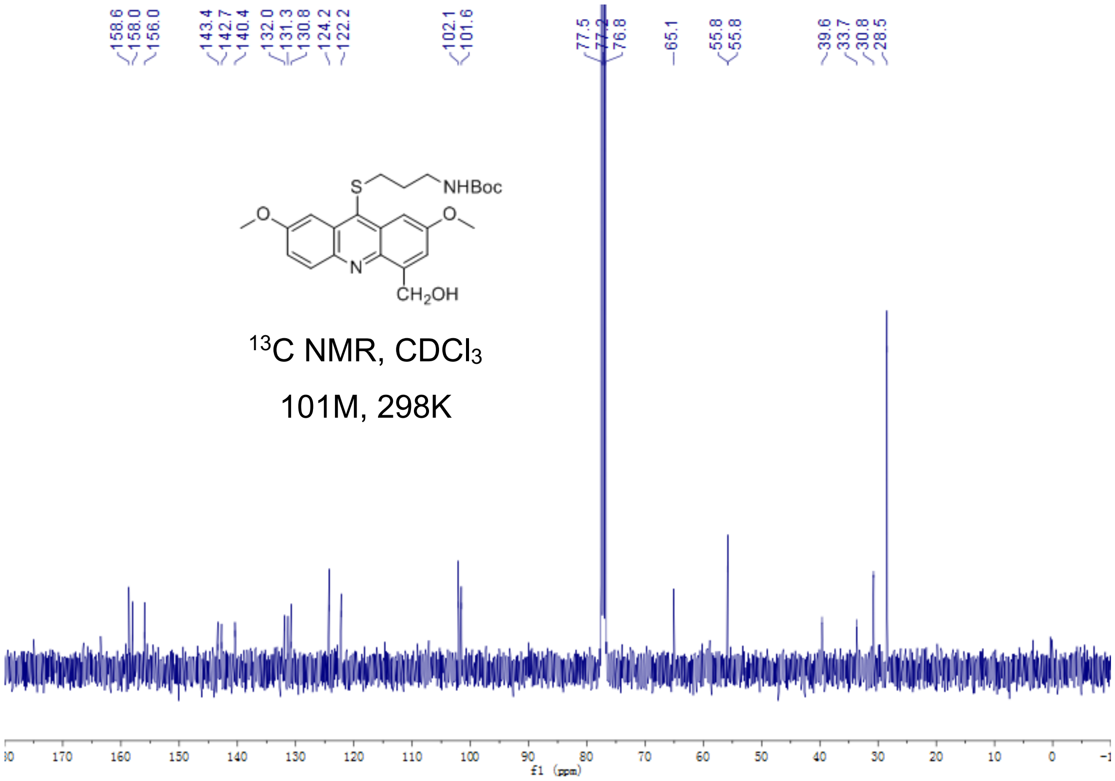


Fig S2.60. $^{13}$C-NMR of **16b**

**16**

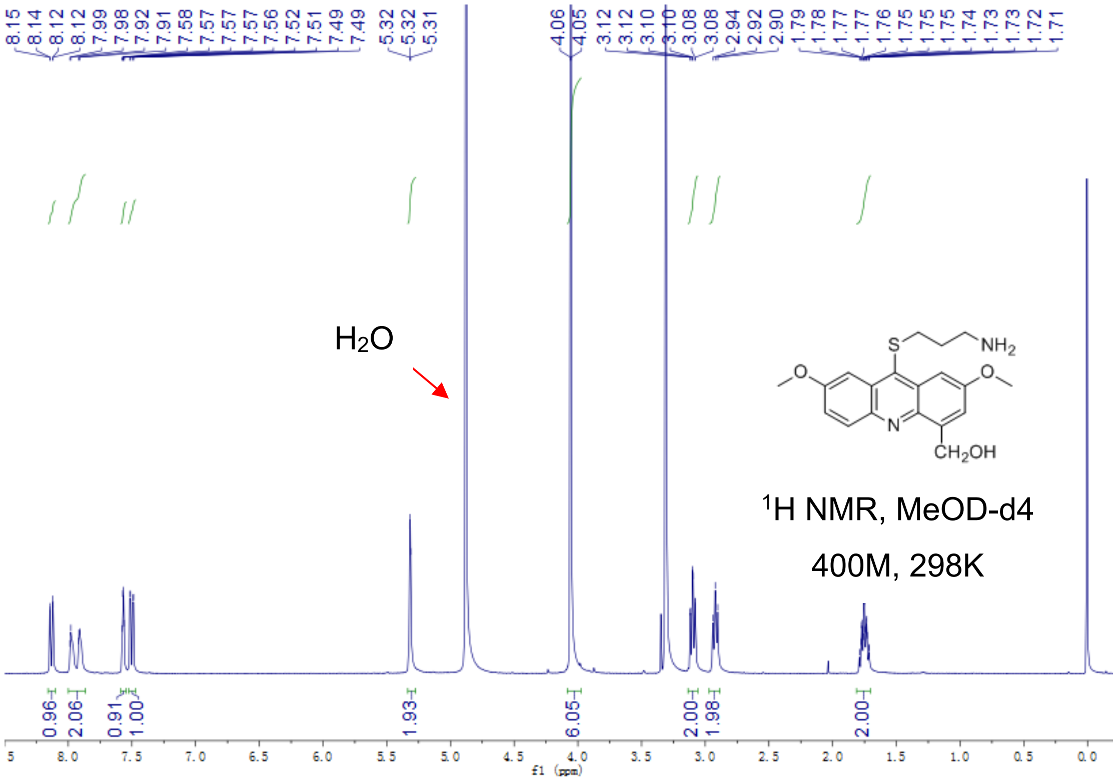


Fig S2.61. $^{1}$H-NMR of **16**

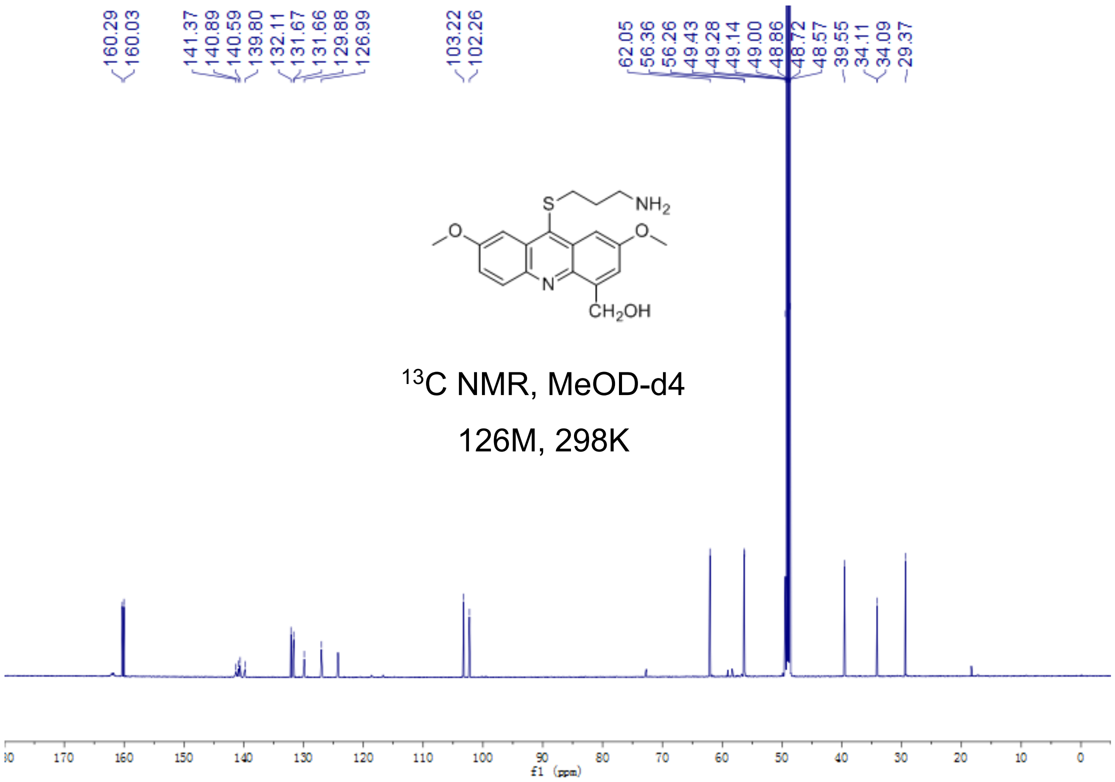

Fig S2.62. $^{13}$C-NMR of **16**

**17a**

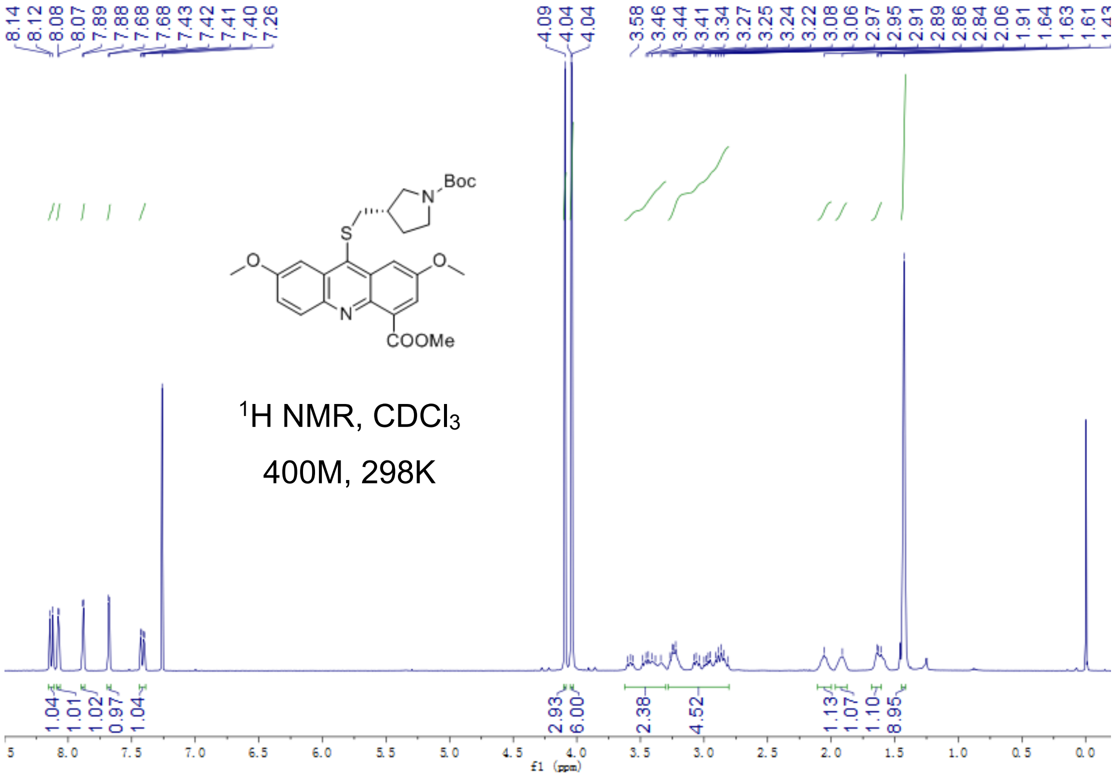


Fig S2.63. $^{1}$H-NMR of **17a**

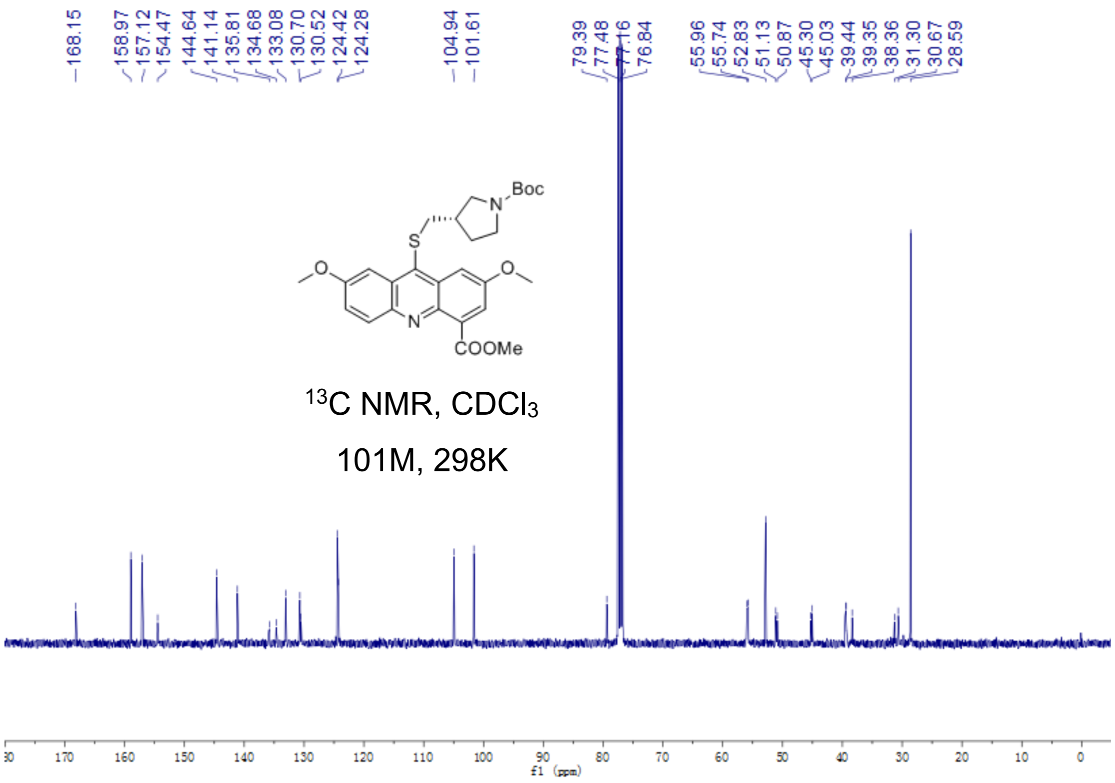


Fig S2.64. $^{13}$C-NMR of **17a**

**17b**

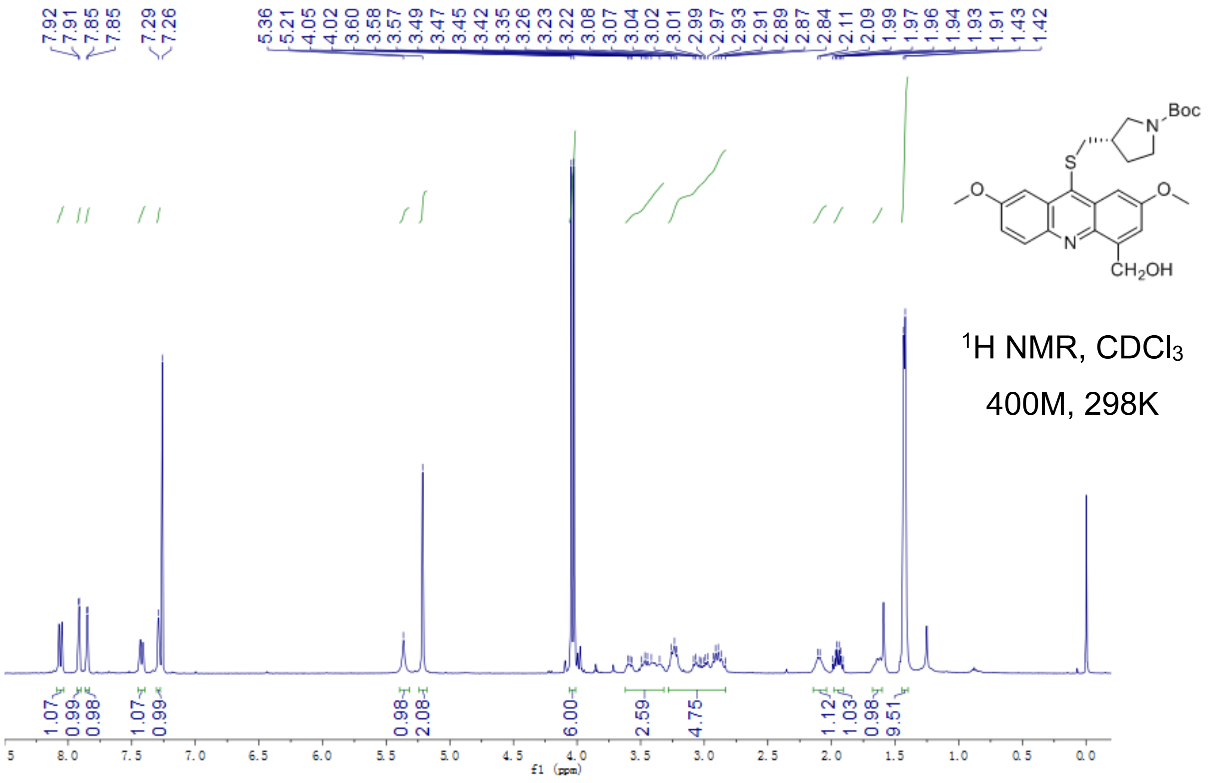


Fig S2.65. $^{1}$H-NMR of **17b**

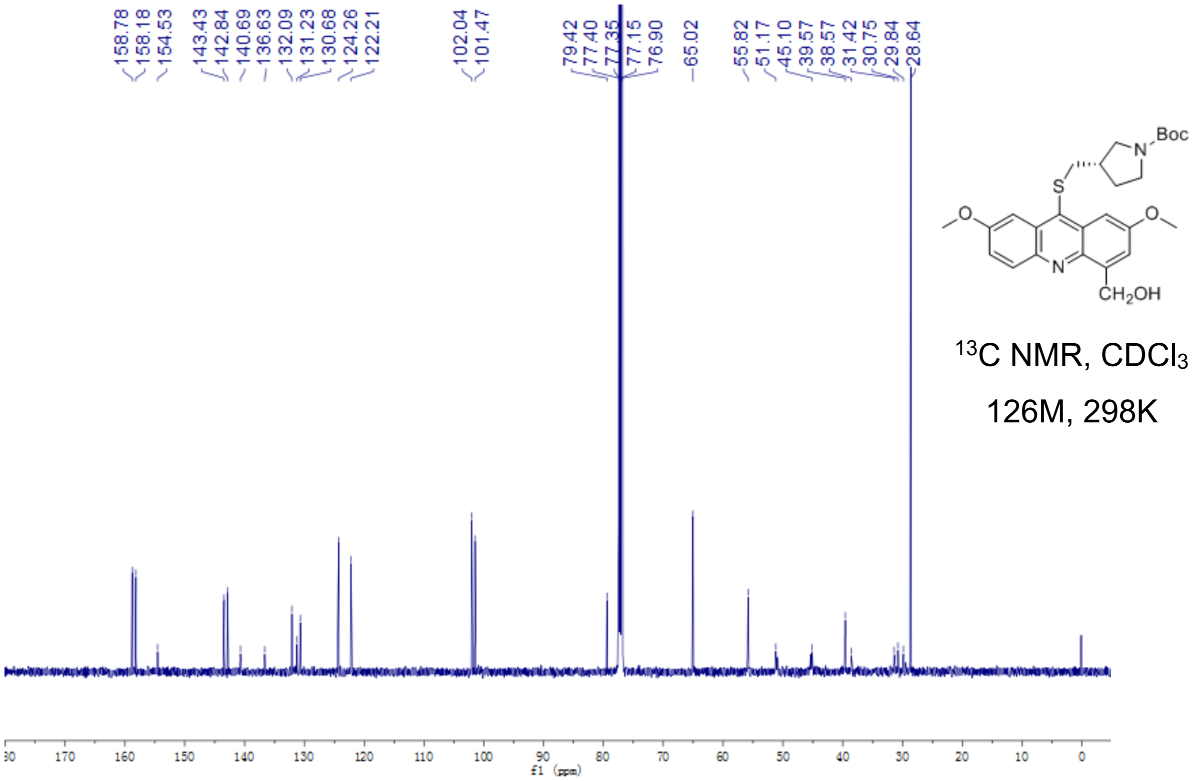


Fig S2.66. $^{13}$C-NMR of **17b**

**17**

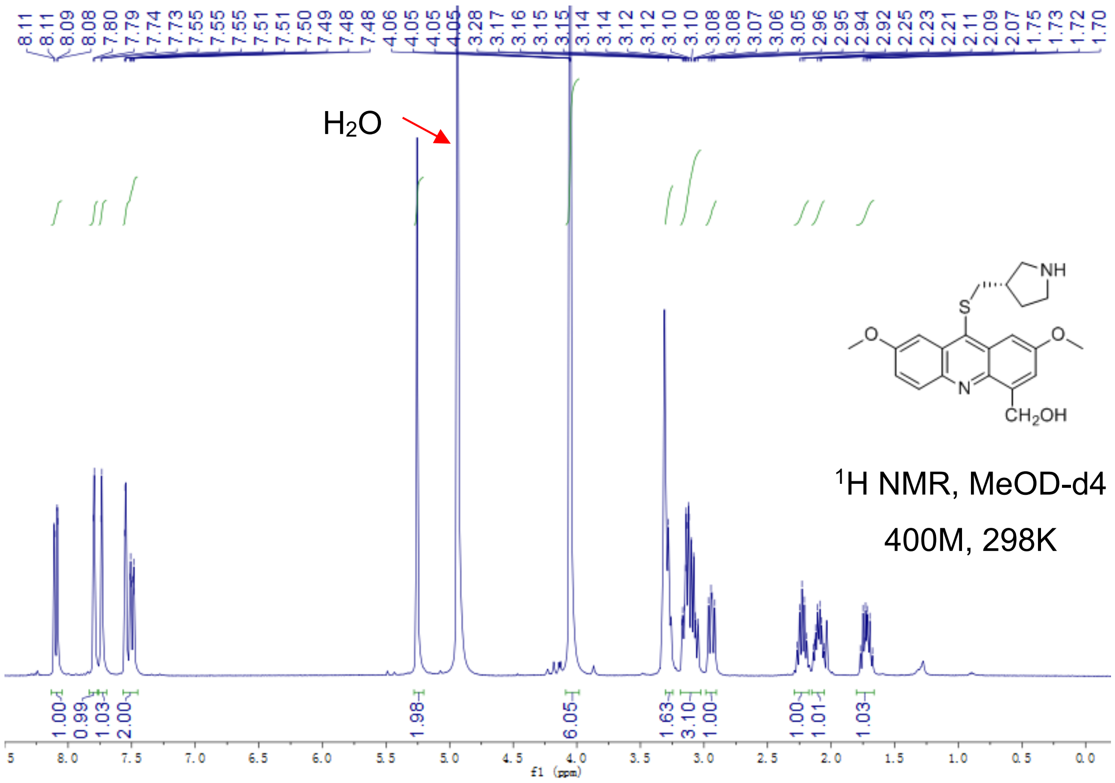

Fig S2.67. $^{1}$H-NMR of **17**

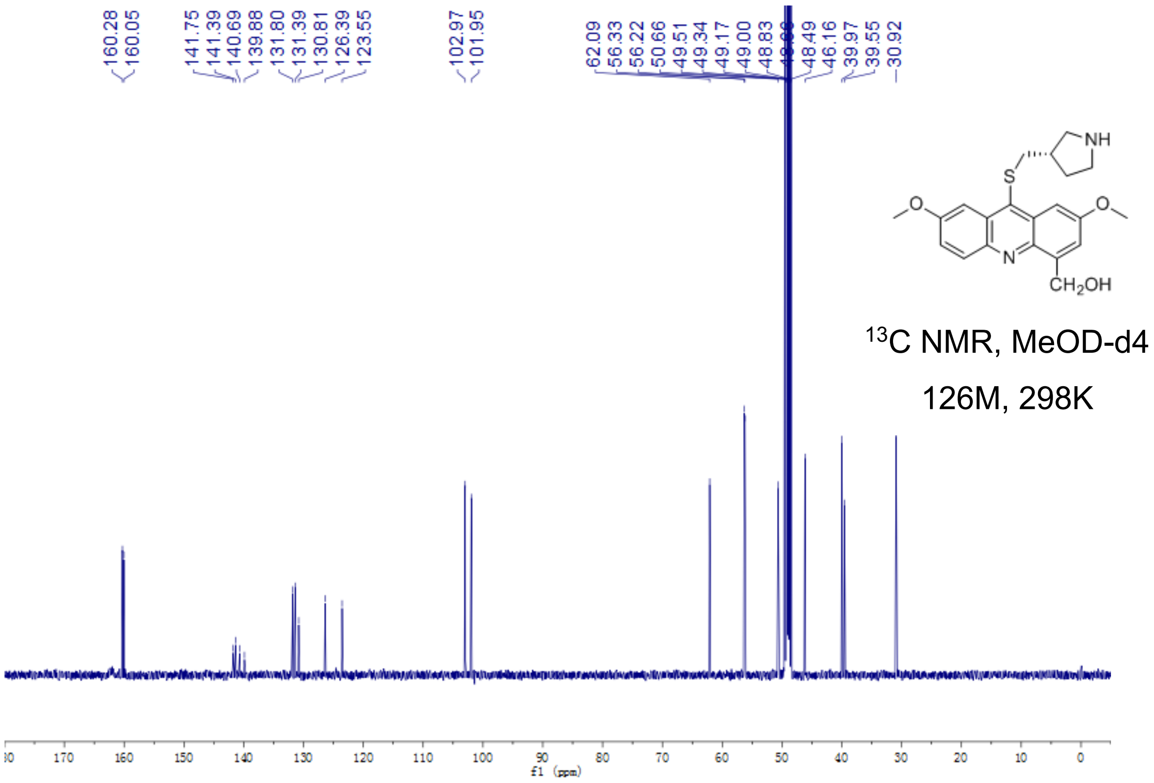


Fig S2.68. $^{13}$C-NMR of **17**

**18a**

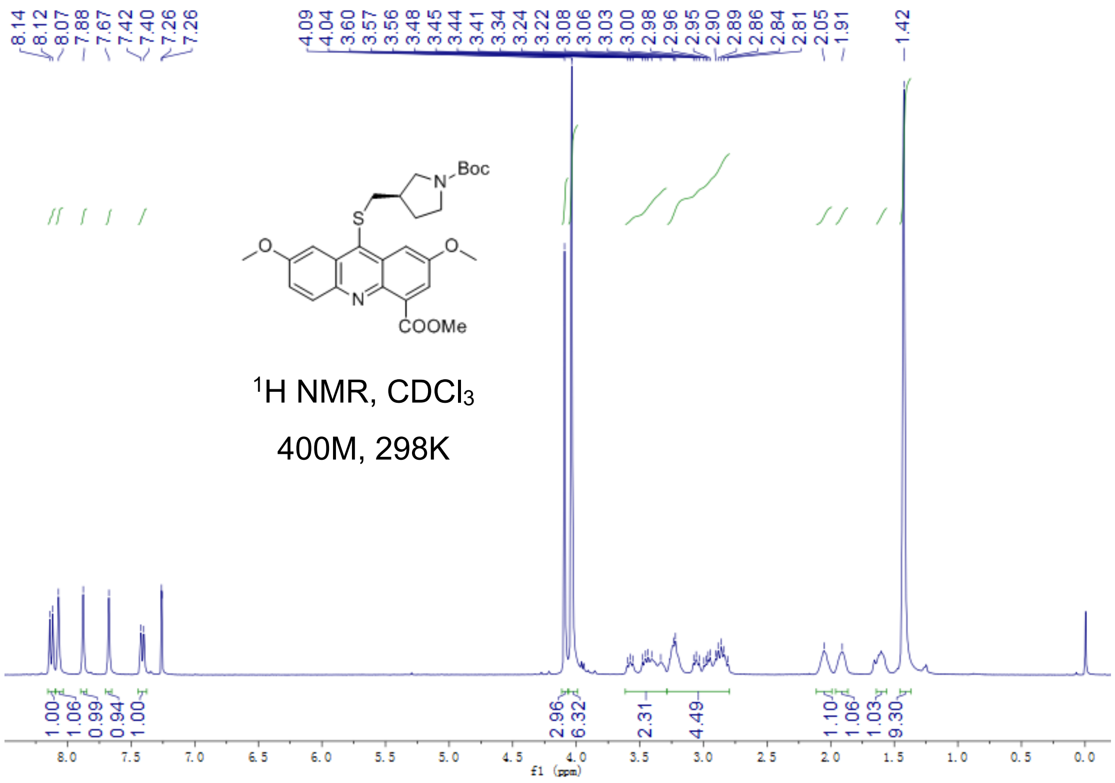


Fig S2.69. $^1$H-NMR of **18a**

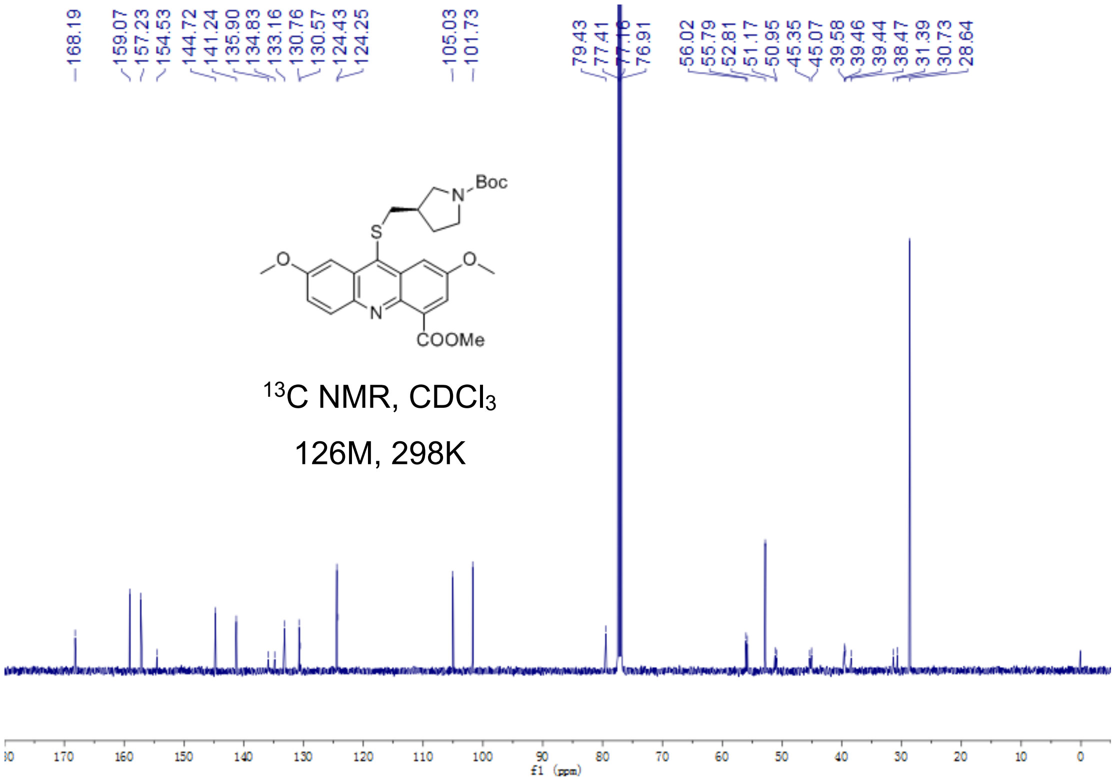


Fig S2.70. $^{13}$C-NMR of **18a**

**18b**

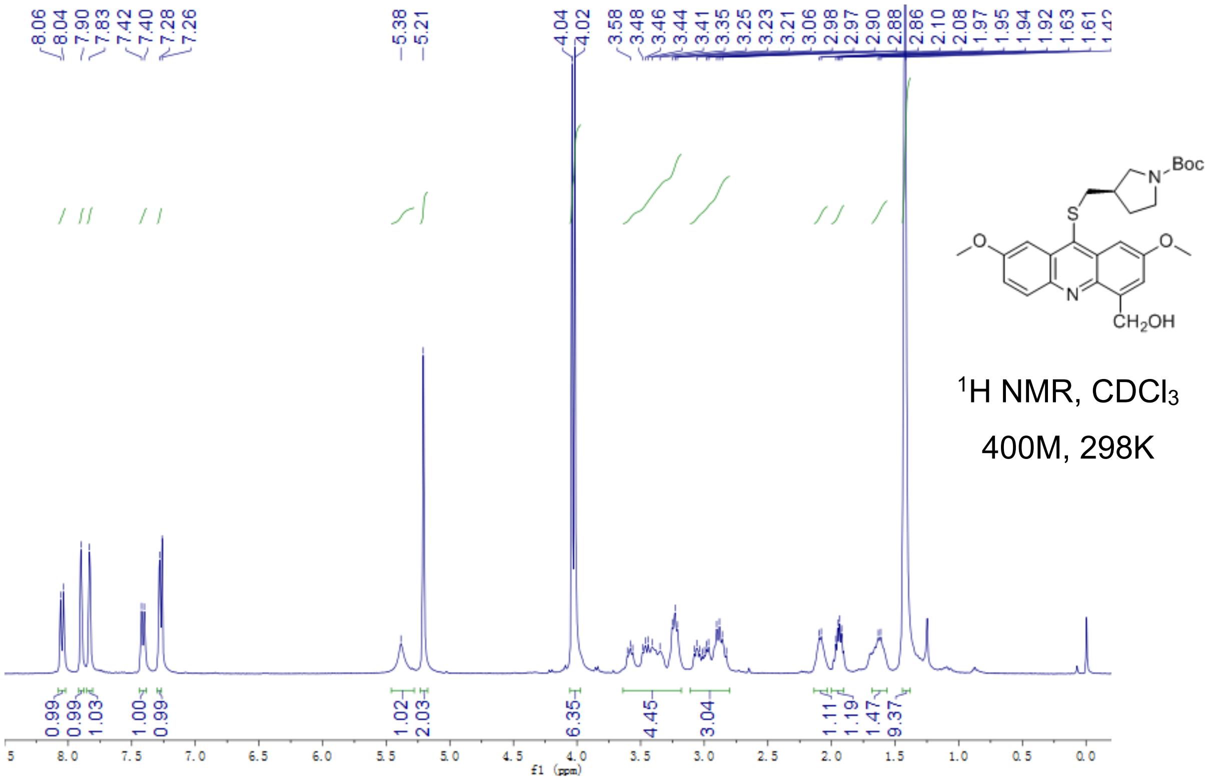


Fig S2.71. $^{1}$H-NMR of **18b**

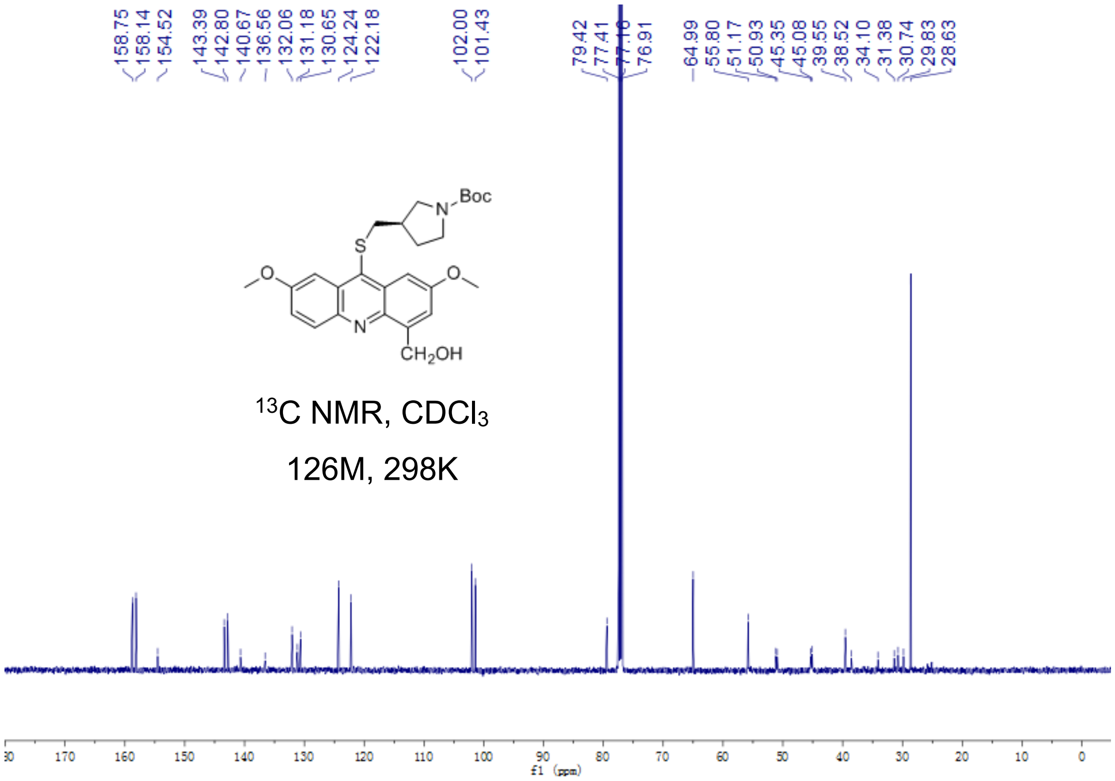


Fig S2.72. $^{13}$C-NMR of **18b**

**18**

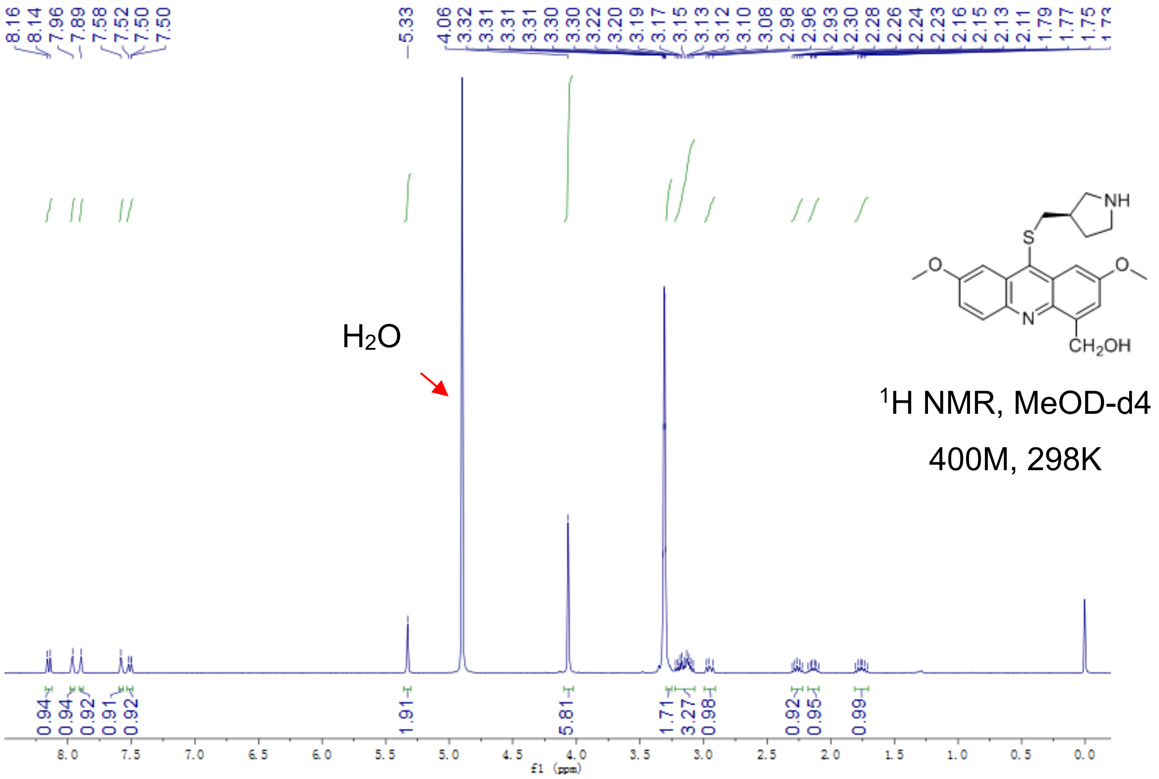


Fig S2.73. ¹H-NMR of **18**

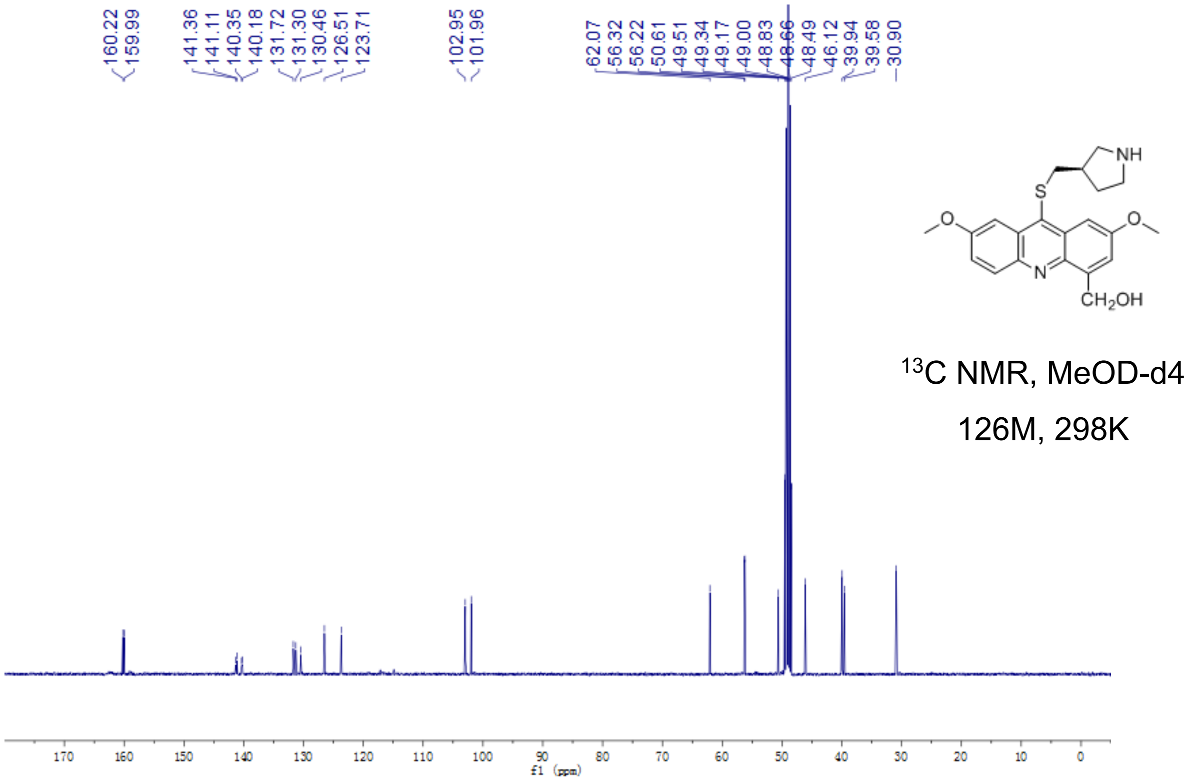


Fig S2.74. ¹³C-NMR of **18**

**19a**

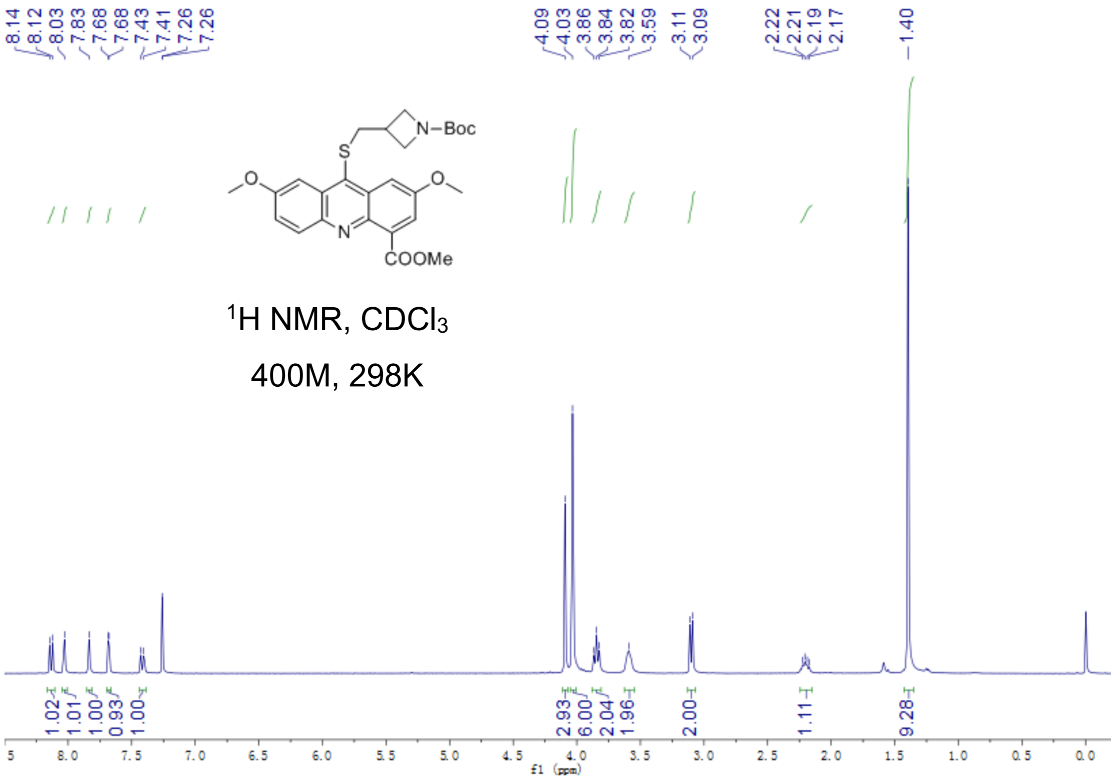


Fig S2.75. $^{1}$H-NMR of **19a**

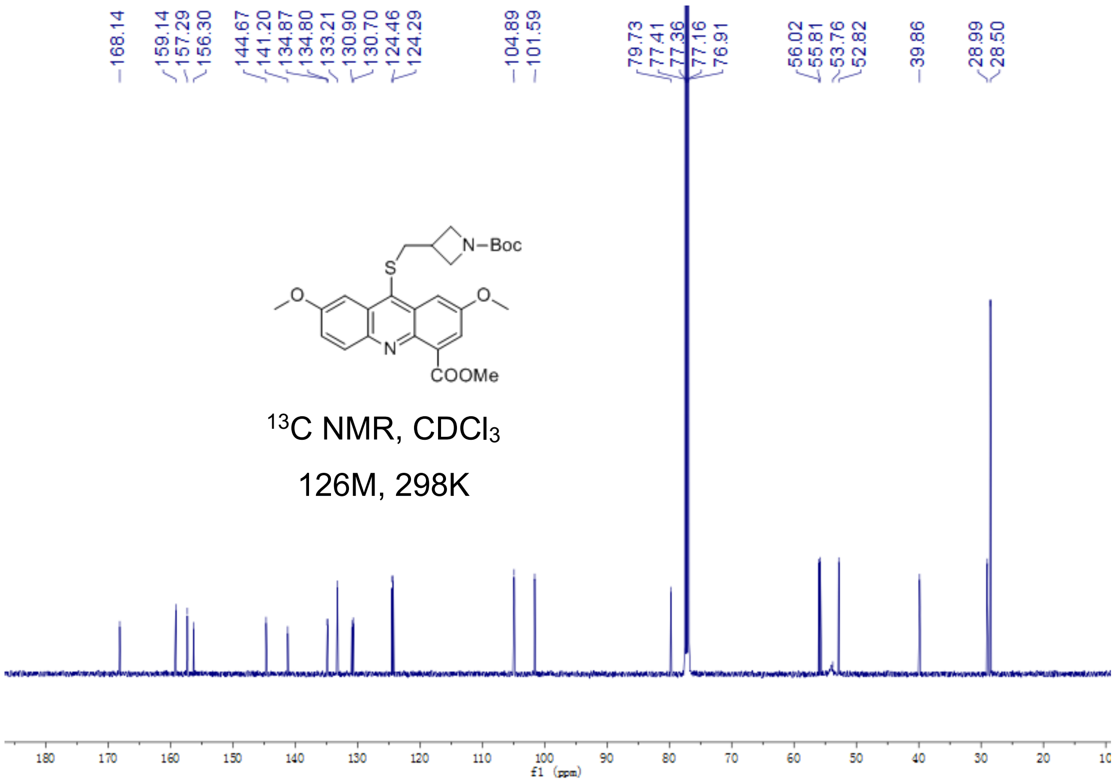

Fig S2.76. $^{13}$C-NMR of **19a**

**19b**

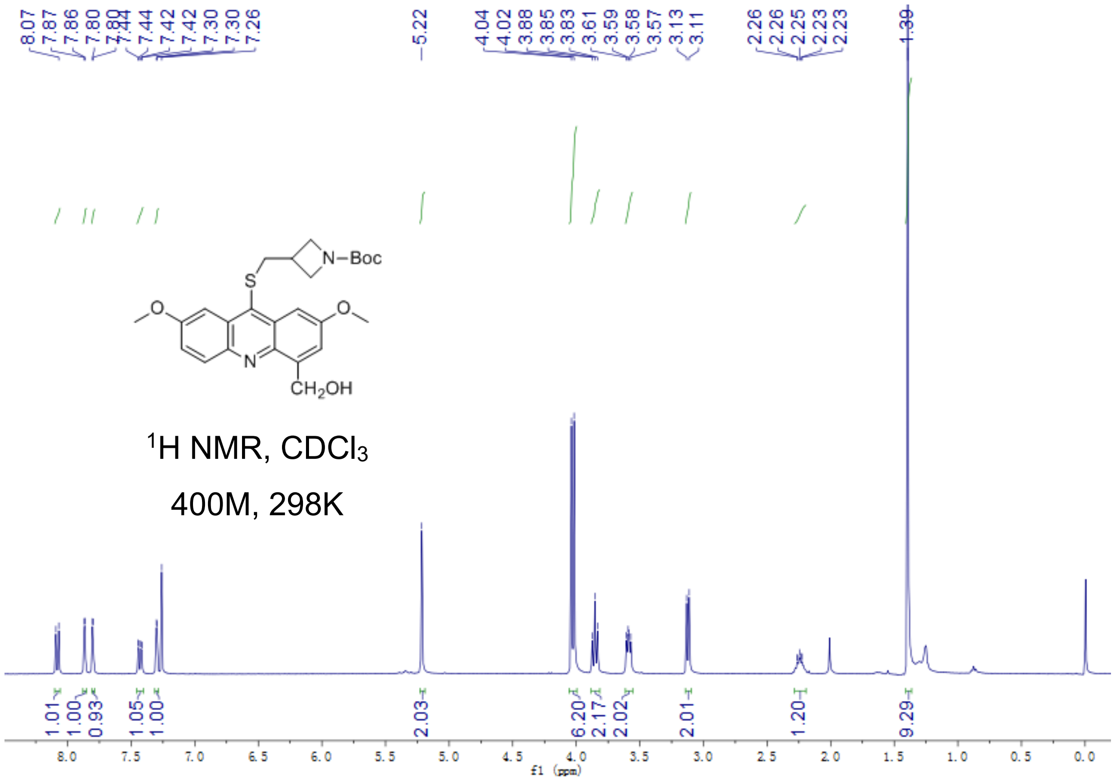


Fig S2.77. $^{1}$H-NMR of **19b**

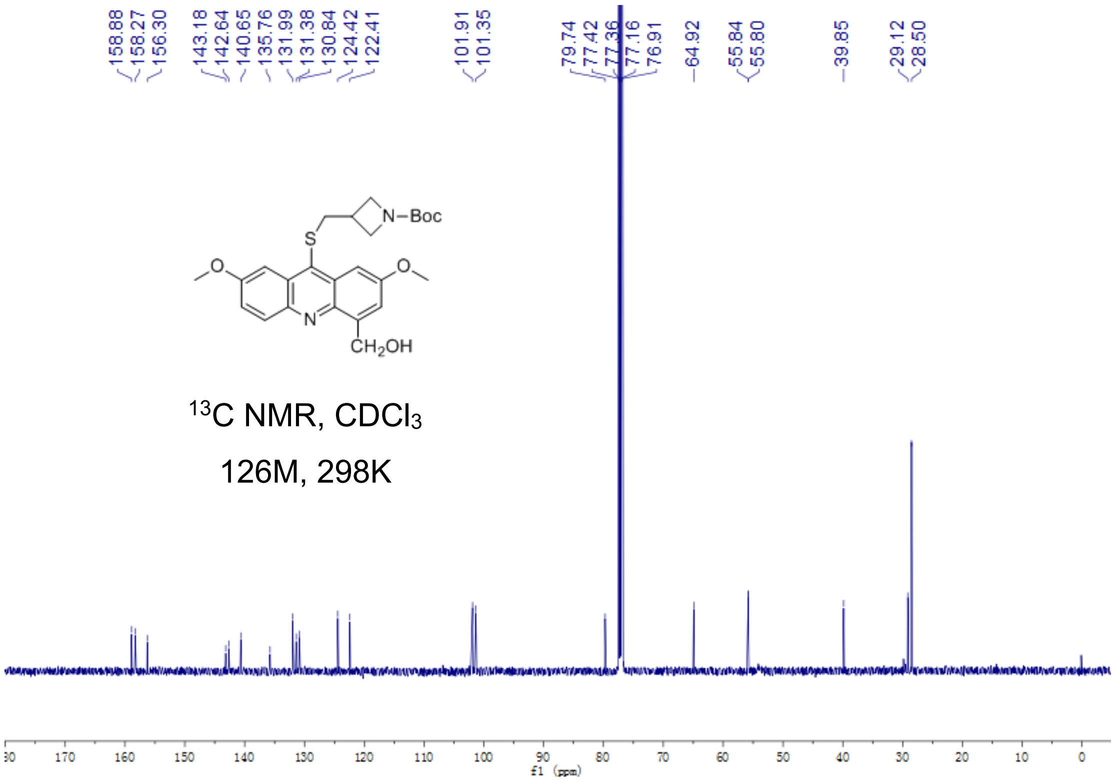


Fig S2.78. $^{13}$C-NMR of **19b**

**19**

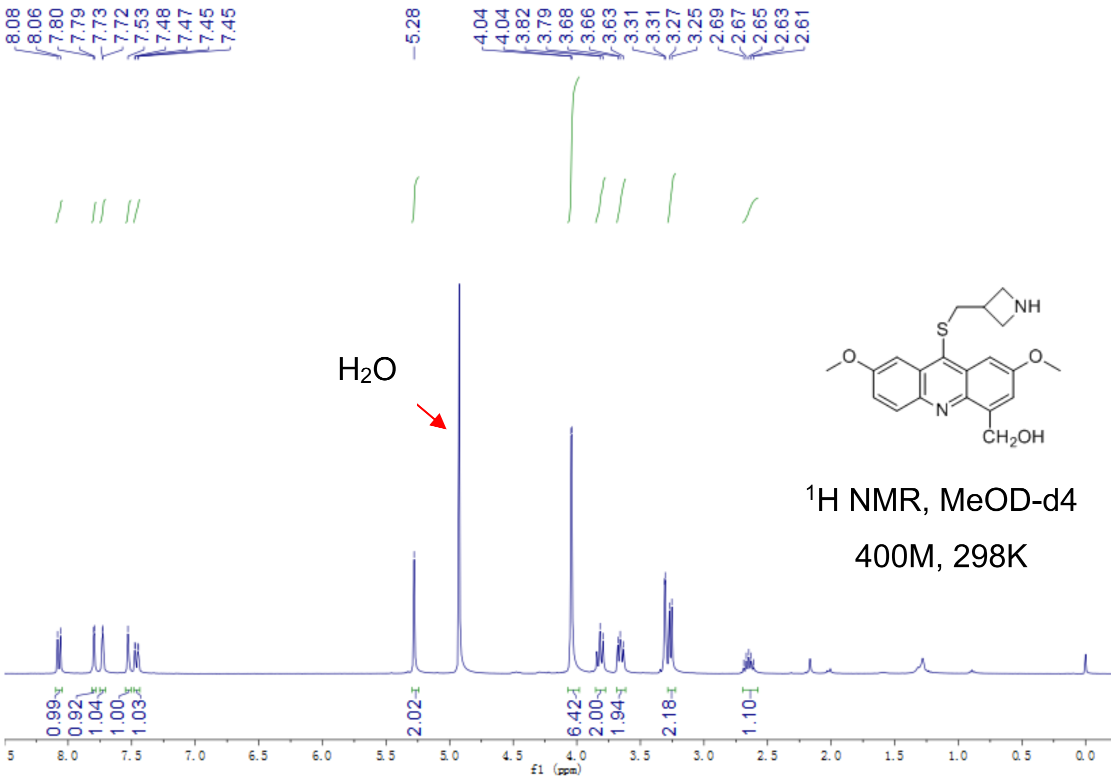


Fig S2.79. $^{1}$H-NMR of **19**

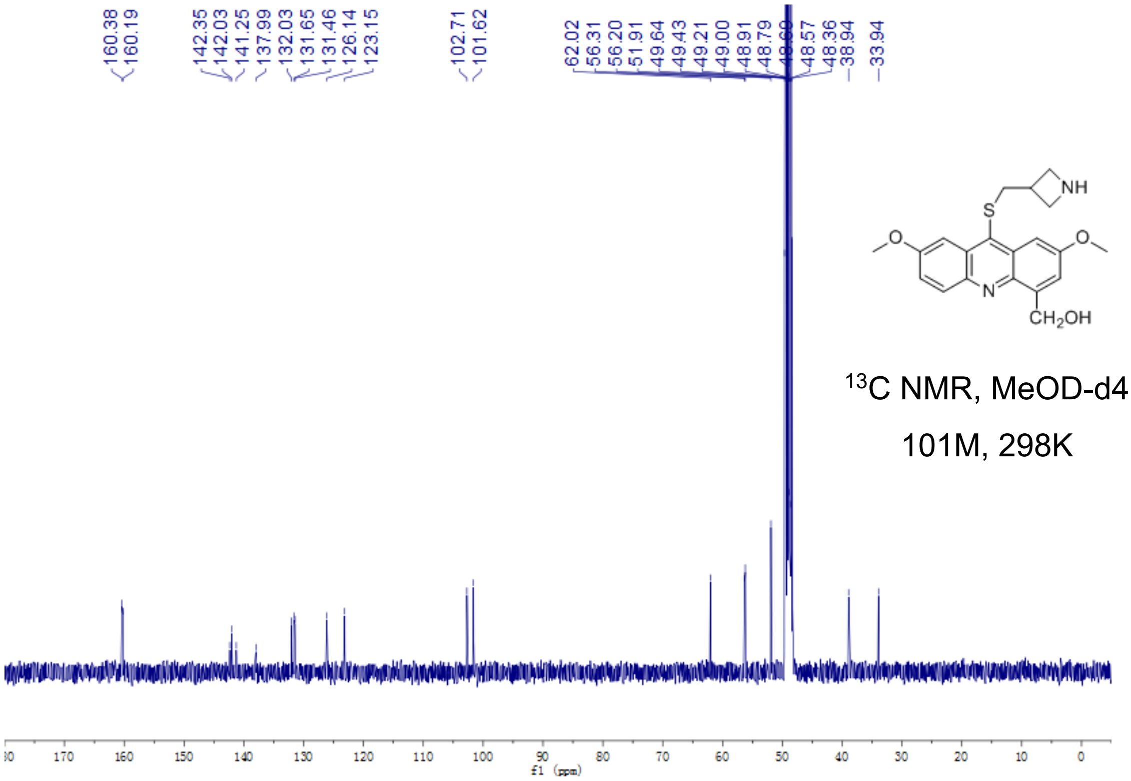


Fig S2.80. $^{13}$C-NMR of **19**

**20a**

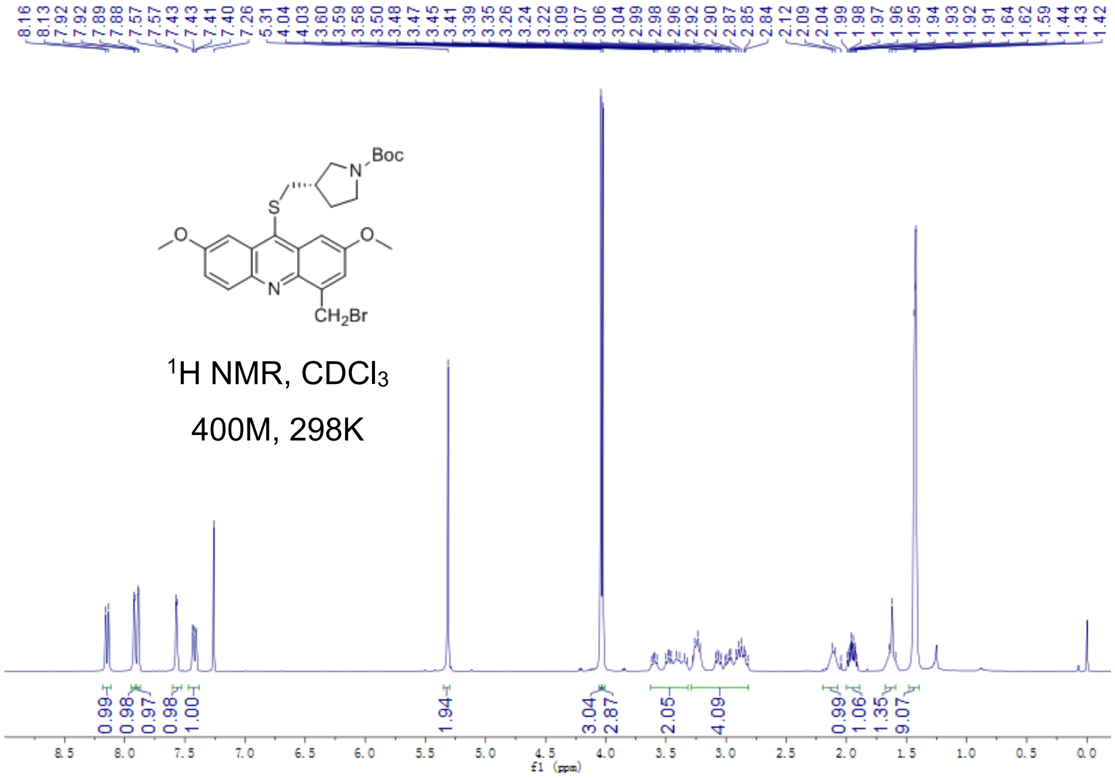

Fig S2.81. [1]H-NMR of **20a**

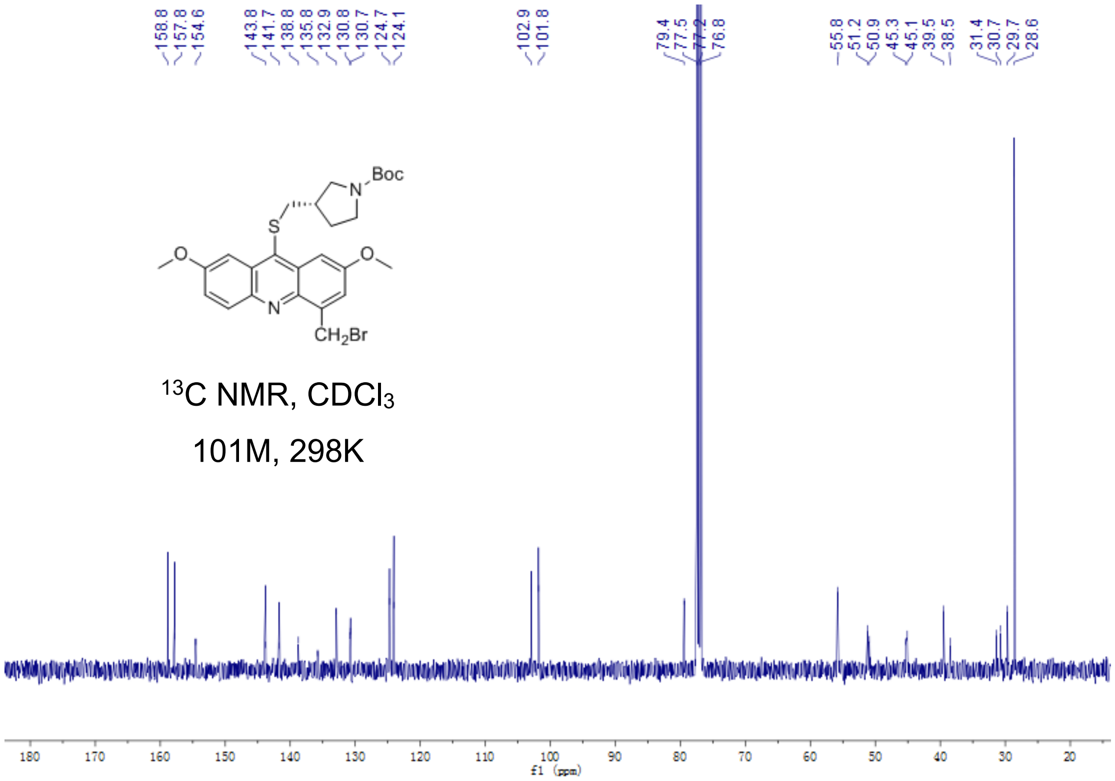


Fig S2.82. [13]C-NMR of **20a**

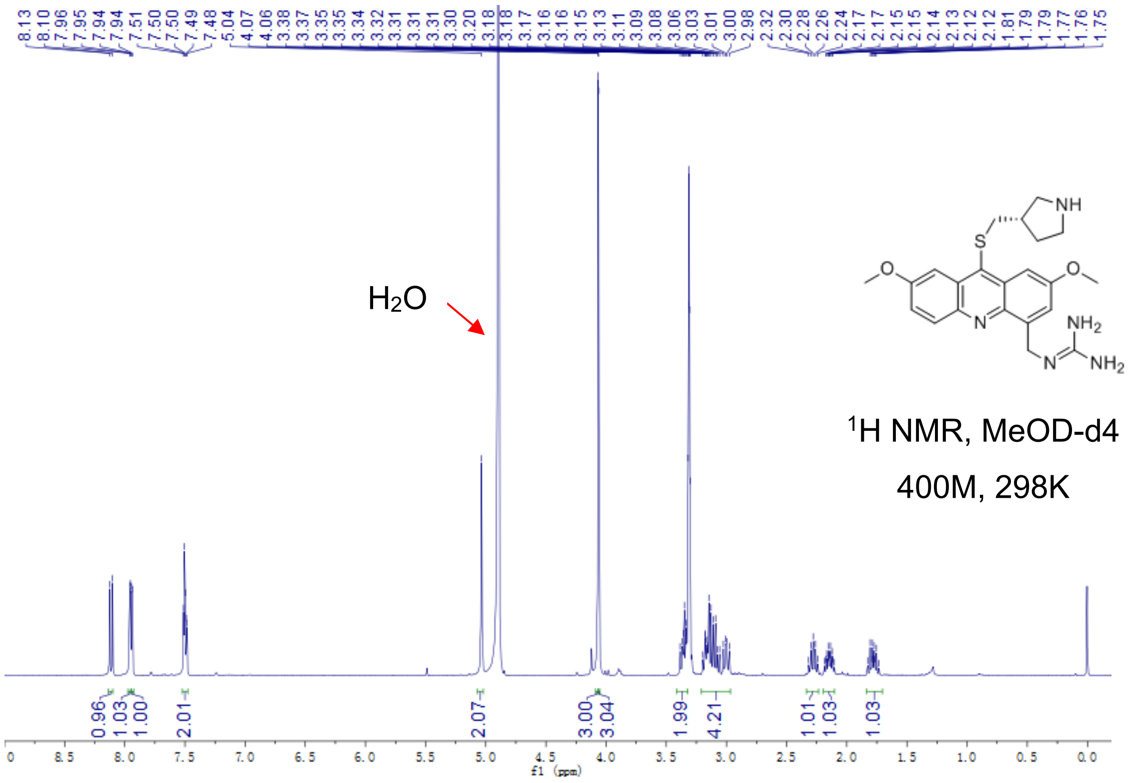


Fig S2.83. [1]H-NMR of **20**

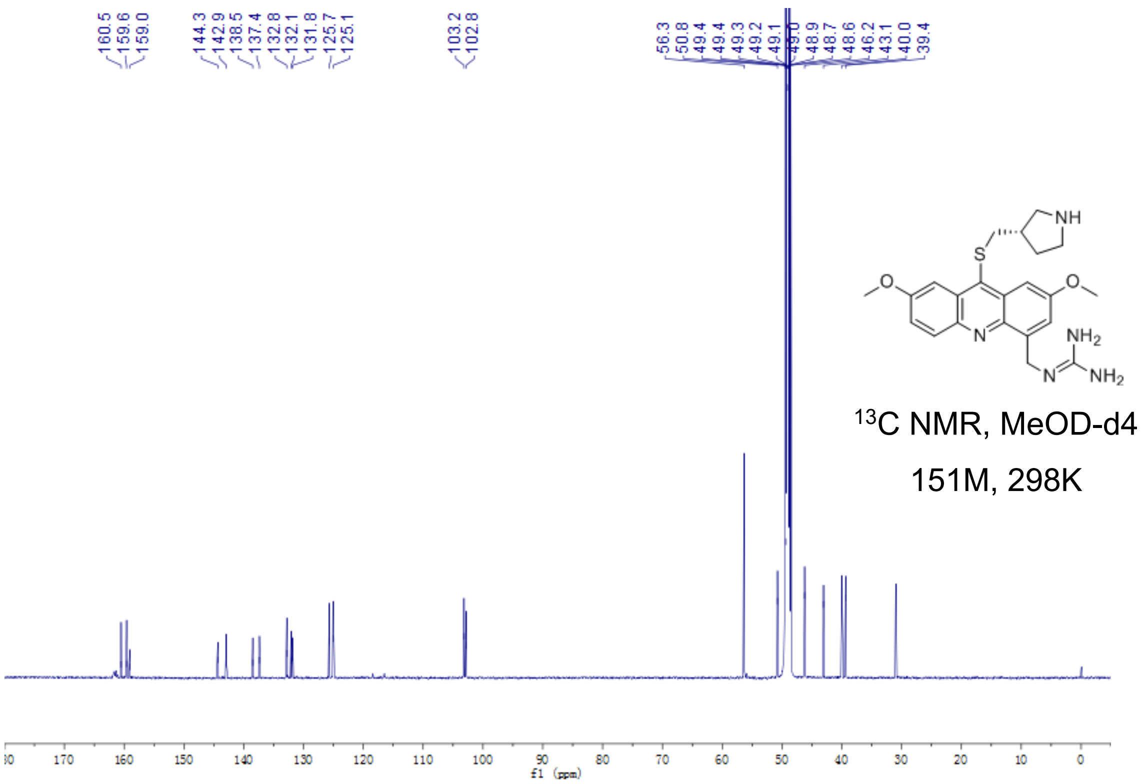


Fig S2.84. [13]C-NMR of **20**

**21a**

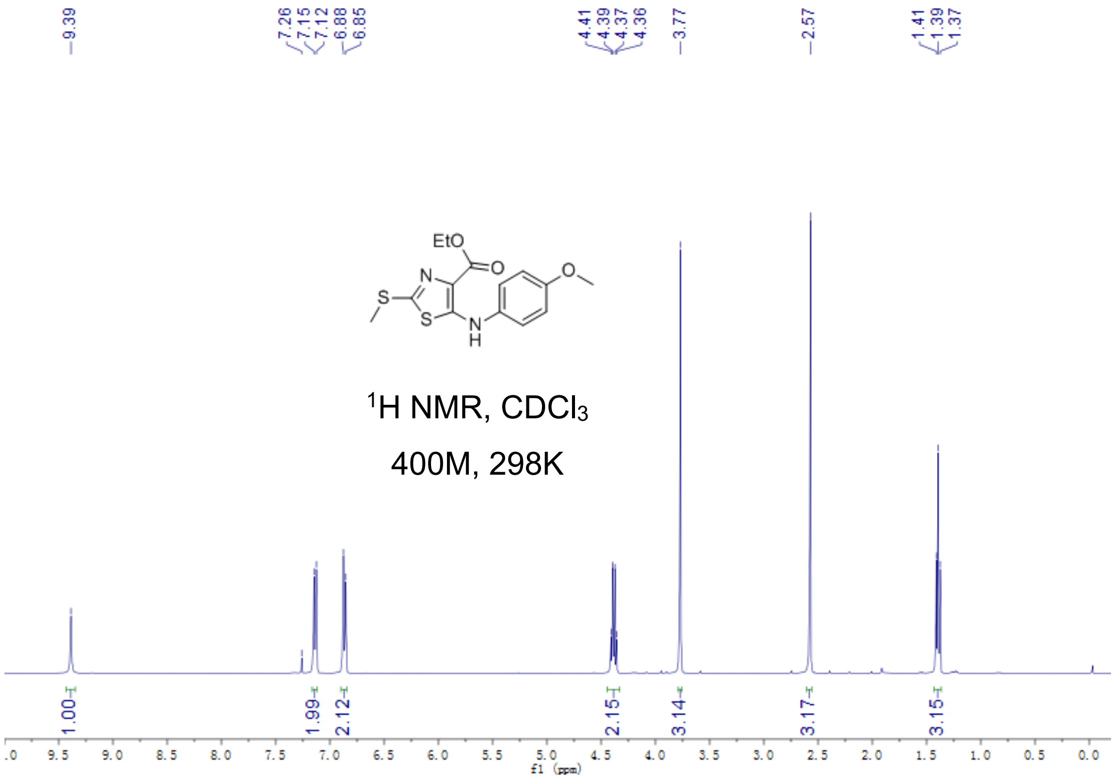


Fig S2.85. $^{1}$H-NMR of **21a**

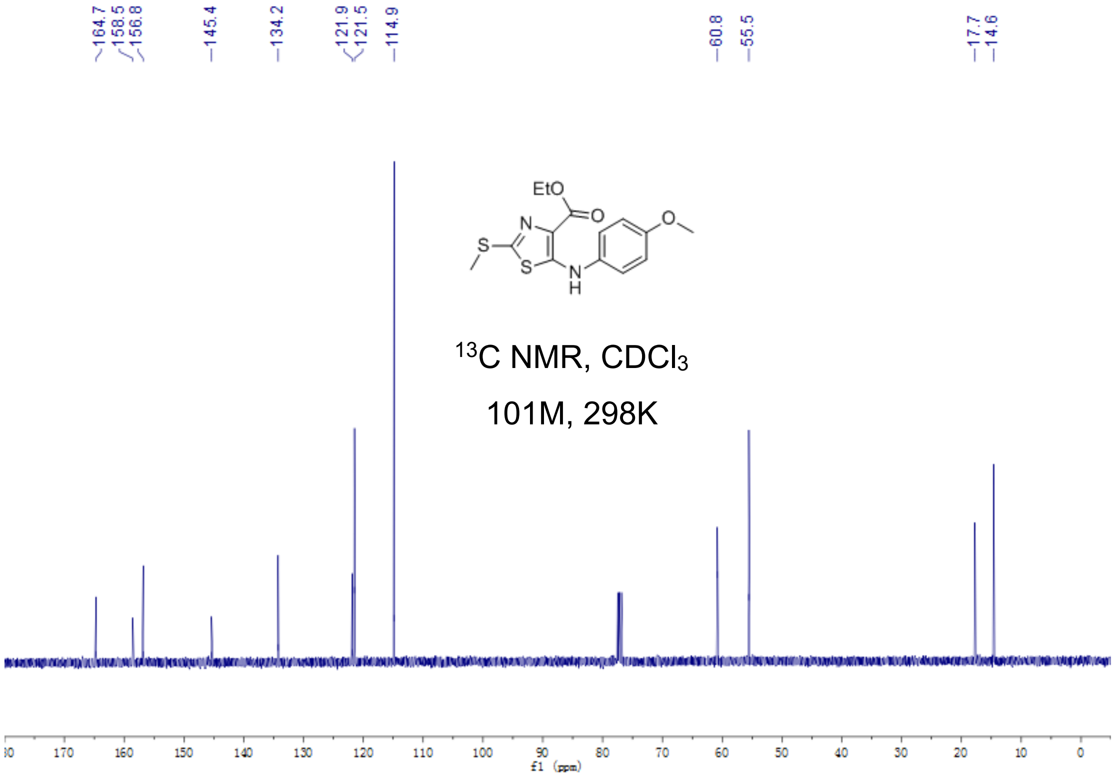


Fig S2.86. $^{13}$C-NMR of **21a**

**21b**

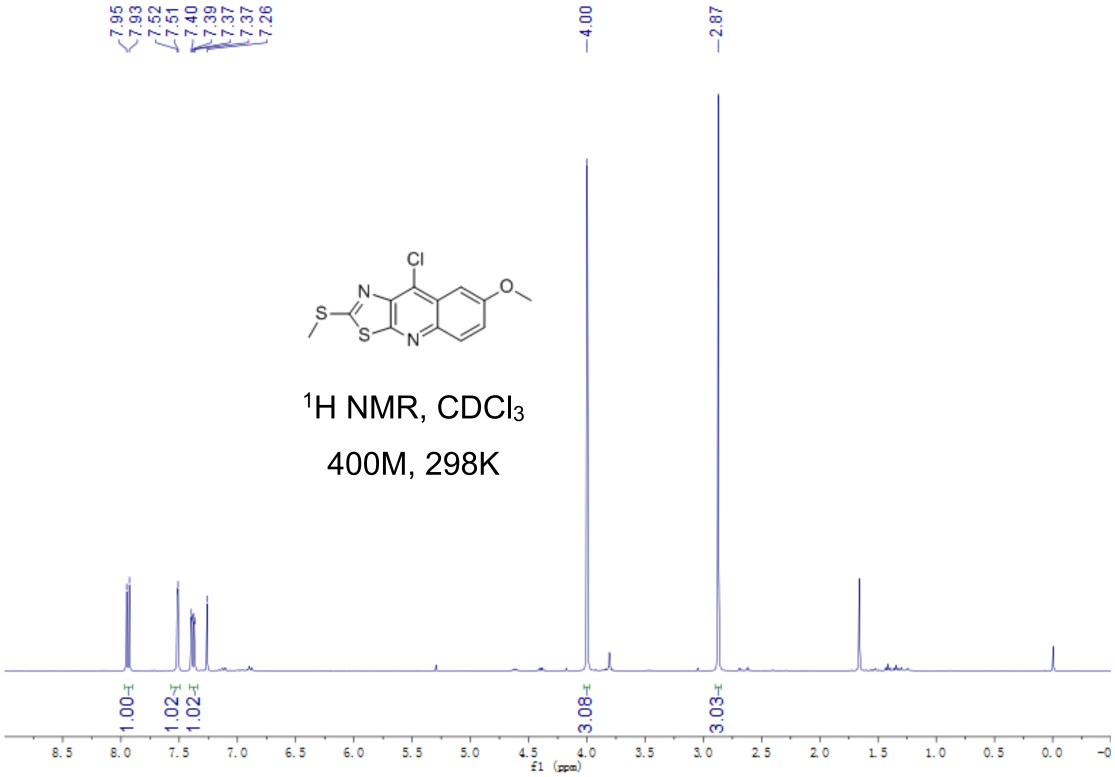


Fig S2.87. $^{1}$H-NMR of **21b**

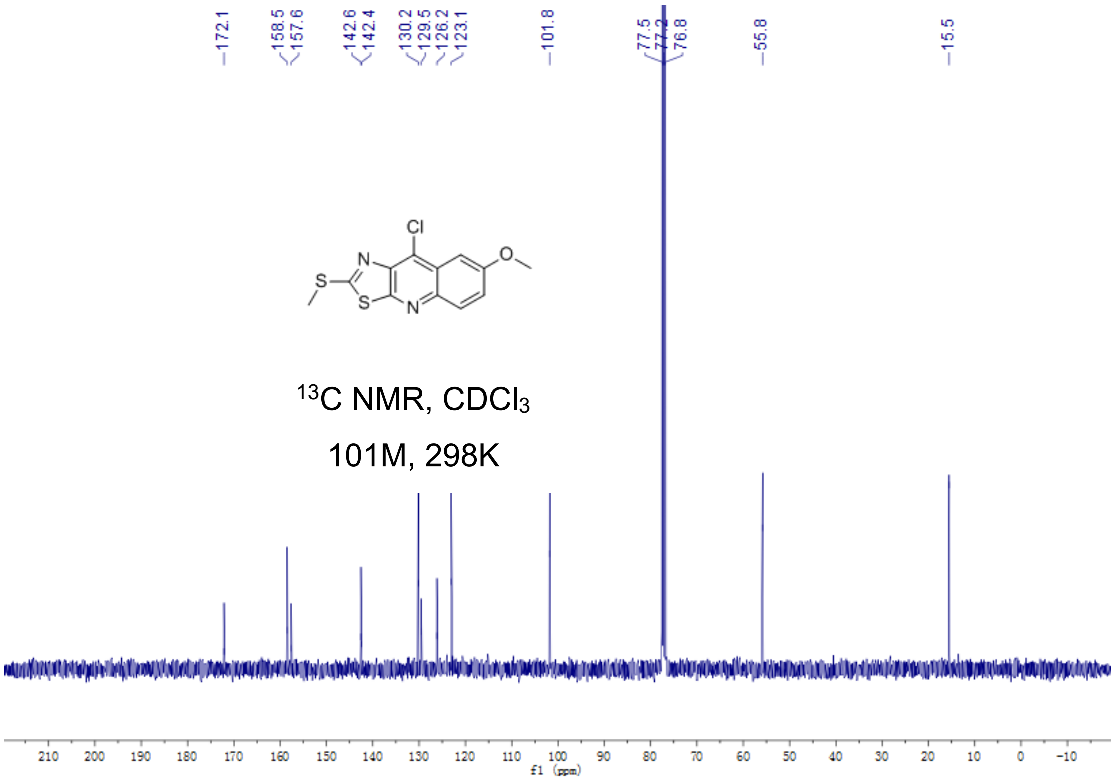


Fig S2.88. $^{13}$C-NMR of **21b**

**21c**

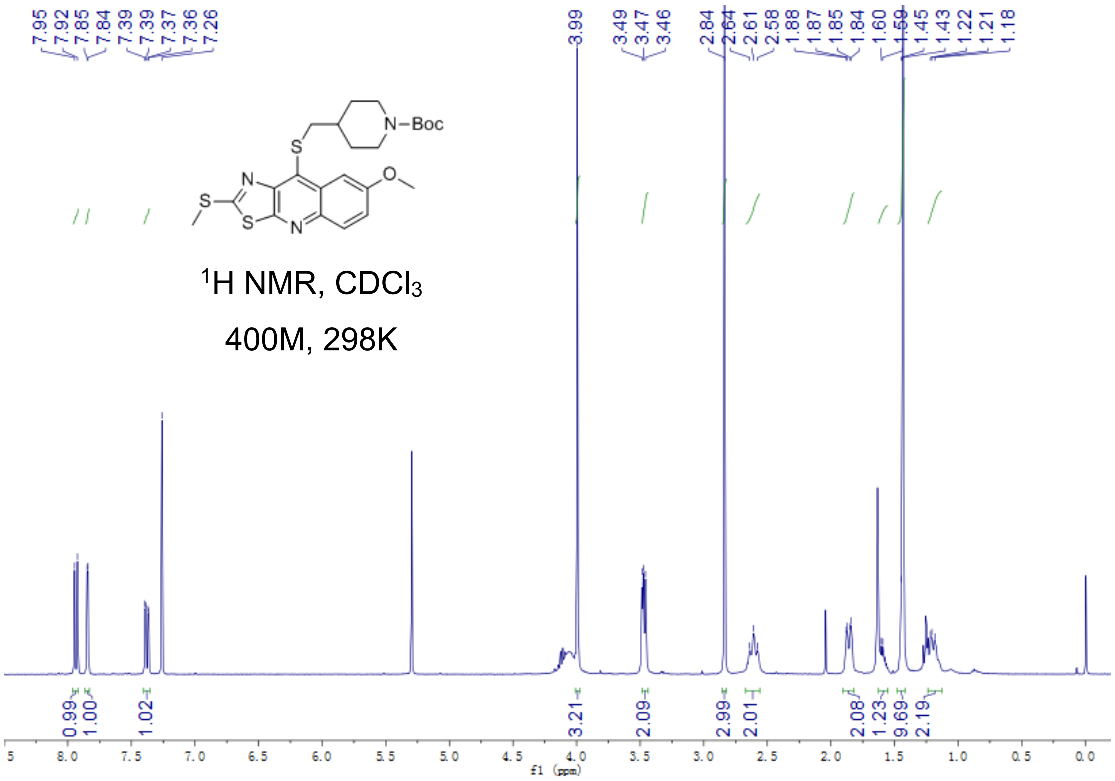


Fig S2.89. $^1$H-NMR of **21c**

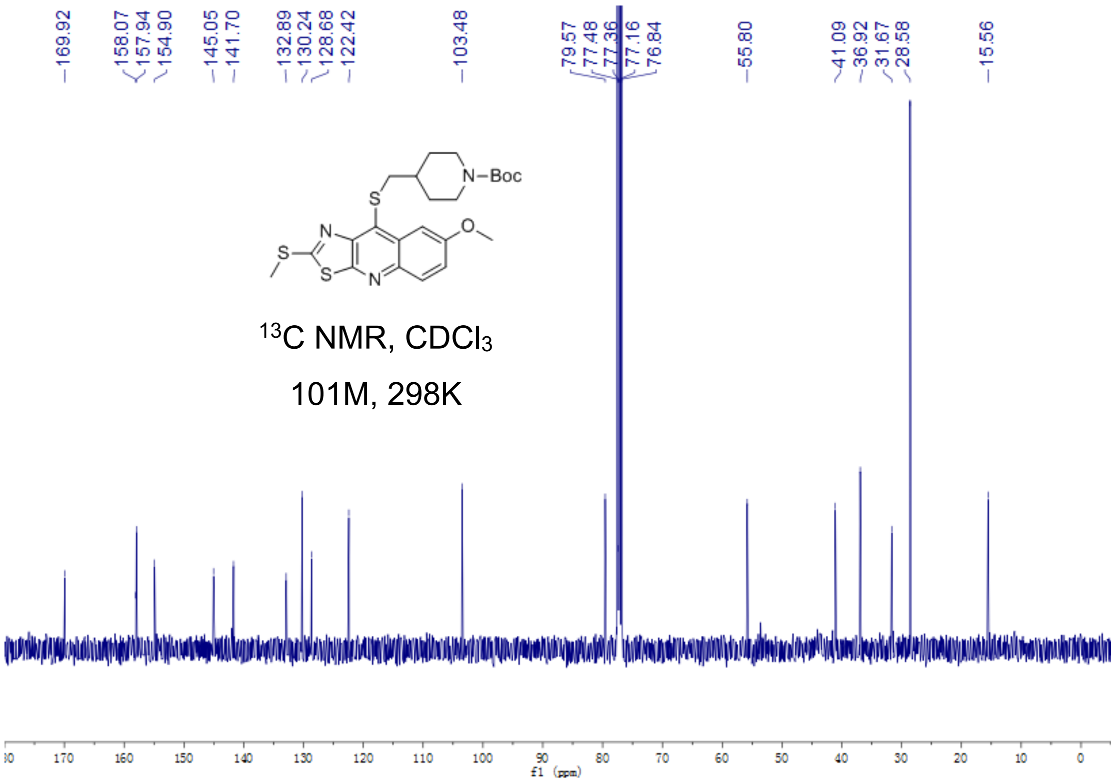

Fig S2.90. $^{13}$C-NMR of **21c**

**21**

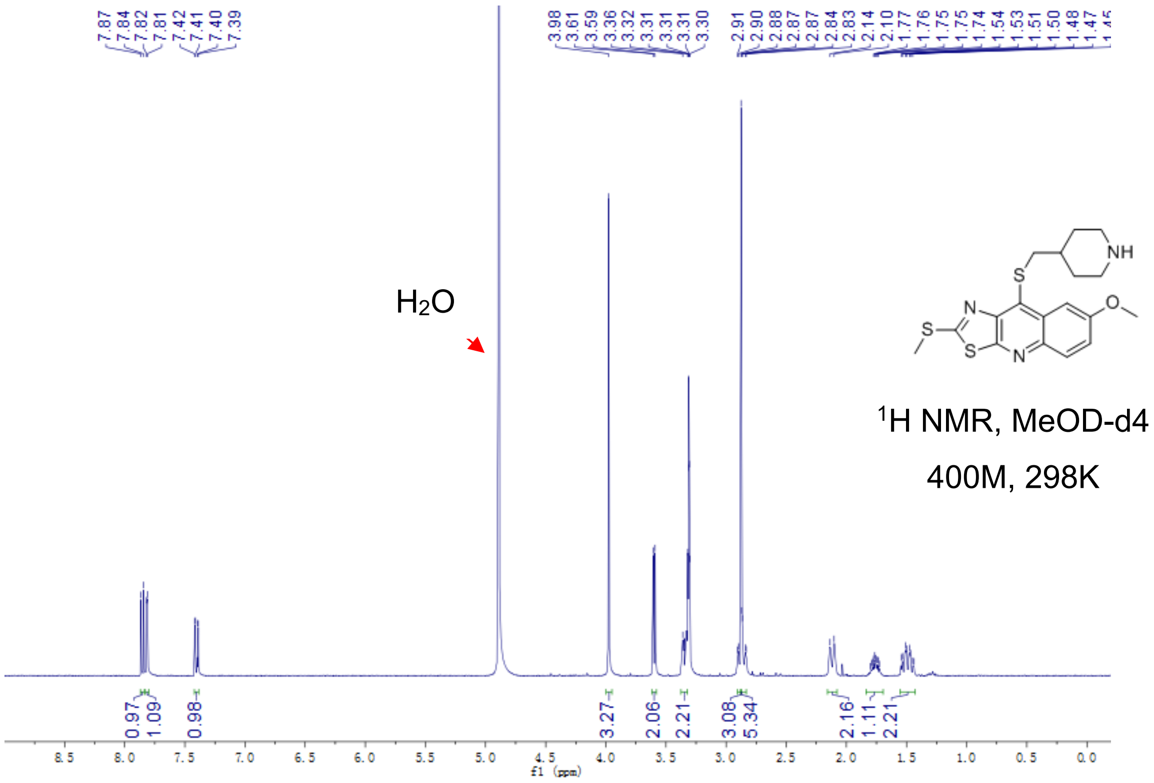


Fig S2.91. $^{1}$H-NMR of **21**

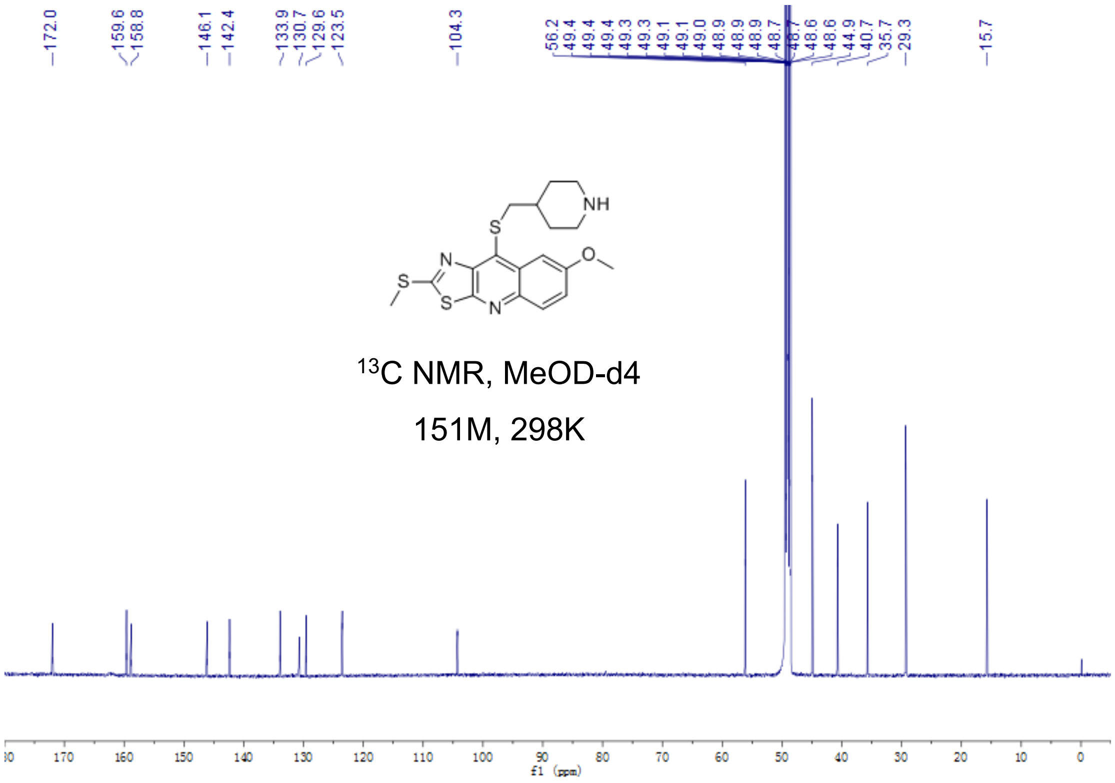


Fig S2.92. $^{13}$C-NMR of **21**

**22a**

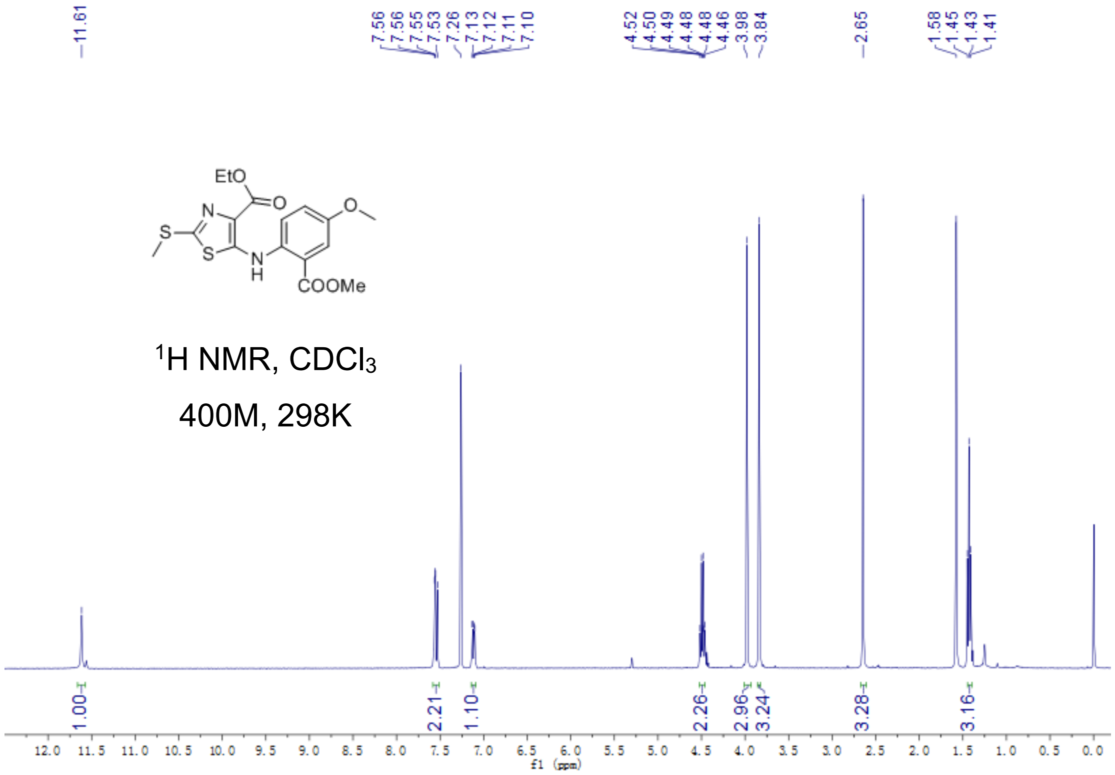


Fig S2.93. $^{1}$H-NMR of **22a**

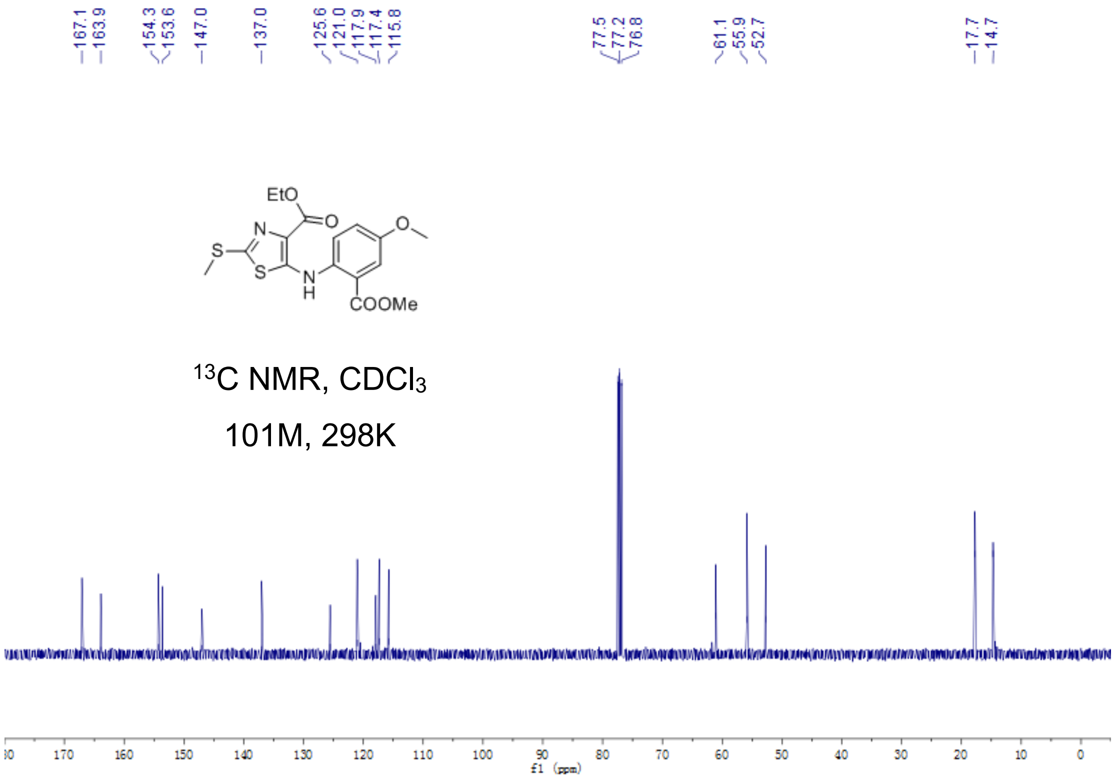


Fig S2.94. $^{13}$C-NMR of **22a**

**22b**

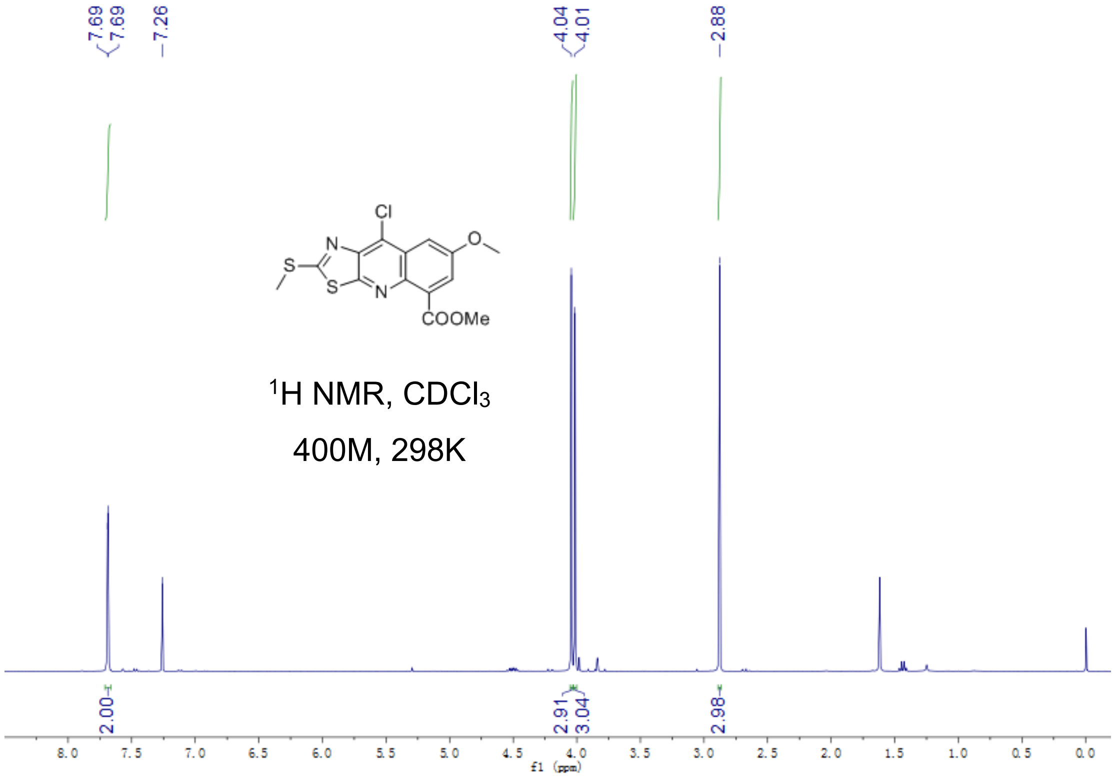

Fig S2.95. $^{1}$H-NMR of **22b**

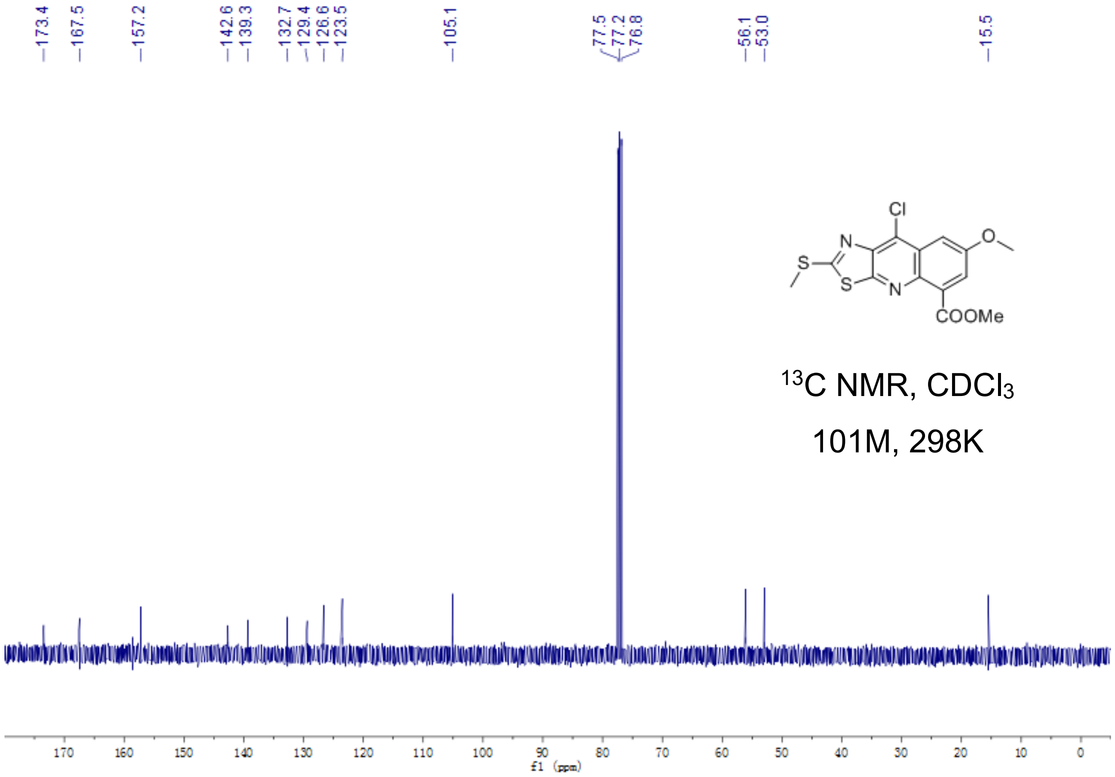


Fig S2.96. $^{13}$C-NMR of **22b**

**22c**

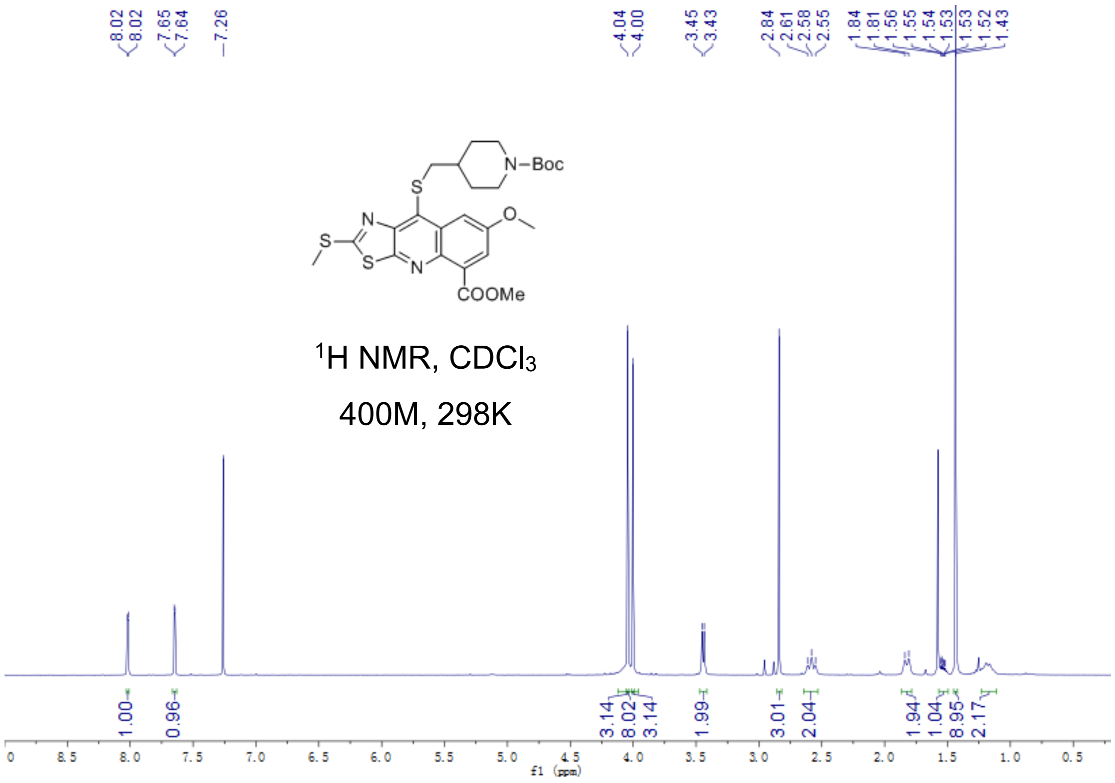


Fig S2.97. $^1$H-NMR of **22c**

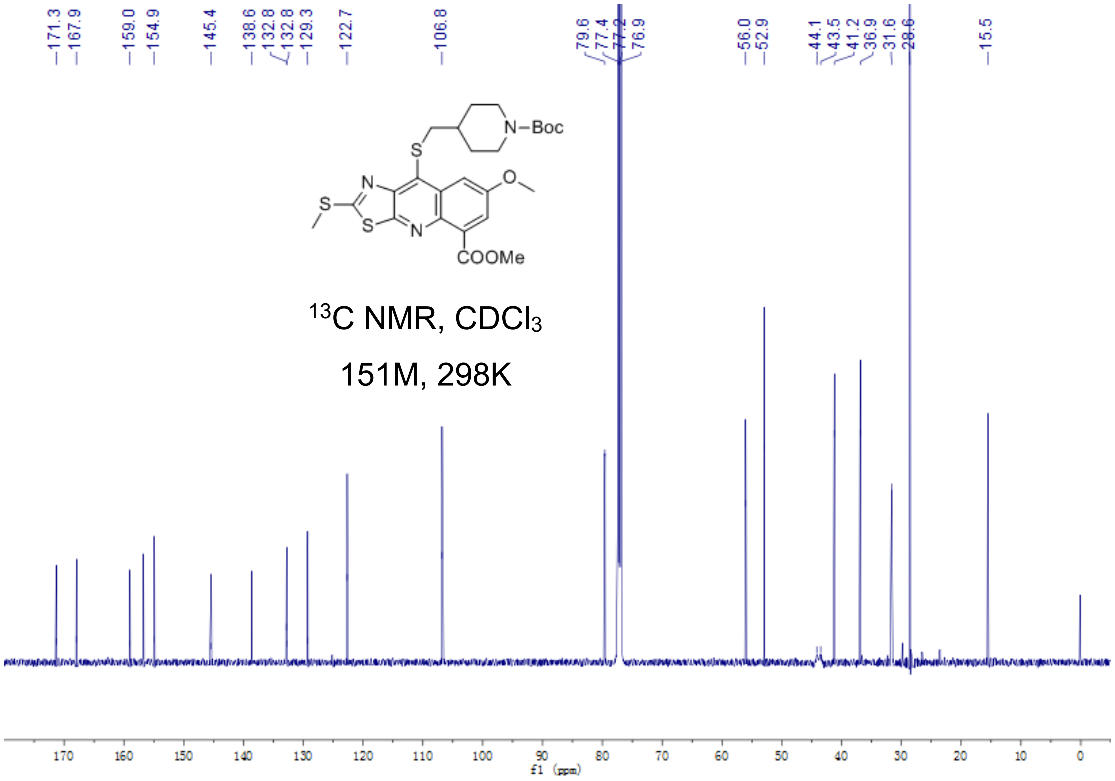


Fig S2.98. $^{13}$C-NMR of **22c**

**22**

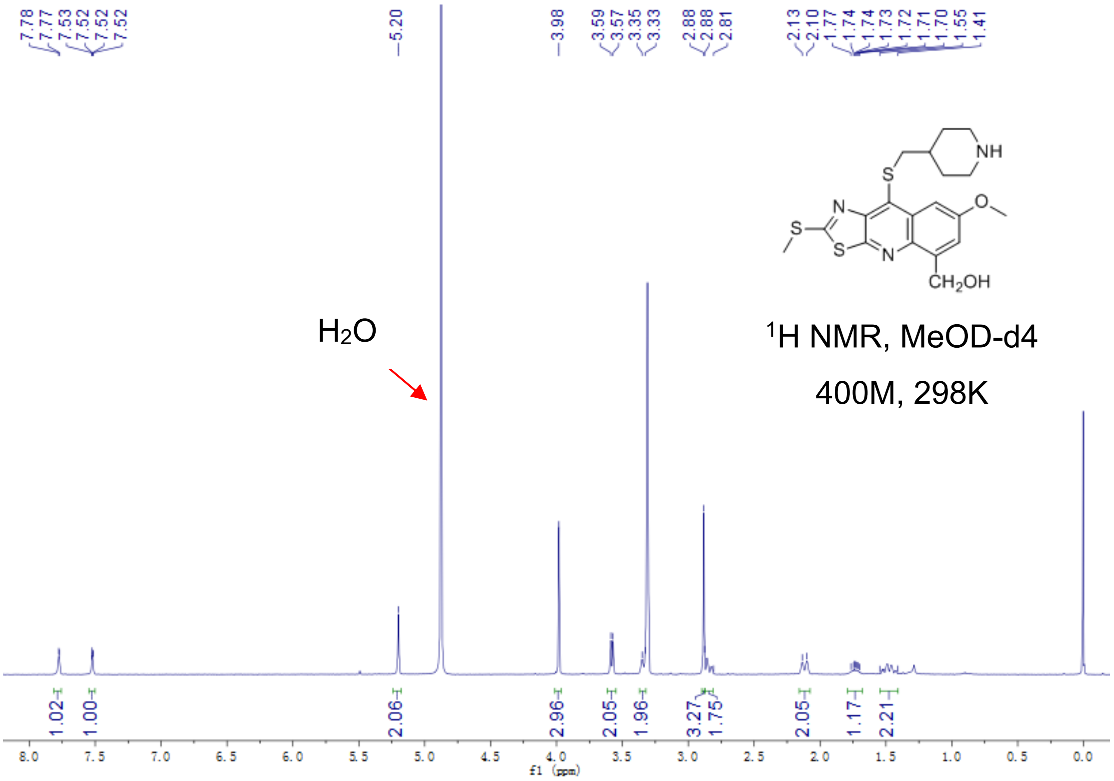


Fig S2.99. [1]H-NMR of **22**

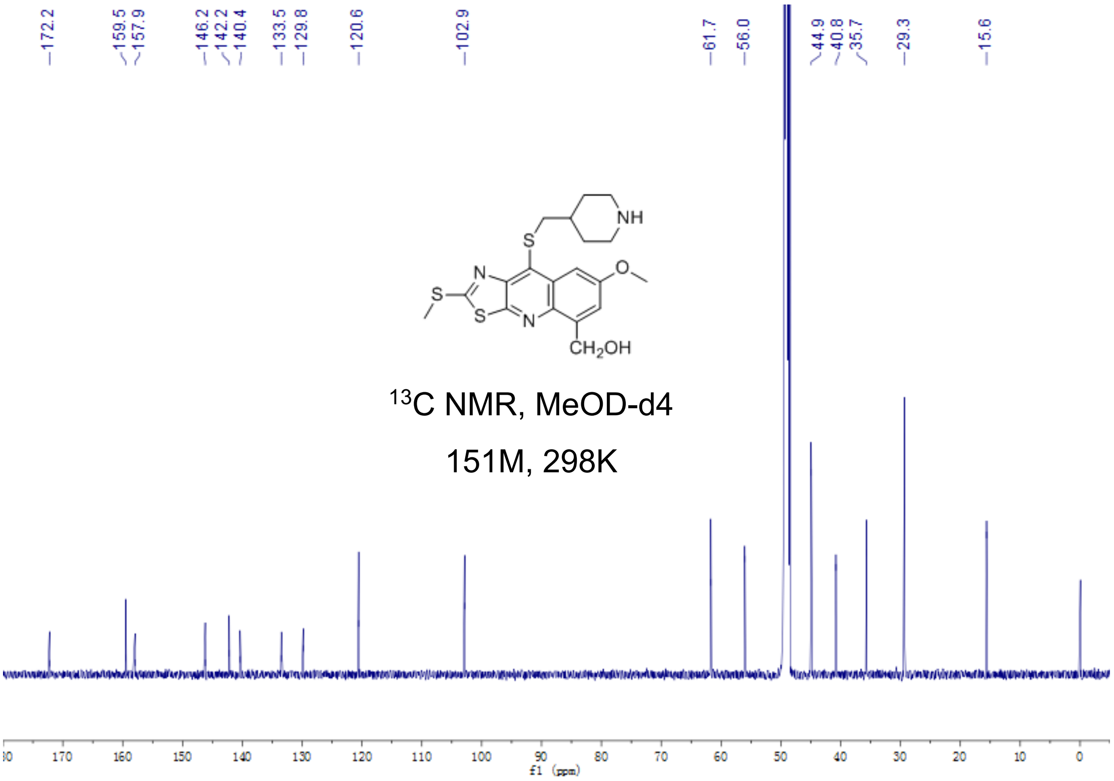


Fig S2.100. [13]C-NMR of **22**
